# Lipidomic signatures in microglial extracellular vesicles during acute inflammation: a gateway to neurological biomarkers

**DOI:** 10.1101/2025.05.27.656354

**Authors:** Nikita Ollen-Bittle, Wenxuan Wang, Kimberly Molina Bean, Shuang Zhao, Adriana Zardini Buzatto, Yifei Dong, Liang Li, Shawn N. Whitehead

**Affiliations:** Department of Anatomy and Cell Biology, Schulich School of Medicine and Dentistry, Western University, London, Ontario, N6A 5C1; Department of Chemistry, University of Alberta, Edmonton, Alberta, T6G 2G2, Canada; The Metabolomics Innovation Centre (TMIC), University of Alberta, Edmonton, Alberta, T6G 2G2, Canada; Department of Biochemistry, Microbiology & Immunology, University of Saskatchewan, Saskatoon, Saskatchewan, Canada

**Keywords:** Lipids, Biomarkers, LC-MS/MS, Microglia, Extracellular Vesicles, Lipidomics, neurological disease

## Abstract

Extracellular vesicles (EVs) are membrane bound vesicles released from all cells throughout the body, including the central nervous system, and are known to carry both membrane-bound proteins and cargo reflective of their cell of origin. EVs show promise as neurological disease biomarkers due to their molecular makeup reflecting their parent-cell composition signature and due to their ability to cross the blood-brain barrier. To-date, the vast majority of research in this field has explored the protein profiles of EVs; however, lipids play an important role not only in the formation of EVs, but also in mediating cellular function and the pathological progression of many neurodegenerative conditions. Herein, we take a critical first step in determining the potential utility of EV lipids as biomarkers in neurological disease. *In vitro* we exposed BV-2 microglia to either control media or media containing lipopolysaccharides (LPS), a known pro-inflammatory stimulus, for 24 hours then isolated both the cells and their EVs and performed LC-MS/MS. For the first time, we reveal distinct lipidomic changes can differentiate resting vs. pro-inflammatory microglia and their EVs, while distinct lipids are preserved between EVs and their parent cell. Moreover, we add to current literature by demonstrating acute pro-inflammatory activation of microglia results in the activation and suppression of distinct lipidomic pathways. Finally, we demonstrate that analysis of lipid-based relationships between parent cells and their EVs may be a useful tool to infer cellular function. This study is the first of its kind to demonstrate that lipidomic analysis can not only differentiate the functional state of cells *in vitro* but can also differentiate their EVs. We lay the first brick in a foundation to support future research into EV lipids as novel and exciting biomarker candidates in neurological disease.

## Introduction

In addition to being a significant cause of both mortality and morbidity, neurological disease is one of the greatest strains on healthcare systems (GBD 2017 US Neurological Disorders Collaborators 2021). Due to the complexity of the nervous system, non-specific symptoms and variable presentations, neurological diseases are considered among the most difficult medical conditions to diagnose. Considering acute pathology, there are cases such as hemorrhagic vs ischemic stroke where conditions present almost indistinguishably clinically yet have opposing treatments that may worsen the alternative diagnosis (Kase *et al*. 2001). In terms of chronic neurodegenerative diseases, there are cases where early intervention can significantly improve patient outcome such as early stem cell replacement therapy in specific leukodystrophies (Dulski *et al*. 2022) or early commencement of cognitive enhancement therapeutics in Alzheimer’s disease (AD) (Dubois *et al*. 2016; Vrahatis *et al*. 2023). Thus, the ability to diagnose neurological diseases in their early phases is critical for identifying optimal therapeutic windows and effective treatments (Rasmussen and and Langerman 2019; Miller 2004; Hansson 2021). Research into biomarkers allowing for the early diagnosis of neurological conditions is continuously emerging. In AD a decrease in amyloid beta (Aβ) 42 has been shown to occur in cerebrospinal fluid (CSF) even prior to abnormal PET imaging (Palmqvist *et al*. 2016; Olsson *et al*. 2016) and a decrease in Aβ42/ Aβ40 within CSF shows promise as a potential diagnostic biomarker (Hansson *et al*. 2019). In the case of Parkinson’s disease, Multiple Systems Atrophy and Dementia with Lewy Bodies α-synuclein decreases in CSF (Hansson 2021; Parnetti *et al*. 2019). While these protein-based biomarkers show early promise, future work aims to utilize biomarker panels, in which additional biomarkers may help improve specificity. Additionally, since lumbar puncture is invasive and not readily accessible outside of specialist centers, there is a need for blood-based biomarkers. Collectively, there is a dire and urgent need for the development of biomarkers to aid in the early diagnosis of neurological diseases. Microglia are the predominant resident immune cell found in the parenchyma of the central nervous system (CNS) and are critical regulators of homeostasis and first responders to CNS injury. Microglia are known to play important roles in neuroprotective processes, but their dysregulation has also been associated with the progression of both acute CNS injury and neurodegenerative disease (Hickman *et al*. 2018). These cells display remarkable heterogeneity to various stimuli, rapidly transitioning between different phenotypes in response to environmental cues and pathological conditions (Tan *et al*. 2020). Their unique ability to sense and respond to subtle changes in brain microenvironment makes them particularly valuable for early detection of neurological dysfunction. Recent single-cell transcriptomic studies have revealed distinct microglial subpopulations with disease-specific signatures, further emphasizing their potential as diagnostic indicators (Olah *et al*. 2020). This makes microglia important targets for both understanding neurological diseases and for therapeutic modulation. In fact, in the last decade genome-wide association studies (GWAS) have identified genetic variants expressed in microglia that increase risk for late onset AD (Romero-Molina *et al*. 2022). There has also been the emergence of microglia modulating therapeutics that are continuing to be explored in a therapeutic context including TREM2 and PPAR-γ agonists (Lewcock *et al*. 2020; Carta and Pisanu 2013). Collectively, since microglia are critical for the maintenance of homeostasis, can respond to but also exacerbate CNS injury and can potentially be modulated therapeutically, there is an increasing effort to discover blood-based microglia specific biomarkers as a peripheral window into neurological disease (Zhang *et al*. 2021; Chen *et al*. 2023; Tirandi *et al*. 2023).

Lipids are dynamic organic molecules that make up more than half of the brains dry weight. They are essential components of cell membranes and axonal sheaths and take part in a vast range of requisite cellular processes in the brain including cell differentiation, energy storage, metabolism and cell signaling (Ioannou *et al*. 2019; Bazan 2005; Montani 2021; Bansal *et al*. 1999). Lipid dysregulation is critically associated with several neurological diseases including AD, Parkinson’s disease, multiple sclerosis (MS) and amyotrophic lateral sclerosis (Phan *et al*. 2023; Wei *et al*. 2024; Podbielska *et al*. 2021). Given the need for biomarkers and the central role lipids play in CNS function, lipids make an intriguing candidate biomarker for many neurological diseases.

Extracellular vesicles (EVs) are small lipid membrane bound vesicles secreted by cells throughout the body and brain into extracellular fluids (Doyle and Wang 2019; Yáñez-Mó *et al*. 2015). EVs were previously thought to be a mechanism of cellular waste removal; however, it is now known they are also important modulators of cell-to-cell signalling (Doyle and Wang 2019; Yáñez-Mó *et al*. 2015). EVs are formed either through the outward budding of the cell membrane (microvesicles) or through the inward budding of the early endosomes as they mature into multivesicular bodies. These multivesicular bodies are then either targeted for destruction by the lysosome or for fusion with the cellular membrane releasing their content of small EVs (exosomes) (Doyle and Wang 2019). Due to their mechanisms of formation, EVs contain membrane, membrane bound proteins and cargo from their parent cells (Doyle and Wang 2019; Yáñez-Mó *et al*. 2015; Raposo and Stoorvogel 2013). Moreover, due to their small size and lipophilic nature, EVs are able to freely cross the blood brain barrier (BBB) (Perets *et al*. 2020; Ramos-Zaldívar *et al*. 2022). The critical role of EVs in cellular communication, their composition being reflective of their parent cells and their ability to cross the BBB have caused EVs to quickly emerge as potential cell-specific biomarkers in neurological disease (Ollen-Bittle *et al*. 2022; Ollen-Bittle *et al*. 2024b; Thompson *et al*. 2016). Most of the research on EV-based biomarkers has focused on protein composition; however, lipids are known to be important players in EV-mediated cell-to-cell communication, cargo sorting and even functional effects on target cells (Fyfe *et al*. 2023). New research on the potential of EV lipids as biomarkers is continuously emerging with recent reports of EV lipids showing promise as biomarkers for both breast (Liu *et al*. 2023) and prostate cancer(Brzozowski *et al*. 2018). Additionally, a recent study has shown dysregulation of lipids in EVs derived from post-mortem AD brains (Su *et al*. 2021). However, little is known about the baseline lipidomic profile of cell-specific EVs, how the EV lipidome relates to their cell of origin, or how/if it changes with changes in cell state. This is all requisite information if EV lipids are to be considered potential blood-based biomarkers for neurological disease and proof of principle work is needed in a controlled *in vitro* setting before moving to more complex experimental models and human samples.

Here, we interrogate the lipidomic profile of microglial EVs in a controlled, *in vitro* setting to establish foundational information regarding the potential future utility of EV lipids as biomarkers in neurological disease. We exposed BV-2 microglia to lipopolysaccharide (LPS), a component of gram-negative bacteria and well-established pro-inflammatory activator of microglia (Nakamura *et al*. 1999) and conducted lipidomic analysis of both the microglia and their secreted EVs. We hypothesized the lipidomic profile of EVs would correlate with the lipidomic profile of the cell group from which they were released, and that the lipidomic profile of EVs would therefore be capable of inferring the cellular activation state.

## Methods

To enhance reproducibility a Lipidomics Minimal Reporting Checklist (Kopczynski *et al*. 2024) from the Lipidomics Standards Initiative can be found in Supplemental Table 8.

### Cell growth conditions

BV-2 cells were purchased from AcceGen (catalogue number: ABC-TC212S). Cells were passaged a maximum of nine times. BV-2 cells were cultured in a humidified atmosphere at 37 °C with 5% CO_2_. Prior to treatment cells were maintained with dulbecco’s Modified Eagle Medium (DMEM) (Gibco) supplemented with 10% EV-depleted fetal bovine serum (FBS), Penicillin (100 U/mL) and Streptomycin (100 μg/mL) (Gibco). EV-depleted FBS was ultra-centrifuged at 100,000xG, 4 °C overnight after which the supernatant was filtered through a 0.2 μm filter.

### Cell treatments

BV-2 microglia were seeded in 150 mm plates at a seeding density of 5.0 × 10^6^ cells per plate and grown to 70% confluency. Cells were then washed with PBS and maintained in serum free media for 12 hours prior to treatment. Cells were then either treated with 40 mL of fresh serum free media (non-treated (NT), n=5) or 40 mL of 500 ng/ml LPS (n=5) in serum free media for 24 hours. At 24 hours both cells and their media were collected, frozen and transported on dry ice for further analysis. Media was collected into 50 mL tubes and cells were scraped off plates and collected in 15 mL tubes. The workflow utilized in this study is summarized in **Figure 1**.

**Figure 1.**
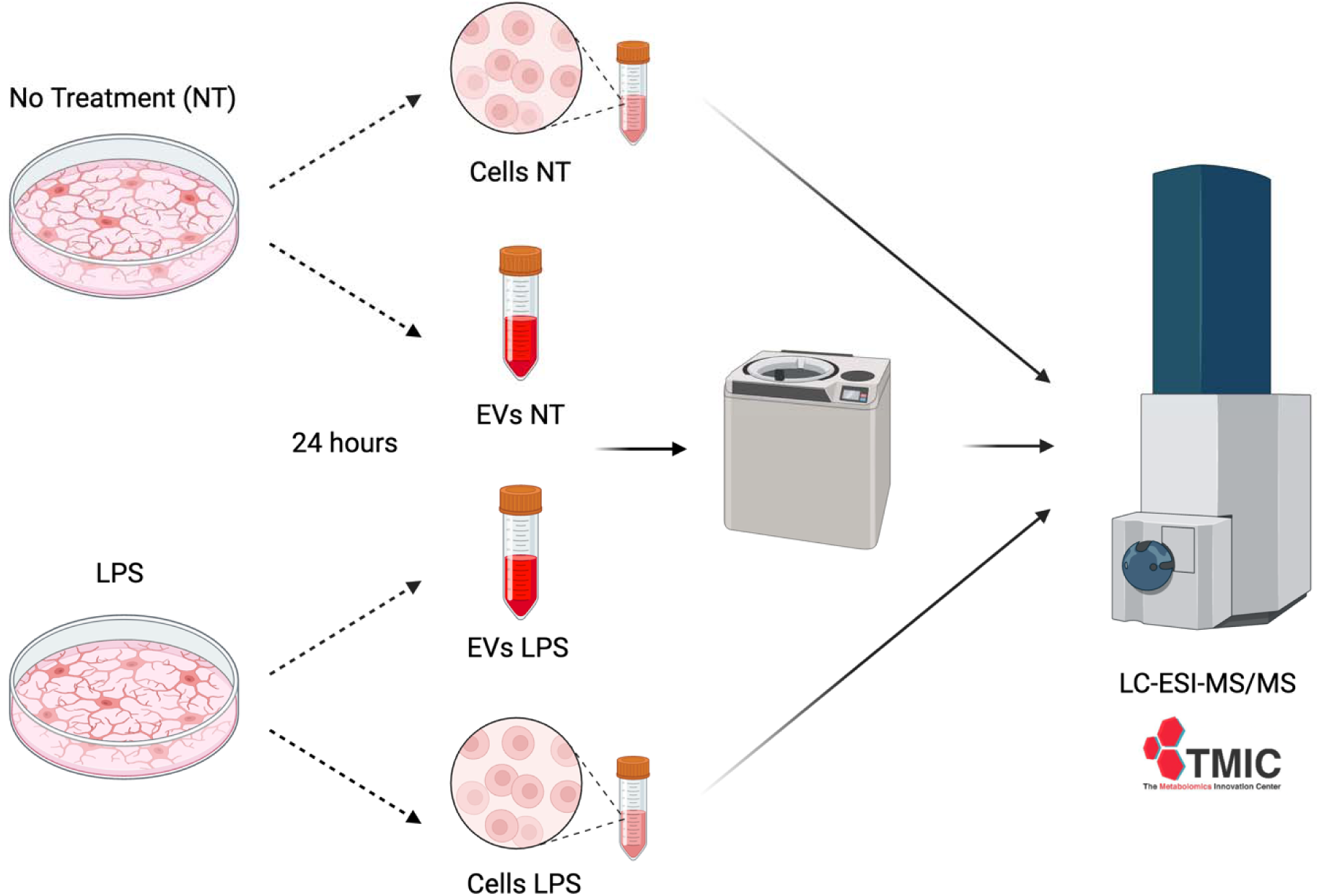
Workflow summary. BV-2 microglia were seeded in 150 mm plates and grown to 70% confluency. Cells were switched to serum free media for 12 hours then non-treated (NT) cells underwent fresh serum free media replacement (NT, n=5), while LPS exposed cells underwent media replacement with serum free media containing 500 ng/ml LPS (LPS, n=5). At 24 hours both cells and their media were collected and frozen. Once all samples were collected media was ultracentrifuged to isolate EVs and both cells and isolated EVs were sent to The Metabolomics Innovation Center (TMIC), Edmonton, Alberta, Canada for LC-MS/MS. Created with BioRender.com.

### EV isolation

EVs were isolated as previously described(Roseborough *et al*. 2023) and isolated EVs were further characterized (Supplemental Figure 1A). Briefly, 50 mL tubes with collected media were centrifuged at 500 RCF for 5 minutes to pellet cellular debris before supernatant was immediately transferred to a new 50 mL tube and placed on ice. Supernatant was then filtered through a 0.2 μm SFPA into Amicon ultra 15 100k centrifuge filter tubes, which were centrifuged to achieve a final concentrated volume of 1 mL supernatant. EVs were then pelleted through ultracentrifugation of the concentrated supernatant at 50,000 rpm (MLA130 rotor) for 1 hour 15 minutes at 4°C. The supernatant was then removed, and the pellet was immediately dried, covered and placed on dry ice for shipment to the LC-MS/MS facility (TMIC, The Metabolomics Innovation Center, Edmonton, Alberta, Canada).

### Data collection and processing by TMIC

Lipid isolation and LC-MS/MS data collection was performed in entirety at TMIC. Samples were analyzed randomized prior to analysis, and technicians at TMIC were blinded to sample group identities and other background information to reduce bias

### Lipid extraction

Lipid extraction was performed based on a modified Folch liquid-liquid extraction protocol in cold conditions. Briefly, the samples were homogenized in 80% ice-cold methanol with ceramic beads (3.2 mm) at 4.6 m/s for 15 s. Samples were aliquoted to 20 nmol of TMC and then mixed with NovaMT LipidRep Internal Standard Basic Mix for Tissue/Cells (Catalog No. IDG-7002-IS). Samples were re-suspended in cold water and sonicated for 5 min. Phase separation was then performed by adding chilled dichloromethane and methanol to achieve a final solvent ratio of 2:1:0.7 (DCM:MeOH:water). The samples were vortexed and equilibrated at 4°C room temperature for 10 min and centrifuged at 16,000 g for 10 min at 4°C. The aqueous and organic layers were collected separately, evaporated to dryness under nitrogen, and stored at –80 °C for subsequent analyses.

To evaluate technical stability, an aggregate of Quality Control (QC) mixture was prepared by pooling aliquots for each the aqueous and organic layers from each sample, then splitting into multiple equal-volume aliquots. These aliquots were dried under nitrogen flow and stored at –80 °C until use. For each randomized batch of samples (two batches of seven samples and one batch of six samples), one QC aliquot was re-suspended and duplicated injected, at the beginning and at the end of the batch sequence to evaluate batch effects. Additionally, multiple QC aliquots were injected both before and after all sample runs.

### LC-MS Analysis

Dried extracts were re-suspended in mobile phase B, vortexed for 1 min, and diluted with mobile phase A. The LC-MS/MS was performed in both positive and negative ionization mode on a Thermo Vanquish UHPLC coupled to Bruker Impact II QTOF Mass Spectrometer. The system included a Waters Acquity CSH C18 column 1.7 μm, with a flow rate of 250 to 200 μL/min and a column oven temperature of 45°C. LC analysis was conducted using 10 mM ammonium formate in methanol/acetonitrile/water (50:40:10 v/v/v) as mobile phase A, and 10 mM ammonium formate in isopropanol/water (95:5 v/v) as mobile phase B. A 20-min gradient was employed, starting at 15% B at 0 min, 43% B at 2.0 min, 46% B at 6.3 min, 75% B at 10.4 min, 85% B at 10.6 min, 86% B at 10.8 min, 88% B at 13.5 min, and 98% B at 14.5 min, which was held until 15.0 min before allowing 5 min of column re-equilibration at the initial conditions. Injection volumes were 4.0 μL for positive ionization mode and 8.0 μL for negative ionization mode. Data were acquired in data-dependent acquisition mode over an m/z range of 150–2000, with MS/MS collision energies set between 10 and 60 eV.

### Data Processing

LC-MS data from 25 injections per extract type (20 sample injection plus 5 QCs) were independently processed in positive and negative ionization using NovaMT LipidScreener software v 1.0.0. Positive- and negative-ion features were merged into a single feature-intensity matrix. Lipid identification was performed based on a three-tier identification approach based on MS/MS spectral similarity, retention time, and accurate mass: 1) Tier 1 (MS/MS): MS/MS match score ≥500; precursor m/z error ≤20.0 ppm and 5.0 mDa; 2) Tier 2 (MS/MS): MS/MS match score <500; precursor m/z error ≤20.0 ppm and 5.0 mDa; and 3) Tier 3 (MS): Mass match with m/z error ≤20.0 ppm and 5.0 mDa. Identified features from all tiers were used for normalization and statistical analysis. Lipids with identical identifications, regardless of retention times, were summed prior to statistical analysis. Unidentified features were not employed for statistics. Identified features were normalized by internal standards and the median intensity ratio prior to any statistical analysis to correct for any sample variations. This ensured slight variations in EV amount between samples did not impact lipidomic analysis.

### Statistical Analysis

All lipids identified were detected in all samples. Statistical analysis was performed using MetaboAnalyst 6.0 Software (Pang *et al*. 2024) without blinding. For multivariate statistics, features with near-constant values between the groups (the 25% features with the lowest relative standard deviation for all samples) were filtered out. The dataset was also auto-scaled. No other filtering, normalization, transformation, or scaling methods were employed before multivariate statistical analysis. Multivariate statistics including PLS-DA, hierarchical clustering heatmaps and correlation heatmaps were employed to assess the ability to differentiate samples by their lipidomic profiles. For univariate statistics and pathways analysis, no extra filtering or data scaling methods were applied. Univariate statistics including t-tests and volcano plots combing fold change and t-test analysis were employed to identify distinct lipidomic changes between groups that may not be apparent with multivariate methods. For pathway analysis BioPAN software was used (Gaud *et al*. 2021). LipidLynxX (Ni and Fedorova 2020) was first used to convert lipid names to BioPAN compatible bulk nomenclature. Due to the size of the dataset all non-BioPAN recognized lipid classes were removed, and pathway analysis was conducted within lipid categories without removing fatty acids. BioPAN groups included: Glycerophospholipids, glycerolipids and fatty acids. BioPAN software was then utilized for reaction pathway analysis at the species and class levels. For Cell:EV ratio analysis of distinct lipid species normality was assessed with a Shapiro-Wilk test then parametric and non-parametric data were analyzed with either an unpaired t-test or a Mann-Whitney test respectively. See **Table 1** for a full list of lipid subclass abbreviations utilized throughout this study. Tables with extended statistics can be found in the supplemental material.

**Table 1.**
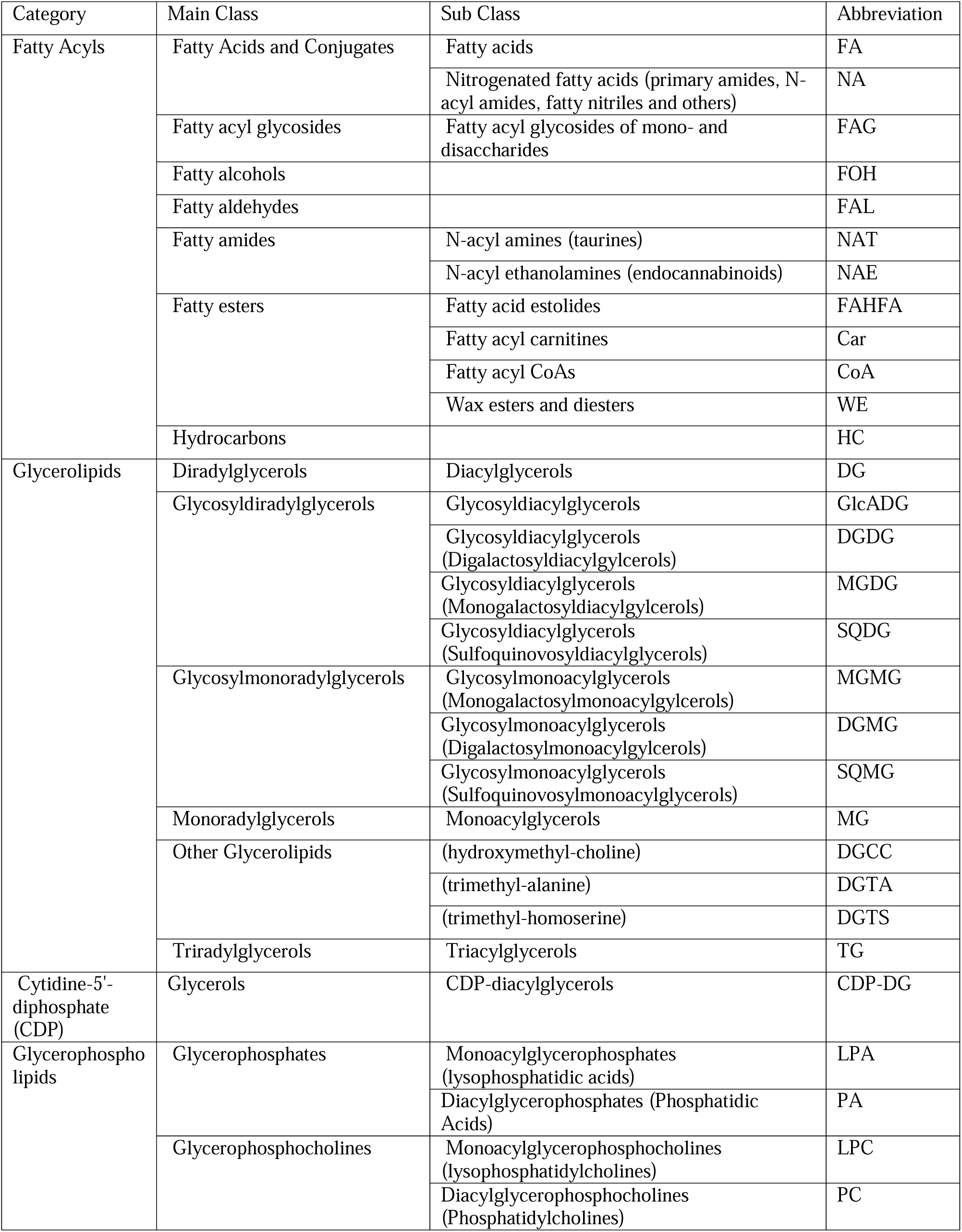
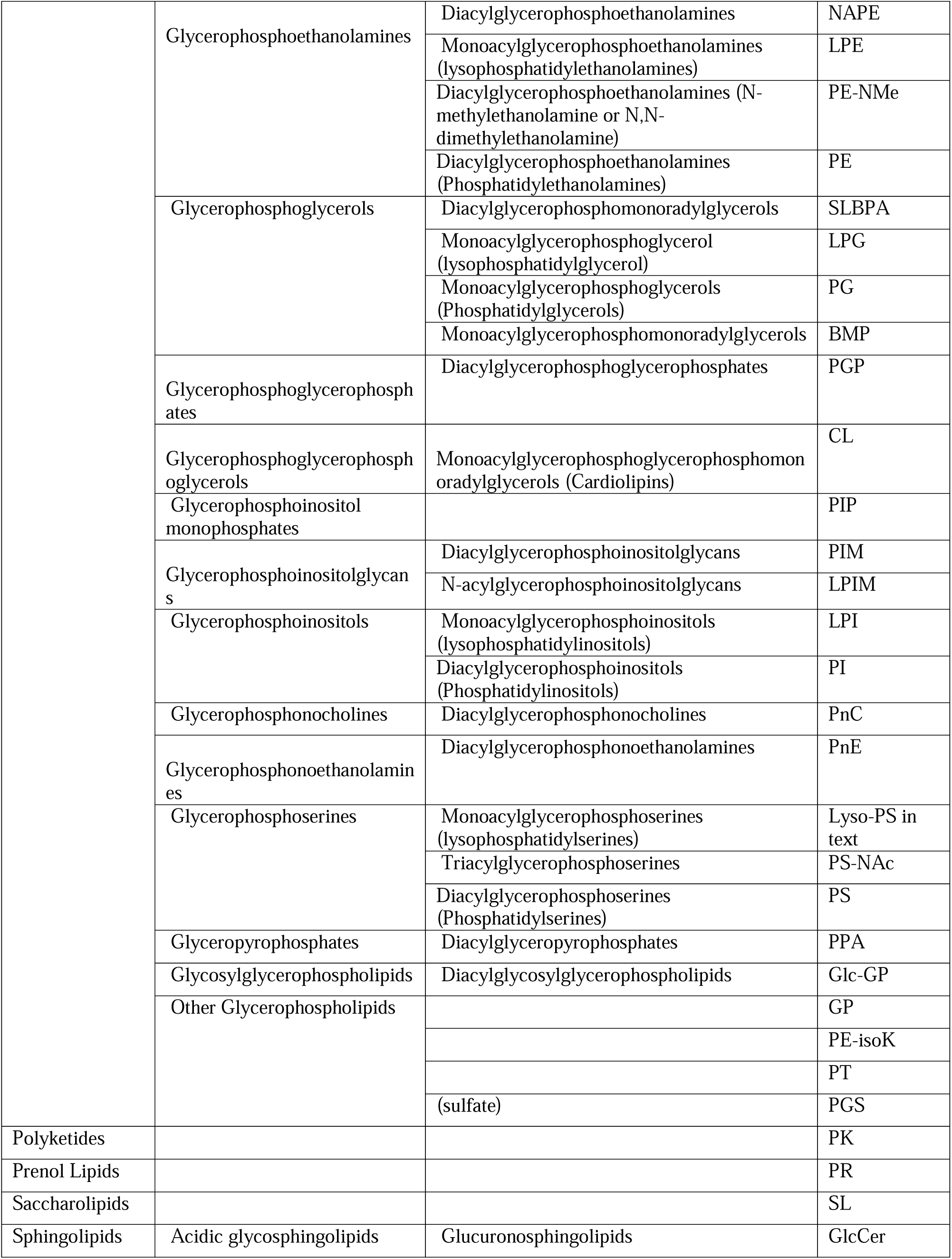
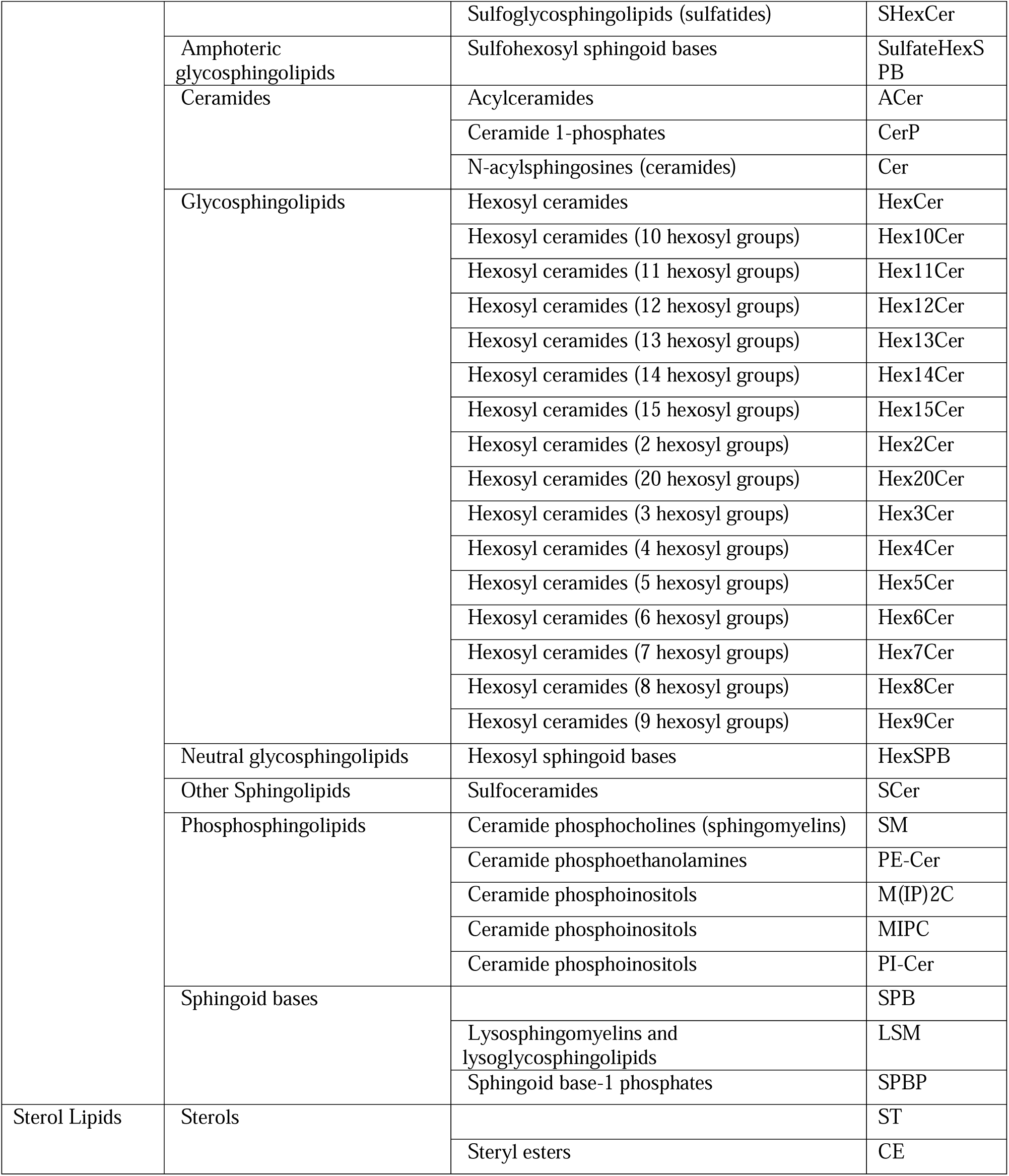
Lipid classifications and abbreviations used in study. The classification and shorthand notation of lipids followed the guidance of LipidMaps (https://www.lipidmaps.org/shorthand_nomenclature), MSDial (http://prime.psc.riken.jp/compms/msdial/lipidnomenclature.html), and Liebisch et al. J. Lipid Res. 2020, 61, 1–17 (DOI: https://doi.org/10.1194/jlr.S120001025) with the exception of lysophosphatidylserine. Since in the manuscript text LPS to refers to lipopolysaccharide, in-text reference to lysophosphatidylserine is abbreviated Lyso-PS. Please note text within figure 4 showcasing BioPAN results does utilize LPS to denote lysophosphatidylserines.

| Category | Main Class | Sub Class | Abbreviation |
| --- | --- | --- | --- |
| Fatty Acyls | Fatty Acids and Conjugates | Fatty acids | FA |
|  |  | Nitrogenated fatty acids (primary amides, N-acyl amides, fatty nitriles and others) | NA |
|  | Fatty acyl glycosides | Fatty acyl glycosides of mono- and disaccharides | FAG |
|  | Fatty alcohols |  | FOH |
|  | Fatty aldehydes |  | FAL |
|  | Fatty amides | N-acyl amines (taurines) | NAT |
|  |  | N-acyl ethanolamines (endocannabinoids) | NAE |
|  | Fatty esters | Fatty acid estolides | FAHFA |
|  |  | Fatty acyl carnitines | Car |
|  |  | Fatty acyl CoAs | CoA |
|  |  | Wax esters and diesters | WE |
|  | Hydrocarbons |  | HC |
| Glycerolipids | Diradylglycerols | Diacylglycerols | DG |
|  | Glycosyldiradylglycerols | Glycosyldiacylglycerols | GlcADG |
|  |  | Glycosyldiacylglycerols (Digalactosyldiacylglycerols) | DGDG |
|  |  | Glycosyldiacylglycerols (Monogalactosyldiacylglycerols) | MGDG |
|  |  | Glycosyldiacylglycerols (Sulfoquinovosyldiacylglycerols) | SQDG |
|  | Glycosylmonoradylglycerols | Glycosylmonoacylglycerols (Monogalactosylmonoacylglycerols) | MGMG |
|  |  | Glycosylmonoacylglycerols (Digalactosylmonoacylglycerols) | DGMG |
|  |  | Glycosylmonoacylglycerols (Sulfoquinovosylmonoacylglycerols) | SQMG |
|  | Monoradylglycerols | Monoacylglycerols | MG |
|  | Other Glycerolipids | (hydroxymethyl-choline) | DGCC |
|  |  | (trimethyl-alanine) | DGTA |
|  |  | (trimethyl-homoserine) | DGTS |
|  | Triradylglycerols | Triacylglycerols | TG |
| Cytidine-5'-diphosphate (CDP) | Glycerols | CDP-diacylglycerols | CDP-DG |
| Glycerophospholipids | Glycerophosphates | Monoacylglycerophosphates (lysophosphatidic acids) | LPA |
|  |  | Diacylglycerophosphates (Phosphatidic Acids) | PA |
|  | Glycerophosphocholines | Monoacylglycerophosphocholines (lysophosphatidylcholines) | LPC |
|  |  | Diacylglycerophosphocholines (Phosphatidylcholines) | PC |
|  | Glycerophosphoethanolamines | Diacylglycerophosphoethanolamines | NAPE |
|  |  | Monoacylglycerophosphoethanolamines (lysophosphatidylethanolamines) | LPE |
|  |  | Diacylglycerophosphoethanolamines (N-methylethanolamine or N,N-dimethylethanolamine) | PE-NMe |
|  |  | Diacylglycerophosphoethanolamines (Phosphatidylethanolamines) | PE |
|  | Glycerophosphoglycerols | Diacylglycerophosphomonoradylglycerols | SLBPA |
|  |  | Monoacylglycerophosphoglycerol (lysophosphatidylglycerol) | LPG |
|  |  | Monoacylglycerophosphoglycerols (Phosphatidylglycerols) | PG |
|  |  | Monoacylglycerophosphomonoradylglycerols | BMP |
|  | Glycerophosphoglycerophosphates | Diacylglycerophosphoglycerophosphates | PGP |
|  | Glycerophosphoglycerophosphoglycerols | Monoacylglycerophosphoglycerophosphomonoradylglycerols (Cardiolipins) | CL |
|  | Glycerophosphoinositol monophosphates |  | PIP |
|  | Glycerophosphoinositolglycans | Diacylglycerophosphoinositolglycans | PIM |
|  |  | N-acylglycerophosphoinositolglycans | LPIM |
|  | Glycerophosphoinositols | Monoacylglycerophosphoinositols (lysophosphatidylinositols) | LPI |
|  |  | Diacylglycerophosphoinositols (Phosphatidylinositols) | PI |
|  | Glycerophosphonocholines | Diacylglycerophosphonocholines | PnC |
|  | Glycerophosphonoethanolamines | Diacylglycerophosphonoethanolamines | PnE |
|  | Glycerophosphoserines | Monoacylglycerophosphoserines (lysophosphatidylserines) | Lyso-PS in text |
|  |  | Triacylglycerophosphoserines | PS-NAc |
|  |  | Diacylglycerophosphoserines (Phosphatidylserines) | PS |
|  | Glyceropyrophosphates | Diacylglyceropyrophosphates | PPA |
|  | Glycosylglycerophospholipids | Diacylglycosylglycerophospholipids | Glc-GP |
|  | Other Glycerophospholipids |  | GP |
|  |  |  | PE-isoK |
|  |  |  | PT |
|  |  | (sulfate) | PGS |
| Polyketides |  |  | PK |
| Prenol Lipids |  |  | PR |
| Saccharolipids |  |  | SL |
| Sphingolipids | Acidic glycosphingolipids | Glucuronosphingolipids | GlcCer |
|  |  | Sulfoglycosphingolipids (sulfatides) | SHexCer |
|  | Amphoteric glycosphingolipids | Sulfohexosyl sphingoid bases | SulfateHexSPB |
|  | Ceramides | Acylceramides | ACer |
|  |  | Ceramide 1-phosphates | CerP |
|  |  | N-acylsphingosines (ceramides) | Cer |
|  | Glycosphingolipids | Hexosyl ceramides | HexCer |
|  |  | Hexosyl ceramides (10 hexosyl groups) | Hex10Cer |
|  |  | Hexosyl ceramides (11 hexosyl groups) | Hex11Cer |
|  |  | Hexosyl ceramides (12 hexosyl groups) | Hex12Cer |
|  |  | Hexosyl ceramides (13 hexosyl groups) | Hex13Cer |
|  |  | Hexosyl ceramides (14 hexosyl groups) | Hex14Cer |
|  |  | Hexosyl ceramides (15 hexosyl groups) | Hex15Cer |
|  |  | Hexosyl ceramides (2 hexosyl groups) | Hex2Cer |
|  |  | Hexosyl ceramides (20 hexosyl groups) | Hex20Cer |
|  |  | Hexosyl ceramides (3 hexosyl groups) | Hex3Cer |
|  |  | Hexosyl ceramides (4 hexosyl groups) | Hex4Cer |
|  |  | Hexosyl ceramides (5 hexosyl groups) | Hex5Cer |
|  |  | Hexosyl ceramides (6 hexosyl groups) | Hex6Cer |
|  |  | Hexosyl ceramides (7 hexosyl groups) | Hex7Cer |
|  |  | Hexosyl ceramides (8 hexosyl groups) | Hex8Cer |
|  |  | Hexosyl ceramides (9 hexosyl groups) | Hex9Cer |
|  | Neutral glycosphingolipids | Hexosyl sphingoid bases | HexSPB |
|  | Other Sphingolipids | Sulfoceramides | SCer |
|  | Phosphosphingolipids | Ceramide phosphocholines (sphingomyelins) | SM |
|  |  | Ceramide phosphoethanolamines | PE-Cer |
|  |  | Ceramide phosphoinositols | M(IP)2C |
|  |  | Ceramide phosphoinositols | MIPC |
|  |  | Ceramide phosphoinositols | PI-Cer |
|  | Sphingoid bases |  | SPB |
|  |  | Lysosphingomyelins and lysoglycosphingolipids | LSM |
|  |  | Sphingoid base-1 phosphates | SPBP |
| Sterol Lipids | Sterols |  | ST |
|  |  | Steryl esters | CE |

## Results

Lipid changes following 24 hours of either no treatment (NT, n= 5) or LPS exposure (n = 5) of both microglia and their released EVs were assessed using LC-MS/MS. A total of 3626 lipid species were detected and identified. **Figure 2A** depicts the relative number of lipid species detected per lipid category in all samples with this protocol. All lipids detected and identified were detected in all samples including both cells and EVs. First, we assessed if the lipid profiles of detected and identified lipids could be used to differentiate the LPS exposed cells, NT cells and their respective EVs using multivariate statistics. Partial least squares discriminant analysis (PLS-DA) indicated that LPS treated microglia could be differentiated from NT microglia and their respective EVs by the component 1 (37.4%), and further by the component 2 (12%) (**Fig. 2B,C**). As shown in **Figure 2D**, further discrimination can be observed in the synchronized 3D plot demonstrating differentiation by three component analysis (component 1: 37.4%, component 2: 16%, component 3: 7.2 %) of treated and non-treated cells and their respective EVs.

**Figure 2.**
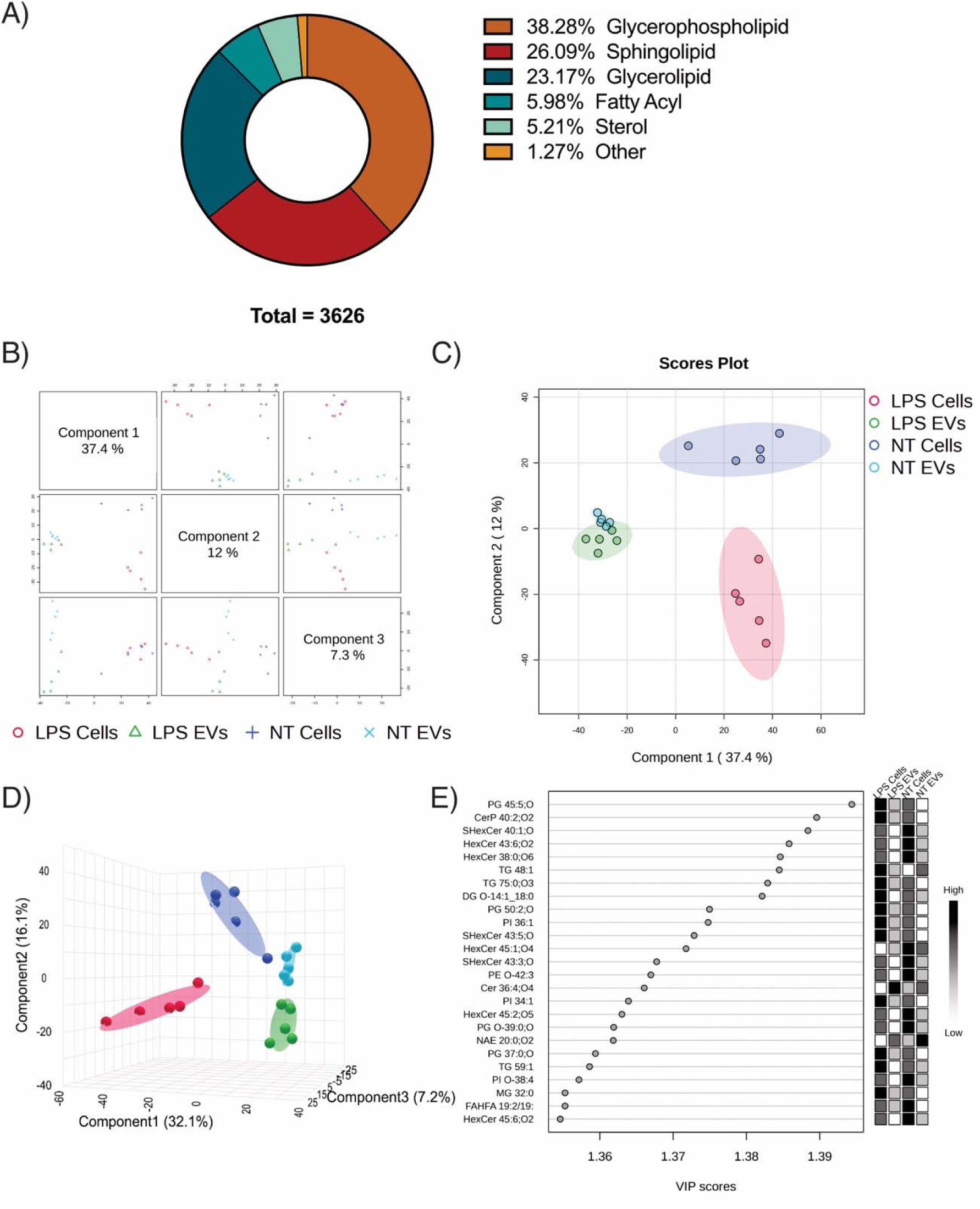
Untargeted lipidomic profiling can differentiate non-treated cells, LPS exposed cells and their respective EVs. A) Visual representation of lipid categories detected in all samples. B) Three component partial least squares discriminant analysis (PLS-DA) overview. C) PLS-DA 2D scores plot shows complete differentiation of cell types with overlap of 95% confidence region between groups of EVs. D) Synchronized 3D plot demonstrating differentiation of treated and non-treated cells and their respective EVs. E) Component three variable importance in projection (VIP) plot showing the 25 most important lipids for discriminating the sample groups with grey scale-coloured boxes depicting the relative abundance of each lipid in each experimental group.

Component three variable importance in projection (VIP) scores from PLS-DA revealed PG 45:5;O, CerP 40:2;O2 and SHexCer 40:1;O to be the three lipids most important in differentiating the experimental groups (**Fig. 2E**). The model was evaluated using cross validation to assess predictive ability (Q2) and goodness of fit (R2), while permutation testing with 100 permutations, 3 components was applied to confirm the significance of the model. Model evaluation revealed a Q2 = 0.80893, R2 = 0.92105 and a p-value < 0.01 indicating that PLS-DA can differentiate LPS treated microglia, NT microglia and their respective EVs.

Next, we aimed to interrogate distinct lipid changes in EV populations and how EVs correlated with their parent cells. First, we utilized an Euclidean heatmap with hierarchical clustering of 120 of the most significant lipid species based on t-test/ANOVA comparison of groups which visualized distinct lipid changes in LPS treated cells and NT cells, as well as the EVs secreted from the LPS treated and NT cells **(Fig. 3A**). To further interrogate differences that may not be apparent in the Euclidean heatmap, we compared the distinct cell groups, and their relation to their respective EVs using volcano plot analysis combining fold change and t-test analysis (absolute fold change ≥ 1.4 and a raw p-value threshold of 0.05). First, we compared NT and LPS cells. A total of 296 significantly decreased and 514 significantly increased lipid species were identified in LPS treated cells (**Fig. 3B**). We then assessed the discrete differences in the abundance of detected lipid species between both cell treatments and their EVs individually. Volcano plot analysis comparing NT and NT EVs identified 859 significantly decreased and 783 significantly increased lipid species in NT EVs (**Fig. 3C**); while, comparing LPS cells relative to LPS EVs revealed 1143 significantly decreased and 822 significantly increased lipid species in LPS EVs (**Fig. 3D**). We then aimed to assess if the overall lipidomic profile of EVs correlated with their parent cells. As visualized in a heatmap (**Fig. 3E**), spearman rank correlation indicated no correlation between groups of EVs and their parent cells. We next considered if there were distinct differences in the lipid profiles of EVs released from LPS treated cells relative to those released from NT cells. Volcano plot analysis identified 167 significantly decreased and 119 significantly increased lipid species in LPS EVs relative to NT EVs (**Fig. 3F**).

**Figure 3.**
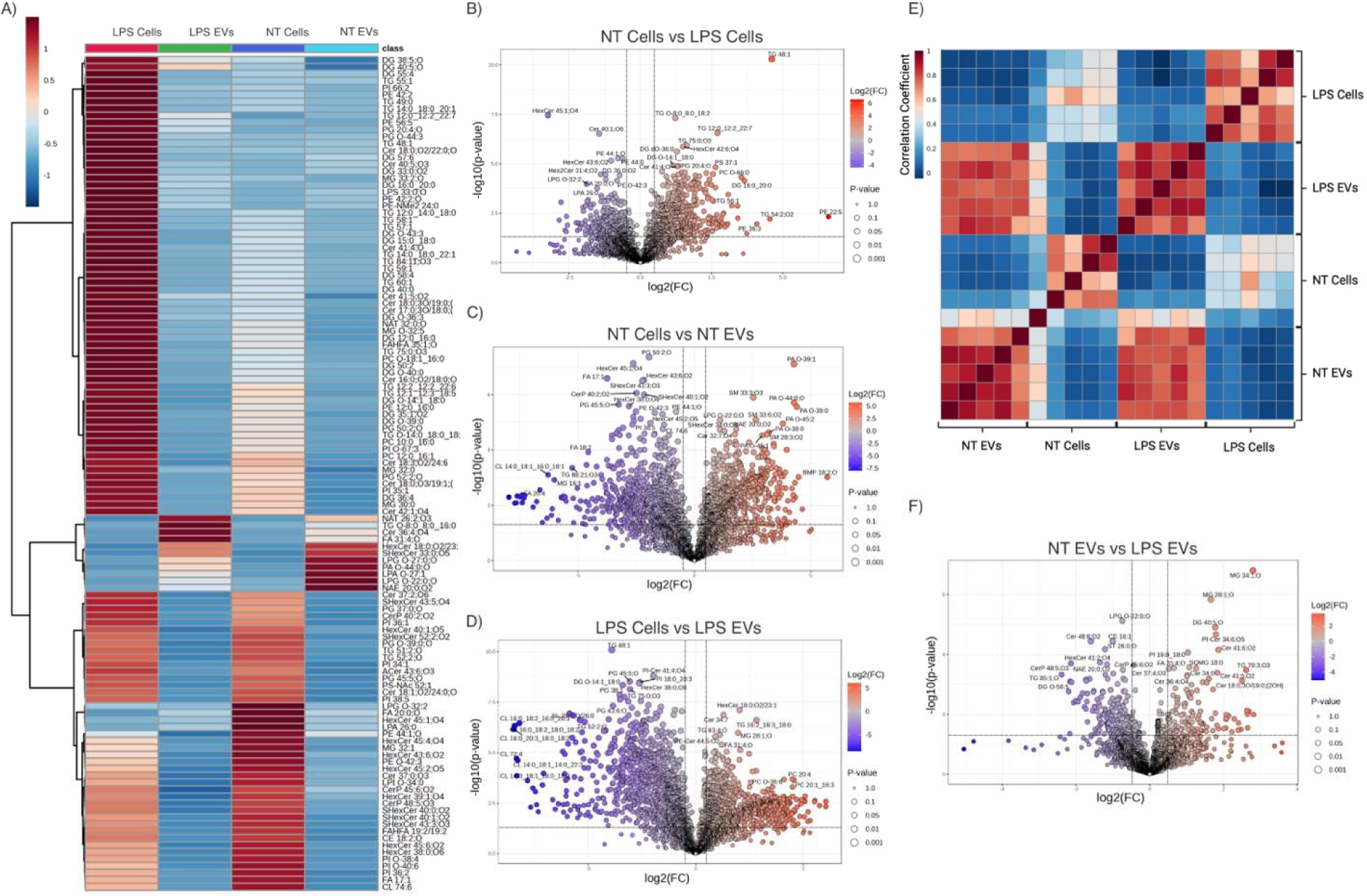
Distinct lipid species differentiate EVs from LPS exposed cells. A) Euclidean heatmap with hierarchical clustering of 120 of the most significant differences based on t-test/ANOVA comparison of groups. B) Volcano plot analysis comparing LPS and NT Cells identified 296 significantly decreased and 514 significantly increased lipid species in LPS treated cells. All volcano plots combine fold change and t-test analysis and represent an absolute fold change ≥ 1.4 and a raw p-value threshold of 0.05. C) Volcano plot analysis comparing NT EVs and NT cells identified 859 significantly decrease and 783 significantly increased lipid species in NT EVs. D) Volcano plot analysis identified 1143 significantly decreased and 822 significantly increased lipid species in LPS EVs relative to LPS cells. E) Spearman rank correlation heatmap. F) Volcano plot analysis identified 167 significantly decreased and 119 significantly increased lipid species in LPS EVs relative to NT EVs. G) Serial t-test analysis of all detected and identified lipid species identified 405 significantly altered lipids between LPS EVs and NT EVs.

We then sought to determine if distinct lipid pathways were activated or suppressed in LPS exposed microglia relative to NT microglia. We conducted BioPAN analysis for reactions involving glycerophospholipids and glycerolipids at the class (**Fig. 4A,E,F**) and species (**Fig. 4. B,C**) level, and for fatty acids (**Fig. 4A,D**). **Table 2**. Summarizes the BioPAN results of all activated and suppressed reactions and the probable genes associated with these changes. Of note, at the class level for glycerolipids, results revealed an increase in diacylglycerol (DG) and triacylglycerol (TG) lipids. At the class level for glycerophospholipids, BioPAN analysis identified an increase in reactions producing Lyso-PS and LPA species with a decrease in LPC formation.

**Figure 4.**
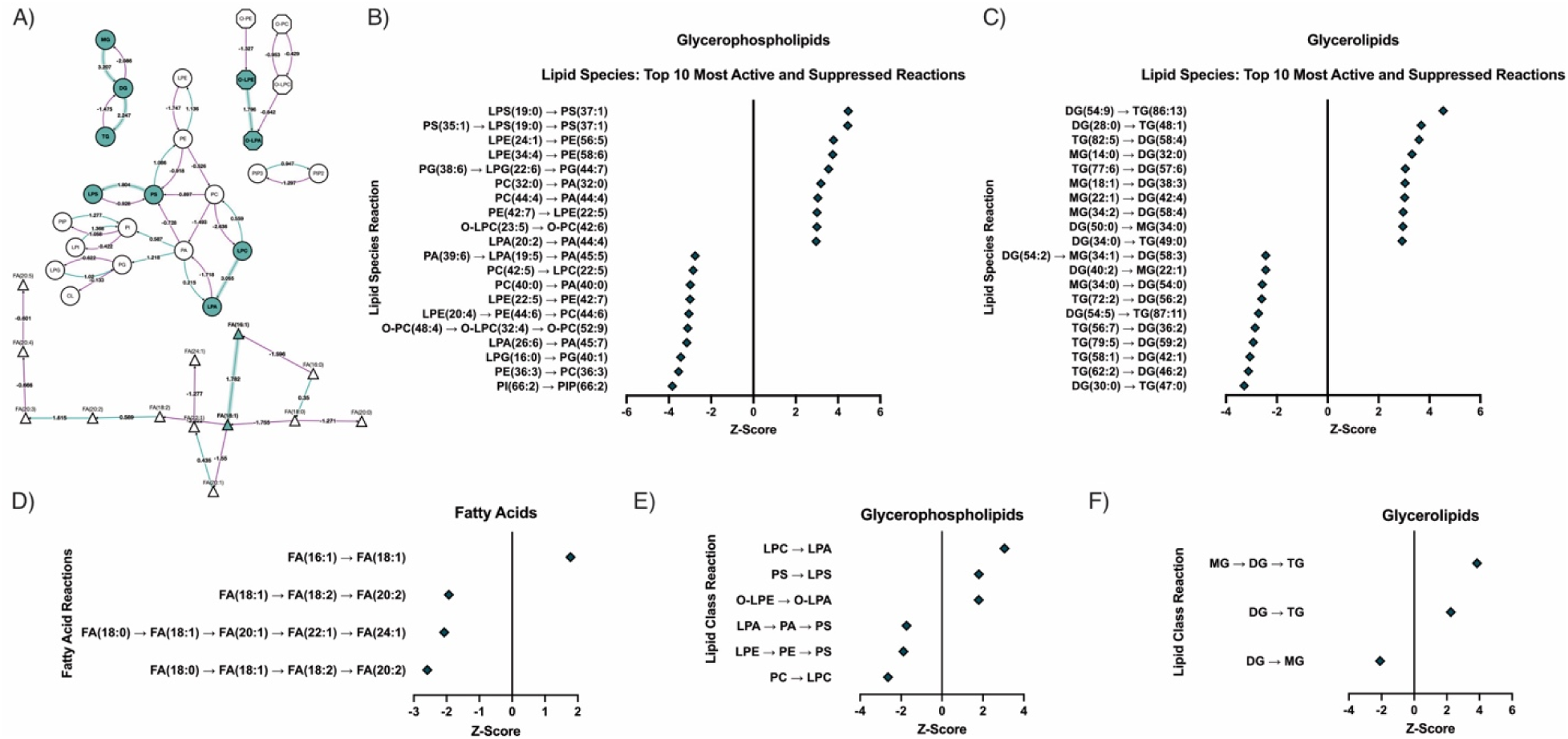
Distinct lipid reactions are activated and suppressed in LPS exposed cells. BioPAN analysis of lipid classes, lipid species and fatty acids revealed distinct reactions are activated and suppressed in LPS treated cells compared to NT cells. A) Diagram of activated reactions in lipid classes and fatty acids. Activated pathways are highlighted in green with respective Z-scores noted in reaction arrows. Negative Z-scores are shown in purple. B) Top 10 activated and suppressed lipid reactions at the species level falling under the glycerophospholipid category. C) Top 10 activated and suppressed lipid reactions at the species level falling under the glycerolipid category. D) Activated and suppressed fatty acid reactions in LPS exposed cells. E) Most activated and suppressed lipid class reactions in the glycerophospholipid category. F) Most activated and suppressed lipid class reactions in the glycerolipid category. *LPS within this figure denotes the lipid class lysophosphatidylserines. See Table 1 For the entire list of lipid subclass abbreviations.

**Table 2.**
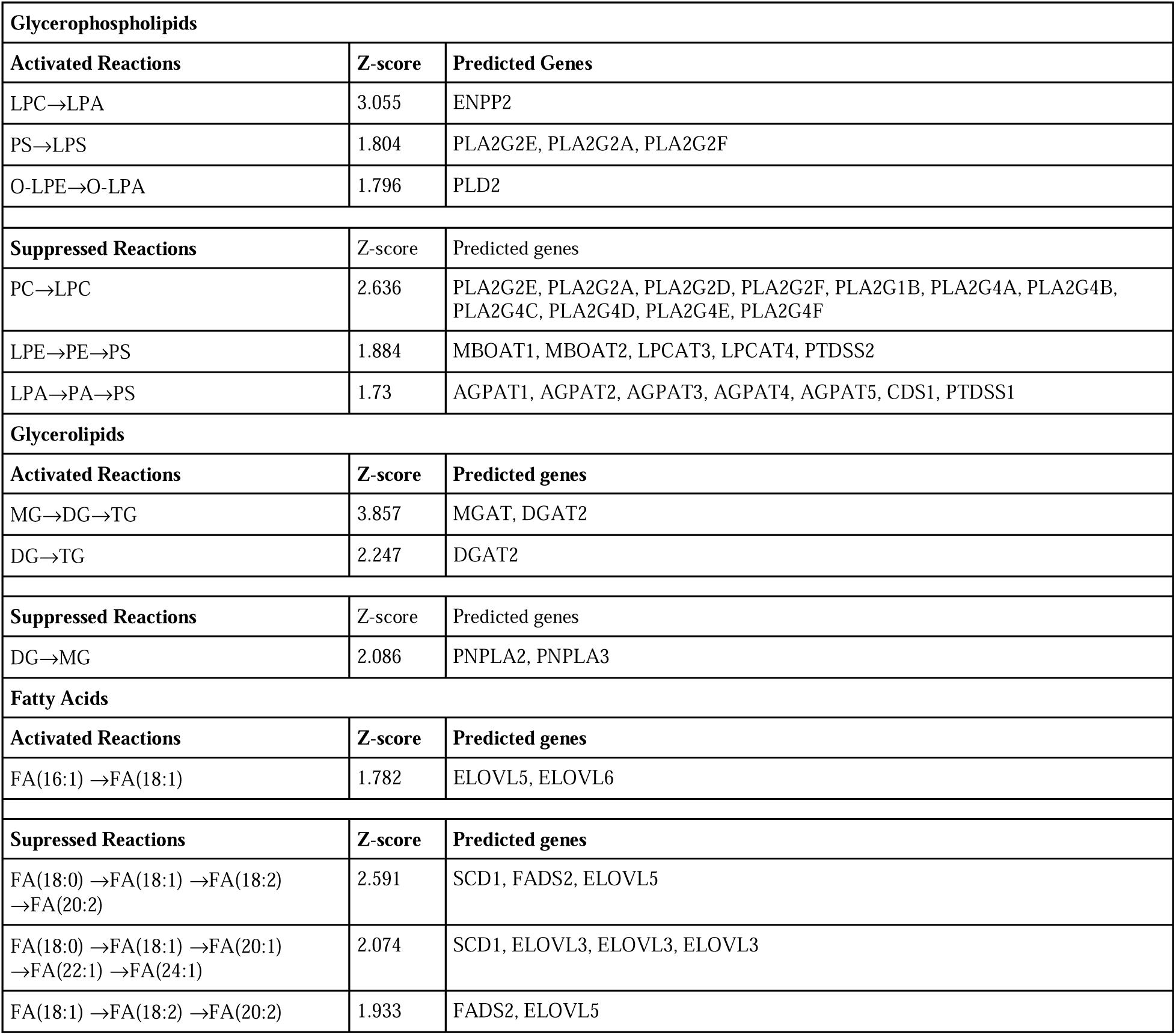
BioPAN analysis revealed activated and suppressed reactions in LPS treated microglia at the lipid class level. Analysis was conducted by lipid categories including glycerophospholipids, glycerolipids and fatty acids.

Given distinct lipid reactions are both activated and suppressed in LPS treated cells, we then assessed Cell:EV lipid ratios as a means provide potential insight into cellular EV based lipid clearance. As shown in **Figure 5A**, volcano plot analysis (fold change and t-test combination, absolute fold change ≥ 1.4 and a raw p-value threshold of 0.05) comparing all lipids identified by LPS Cell:EV and NT Cell:EV ratios identified 179 significantly decreased and 397 significantly increased lipid ratios with LPS exposure. Further visualization of this comparison through t-tests analysis comparing the Cell:EV ratio of all identified lipids in LPS and NT groups identified 582 lipids with significantly altered ratios (**Fig. 5B**). We then interrogated the ratio analysis of distinct lipid species thought to exert neurotoxic effects. The simple ganglioside species GM3 (d18:1/16:0) was among the lipid species significantly increased (p = 0.0491,, t = 2.317, degrees of freedom (df) = 8) in the Cell:EV ratio of LPS treated cells (**Fig. 5C**). We further interrogated lyso-phosphatidylcholines (LPCs) and oxidized phosphatidylcholines (OxPCs). Two LPCs were identified to have significantly altered ratios in the LPS treated group. LPC 30:4 was found to have a significantly increased ratio (p < 0.0001, t = 7.205, df = 8) with LPS treatment (**Fig. 5D);** while LPC 22:5 was found to have a significantly decreased (p = 0.0317, U = 2) ratio with LPS treatment (**Fig. 5E**). Two OxPCs were also found to have altered ratios with LPS treatment. PC 30:0;O was found to have an increased ratio (p = 0.0360, t = 2.517, df = 8) with LPS treatment (**Fig. 5F**); while PC 43:11;O was shown to have a significantly decreased (p = 0.0079, U = 0) ratio with LPS treatment (**Fig. 5G**).

**Figure 5.**
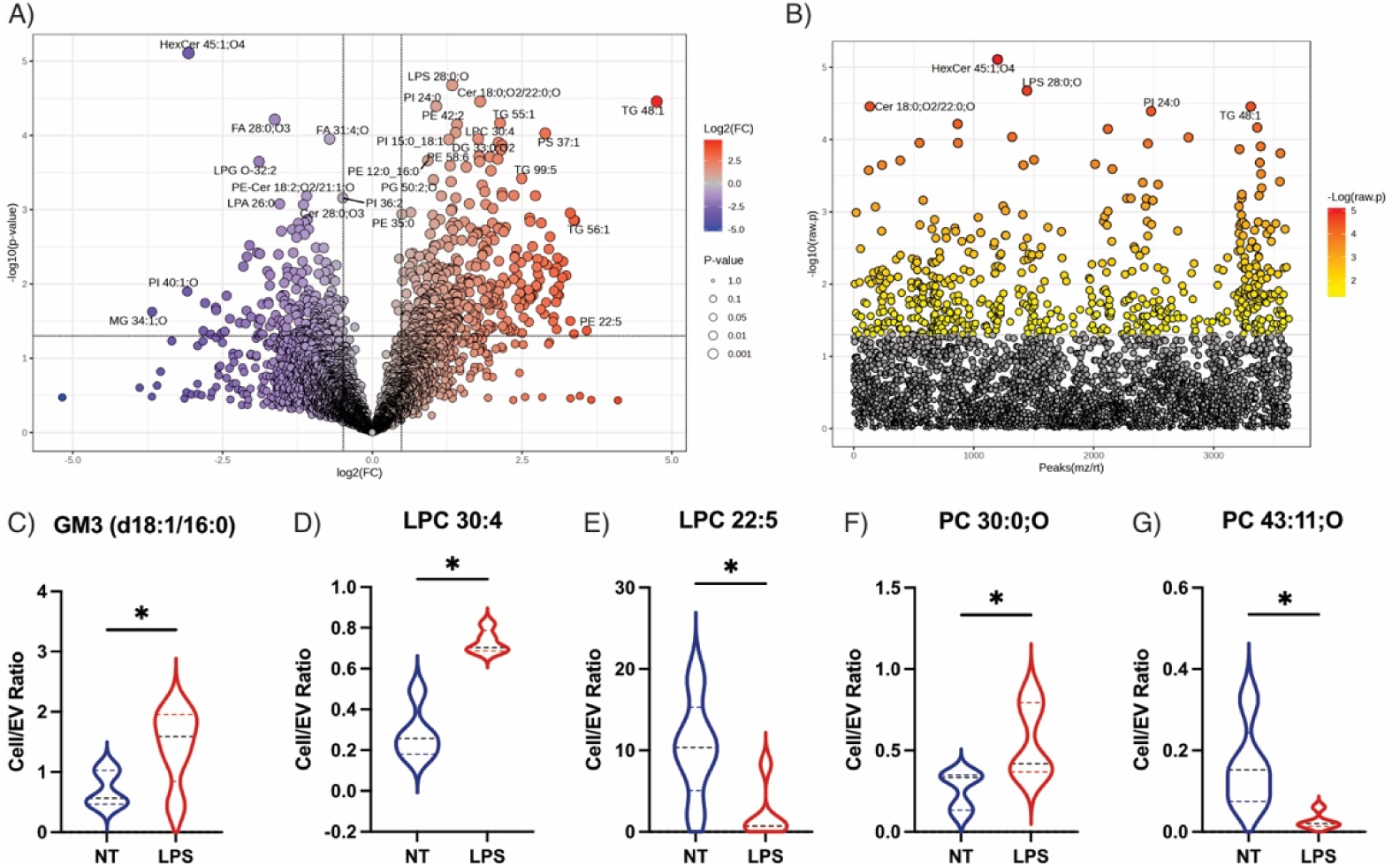
The Cell:EV ratio of distinct lipid species differs between LPS and non-treated cells. A) Volcano plot analysis (fold change and t-test combination, absolute fold change ≥ 1.4 and a raw p-value threshold of 0.05) comparing LPS Cell:EV and NT Cell:EV ratios identified 179 significantly decreased and 397 significantly increased lipid ratios with LPS treatment. B) T-tests comparing the Cell:EV ratio of all identified lipids in LPS and NT groups identified 582 lipids with significantly altered ratios. C) GM3 (d18:1/16:0) was among the lipid species significantly increased (p = 0.0491, t = 2.317, degrees of freedom (df) = 8) in the Cell:EV ratio of LPS treated cells. D) Two significantly altered lyso-phosphatidylcholines (LPCs) were detected between treatments. LPC 30:4 was found to have a significantly increased ratio (p < 0.0001, t = 7.205, df = 8) with LPS treatment; while, E) LPC 22:5 was found to have a significantly decreased (p = 0.0317, U = 2) ratio with LPS treatment. F) Additionally, two OxPCs species were found to have altered ratios with LPS treatment. PC 30:0;O was found to have an increased ratio (p = 0.0360, t = 2.517, df = 8) with LPS treatment; while, G) PC 43:11;O was shown to have a significantly decreased (p = 0.0079, U = 0) ratio with LPS treatment.

## Discussion

This study marks the first of its kind to interrogate the lipidomic relationship between microglia and their secreted EVs. The significance of this study is three-fold: 1) in a controlled *in vitro* experiment, we reveal that distinct lipidomic changes can differentiate microglia exposed to LPS from NT microglia and their EVs; while also demonstrating that distinct lipids are preserved, and others change with cell activation. Elucidating this relationship validates EV lipids as a potential future lipid-based biomarker capable of serving as a non-invasive window into changes in microglial function in the CNS. 2) We added to current literature on acute microglial activation with LPS by demonstrating acute pro-inflammatory activation of microglia results in the activation and suppression of distinct lipidomic pathways. This provides mechanistic insight into biological mechanisms and metabolic changes occurring with acute pro-inflammatory activation. Finally, 3) through analyzing the relationships between parent cells and their secreted EVs, we reveal lipid-specific changes and complex relationships within lipid classes in response to pro-inflammatory activation. These results demonstrate that lipid changes occurring with pro-inflammatory activation in microglia and their secreted EVs are highly complex and encourages future research investigating microglial/EV lipid changes with different experimental models and disease profiles. Collectively, this study utilized a controlled in vitro approach to elucidate foundational information on the lipidome of both microglia and their secreted EVs. This validates emerging research into EV lipids as novel and exciting potential biomarkers for neurological disease.

Through an untargeted approach and multi-variate analysis we were able to differentiate LPS-exposed BV-2 microglia, NT microglia and their respective EVs (**Fig. 2B-D**). **Figure 2E** demonstrates the top 25 most important lipids in differentiating these groups based on component 3 of PLS-DA which further differentiated EV groups. Of the top 25 lipids, 10 were noted to contain a ceramide backbone. Exosomes (small EVs) have previously been shown to be ceramide-rich and ceramides have been shown to be responsible for triggering the budding of exosomes into MVBs (Trajkovic *et al*. 2008). Acute LPS exposure in macrophages has also been shown to increase ceramides (MacKichan and DeFranco 1999) and a similar increase was recently observed in microglia (Blank *et al*. 2022). It is possible in this LPS paradigm, BV-2 microglia release smaller EVs possibly including exosomes, which may explain the important role of ceramides. We performed nanoflow cytometry to further characterize the size of EVs released from BV2s and demonstrated BV2s release EVs with an average size of 179.8 +/- 19.6 nm (Supplemental Figure 1A), further supporting this hypothesis. However, more work is needed to determine if this is stimulus, cell type and temporally dependent. We further interrogated the distinct lipid species changing between the experimental groups (**Fig. 3A,B**) and similarly found a prominent enrichment of distinct ceramide species in LPS treated microglia. Our results support ceramides as a critically important lipid for the differentiation of microglial activation phenotypes and their EVs.

From a neurological disease perspective accumulation of DG has also previously been shown in AD and PD brains (Wood *et al*. 2018; Wood *et al*. 2015; Chan *et al*. 2012). DG lipids are known to serve as lipid mediators of immune activation (Brose *et al*. 2004) and are also precursors for major glycerophospholipids and TG lipids (Coleman and Mashek 2011), potentially making both DG and TG lipids highly relevant in microglial activation. We found DG and TG lipids to be upregulated in LPS cells (**Fig. 3A,B**). This was further validated with BioPAN analysis (**Fig. 4A,C,F**) that revealed activated reactions metabolically shifting LPS treated cells towards TG synthesis. This was similarly shown in a recent publication utilizing a whole-cell matrix-assisted laser desorption/ionization (MALDI) MS whole cell fingerprinting workflow (Blank *et al*. 2022). Activated microglia also undergo alterations in mitochondrial function and the tricarboxylic acid cycle that can lead to TG accumulation within lipid droplets (Khatchadourian *et al*. 2012) and LPS treated microglia have been shown to accumulate TG-enriched lipid droplets (Marschallinger *et al*. 2020), further validating our findings. Intriguingly, our results showed that while specific TG species increased in LPS treated cells, this was not preserved with an equal abundance in EVs. Compared to NT microglia, TG (48:1) was significantly increased in LPS treated cells (**Fig. 3A,B**); however, TG (48:1) was significantly lower in LPS treated EVs compared to LPS cells (**Fig. 3A,D**). This was in keeping with our correlation analysis (**Fig. 3E**) which showed no relationship between the overall lipidomic profile of EVs and their parent cells, and with our comparison between LPS EVs and NT EVs showing TG lipids as one of the top three lipid classes significantly decreased in LPS EVs (**Fig. 3G,H**). Collectively, our results suggest that rather than correlation of overall lipidomic profiles, distinct lipids species are preserved between EVs and their parent cells, while distinct species change based on cellular activation.

Utilizing BioPAN software to analyze glycerophospholipids, we identified an increase in reactions producing Lyso-PS and LPA species with a decrease in LPC formation. LPS stimulation has previously been shown to upregulate LPA production through Autotaxin (ATX), a phospholipase D that converts LPC into LPA (Awada *et al*. 2014). This same pathway demonstrated activation based on our BioPAN analysis (**Fig. 4A**). Lyso-PS has also been shown to function in cellular morphology changes in microglia *in vitro* (Tokizane *et al*. 2017). In contrast to our BioPAN analysis result however, previous work has shown an increase in distinct LPC species with LPS treatment of microglia (Blank *et al*. 2022). It is important to note our overall dataset supports this as distinct LPC species were upregulated in LPS treated cells; however, this serves to highlight the importance of pathway and reaction-based assessment of lipidomic data. Limitations of these results should also be considered, as currently BioPAN software does not factor in oxidized lipid species which may alter this finding. Future work especially when assessing disease specific paradigms should factor lipid oxidation into analysis.

To deepen our analysis, we examined the Cell:EV ratio of specific lipid species with known neurotoxic properties, including simple gangliosides, LPCs, and OxPCs (Dong *et al*. 2021; Plemel *et al*. 2018; Wang and Whitehead 2020). Previous work has demonstrated GM3 colocalizes with plaques both in animal models (Wang *et al*. 2025; Kaya *et al*. 2018) and in the AD brain (Ollen-Bittle *et al*. 2024a; Michno *et al*. 2024). OxPCs have also been shown to accumulate in MS lesions (Dong *et al*. 2021) and can be converted either through hydrolysis reactions or enzymatic processes to LPCs, which are known to cause demyelination and are even used as a means of inducing spinal cord injury in animal models (Plemel *et al*. 2018). We hypothesized that we may see a decrease in the Cell:EV ratio of these neurotoxic species in LPS treated cells as they take on a proinflammatory phenotype and accumulate lipids, potentially indicating reduced clearance through EVs. While we did see this relationship with GM3 (**Fig. 5C**), of the two LPCs and OxPCs identified to have significant differences with LPS treatment, LPC 30:4 and PC 30:0;O were found to have a significantly increased ratio with LPS treatment, while LPC 22:5 and PC 43:11;O were shown to have a significantly decreased ratios (**Fig. 5D-G**). This suggests a dynamic relationship of these neurotoxic species in which cell activation and lipid class are not the only influences on the Cell:EV ratio. It remains probable physiologically that the type of pro-inflammatory exposure, extracellular environment, genetics and chronicity of pro-inflammatory state all factor into the Cell:EV ratios of distinct lipid species. Future work should consider assessment of these relationships when characterizing cell-specific changes in EV lipids.

Additional limitations and future directions should also be noted. While EV lipids may one day be promising biomarkers for neurological diseases important logistical challenges exist. Analysis of EVs requires collection, isolation, extraction and analysis processes that may take too long to be useful depending on the clinical setting. For instance, with current technology EV lipids would not likely be useful for acutely differentiating stroke subtype in an emergency room setting, until technical advances improve the temporal limitations (Ollen-Bittle *et al*. 2022). Additionally, this study serves as a critical foundation to a plethora of research to come before clinical utility even in more chronic conditions. Future research is needed to determine the effects of differential stimuli, different cell lines and primary cells, and different diseases both in experimental models and samples from human patients. This work provides foundational evidence to continue the pursuit of microglia lipid EV-based biomarkers in neurological disease.

Overall, this study provides the critical foundation requisite for the future study of lipid-based EV biomarkers in neurological diseases. For the first time, we validate that distinct lipid changes can differentiate LPS exposed microglia from NT microglia and their EVs; while demonstrating distinct lipids are preserved between the EVs and their parent cells, and others change with cell activation. Collectively, we validate a workflow for the *in vitro* characterization of EVs and most importantly demonstrate that microglial EV lipidome holds potential as a non-invasive window into functional changes in microglia.

## Acknowledgements

We would like to thank Dr. Lin Zhao for his guidance and expertise surrounding cell-culture and the isolation of EVs. We would also like to thank him for his work running the western blots and nanoflow cytometry used for characterizing the EVs. We would further like to acknowledge Reza Khazaee for his expertise and electron microscopy imaging of EVs. These figures can be found in the Supplemental Material.

## Data Availability

Data available upon request.

## Conflicts of Interest

No conflicts to report.

## Funding Sources

This study was supported by the Canadian Institutes for Health Research (CIHR) [202104PJT-461038] and the Natural Sciences and Engineering Research Council of Canada (NSERC) [RGPIN-2019-04742] to SNW. Additionally, NO and WW were supported by CIHR Canada Graduate Scholarships Doctoral Awards.

## List of Non-Lipid Abbreviations

EVs: Extracellular vesicles
LPS: lipopolysaccharides
AD: Alzheimer’s disease
Aβ: Amyloid beta
CSF: Cerebrospinal fluid
CNS: Central nervous system
BBB: Blood brain barrier
DMEM: Dulbecco’s Modified Eagle Medium
FBS: Fetal bovine serum
NT: Non-treated
TMIC: The Metabolomics Innovation Center
QC: Quality Control
PLS-DA: Partial least squares discriminant analysis
VIP: Variable importance in projection

## Supplemental Material

**Supplemental Figure 1.**
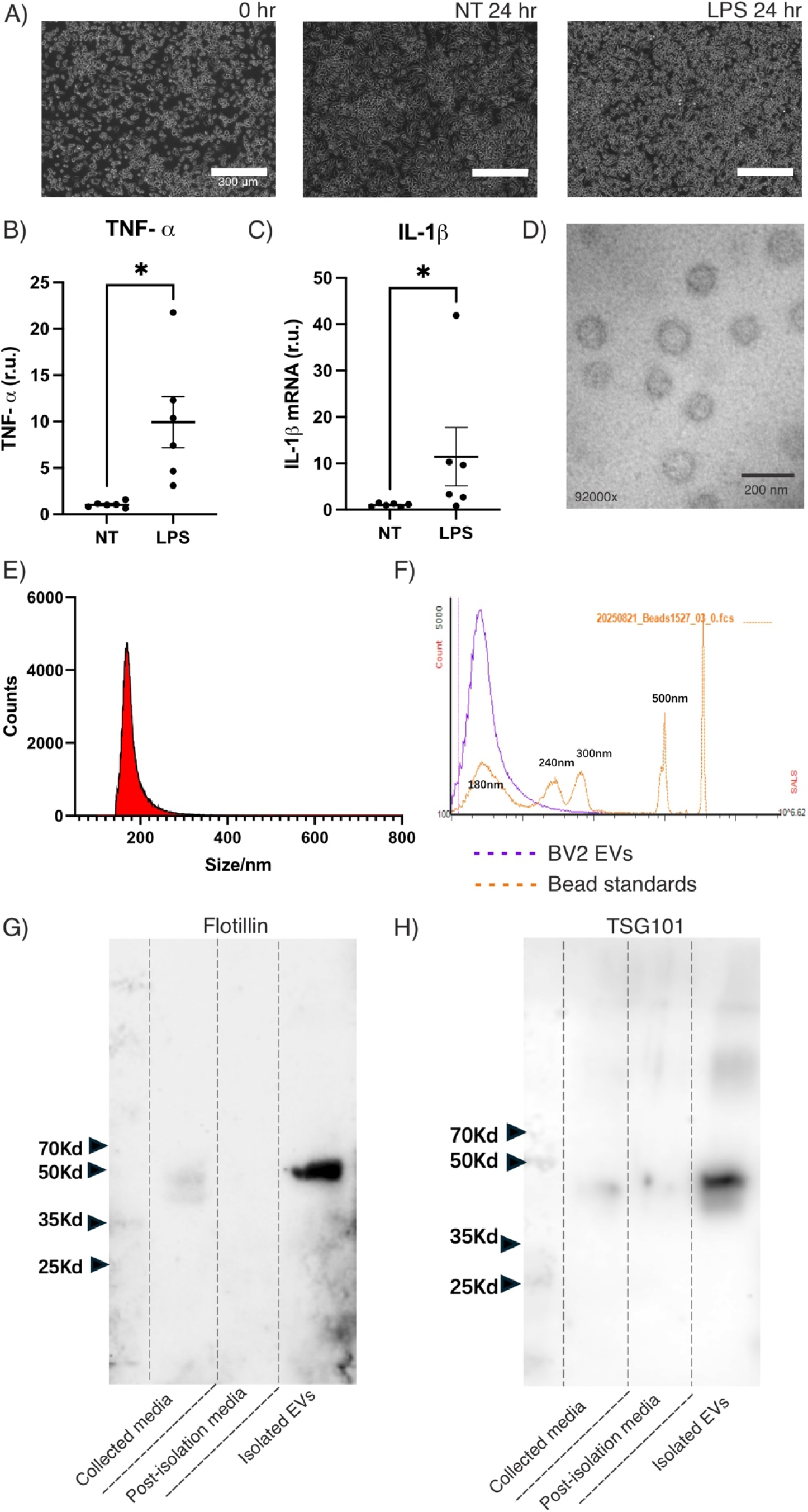
Cell and EV characterization. A) Microscope images of BV2 microglia pretreatment (0 hr), and after 24 hours of either control treatment with just media (NT) or 500 ng/ml LPS. B) Upregulation of TNF-α (p = 0.0090, n =6) and, C) IL-1β (p = 0.0260, n = 6) in LPS treated cells after 24 hours with this protocol. D) Electron microscopy 92000x image of BV2 EVs isolated using this protocol. E) Nanoflow cytometry showing the average size of isolated BV2 EVs is 179.8 +/- 19.6 nm. F) Nanoflow cytometry showing the average size of isolated BV2 EVs relative to bead standards of known size. G) Western blot showing an upregulation of flotillin in isolated BV-2 EVs relative to the initial filtered media and the media post isolation. All samples loaded with the same protein content. H) Western blot showing an upregulation of TSG101 in isolated BV-2 EVs relative to the initial filtered media and the media post isolation. Note: all Z scores shown as absolute values reflecting either activated or suppressed pathways comparing LPS vs NT.

**Supplemental Table 1.**
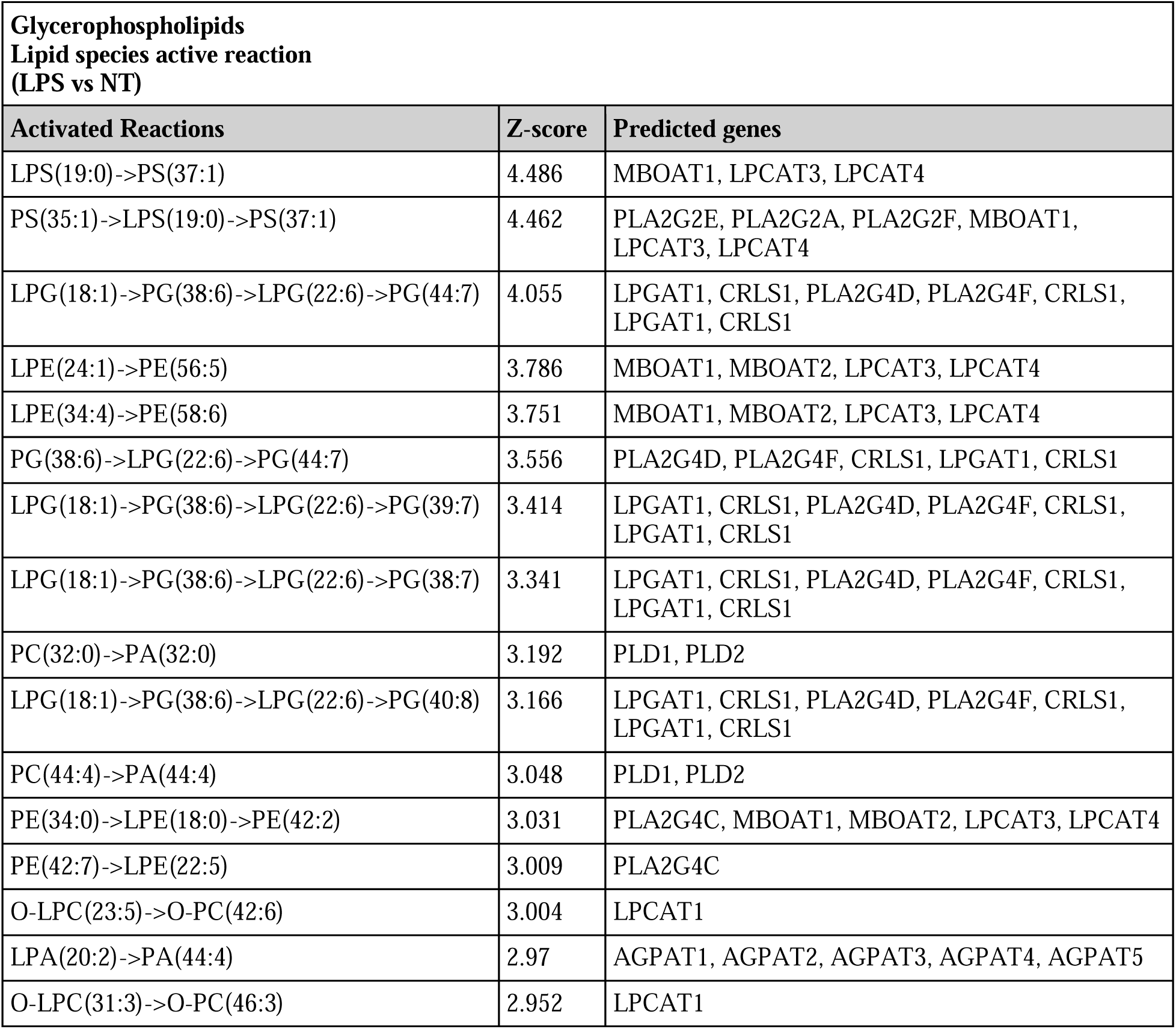
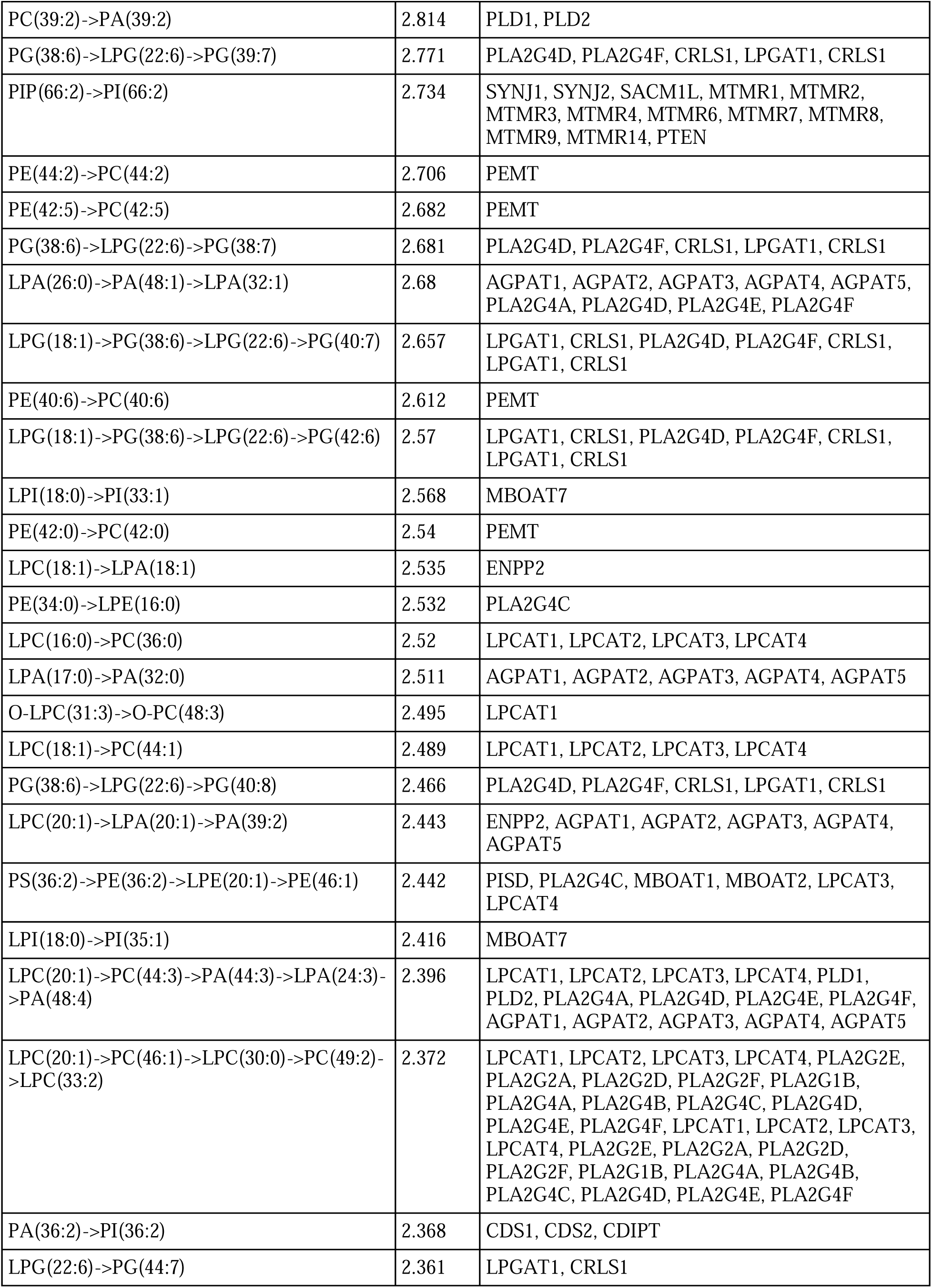
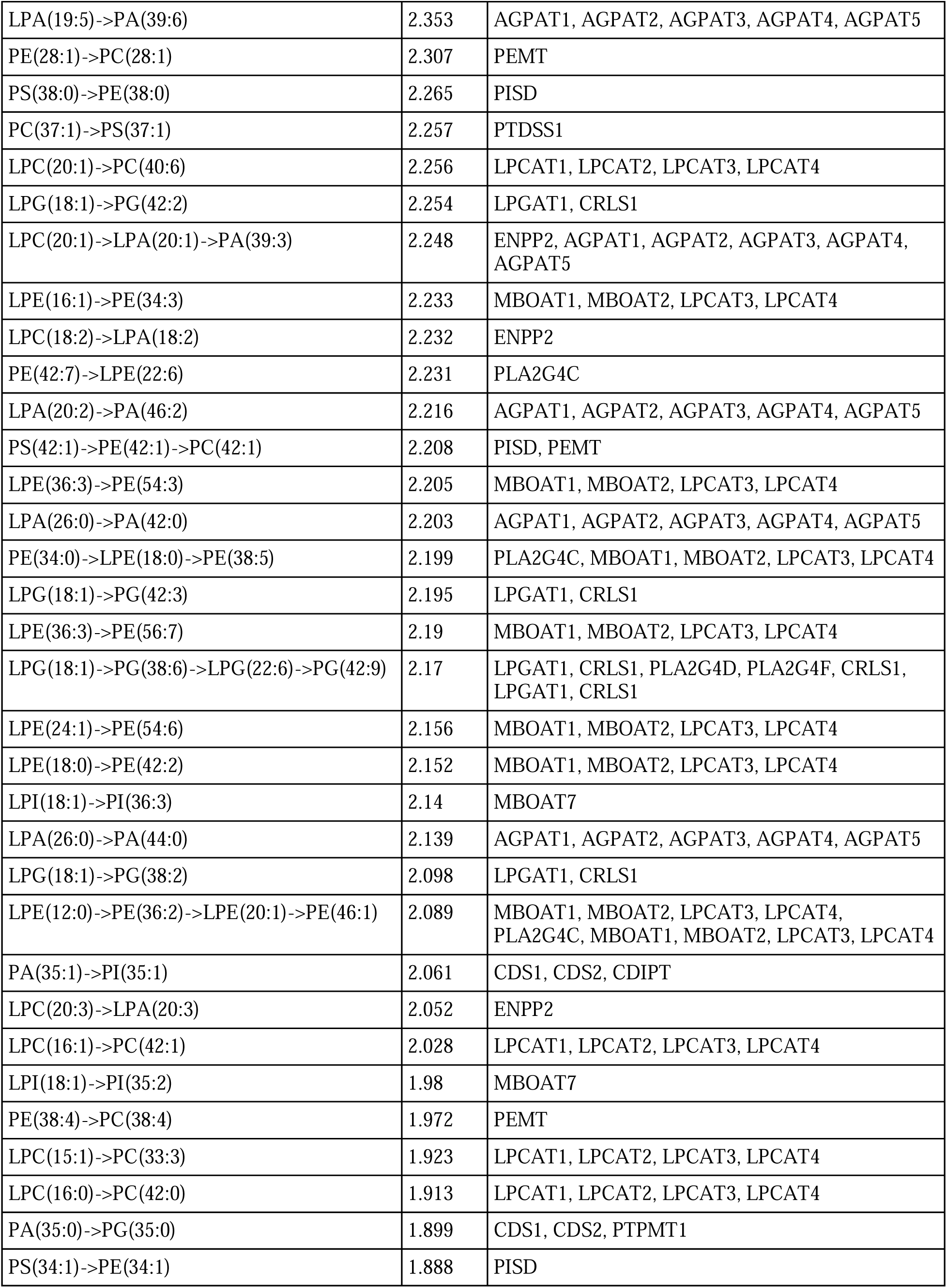
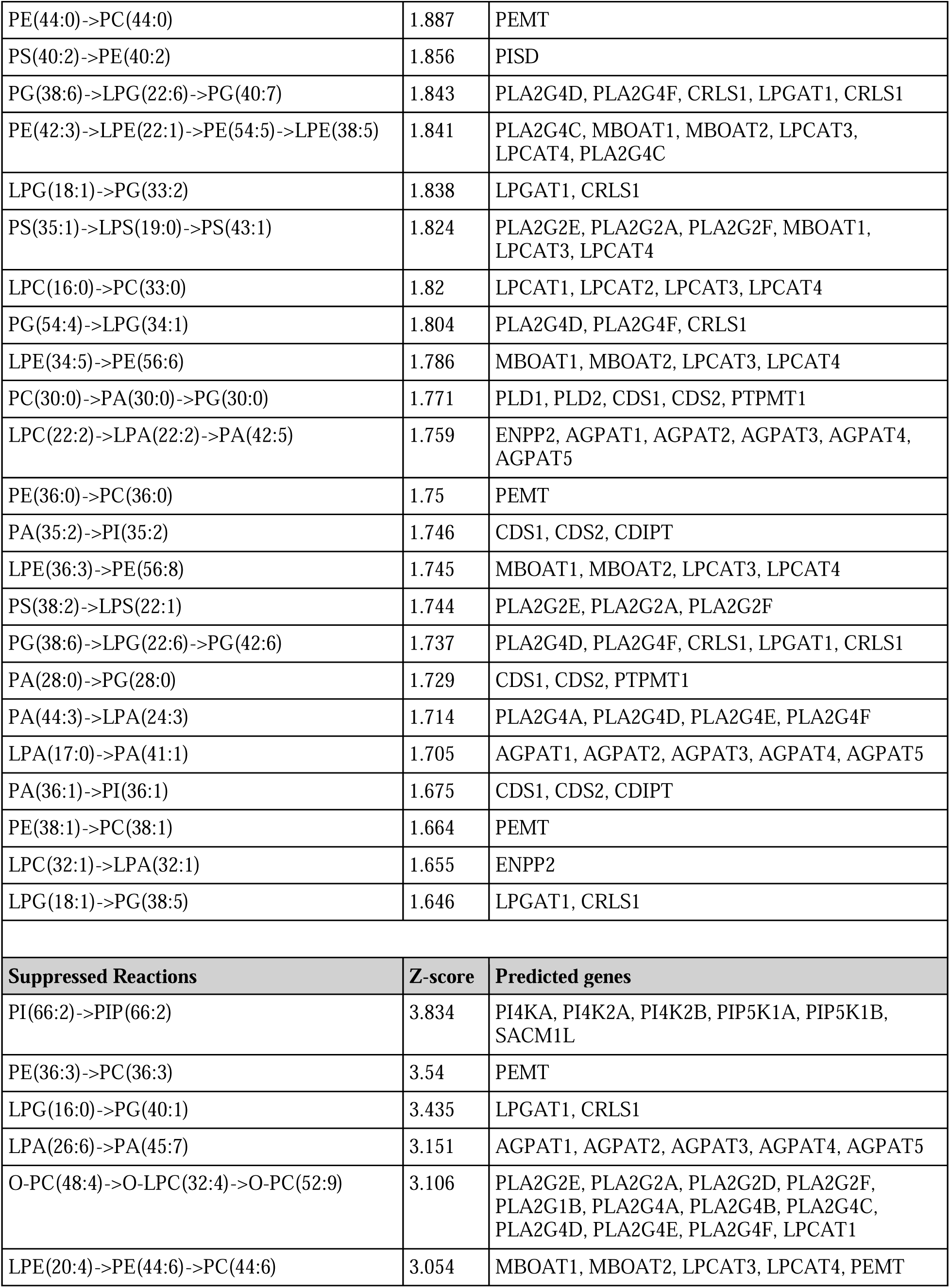
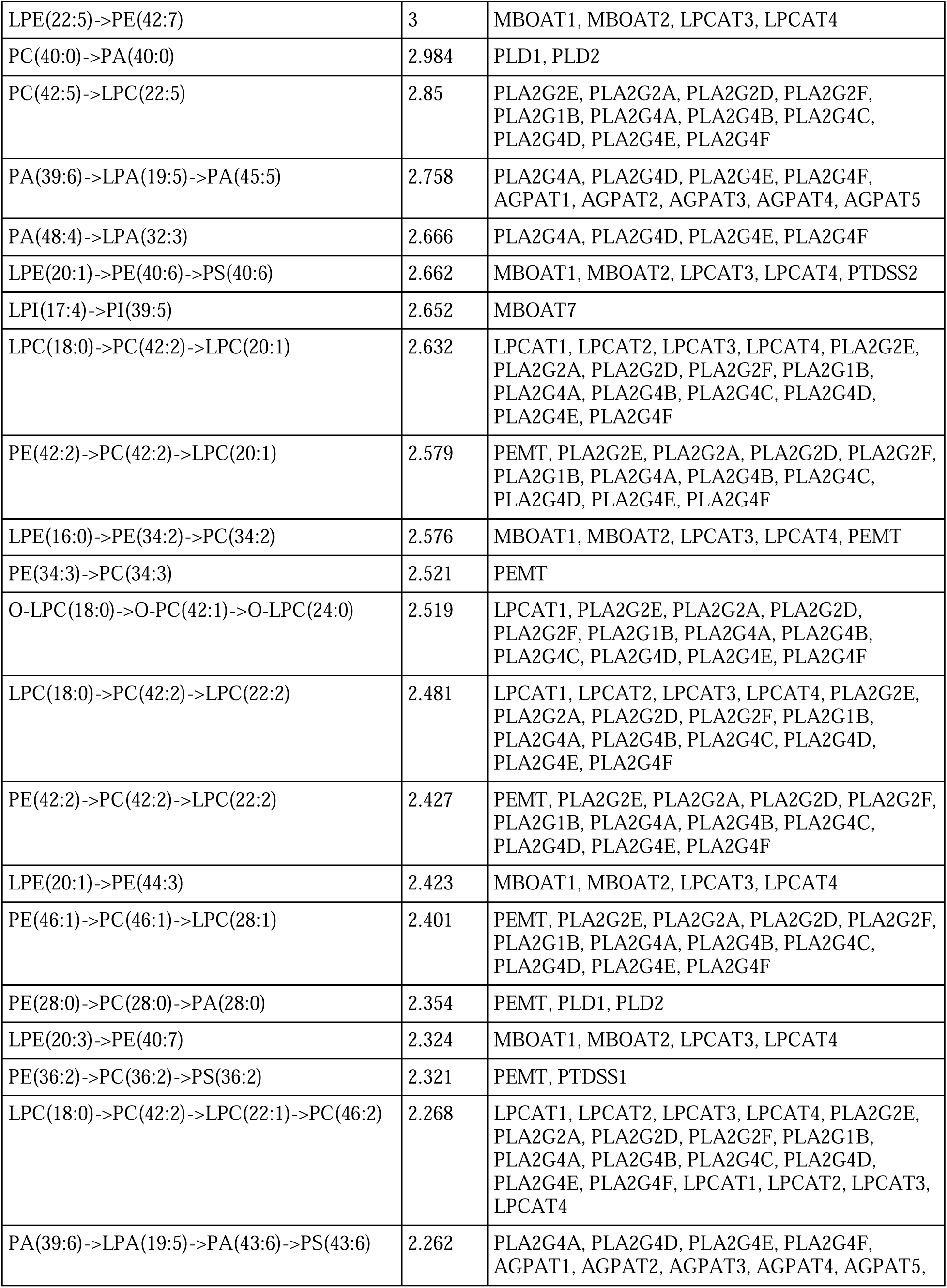
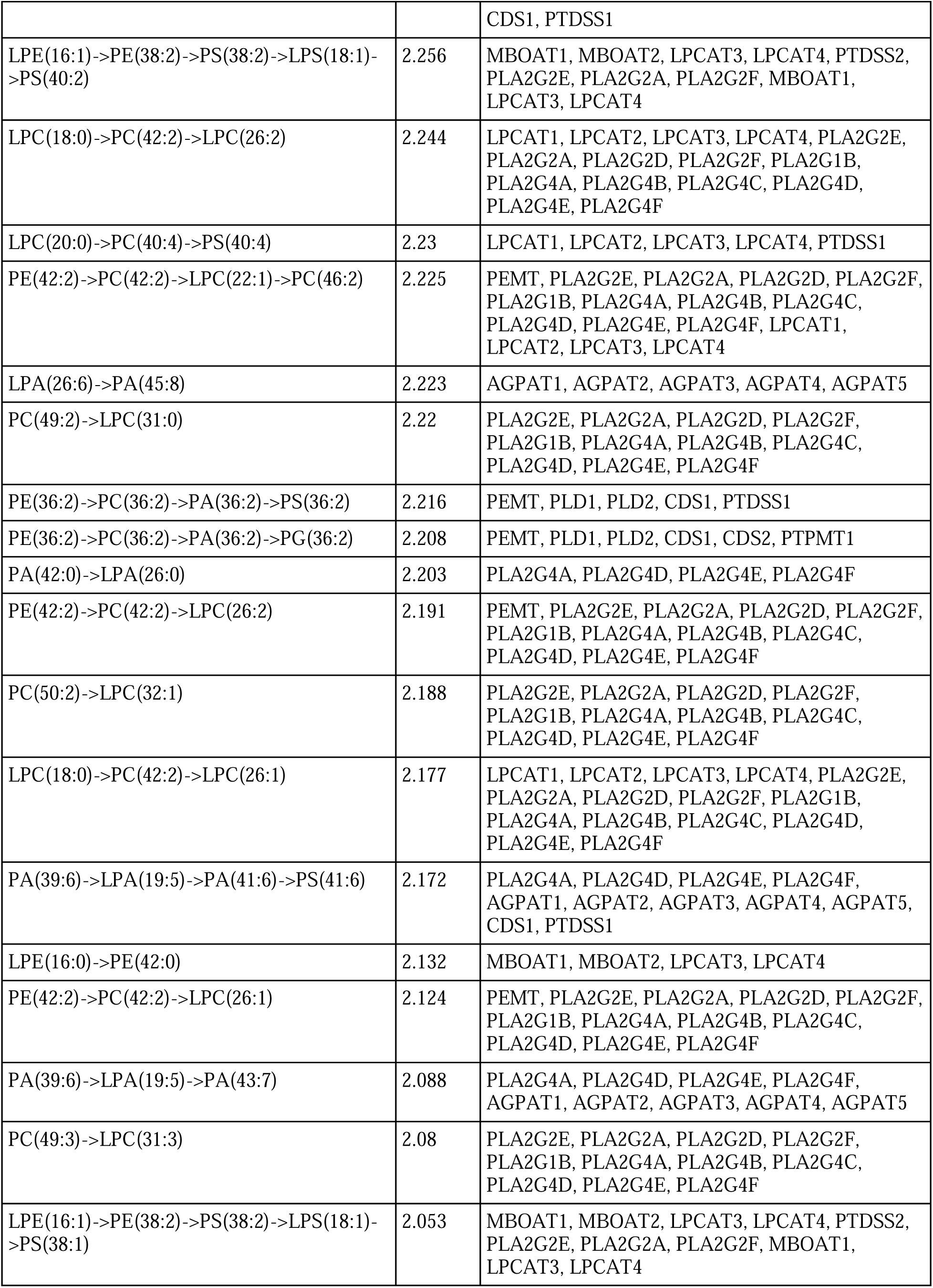
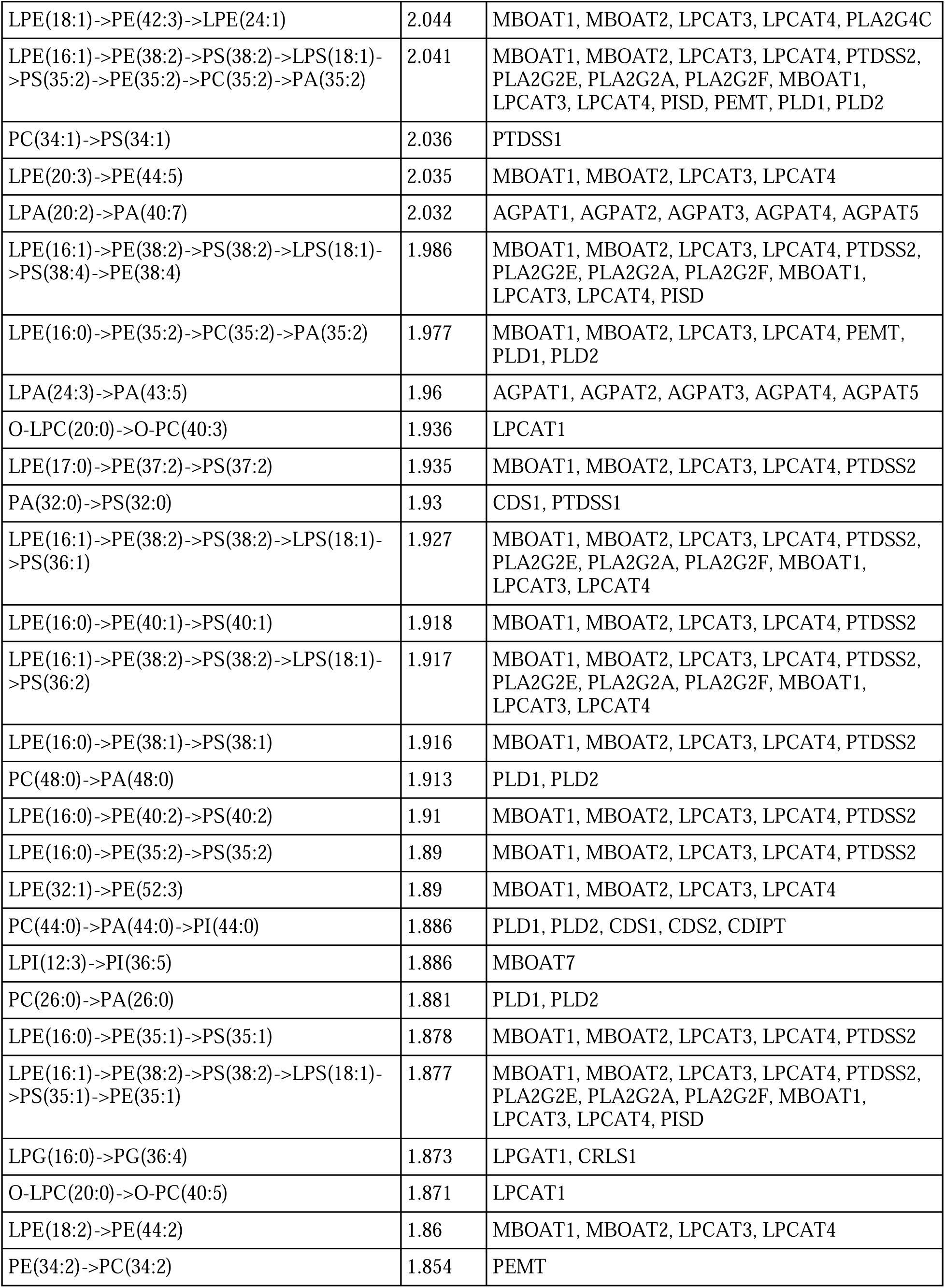
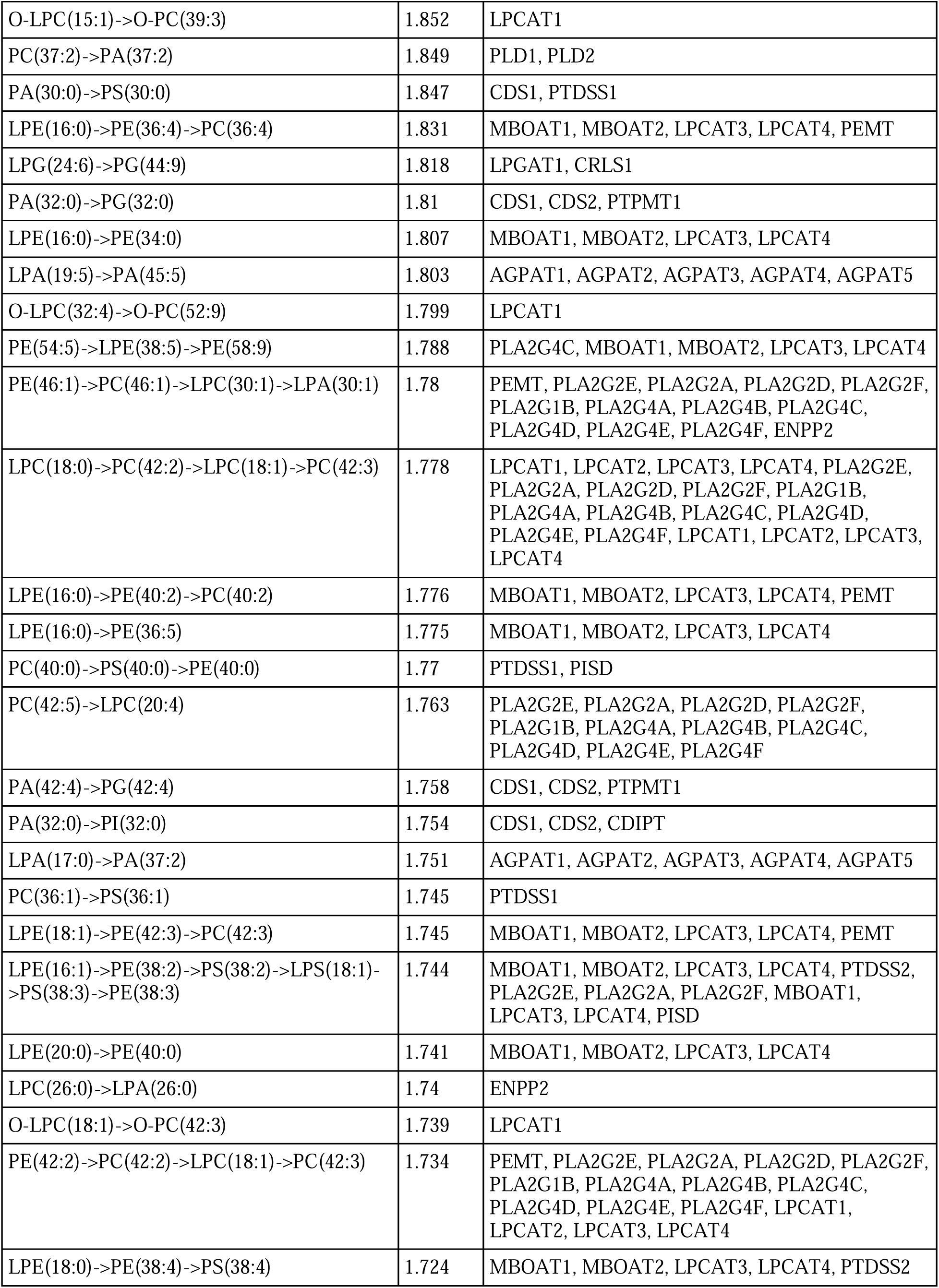
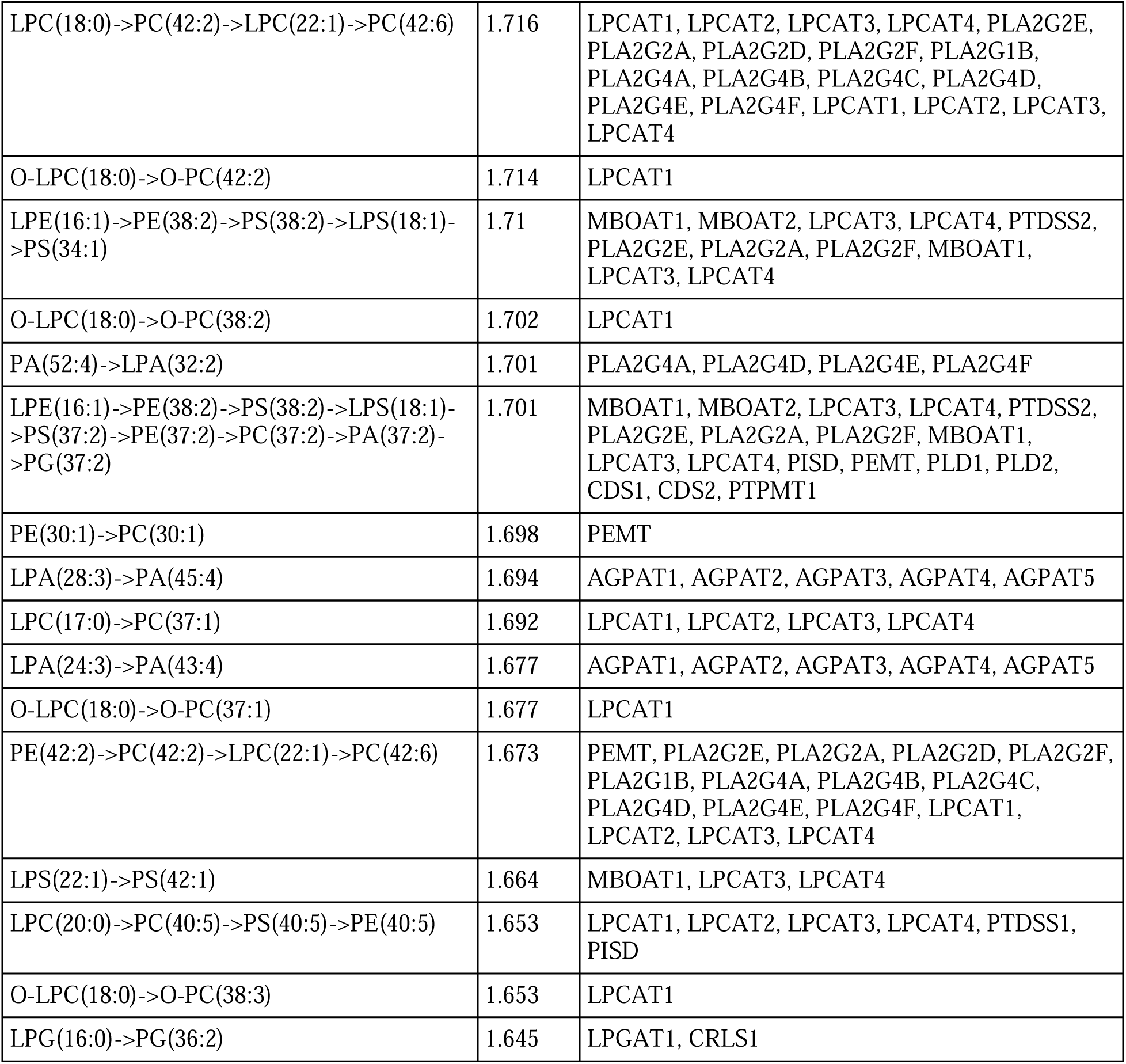
BioPAN analysis of Glycerophospholipids. BioPAN analysis revealed activated and suppressed reactions in LPS treated microglia at the lipid species level.

**Supplemental Table 2.**
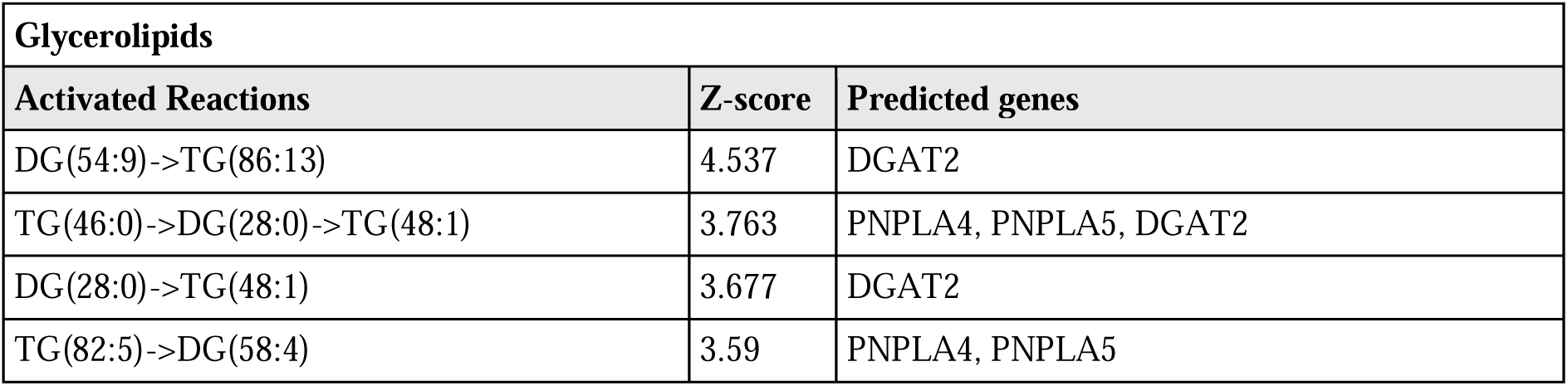
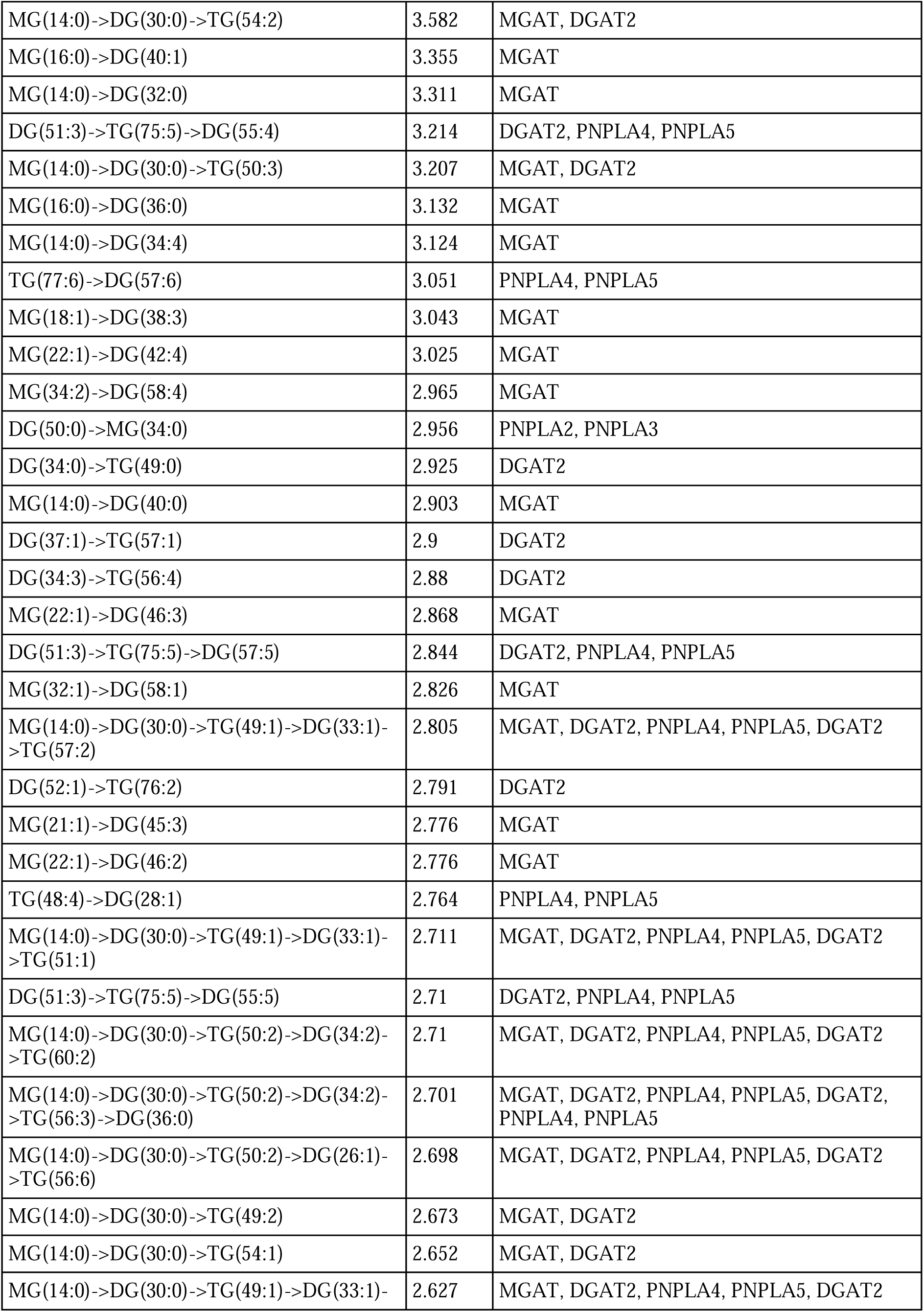
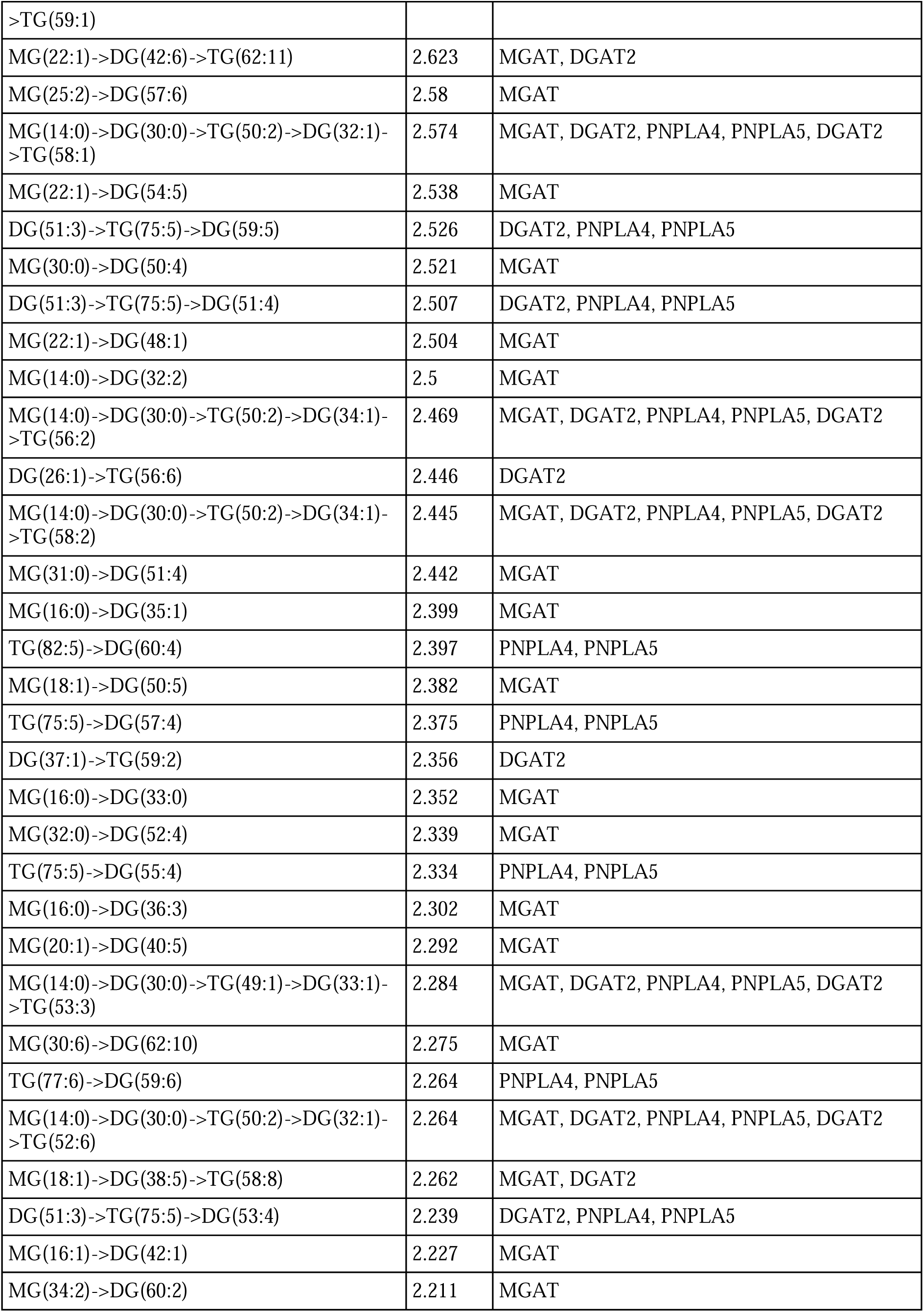
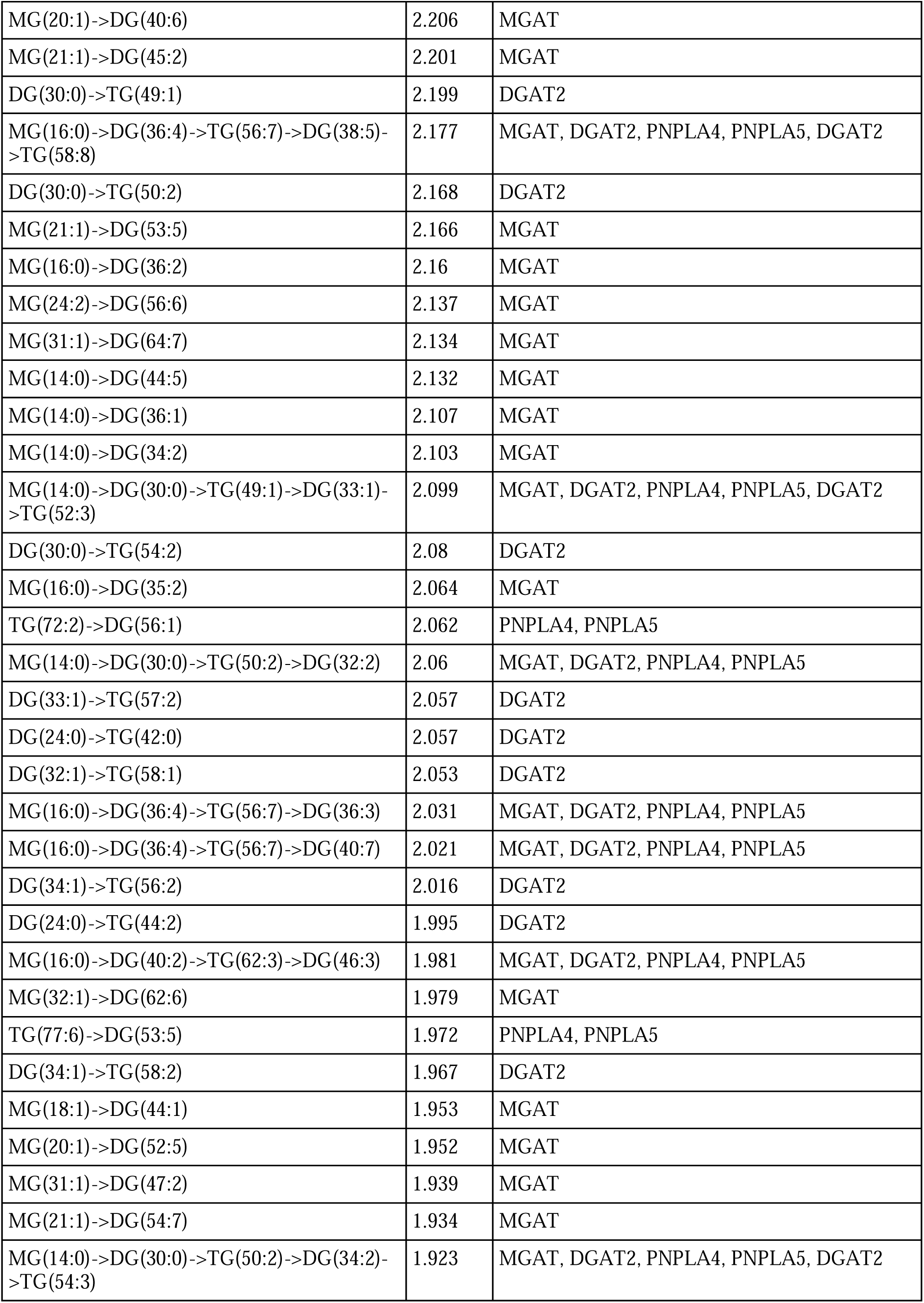
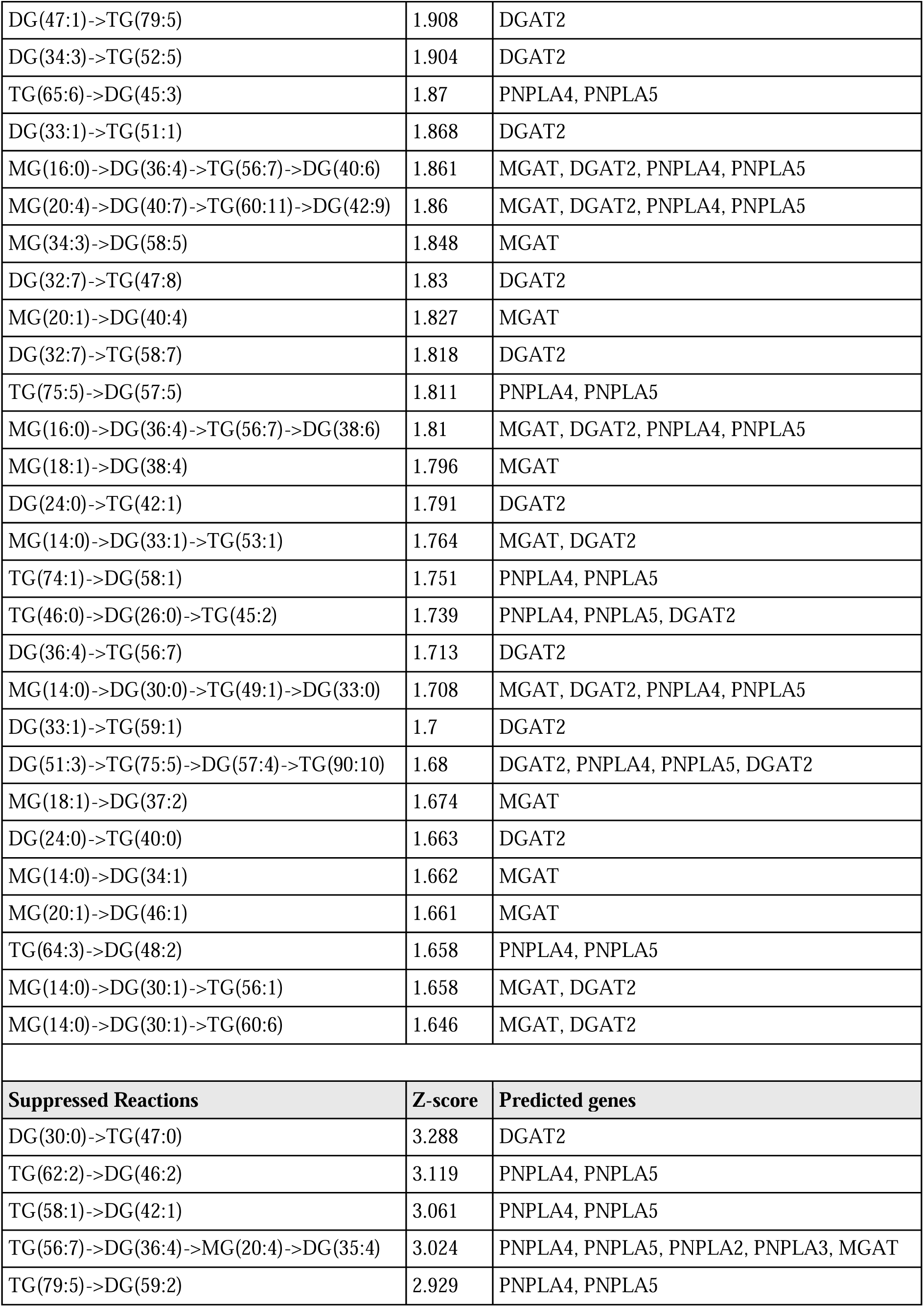
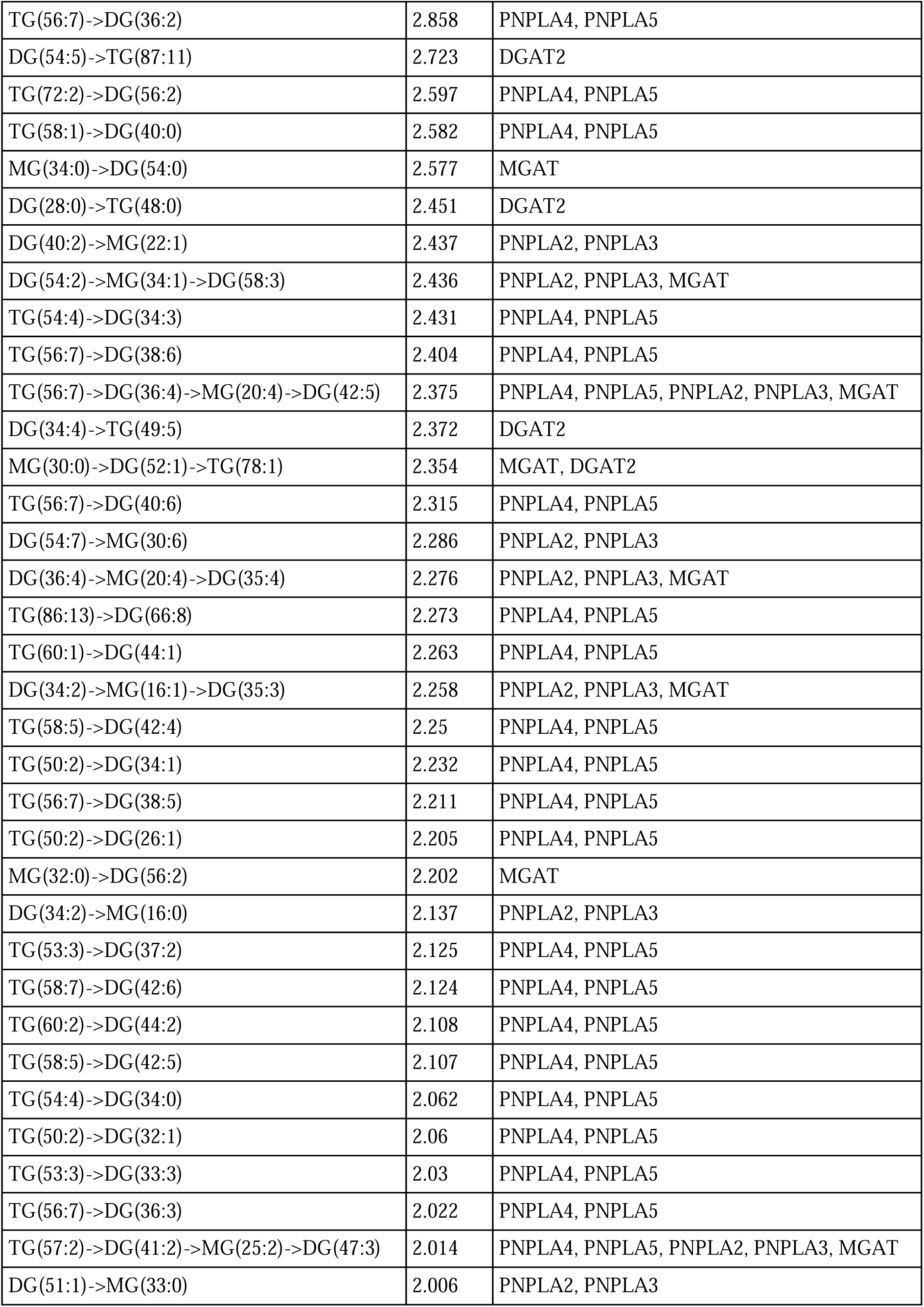
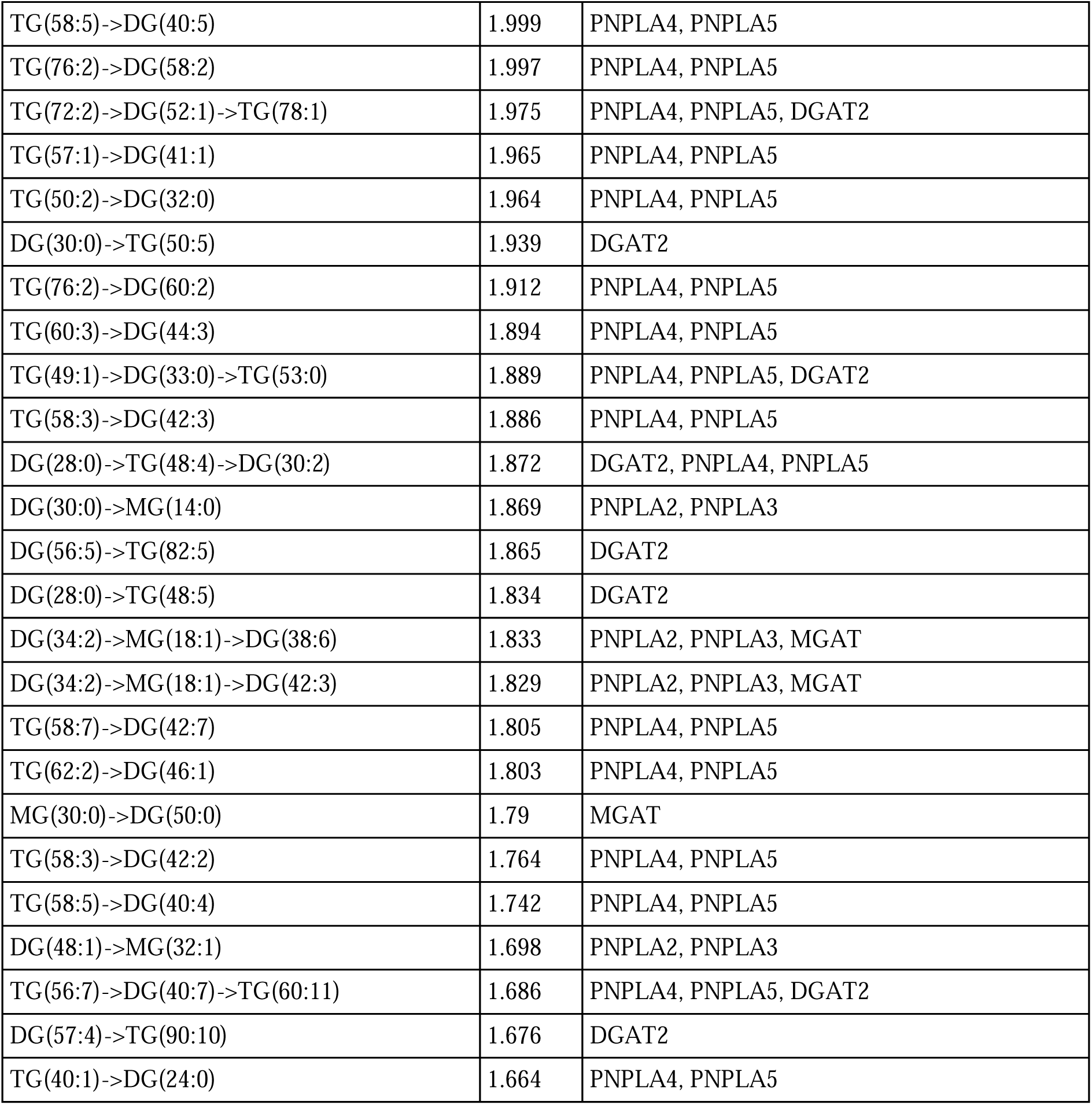
BioPAN analysis of Glycerolipids. BioPAN analysis revealed activated and suppressed reactions in LPS treated microglia at the lipid species level.

**Supplemental Table 3.** NT cells vs LPS cells volcano plot extended statistics. Extended statistics corresponding to Figure 3B in main text. Extended statistics include fold change (FC), log 2 fold change (log2(FC)), the raw p value (raw.pval) and the -log10 of the p-value (- log10(p)).

| Lipid | FC | log2(FC) | raw.pval | -log10(p) |
| --- | --- | --- | --- | --- |
| **TG 48:1** | 24.44 | 4.6111 | 5.0792E-11 | 10.294 |
| **HexCer 45:1;O4** | 0.10565 | -3.2426 | 3.5647E-08 | 7.448 |
| **TG O-8:0_8:0_18:2** | 2.3378 | 1.2251 | 4.8452E-08 | 7.3147 |
| **TG 12:0_12:2_22:7** | 6.524 | 2.7058 | 2.7466E-07 | 6.5612 |
| **Cer 40:1;O6** | 0.36671 | -1.4473 | 2.9644E-07 | 6.5281 |
| **TG 75:0;O3** | 3.0904 | 1.6278 | 1.1756E-06 | 5.9297 |
| **HexCer 42:6;O4** | 2.8107 | 1.4909 | 1.3581E-06 | 5.8671 |
| **DG dO-36:0** | 2.4332 | 1.2828 | 2.3996E-06 | 5.6199 |
| **PE 44:1;O** | 0.57925 | -0.78773 | 5.3329E-06 | 5.273 |
| **PE 44:0** | 0.64975 | -0.62205 | 5.4784E-06 | 5.2613 |
| **HexCer 43:6;O2** | 0.49126 | -1.0254 | 7.0532E-06 | 5.1516 |
| **DG O-14:1_18:0** | 2.2107 | 1.1445 | 1.1919E-05 | 4.9238 |
| **PG 20:4;O** | 2.5054 | 1.325 | 1.3005E-05 | 4.8859 |
| **Cer 41:4;O** | 2.452 | 1.294 | 1.3731E-05 | 4.8623 |
| **PS 37:1** | 6.2131 | 2.6353 | 1.4932E-05 | 4.8259 |
| **HexCer 48:1;O4** | 1.6961 | 0.76222 | 2.4994E-05 | 4.6022 |
| **DG O-16:0_18:1** | 2.7279 | 1.4478 | 2.7349E-05 | 4.5631 |
| **PI 66:2** | 2.9553 | 1.5633 | 2.7409E-05 | 4.5621 |
| **PE 56:5** | 4.9699 | 2.3132 | 2.954E-05 | 4.5296 |
| **PC 68:0** | 5.8519 | 2.5489 | 3.3357E-05 | 4.4768 |
| **Hex2Cer 31:4;O2** | 0.38046 | -1.3942 | 3.4156E-05 | 4.4665 |
| **DG 39:5;O** | 2.5092 | 1.3272 | 3.4747E-05 | 4.4591 |
| **DG 36:0;O2** | 0.42861 | -1.2223 | 3.5841E-05 | 4.4456 |
| **Hex(4)-HexNAc-Fuc-Cer 42:1;O2** | 0.60302 | -0.72972 | 3.8797E-05 | 4.4112 |
| **TG 59:1** | 3.8009 | 1.9263 | 3.9446E-05 | 4.404 |
| **TG 75:5** | 2.2566 | 1.1741 | 4.202E-05 | 4.3765 |
| **TG 12:0_14:0_18:0** | 4.1841 | 2.0649 | 4.3158E-05 | 4.3649 |
| **FOH 10:2** | 1.8766 | 0.90813 | 4.3717E-05 | 4.3593 |
| **DG O-43:3** | 2.8429 | 1.5074 | 4.3927E-05 | 4.3573 |
| **PC O-66:0** | 11.646 | 3.5418 | 4.7996E-05 | 4.3188 |
| **Cer 40:5;O3** | 3.1601 | 1.66 | 5.1987E-05 | 4.2841 |
| **PG 50:2;O** | 2.3218 | 1.2152 | 5.721E-05 | 4.2425 |
| **TG 58:1** | 5.3697 | 2.4248 | 6.1434E-05 | 4.2116 |
| **FA 20:0;O** | 0.43105 | -1.2141 | 6.1679E-05 | 4.2099 |
| **DG 40:0** | 2.9331 | 1.5524 | 6.7574E-05 | 4.1702 |
| **PE O-42:3** | 0.56591 | -0.82135 | 7.1172E-05 | 4.1477 |
| **Cer 17:0;3O/18:0;(2OH)** | 3.7721 | 1.9153 | 7.1313E-05 | 4.1468 |
| **DG 16:0_20:0** | 12.538 | 3.6483 | 7.169E-05 | 4.1445 |
| **DG 14:0_16:0** | 1.9517 | 0.96471 | 7.8941E-05 | 4.1027 |
| **PE 42:2;O** | 2.6774 | 1.4208 | 8.337E-05 | 4.079 |
| **CerP 52:2;O2** | 3.6339 | 1.8615 | 8.4292E-05 | 4.0742 |
| **DG O-37:3** | 2.4494 | 1.2924 | 8.4645E-05 | 4.0724 |
| TG 71:0;O3 | 4.1193 | 2.0424 | 8.977E-05 | 4.0469 |
| TG 51:3;O3 | 3.663 | 1.873 | 9.0679E-05 | 4.0425 |
| TG 60:1 | 3.6463 | 1.8665 | 9.3235E-05 | 4.0304 |
| Cer 38:0;O4 | 2.0219 | 1.0157 | 0.00010159 | 3.9932 |
| PE-NMe2 24:0 | 2.3733 | 1.2469 | 0.00010326 | 3.986 |
| PG O-44:3 | 5.0142 | 2.326 | 0.00010506 | 3.9786 |
| LPG O-32:2 | 0.26962 | -1.891 | 0.00010638 | 3.9732 |
| Cer 18:0;O2/22:0;O | 3.4279 | 1.7773 | 0.0001091 | 3.9622 |
| ACer 71:3;O6 | 2.3275 | 1.2188 | 0.00011042 | 3.9569 |
| LPA O-27:3 | 4.039 | 2.014 | 0.00011395 | 3.9433 |
| TG 57:1 | 5.6202 | 2.4906 | 0.00011768 | 3.9293 |
| PE 58:6 | 2.9701 | 1.5705 | 0.00012381 | 3.9072 |
| PE 34:1 | 1.7379 | 0.79734 | 0.00012458 | 3.9046 |
| PE 42:2 | 2.7406 | 1.4545 | 0.00012581 | 3.9003 |
| MG 33:2;O | 1.9488 | 0.96255 | 0.00014218 | 3.8471 |
| DG O-36:3 | 3.8724 | 1.9532 | 0.00014308 | 3.8444 |
| DG 34:4 | 2.5113 | 1.3284 | 0.0001449 | 3.8389 |
| LPS 33:0;O | 3.1519 | 1.6562 | 0.00014598 | 3.8357 |
| SM 45:5;O2 | 2.3185 | 1.2132 | 0.0001477 | 3.8306 |
| DG 37:0;O2 | 2.6932 | 1.4293 | 0.00016037 | 3.7949 |
| DG 57:6 | 6.2151 | 2.6358 | 0.00016131 | 3.7923 |
| DG 58:4 | 4.2292 | 2.0804 | 0.00016342 | 3.7867 |
| LPA 26:0 | 0.3492 | -1.5179 | 0.00017284 | 3.7623 |
| LPE 22:1 | 2.5664 | 1.3597 | 0.00018766 | 3.7266 |
| DG 33:0;O2 | 3.2662 | 1.7076 | 0.0001927 | 3.7151 |
| FA 30:5 | 3.6134 | 1.8534 | 0.00019496 | 3.7101 |
| TG 49:0 | 3.9026 | 1.9645 | 0.00019978 | 3.6994 |
| TG 14:0_18:0_22:1 | 4.6642 | 2.2216 | 0.00020468 | 3.6889 |
| PI 15:0_18:1 | 2.612 | 1.3852 | 0.0002075 | 3.683 |
| Cer 18:0;3O/19:0;(2OH) | 4.8763 | 2.2858 | 0.00021737 | 3.6628 |
| DG 38:5;O | 2.456 | 1.2963 | 0.0002527 | 3.5974 |
| FAHFA 35:1;O | 2.171 | 1.1184 | 0.00028671 | 3.5426 |
| BMP 19:4;O | 2.4485 | 1.2919 | 0.00029433 | 3.5312 |
| DG O-40:0 | 2.2614 | 1.1772 | 0.00029815 | 3.5256 |
| TG O-18:3_8:0_12:0 | 4.8821 | 2.2875 | 0.0003008 | 3.5217 |
| LPE 22:5 | 2.5011 | 1.3226 | 0.00030543 | 3.5151 |
| PC O-18:1_16:0 | 1.9902 | 0.99293 | 0.00031374 | 3.5034 |
| PE O-18:1_21:2 | 0.53073 | -0.91394 | 0.00031943 | 3.4956 |
| PA 32:0 | 2.796 | 1.4833 | 0.00032021 | 3.4946 |
| TG 55:1 | 3.7455 | 1.9051 | 0.00032152 | 3.4928 |
| MG O-30:0;O | 1.4282 | 0.51424 | 0.00033443 | 3.4757 |
| TG 14:0_18:0_20:1 | 8.4639 | 3.0813 | 0.00036181 | 3.4415 |
| PC 30:0;O | 2.4117 | 1.2701 | 0.00036978 | 3.4321 |
| TG 84:11;O3 | 2.7757 | 1.4728 | 0.0003842 | 3.4154 |
| DG 15:0_18:0 | 3.0763 | 1.6212 | 0.00038501 | 3.4145 |
| SHexCer 38:1;O2 | 0.49042 | -1.0279 | 0.00038764 | 3.4116 |
| HexCer 43:2;O4 | 0.3853 | -1.3759 | 0.00040348 | 3.3942 |
| Cer 41:5;O2 | 2.6021 | 1.3797 | 0.00042337 | 3.3733 |
| PC 12:0_14:0 | 2.4645 | 1.3013 | 0.00045648 | 3.3406 |
| DG 55:4 | 5.2186 | 2.3837 | 0.00045691 | 3.3402 |
| TG 96:8;O | 1.9808 | 0.98608 | 0.00046132 | 3.336 |
| MG O-32:5 | 2.8368 | 1.5043 | 0.00046734 | 3.3304 |
| TG O-14:0_18:0_18:1 | 2.6528 | 1.4075 | 0.00046751 | 3.3302 |
| TG 60:5;O2 | 3.7282 | 1.8985 | 0.00046815 | 3.3296 |
| Cer 51:1;O4 | 1.5153 | 0.5996 | 0.00049494 | 3.3054 |
| SHexCer 38:1;O3 | 0.55207 | -0.85709 | 0.00050182 | 3.2995 |
| DG 26:1 | 0.48897 | -1.0322 | 0.00053588 | 3.2709 |
| PE-Cer 18:2;O2/21:1;O | 0.45732 | -1.1287 | 0.0005379 | 3.2693 |
| PG 40:4 | 0.39207 | -1.3508 | 0.00053879 | 3.2686 |
| DG O-38:0 | 1.5726 | 0.65318 | 0.00054028 | 3.2674 |
| MG 34:2;O | 1.5809 | 0.66073 | 0.0005565 | 3.2545 |
| CE 32:3;O | 1.6657 | 0.73616 | 0.00056006 | 3.2518 |
| PC 10:0_16:0 | 2.5244 | 1.336 | 0.00057135 | 3.2431 |
| TG 99:5 | 5.1534 | 2.3655 | 0.0005742 | 3.2409 |
| TG O-42:1 | 1.9678 | 0.97657 | 0.00057598 | 3.2396 |
| DG 12:0_16:0 | 2.8001 | 1.4855 | 0.00058011 | 3.2365 |
| DG 40:5;O | 2.4834 | 1.3123 | 0.00059777 | 3.2235 |
| TG 46:3 | 2.1411 | 1.0983 | 0.00059829 | 3.2231 |
| Cer 50:2;O3 | 1.5418 | 0.62458 | 0.00061979 | 3.2078 |
| HexCer 52:4;O3 | 3.9016 | 1.9641 | 0.00063844 | 3.1949 |
| FA 19:0;O2 | 2.4467 | 1.2909 | 0.00065113 | 3.1863 |
| PE O-40:3 | 0.51683 | -0.95223 | 0.00066276 | 3.1786 |
| NA 18:0;O | 0.28148 | -1.8289 | 0.00066432 | 3.1776 |
| TG 16:1_18:1_18:1 | 5.725 | 2.5173 | 0.0006731 | 3.1719 |
| MG O-32:0;O | 3.7027 | 1.8886 | 0.00070429 | 3.1522 |
| DG 53:3 | 3.6718 | 1.8765 | 0.0007052 | 3.1517 |
| PC 16:0_20:0 | 2.1748 | 1.1209 | 0.00072584 | 3.1392 |
| Cer 42:1;O3 | 0.59463 | -0.74993 | 0.00075369 | 3.1228 |
| PG 42:4 | 0.42937 | -1.2197 | 0.00075502 | 3.122 |
| TG 14:0_18:0_19:1 | 5.9296 | 2.5679 | 0.00077231 | 3.1122 |
| PE 40:6 | 0.45276 | -1.1432 | 0.00077799 | 3.109 |
| LPE 20:4 | 2.5254 | 1.3365 | 0.00078491 | 3.1052 |
| PA 26:0 | 0.27297 | -1.8732 | 0.00079701 | 3.0985 |
| PG 40:1 | 0.38506 | -1.3768 | 0.00080319 | 3.0952 |
| DG 41:3 | 0.41723 | -1.2611 | 0.00080852 | 3.0923 |
| MGDG 17:0_18:2 | 0.65489 | -0.61067 | 0.00081995 | 3.0862 |
| Cer 16:0;O2/18:0;O | 2.8217 | 1.4965 | 0.00084176 | 3.0748 |
| NAT 32:0;O | 1.9214 | 0.94217 | 0.00085676 | 3.0671 |
| DG 28:1 | 4.4854 | 2.1652 | 0.00087649 | 3.0573 |
| HBMP 60:2; HBMP 20:0-22:1-18:1; [M+NH4]<sup>+</sup> | 0.1451 | -2.7849 | 0.00088191 | 3.0546 |
| PIP3 37:6 | 0.61951 | -0.69079 | 0.00089314 | 3.0491 |
| HexCer 32:5;O5 | 0.565 | -0.82368 | 0.00089682 | 3.0473 |
| TG 79:0;O3 | 2.9578 | 1.5645 | 0.00091049 | 3.0407 |
| DG O-35:3 | 2.3635 | 1.2409 | 0.00091579 | 3.0382 |
| TG O-83:15 | 1.5208 | 0.6048 | 0.00092905 | 3.032 |
| PI 31:4;O | 0.53702 | -0.89695 | 0.00093088 | 3.0311 |
| HexCer 45:2;O5 | 0.68277 | -0.55053 | 0.00094679 | 3.0237 |
| PI O-67:3 | 2.6399 | 1.4005 | 0.00095413 | 3.0204 |
| DG 40:7;O | 2.0206 | 1.0148 | 0.00096607 | 3.015 |
| HexCer 54:5;O3 | 2.6576 | 1.4101 | 0.0010078 | 2.9966 |
| Cer 41:4;O2 | 1.879 | 0.90998 | 0.0010484 | 2.9795 |
| DG 40:5 | 1.7495 | 0.80696 | 0.0010809 | 2.9662 |
| TG 52:7;O2 | 0.31823 | -1.6519 | 0.0011095 | 2.9549 |
| LPS 18:1 | 0.59248 | -0.75516 | 0.0011105 | 2.9545 |
| DG 38:4 | 2.2454 | 1.167 | 0.0011236 | 2.9494 |
| DG 38:6;O | 2.57 | 1.3618 | 0.0011392 | 2.9434 |
| PA 44:4 | 2.1907 | 1.1314 | 0.0011583 | 2.9362 |
| SM 43:5;O2 | 1.6902 | 0.7572 | 0.0011865 | 2.9257 |
| TG 12:2_12:2_22:6 | 2.2487 | 1.1691 | 0.0011909 | 2.9241 |
| TG 16:0_18:1_18:1 | 3.9146 | 1.9689 | 0.001191 | 2.9241 |
| PA O-54:2 | 2.1213 | 1.0849 | 0.0011959 | 2.9223 |
| TG O-45:2 | 0.29715 | -1.7507 | 0.0012006 | 2.9206 |
| Hex2Cer 40:6;O2 | 0.62608 | -0.67558 | 0.0012035 | 2.9196 |
| TG 86:13 | 1.7398 | 0.79894 | 0.0012424 | 2.9057 |
| TG 56:4 | 1.9454 | 0.96005 | 0.0012466 | 2.9043 |
| PE 35:0 | 1.5428 | 0.62553 | 0.0012636 | 2.8984 |
| TG O-75:0;O3 | 1.9889 | 0.99196 | 0.0012658 | 2.8976 |
| PE-Cer 32:1;O;P | 1.8273 | 0.8697 | 0.0012664 | 2.8974 |
| Hex(5)-HexNAc(3)-Cer 34:1;O2 | 1.5086 | 0.59321 | 0.0012751 | 2.8945 |
| TG 53:1 | 3.219 | 1.6866 | 0.001281 | 2.8924 |
| DG 26:0 | 3.6757 | 1.878 | 0.0012822 | 2.892 |
| PG O-39:0 | 1.6909 | 0.75779 | 0.0012871 | 2.8904 |
| TG 97:6;O2 | 2.3191 | 1.2136 | 0.0012885 | 2.8899 |
| DG dO-40:2 | 2.7801 | 1.4751 | 0.0012938 | 2.8881 |
| DG 50:2 | 3.5189 | 1.8151 | 0.0013111 | 2.8824 |
| TG 56:1 | 9.2115 | 3.2034 | 0.0013114 | 2.8823 |
| HexCer 48:2;O4 | 1.6794 | 0.74796 | 0.001352 | 2.869 |
| PG 58:4;O | 0.26342 | -1.9246 | 0.0013546 | 2.8682 |
| TG 14:0_18:0_18:1 | 10.605 | 3.4067 | 0.0013747 | 2.8618 |
| PC 36:0 | 2.1311 | 1.0916 | 0.0014019 | 2.8533 |
| SM 44:4;O4 | 3.2513 | 1.701 | 0.0014107 | 2.8506 |
| Cer 37:6;O2 | 2.814 | 1.4926 | 0.0014715 | 2.8323 |
| DG 30:2 | 1.5694 | 0.65026 | 0.0015051 | 2.8224 |
| MG 30:0 | 1.897 | 0.92372 | 0.0015882 | 2.7991 |
| TG O-61:0;O3 | 0.4159 | -1.2657 | 0.0015899 | 2.7986 |
| TG 54:2 | 3.3383 | 1.7391 | 0.0015924 | 2.798 |
| LPG O-32:0 | 0.24837 | -2.0095 | 0.0016068 | 2.794 |
| LPS 32:0;O | 0.46045 | -1.1189 | 0.0016187 | 2.7908 |
| TG 96:18 | 2.4307 | 1.2813 | 0.0016197 | 2.7906 |
| MG 34:0 | 3.0996 | 1.6321 | 0.0016436 | 2.7842 |
| PI 64:1 | 2.416 | 1.2726 | 0.0016439 | 2.7841 |
| HexCer 39:3;O3 | 2.1564 | 1.1086 | 0.0016473 | 2.7832 |
| MG 32:0 | 1.6736 | 0.743 | 0.0016965 | 2.7705 |
| HexCer 40:4;O4 | 2.2266 | 1.1548 | 0.001721 | 2.7642 |
| HexCer 53:5;O6 | 0.29844 | -1.7445 | 0.0017331 | 2.7612 |
| TG 12:1_12:3_18:5 | 1.8711 | 0.90389 | 0.0017512 | 2.7567 |
| PG 52:2;O | 1.8945 | 0.92178 | 0.0018291 | 2.7378 |
| DG 39:6;O | 1.6647 | 0.73524 | 0.0018498 | 2.7329 |
| HexCer 41:4;O2 | 2.0556 | 1.0396 | 0.0018601 | 2.7305 |
| TG 98:9;O2 | 2.2982 | 1.2005 | 0.0019077 | 2.7195 |
| PE 54:2;O | 2.7264 | 1.447 | 0.0019429 | 2.7115 |
| DG O-39:0 | 2.517 | 1.3317 | 0.0020039 | 2.6981 |
| Cer 40:2;O3 | 2.1981 | 1.1363 | 0.0020063 | 2.6976 |
| DG O-16:0_18:0 | 3.1244 | 1.6436 | 0.0020575 | 2.6867 |
| LPA O-27:6 | 3.4164 | 1.7725 | 0.0020622 | 2.6857 |
| HexCer 45:4;O4 | 0.51293 | -0.96317 | 0.0021201 | 2.6736 |
| DG 41:6;O | 1.6464 | 0.7193 | 0.0021365 | 2.6703 |
| LPC 20:1 | 0.4879 | -1.0353 | 0.0021372 | 2.6702 |
| DG 18:1_34:1 | 2.0067 | 1.0048 | 0.0021869 | 2.6602 |
| PI O-53:3 | 2.1571 | 1.1091 | 0.0021894 | 2.6597 |
| HBMP 16:1_18:0_18:0 | 2.3672 | 1.2432 | 0.002194 | 2.6588 |
| Cer 40:6;O3 | 2.4487 | 1.292 | 0.0022269 | 2.6523 |
| PE 38:5 | 1.6339 | 0.70835 | 0.0022437 | 2.649 |
| PE O-42:4 | 0.53367 | -0.90599 | 0.0022442 | 2.6489 |
| HexCer 41:0;O5 | 2.1648 | 1.1142 | 0.002248 | 2.6482 |
| TG 56:5 | 3.1908 | 1.6739 | 0.0022983 | 2.6386 |
| PA 55:0 | 0.40788 | -1.2938 | 0.0023127 | 2.6359 |
| TG 98:18 | 1.8825 | 0.91266 | 0.0023204 | 2.6344 |
| PE 30:1 | 0.49919 | -1.0023 | 0.0023577 | 2.6275 |
| PE 12:0_16:0 | 1.7571 | 0.8132 | 0.0023974 | 2.6203 |
| AHexCer (O-17:3)18:1;2O/20:2;O | 3.0137 | 1.5916 | 0.0024163 | 2.6168 |
| TG 84:5;O3 | 0.25018 | -1.999 | 0.0024732 | 2.6067 |
| Cer 42:6;O3 | 1.7607 | 0.81614 | 0.002516 | 2.5993 |
| TG 56:6 | 4.6844 | 2.2279 | 0.002555 | 2.5926 |
| TG 83:0;O3 | 2.4007 | 1.2635 | 0.0025843 | 2.5877 |
| DG O-43:4 | 1.4075 | 0.4931 | 0.002592 | 2.5864 |
| LPC O-22:0 | 0.40252 | -1.3129 | 0.0026062 | 2.584 |
| Car 34:5;O | 2.6332 | 1.3968 | 0.0026244 | 2.581 |
| DG 16:0_32:2 | 2.6337 | 1.3971 | 0.0026452 | 2.5775 |
| HexCer 38:0;O6 | 0.69575 | -0.52335 | 0.0027062 | 2.5676 |
| PC 30:1;O | 1.6704 | 0.74016 | 0.0027372 | 2.5627 |
| DG 60:5 | 2.5814 | 1.3682 | 0.0028175 | 2.5501 |
| PA 45:7 | 0.34176 | -1.5489 | 0.0028181 | 2.55 |
| SL 10:0;O/14:1;O | 2.8515 | 1.5117 | 0.0028549 | 2.5444 |
| PI O-18:4_16:1 | 0.64051 | -0.6427 | 0.0028556 | 2.5443 |
| PA O-39:4 | 0.67827 | -0.56007 | 0.0028897 | 2.5392 |
| TG 82:4;O3 | 0.46063 | -1.1183 | 0.0028905 | 2.539 |
| DG 39:2 | 2.4668 | 1.3026 | 0.0029065 | 2.5366 |
| DG 16:0_20:3 | 2.2107 | 1.1445 | 0.0029171 | 2.5351 |
| TG 12:3_12:3_18:4 | 2.5079 | 1.3265 | 0.0029288 | 2.5333 |
| DG 31:2 | 2.1723 | 1.1192 | 0.002932 | 2.5328 |
| DG O-40:6 | 2.3082 | 1.2067 | 0.0029525 | 2.5298 |
| Cer 39:6;O2 | 2.103 | 1.0724 | 0.0029572 | 2.5291 |
| PE 56:5;O | 2.4509 | 1.2933 | 0.0030023 | 2.5225 |
| MG 21:1 | 0.56688 | -0.81888 | 0.0030177 | 2.5203 |
| HexCer 40:6;O4 | 2.2865 | 1.1931 | 0.0030407 | 2.517 |
| FA 17:0 | 0.62559 | -0.6767 | 0.0030411 | 2.517 |
| PA O-45:1 | 0.63534 | -0.6544 | 0.0030448 | 2.5164 |
| CerP 54:6;O5 | 2.2575 | 1.1747 | 0.0031031 | 2.5082 |
| DG 57:4 | 7.5881 | 2.9237 | 0.0031043 | 2.508 |
| DG 35:1;O2 | 2.3517 | 1.2337 | 0.003137 | 2.5035 |
| PC O-22:3_26:0 | 2.4199 | 1.2749 | 0.0031588 | 2.5005 |
| Cer 49:1;O4 | 1.8335 | 0.87462 | 0.0031851 | 2.4969 |
| SM 18:1;O2/26:2 | 0.50648 | -0.98142 | 0.0031924 | 2.4959 |
| HexCer 35:2;O2 | 1.4067 | 0.49228 | 0.0032402 | 2.4894 |
| Cer 38:0;O6 | 1.5608 | 0.64229 | 0.0033067 | 2.4806 |
| PI 14:0_18:2 | 0.48332 | -1.0489 | 0.0033357 | 2.4768 |
| DG 40:1 | 1.9556 | 0.96764 | 0.003379 | 2.4712 |
| DG O-41:4 | 1.8458 | 0.88425 | 0.0034205 | 2.4659 |
| Car 30:1 | 0.52302 | -0.93507 | 0.0035484 | 2.45 |
| DG 42:1 | 1.8616 | 0.89651 | 0.0035661 | 2.4478 |
| DG 52:4 | 3.5718 | 1.8366 | 0.0035699 | 2.4473 |
| FA 33:3;O2 | 2.986 | 1.5782 | 0.0035712 | 2.4472 |
| PC 18:0_20:0 | 1.6286 | 0.70366 | 0.0036253 | 2.4407 |
| TG 48:2 | 7.3938 | 2.8863 | 0.0036723 | 2.4351 |
| PC 15:0_16:2 | 2.3524 | 1.2341 | 0.0036833 | 2.4338 |
| PC 33:3;O | 3.055 | 1.6112 | 0.0036931 | 2.4326 |
| TG 14:0_16:0_19:1 | 6.4056 | 2.6793 | 0.0037676 | 2.4239 |
| DG 40:6;O | 1.6628 | 0.73365 | 0.0037799 | 2.4225 |
| Cer 39:0;O3 | 1.9202 | 0.94123 | 0.0037819 | 2.4223 |
| DG 36:4 | 1.7791 | 0.83111 | 0.0038076 | 2.4194 |
| LPE 20:0 | 1.677 | 0.74584 | 0.0038759 | 2.4116 |
| CE 20:5 | 0.62031 | -0.68893 | 0.003915 | 2.4073 |
| LPA 26:6 | 1.5742 | 0.6546 | 0.0039167 | 2.4071 |
| PG 14:0_18:0 | 3.2299 | 1.6915 | 0.0039519 | 2.4032 |
| SM 16:0;O2/18:0 | 1.6759 | 0.74495 | 0.003978 | 2.4003 |
| PI-Cer 40:5;O6 | 2.167 | 1.1157 | 0.004013 | 2.3965 |
| DG O-38:4 | 1.6076 | 0.68493 | 0.0040607 | 2.3914 |
| DG 53:4 | 2.8961 | 1.5341 | 0.0040846 | 2.3889 |
| HexCer 18:1;O2/24:1 | 0.49302 | -1.0203 | 0.00415 | 2.382 |
| TG 54:5 | 3.7345 | 1.9009 | 0.0042644 | 2.3701 |
| Cer 44:6;O | 0.29858 | -1.7438 | 0.0042672 | 2.3699 |
| ST 28:3;O4 | 0.34492 | -1.5357 | 0.004297 | 2.3668 |
| PE O-18:0_20:3 | 0.43442 | -1.2028 | 0.0043204 | 2.3645 |
| MG O-18:0 | 2.8279 | 1.4997 | 0.0043645 | 2.3601 |
| DG 26:0_18:1 | 2.3769 | 1.2491 | 0.0043669 | 2.3598 |
| DG 56:8 | 3.391 | 1.7617 | 0.0043762 | 2.3589 |
| PIP2 62:4 | 0.51398 | -0.96022 | 0.0043854 | 2.358 |
| PA O-58:6 | 3.5171 | 1.8144 | 0.004415 | 2.3551 |
| Cer 40:0;O6 | 0.58827 | -0.76546 | 0.0044222 | 2.3544 |
| DG O-44:8 | 2.3536 | 1.2348 | 0.0044443 | 2.3522 |
| Car 22:3;O2 | 0.48801 | -1.035 | 0.0044823 | 2.3485 |
| TG 70:12;O2 | 0.16551 | -2.595 | 0.0045722 | 2.3399 |
| DG 54:3 | 1.5443 | 0.62698 | 0.0046078 | 2.3365 |
| PA 45:6 | 0.42766 | -1.2255 | 0.0046165 | 2.3357 |
| MG 32:1 | 0.56468 | -0.82449 | 0.0046189 | 2.3355 |
| DG 51:4 | 3.1818 | 1.6699 | 0.004623 | 2.3351 |
| HexCer 45:2;O4 | 0.40774 | -1.2943 | 0.0047063 | 2.3273 |
| TG 44:6;O2 | 5.322 | 2.412 | 0.0047434 | 2.3239 |
| PE 22:5 | 97.134 | 6.6019 | 0.0047464 | 2.3236 |
| TG 60:6 | 2.2986 | 1.2008 | 0.0048956 | 2.3102 |
| PI O-38:5 | 0.49051 | -1.0277 | 0.0049016 | 2.3097 |
| DG 18:0_22:6 | 2.1163 | 1.0816 | 0.0049352 | 2.3067 |
| Cer 43:1;O3 | 0.41653 | -1.2635 | 0.0049641 | 2.3042 |
| PG 48:2;O | 2.2364 | 1.1611 | 0.0049904 | 2.3019 |
| Hex2Cer 43:0;O2 | 1.4061 | 0.49167 | 0.0050307 | 2.2984 |
| TG 100:10;O2 | 2.0203 | 1.0145 | 0.0050532 | 2.2964 |
| PI 14:0_16:0 | 2.3699 | 1.2448 | 0.0051599 | 2.2874 |
| Cer 41:6;O3 | 1.5375 | 0.62054 | 0.0051803 | 2.2856 |
| PE 34:3 | 3.8076 | 1.9289 | 0.0052109 | 2.2831 |
| TG O-97:5 | 2.9327 | 1.5522 | 0.0052449 | 2.2803 |
| PA O-35:1 | 0.42123 | -1.2473 | 0.0052541 | 2.2795 |
| DG 44:8;O | 1.6102 | 0.68726 | 0.0052877 | 2.2767 |
| HexCer 35:2;O5 | 5.4917 | 2.4573 | 0.0052937 | 2.2762 |
| PC O-38:2 | 0.68314 | -0.54975 | 0.0053758 | 2.2696 |
| Cer 44:0;O2 | 0.67449 | -0.56813 | 0.0054176 | 2.2662 |
| TG 15:0_16:0_17:1 | 10.686 | 3.4176 | 0.0055967 | 2.2521 |
| DG 50:4 | 7.1644 | 2.8408 | 0.0056392 | 2.2488 |
| TG 84:6;O2 | 1.6034 | 0.68111 | 0.0056563 | 2.2475 |
| TG 62:10 | 2.1101 | 1.0773 | 0.00566 | 2.2472 |
| Cer 42:1;O4 | 1.8243 | 0.86738 | 0.0057237 | 2.2423 |
| TG 100:17 | 0.38396 | -1.381 | 0.0057393 | 2.2411 |
| SM 48:1;3O | 1.891 | 0.91916 | 0.0057878 | 2.2375 |
| PE 16:1_20:1 | 5.1805 | 2.3731 | 0.0058664 | 2.2316 |
| TG 18:1_19:0_21:4 | 5.8525 | 2.549 | 0.0058864 | 2.2302 |
| DG 28:4;O2 | 1.7037 | 0.76863 | 0.0059564 | 2.225 |
| FA 17:0;O2 | 1.9061 | 0.9306 | 0.0059918 | 2.2224 |
| TG 14:0_18:0_18:2 | 6.1336 | 2.6167 | 0.0060193 | 2.2205 |
| PI O-40:6 | 0.59266 | -0.75473 | 0.0060221 | 2.2203 |
| SM 40:1;O6 | 2.8542 | 1.5131 | 0.0060923 | 2.2152 |
| HexCer 28:0;O4 | 1.9813 | 0.98648 | 0.0061514 | 2.211 |
| LPI 18:1 | 0.54395 | -0.87846 | 0.0061652 | 2.2101 |
| SPBP 16:1;O3 | 2.0108 | 1.0077 | 0.0062184 | 2.2063 |
| TG 54:2;O2 | 23.303 | 4.5425 | 0.0062263 | 2.2058 |
| TG 79:0;O2 | 4.0669 | 2.0239 | 0.0063483 | 2.1973 |
| HexCer 33:2;O6 | 0.30073 | -1.7335 | 0.0063518 | 2.1971 |
| HexCer 44:2;O5 | 0.44794 | -1.1586 | 0.0064044 | 2.1935 |
| DG 56:1 | 2.6509 | 1.4065 | 0.0064446 | 2.1908 |
| PC 12:0_16:1 | 1.8024 | 0.84996 | 0.0064577 | 2.1899 |
| DG 40:1;O2 | 1.6894 | 0.75647 | 0.0064589 | 2.1898 |
| SHexCer 34:1;O3 | 0.69048 | -0.53433 | 0.0065913 | 2.181 |
| WE 24:1 | 0.40238 | -1.3134 | 0.0066036 | 2.1802 |
| Cer 17:0;3O/18:1;(2OH) | 2.4272 | 1.2793 | 0.0066406 | 2.1778 |
| DG 32:0 | 2.662 | 1.4125 | 0.0066604 | 2.1765 |
| DG 59:5 | 3.4037 | 1.7671 | 0.0066735 | 2.1756 |
| DG O-43:5 | 1.7704 | 0.82411 | 0.0066824 | 2.1751 |
| TG 16:0_19:3_19:3 | 6.1389 | 2.618 | 0.0067588 | 2.1701 |
| TG 44:2 | 1.649 | 0.72161 | 0.0068741 | 2.1628 |
| PC 15:0_18:4 | 1.5699 | 0.65064 | 0.0068786 | 2.1625 |
| FAHFA 33:1;O | 1.4615 | 0.54744 | 0.0069446 | 2.1584 |
| SCer 33:0;O | 2.5202 | 1.3336 | 0.0069597 | 2.1574 |
| Hex2Cer 40:5;O2 | 0.70388 | -0.50659 | 0.0070292 | 2.1531 |
| TG 13:0_14:0_16:0 | 1.597 | 0.67532 | 0.0070881 | 2.1495 |
| PC O-38:3 | 0.46619 | -1.101 | 0.0070921 | 2.1492 |
| Cer 18:0;O3/19:1;(2OH) | 2.122 | 1.0854 | 0.0071077 | 2.1483 |
| PC O-18:1_22:1 | 0.58782 | -0.76656 | 0.0071441 | 2.1461 |
| PI 40:1;O | 0.13887 | -2.8482 | 0.0071691 | 2.1445 |
| ACer 58:0;O3 | 2.537 | 1.3431 | 0.007174 | 2.1442 |
| Cer 18:0;O3/21:1;(2OH) | 1.8854 | 0.91487 | 0.0071963 | 2.1429 |
| MG 29:3;O | 1.8517 | 0.88884 | 0.0072111 | 2.142 |
| SM 44:5;O4 | 2.3079 | 1.2066 | 0.007253 | 2.1395 |
| Hex2Cer 30:0;O3 | 1.5723 | 0.65286 | 0.007272 | 2.1383 |
| FA 18:2 | 0.4273 | -1.2267 | 0.0073307 | 2.1349 |
| SHexCer 43:4;O3 | 0.40157 | -1.3163 | 0.0073534 | 2.1335 |
| PE 30:0 | 0.67066 | -0.57636 | 0.0073669 | 2.1327 |
| DG 30:1 | 2.4186 | 1.2741 | 0.0074064 | 2.1304 |
| PC 10:0_18:1 | 1.7349 | 0.79488 | 0.0074204 | 2.1296 |
| PI O-63:3 | 1.915 | 0.93733 | 0.0074427 | 2.1283 |
| PR 20:2;O3 | 0.26388 | -1.9221 | 0.0075318 | 2.1231 |
| TG 12:0_14:0_16:0 | 3.0476 | 1.6077 | 0.0075954 | 2.1194 |
| PI O-55:3 | 0.596 | -0.74663 | 0.0076749 | 2.1149 |
| PE 56:8 | 1.9271 | 0.9464 | 0.0076911 | 2.114 |
| TG 15:0_16:0_16:1 | 3.6966 | 1.8862 | 0.0077176 | 2.1125 |
| LPC 22:2 | 0.54759 | -0.86882 | 0.0077337 | 2.1116 |
| PI 38:10; PI 18:5-20:5; [M-H]- | 0.55718 | -0.84378 | 0.0077723 | 2.1094 |
| TG 16:0_18:1_22:4 | 4.5994 | 2.2014 | 0.0078158 | 2.107 |
| DG O-40:4 | 2.3118 | 1.209 | 0.0078541 | 2.1049 |
| PI 35:1 | 1.7432 | 0.80171 | 0.0078604 | 2.1046 |
| GlcADG 14:0_18:0 | 1.8275 | 0.8699 | 0.0079321 | 2.1006 |
| Car 22:4;O4 | 3.0515 | 1.6095 | 0.0079677 | 2.0987 |
| DG 57:5 | 5.3949 | 2.4316 | 0.0079743 | 2.0983 |
| DG O-47:5 | 0.43867 | -1.1888 | 0.0080101 | 2.0964 |
| DG 46:3 | 2.8319 | 1.5018 | 0.008036 | 2.095 |
| PE 40:1;O | 0.33791 | -1.5653 | 0.0080433 | 2.0946 |
| DG 59:6 | 6.7765 | 2.7605 | 0.0080611 | 2.0936 |
| NAE 19:0;O | 0.40128 | -1.3173 | 0.0081011 | 2.0915 |
| LPC 18:1 | 0.58405 | -0.77583 | 0.0083015 | 2.0808 |
| Cer 34:1;O4 | 0.64446 | -0.63384 | 0.0083265 | 2.0795 |
| ST 27:3;F | 0.36412 | -1.4575 | 0.0083804 | 2.0767 |
| PG 56:4;O | 0.59111 | -0.75851 | 0.0084912 | 2.071 |
| PC O-18:1_22:2 | 0.39621 | -1.3357 | 0.0084969 | 2.0707 |
| PA O-36:0;O | 0.34679 | -1.5279 | 0.0085408 | 2.0685 |
| HexCer 45:0;O5 | 2.0318 | 1.0227 | 0.0085636 | 2.0673 |
| DG 44:9 | 1.7755 | 0.8282 | 0.0085959 | 2.0657 |
| DG 56:2 | 0.50106 | -0.99696 | 0.0087023 | 2.0604 |
| DG 53:5 | 2.7468 | 1.4578 | 0.0089129 | 2.05 |
| Cer 39:5;O2 | 1.8883 | 0.9171 | 0.008992 | 2.0461 |
| CerP 54:6;O3 | 0.69633 | -0.52216 | 0.0091159 | 2.0402 |
| TG 42:0 | 2.5825 | 1.3687 | 0.0091546 | 2.0384 |
| MG O-20:1 | 0.36764 | -1.4436 | 0.0093063 | 2.0312 |
| HexCer 42:1;O5 | 0.60098 | -0.73461 | 0.0093796 | 2.0278 |
| TG 12:0_16:0_16:1 | 5.2791 | 2.4003 | 0.0094651 | 2.0239 |
| HexCer 36:0;O6 | 0.2203 | -2.1824 | 0.0094949 | 2.0225 |
| PI 14:0_18:0 | 2.3548 | 1.2356 | 0.0095293 | 2.0209 |
| PG 14:0_16:0 | 2.8269 | 1.4992 | 0.009546 | 2.0202 |
| DG 58:5 | 7.2081 | 2.8496 | 0.0095997 | 2.0177 |
| PA 45:4 | 0.55721 | -0.84371 | 0.0096123 | 2.0172 |
| DG O-18:1_20:4 | 2.4604 | 1.2989 | 0.0096822 | 2.014 |
| HexCer 47:1;O4 | 0.20916 | -2.2573 | 0.0097499 | 2.011 |
| HexCer 45:5;O2 | 0.33956 | -1.5583 | 0.0097676 | 2.0102 |
| SM 48:4;O3 | 0.32344 | -1.6284 | 0.00978 | 2.0097 |
| LPE 20:1 | 2.4287 | 1.2802 | 0.0098005 | 2.0088 |
| ACer 64:2;O4 | 2.9815 | 1.576 | 0.0099172 | 2.0036 |
| PE O-40:1 | 0.52448 | -0.93104 | 0.0099173 | 2.0036 |
| HexCer 39:0;O5 | 1.6811 | 0.7494 | 0.0099443 | 2.0024 |
| ST 27:3;O3 | 2.6207 | 1.3899 | 0.0099891 | 2.0005 |
| TG 88:13;O2 | 0.24288 | -2.0417 | 0.010012 | 1.9995 |
| MG 22:1 | 0.40913 | -1.2894 | 0.010041 | 1.9982 |
| HexCer 45:6;O6 | 0.71068 | -0.49274 | 0.010127 | 1.9945 |
| PC 15:0_18:3 | 1.7071 | 0.77154 | 0.010237 | 1.9898 |
| TG O-83:5 | 3.2263 | 1.6899 | 0.010319 | 1.9864 |
| PE-Cer 54:0;O5 | 1.6091 | 0.68627 | 0.010354 | 1.9849 |
| PC 18:0_26:1 | 1.9438 | 0.95889 | 0.010355 | 1.9848 |
| LPC O-34:0;O | 0.55827 | -0.84096 | 0.010386 | 1.9836 |
| TG 59:2 | 5.4327 | 2.4417 | 0.010396 | 1.9831 |
| Hex(8)-Cer 36:1;O2 | 0.69156 | -0.53207 | 0.010434 | 1.9816 |
| TG 46:2 | 6.5173 | 2.7043 | 0.010439 | 1.9814 |
| TG 50:0;O | 2.8105 | 1.4908 | 0.010482 | 1.9796 |
| TG 51:1 | 7.4452 | 2.8963 | 0.010496 | 1.979 |
| TG 14:0_15:0_17:0 | 1.9058 | 0.93039 | 0.010499 | 1.9788 |
| PE O-18:2_21:1 | 0.48988 | -1.0295 | 0.010522 | 1.9779 |
| DG 60:10 | 6.3051 | 2.6565 | 0.010601 | 1.9746 |
| TG 60:2 | 5.2668 | 2.3969 | 0.010617 | 1.974 |
| TG 14:0_16:0_16:1 | 8.3854 | 3.0679 | 0.010635 | 1.9733 |
| PE 40:2 | 2.9965 | 1.5833 | 0.010841 | 1.9649 |
| PE 36:3 | 16.98 | 4.0857 | 0.011031 | 1.9574 |
| Cer 40:0;O3 | 0.38908 | -1.3619 | 0.011037 | 1.9572 |
| TG 51:3;O2 | 1.6544 | 0.72632 | 0.011099 | 1.9547 |
| SM 42:2;O4 | 2.5127 | 1.3292 | 0.011147 | 1.9528 |
| MG 19:0;O | 2.8675 | 1.5198 | 0.011207 | 1.9505 |
| DG 54:5 | 3.3351 | 1.7377 | 0.011317 | 1.9463 |
| DG 33:0;O | 0.62271 | -0.68336 | 0.011334 | 1.9456 |
| HexCer 44:1;O6 | 0.63314 | -0.6594 | 0.011415 | 1.9425 |
| Cer 18:0;O3/35:1;(2OH) | 3.0636 | 1.6152 | 0.011589 | 1.9359 |
| Cer 46:1;O3 | 0.22777 | -2.1344 | 0.011856 | 1.9261 |
| PI-Cer 49:2;O3 | 0.24186 | -2.0478 | 0.011857 | 1.926 |
| PA 61:7 | 0.45429 | -1.1383 | 0.011914 | 1.924 |
| PC 18:0_24:0 | 1.929 | 0.94786 | 0.011914 | 1.9239 |
| TG 16:0_18:1_22:6 | 6.4907 | 2.6984 | 0.011968 | 1.922 |
| GM3(d18:1/16:0) | 2.5083 | 1.3267 | 0.011969 | 1.9219 |
| DG 17:0_18:1 | 2.0932 | 1.0657 | 0.011989 | 1.9212 |
| SM 44:6;O2 | 1.603 | 0.68076 | 0.011994 | 1.921 |
| SM 18:1;O2/25:1 | 0.55317 | -0.85421 | 0.012046 | 1.9192 |
| TG 16:0_18:0_22:2 | 5.6516 | 2.4987 | 0.012105 | 1.917 |
| DG O-39:4 | 2.1029 | 1.0724 | 0.012117 | 1.9166 |
| HexCer 50:2;O3 | 1.6996 | 0.76517 | 0.012142 | 1.9157 |
| TG 81:1;O3 | 1.6661 | 0.73646 | 0.012256 | 1.9117 |
| TG 49:0;O | 0.40695 | -1.2971 | 0.01229 | 1.9105 |
| LPG 18:1 | 0.40141 | -1.3168 | 0.012362 | 1.9079 |
| TG O-52:0;O | 1.7616 | 0.81692 | 0.012425 | 1.9057 |
| DG 45:3 | 1.4894 | 0.57469 | 0.012438 | 1.9052 |
| PC O-18:1_20:2 | 0.45763 | -1.1278 | 0.012645 | 1.8981 |
| LPE 20:3 | 1.7636 | 0.81852 | 0.012651 | 1.8979 |
| Hex(4)-HexNAc(2)-Fuc(2)-Cer 34:1;O2 | 0.37828 | -1.4025 | 0.012672 | 1.8972 |
| PC O-42:0 | 0.16998 | -2.5565 | 0.012777 | 1.8936 |
| Car 20:4 | 0.51883 | -0.94665 | 0.012801 | 1.8928 |
| TG 16:0_18:1_20:1 | 8.1996 | 3.0356 | 0.012894 | 1.8896 |
| PI 40:3 | 0.70735 | -0.49949 | 0.012901 | 1.8894 |
| DG O-38:5 | 1.7128 | 0.77637 | 0.012924 | 1.8886 |
| CE 20:4 | 0.44206 | -1.1777 | 0.012959 | 1.8874 |
| Cer 48:6;O3 | 0.52242 | -0.93671 | 0.012968 | 1.8871 |
| TG 33:1;O3 | 1.8748 | 0.90672 | 0.013041 | 1.8847 |
| PA 41:4 | 0.57481 | -0.79884 | 0.013049 | 1.8844 |
| PA 45:5 | 0.26272 | -1.9284 | 0.013061 | 1.884 |
| DG 32:2 | 2.0298 | 1.0213 | 0.013101 | 1.8827 |
| DG 55:5 | 4.2902 | 2.101 | 0.013109 | 1.8824 |
| TG 70:3;O3 | 2.0828 | 1.0586 | 0.013125 | 1.8819 |
| FA 24:1 | 0.38246 | -1.3866 | 0.013132 | 1.8817 |
| DG 62:10 | 3.8916 | 1.9604 | 0.01315 | 1.8811 |
| DG 42:4 | 1.4306 | 0.5166 | 0.013255 | 1.8776 |
| Cer 49:6;O4 | 1.4586 | 0.54461 | 0.013329 | 1.8752 |
| NAT 20:2;O | 3.3353 | 1.7378 | 0.013385 | 1.8734 |
| DG O-35:0;O2 | 0.55299 | -0.85468 | 0.013477 | 1.8704 |
| PC 18:0_22:0 | 1.5503 | 0.63254 | 0.013604 | 1.8663 |
| Cer 21:3;O2/24:2;O | 0.31143 | -1.683 | 0.013857 | 1.8583 |
| TG 95:5;O | 1.6543 | 0.72624 | 0.013858 | 1.8583 |
| PE 52:3 | 0.53609 | -0.89946 | 0.013896 | 1.8571 |
| TG 45:1;O | 0.55365 | -0.85296 | 0.013902 | 1.8569 |
| LPE 22:6 | 2.3083 | 1.2068 | 0.01392 | 1.8563 |
| DG 50:5 | 3.5563 | 1.8304 | 0.013936 | 1.8559 |
| TG 51:4;O2 | 1.5954 | 0.6739 | 0.013963 | 1.855 |
| PC O-22:3_26:1 | 3.3278 | 1.7346 | 0.013998 | 1.8539 |
| SHexCer 34:0;O2 | 0.66295 | -0.59303 | 0.014092 | 1.851 |
| PI-Cer 39:2;O2 | 0.42764 | -1.2255 | 0.014147 | 1.8493 |
| PR 58:14;O3 | 3.2384 | 1.6953 | 0.014176 | 1.8485 |
| DG 43:3 | 0.30702 | -1.7036 | 0.014465 | 1.8397 |
| PA 41:5 | 0.68871 | -0.53803 | 0.014485 | 1.8391 |
| HexCer 41:3;O4 | 1.4878 | 0.57314 | 0.014572 | 1.8365 |
| PA O-44:5 | 1.8661 | 0.90006 | 0.014641 | 1.8344 |
| NAE 19:2 | 0.3653 | -1.4529 | 0.01479 | 1.83 |
| TG 42:1 | 2.2474 | 1.1683 | 0.014793 | 1.8299 |
| Cer 42:2;O | 0.52425 | -0.93169 | 0.014881 | 1.8274 |
| TG 14:0_16:0_22:6 | 4.0937 | 2.0334 | 0.014898 | 1.8269 |
| PG 39:4;O | 2.2482 | 1.1688 | 0.014908 | 1.8266 |
| TG 58:2 | 6.5491 | 2.7113 | 0.014993 | 1.8241 |
| PI-Cer 37:1;O2 | 1.7287 | 0.78971 | 0.015032 | 1.823 |
| HexCer 44:2;O4 | 1.4737 | 0.55945 | 0.015076 | 1.8217 |
| TG O-14:0_18:0_20:2 | 1.897 | 0.92372 | 0.015143 | 1.8198 |
| SM 48:5;O2 | 3.0453 | 1.6066 | 0.01516 | 1.8193 |
| PC 15:0_18:0 | 1.4867 | 0.57214 | 0.01524 | 1.817 |
| TG 57:2 | 6.914 | 2.7895 | 0.015338 | 1.8142 |
| TG 84:6;O | 0.39076 | -1.3557 | 0.015427 | 1.8117 |
| MGDG 15:0_18:1 | 3.54 | 1.8238 | 0.015512 | 1.8093 |
| PIP3 24:0 | 0.23144 | -2.1113 | 0.01555 | 1.8083 |
| Cer 38:0;O | 2.0094 | 1.0068 | 0.015555 | 1.8081 |
| PC O-52:9 | 0.28342 | -1.819 | 0.01571 | 1.8038 |
| MIPC 32:2;O5 | 0.45546 | -1.1346 | 0.015803 | 1.8013 |
| DG 56:6 | 2.4625 | 1.3001 | 0.015838 | 1.8003 |
| TG 16:1_18:0_18:1 | 7.822 | 2.9675 | 0.01584 | 1.8002 |
| FA 21:2;O | 1.8314 | 0.87298 | 0.015851 | 1.7999 |
| PA 40:7 | 0.65436 | -0.61185 | 0.016002 | 1.7958 |
| PE 54:5;O | 2.1916 | 1.132 | 0.016073 | 1.7939 |
| TG 54:7 | 3.3356 | 1.7379 | 0.016378 | 1.7857 |
| PR 53:11;O2 | 0.23179 | -2.1091 | 0.016399 | 1.7852 |
| DG 52:1 | 0.65593 | -0.60839 | 0.016415 | 1.7847 |
| TG 55:2 | 2.7424 | 1.4554 | 0.016508 | 1.7823 |
| DG 58:6 | 3.8344 | 1.939 | 0.016539 | 1.7815 |
| SHexCer 40:1;O4 | 0.47757 | -1.0662 | 0.01656 | 1.7809 |
| DG 36:2 | 1.5566 | 0.63842 | 0.016608 | 1.7797 |
| PC O-22:3_24:0 | 2.4678 | 1.3032 | 0.016675 | 1.7779 |
| PG 56:7;O | 1.8845 | 0.91415 | 0.016803 | 1.7746 |
| Cer 41:6;O2 | 3.1427 | 1.652 | 0.0171 | 1.767 |
| TG 52:5 | 3.4188 | 1.7735 | 0.017178 | 1.765 |
| DG 60:7 | 3.3307 | 1.7358 | 0.017216 | 1.7641 |
| TG 56:2 | 2.5968 | 1.3767 | 0.017256 | 1.7631 |
| CerP 51:3;O2 | 3.9061 | 1.9657 | 0.017466 | 1.7578 |
| MG O-28:0;O | 1.4107 | 0.49638 | 0.017507 | 1.7568 |
| DG 61:8 | 2.1234 | 1.0863 | 0.017561 | 1.7554 |
| DG 36:3 | 2.5512 | 1.3512 | 0.017682 | 1.7525 |
| PI 36:5 | 0.35452 | -1.496 | 0.017858 | 1.7482 |
| PIP3 35:6 | 1.7507 | 0.8079 | 0.017874 | 1.7478 |
| TG 58:2;O2 | 2.9915 | 1.5809 | 0.017877 | 1.7477 |
| PA O-41:6 | 0.45069 | -1.1498 | 0.017896 | 1.7472 |
| SPB 20:2;O6 | 0.49114 | -1.0258 | 0.01804 | 1.7438 |
| PA 48:4 | 1.6173 | 0.69354 | 0.018045 | 1.7436 |
| TG O-18:0_14:0_16:1 | 1.587 | 0.66626 | 0.018096 | 1.7424 |
| TG 17:1_18:0_18:1 | 6.4015 | 2.6784 | 0.018096 | 1.7424 |
| TG 54:3 | 2.7608 | 1.4651 | 0.018146 | 1.7412 |
| PE 25:1;O | 1.5479 | 0.63034 | 0.018234 | 1.7391 |
| PA 45:8 | 0.5561 | -0.8466 | 0.018432 | 1.7344 |
| TG 50:3 | 3.49 | 1.8032 | 0.018441 | 1.7342 |
| LPE 24:0 | 0.52396 | -0.93247 | 0.01846 | 1.7338 |
| PI-Cer 46:0;O3 | 0.54977 | -0.86311 | 0.018472 | 1.7335 |
| PI O-38:4 | 0.66384 | -0.5911 | 0.018577 | 1.731 |
| PG 44:9 | 0.48161 | -1.0541 | 0.018636 | 1.7296 |
| PI-Cer 38:0;O4 | 0.40085 | -1.3189 | 0.018748 | 1.727 |
| DG O-39:3 | 1.4542 | 0.54023 | 0.018845 | 1.7248 |
| HexCer 43:4;O2 | 0.47786 | -1.0653 | 0.018872 | 1.7242 |
| HexCer 41:5;O4 | 0.37579 | -1.412 | 0.018908 | 1.7233 |
| CerP 52:0;O5 | 2.5153 | 1.3307 | 0.019026 | 1.7207 |
| Cer 45:6;O3 | 0.56774 | -0.8167 | 0.019115 | 1.7186 |
| Hex2Cer 34:1;O3 | 0.37091 | -1.4309 | 0.019175 | 1.7173 |
| Cer 44:6;O2 | 1.8454 | 0.88394 | 0.0192 | 1.7167 |
| TG 86:12;O2 | 0.31817 | -1.6521 | 0.019238 | 1.7158 |
| FA 17:1 | 0.63371 | -0.6581 | 0.019243 | 1.7157 |
| DG 19:0_18:1 | 1.8829 | 0.91292 | 0.019396 | 1.7123 |
| CerP 54:1;O5 | 2.6956 | 1.4306 | 0.01942 | 1.7118 |
| CE 22:5 | 0.66093 | -0.59744 | 0.019557 | 1.7087 |
| DG 45:2;O | 2.2632 | 1.1784 | 0.019592 | 1.7079 |
| MG 31:0 | 1.5476 | 0.63007 | 0.019601 | 1.7077 |
| LPC 18:0 | 2.0223 | 1.016 | 0.019632 | 1.707 |
| DG 16:0_18:2 | 2.642 | 1.4016 | 0.019703 | 1.7055 |
| Cer 18:0;O3/42:1;(2OH) | 2.2075 | 1.1424 | 0.019992 | 1.6991 |
| SM 44:4;O5 | 2.4991 | 1.3214 | 0.020123 | 1.6963 |
| ACer 56:2;O2 | 0.68965 | -0.53605 | 0.020174 | 1.6952 |
| PA 48:0 | 0.22886 | -2.1275 | 0.02018 | 1.6951 |
| DG 52:5 | 2.6761 | 1.4201 | 0.020186 | 1.695 |
| TG 62:2 | 2.8727 | 1.5224 | 0.020241 | 1.6938 |
| HexCer 39:2;O5 | 0.29772 | -1.748 | 0.020258 | 1.6934 |
| Car 34:2;O | 2.0079 | 1.0057 | 0.020381 | 1.6908 |
| TG 98:19 | 2.511 | 1.3283 | 0.020405 | 1.6903 |
| PG 18:1_20:1 | 0.37484 | -1.4157 | 0.020509 | 1.6881 |
| PC 34:2 | 1.6208 | 0.69668 | 0.020518 | 1.6879 |
| PI O-67:6 | 1.6156 | 0.69204 | 0.020567 | 1.6868 |
| PG 44:7 | 1.6018 | 0.67969 | 0.020579 | 1.6866 |
| LPE 24:1 | 0.49337 | -1.0192 | 0.020682 | 1.6844 |
| Cer 39:4;O | 1.4461 | 0.53215 | 0.020706 | 1.6839 |
| PS 20:0_16:2 | 0.61067 | -0.71154 | 0.020729 | 1.6834 |
| PI O-65:4 | 2.2658 | 1.18 | 0.02074 | 1.6832 |
| PC O-40:3 | 0.51469 | -0.95823 | 0.020978 | 1.6782 |
| PC 28:0;O | 1.7444 | 0.80274 | 0.020984 | 1.6781 |
| SHEXcer 32:5;O3 | 0.64122 | -0.64112 | 0.021212 | 1.6734 |
| PG 42:2;O | 0.62226 | -0.6844 | 0.021293 | 1.6718 |
| TG 82:1;O3 | 0.40523 | -1.3032 | 0.021315 | 1.6713 |
| PC O-14:1_10:0 | 0.61259 | -0.707 | 0.02137 | 1.6702 |
| DG O-16:0_20:3 | 0.40627 | -1.2995 | 0.021751 | 1.6625 |
| TG 14:0_16:1_18:1 | 5.1182 | 2.3556 | 0.021819 | 1.6612 |
| PE-Cer 28:2;O2 | 2.7247 | 1.4461 | 0.021835 | 1.6609 |
| PA O-44:0;O | 0.52884 | -0.91909 | 0.021921 | 1.6591 |
| ST 23:2;O4;S | 0.58655 | -0.76968 | 0.022251 | 1.6527 |
| TG 57:9;O3 | 0.30375 | -1.7191 | 0.022697 | 1.644 |
| LPI 30:5 | 0.58135 | -0.78253 | 0.022697 | 1.644 |
| SHEXcer 43:4;O4 | 2.6124 | 1.3854 | 0.022724 | 1.6435 |
| PA 43:4 | 0.67359 | -0.57005 | 0.022768 | 1.6427 |
| LPS 28:0;O | 1.5087 | 0.59331 | 0.022806 | 1.642 |
| MG 18:2;O | 0.15982 | -2.6455 | 0.022822 | 1.6417 |
| TG 52:1;O | 2.0466 | 1.0332 | 0.022864 | 1.6408 |
| TG O-85:10 | 0.52152 | -0.93921 | 0.023118 | 1.6361 |
| Cer 15:2;2O/38:10 | 4.8657 | 2.2826 | 0.023187 | 1.6348 |
| Hex2Cer 30:1;O3 | 0.48536 | -1.0429 | 0.023349 | 1.6317 |
| DG O-18:1_22:6 | 1.7685 | 0.82252 | 0.023518 | 1.6286 |
| PI 18:0_22:2 | 0.70941 | -0.49531 | 0.023548 | 1.628 |
| PG O-38:0 | 2.0745 | 1.0528 | 0.023653 | 1.6261 |
| DG 68:9;O | 0.61607 | -0.69883 | 0.02377 | 1.624 |
| LPC 22:5 | 0.21582 | -2.2121 | 0.023932 | 1.621 |
| Hex2Cer 30:4;O4 | 1.976 | 0.98256 | 0.024175 | 1.6166 |
| ST 29:7;O | 3.1915 | 1.6743 | 0.024319 | 1.6141 |
| PI-Cer 35:0;O2 | 1.718 | 0.78076 | 0.024364 | 1.6133 |
| PC O-18:1_24:1 | 0.40732 | -1.2958 | 0.024529 | 1.6103 |
| MG 31:1 | 0.57352 | -0.80207 | 0.024593 | 1.6092 |
| TG 14:0_18:1_19:1 | 5.7694 | 2.5284 | 0.024778 | 1.6059 |
| PI 40:10 | 0.48307 | -1.0497 | 0.024787 | 1.6058 |
| FAHFA 26:2;O | 1.8028 | 0.85027 | 0.024874 | 1.6042 |
| PE 18:1_26:4 | 0.58901 | -0.76363 | 0.024895 | 1.6039 |
| PE O-18:2_24:2 | 0.39115 | -1.3542 | 0.024899 | 1.6038 |
| TG 15:0_16:1_18:1 | 2.6021 | 1.3797 | 0.02506 | 1.601 |
| TG 50:4;O2 | 1.579 | 0.65905 | 0.025095 | 1.6004 |
| HexCer 18:0;O2/20:1 | 1.4932 | 0.57843 | 0.025166 | 1.5992 |
| PA O-56:5 | 3.3237 | 1.7328 | 0.025184 | 1.5989 |
| ST 27:3 | 0.26178 | -1.9336 | 0.0252 | 1.5986 |
| DG O-35:0 | 1.8075 | 0.85397 | 0.025253 | 1.5977 |
| DG 64:7 | 1.961 | 0.97158 | 0.025301 | 1.5969 |
| HexCer 45:5;O5 | 1.9194 | 0.94063 | 0.025398 | 1.5952 |
| CE 28:1;O | 2.1627 | 1.1129 | 0.025524 | 1.593 |
| PG 18:1_20:2 | 0.45535 | -1.1349 | 0.025604 | 1.5917 |
| TG 14:0_18:1_18:2 | 3.9448 | 1.98 | 0.025647 | 1.591 |
| Cer 36:0;O | 1.5804 | 0.66025 | 0.025777 | 1.5888 |
| PE-NMe2 36:2 | 1.5123 | 0.59676 | 0.025809 | 1.5882 |
| CerP 51:6;O2 | 0.6191 | -0.69176 | 0.025836 | 1.5878 |
| Cer 45:6;O | 0.30564 | -1.7101 | 0.026007 | 1.5849 |
| TG 45:2 | 5.7472 | 2.5229 | 0.026131 | 1.5828 |
| TG 58:3;O2 | 4.6981 | 2.2321 | 0.026142 | 1.5827 |
| FA 24:2 | 0.39684 | -1.3334 | 0.026186 | 1.5819 |
| PA 37:8;O | 0.53905 | -0.89152 | 0.026215 | 1.5815 |
| HexCer 39:6;O3 | 0.54155 | -0.88484 | 0.026216 | 1.5814 |
| PE O-40:5 | 0.51456 | -0.9586 | 0.026229 | 1.5812 |
| DG 58:1 | 2.2759 | 1.1864 | 0.026248 | 1.5809 |
| DG 15:0_18:1 | 1.9756 | 0.98231 | 0.026621 | 1.5748 |
| NAE 19:1 | 0.16461 | -2.6028 | 0.026937 | 1.5696 |
| DG 41:7;O | 1.6092 | 0.68636 | 0.027198 | 1.5655 |
| PC O-68:1 | 1.7472 | 0.80501 | 0.02739 | 1.5624 |
| PI 18:0_19:2 | 0.55682 | -0.84472 | 0.0277 | 1.5575 |
| PA O-37:1 | 0.47341 | -1.0788 | 0.027733 | 1.557 |
| PA 40:0 | 0.67031 | -0.5771 | 0.027745 | 1.5568 |
| MG O-32:1 | 0.63483 | -0.65556 | 0.028128 | 1.5509 |
| PA O-43:1 | 0.48062 | -1.057 | 0.028286 | 1.5484 |
| TG 77:1;O3 | 1.6641 | 0.73476 | 0.028306 | 1.5481 |
| MG O-19:1 | 0.63363 | -0.65828 | 0.028363 | 1.5472 |
| ST 28:2;O4 | 0.33583 | -1.5742 | 0.028465 | 1.5457 |
| DG 62:11 | 3.3606 | 1.7487 | 0.028512 | 1.545 |
| PA 30:0 | 2.4837 | 1.3125 | 0.028582 | 1.5439 |
| TG 52:1 | 1.4122 | 0.49794 | 0.028641 | 1.543 |
| PI O-16:0_18:2 | 0.4522 | -1.145 | 0.028823 | 1.5403 |
| PA O-38:0;O | 0.58041 | -0.78486 | 0.029107 | 1.536 |
| PC O-36:3 | 2.4391 | 1.2864 | 0.029383 | 1.5319 |
| PI O-38:2 | 0.30327 | -1.7213 | 0.029597 | 1.5288 |
| PE 54:5 | 4.4096 | 2.1406 | 0.029633 | 1.5282 |
| PE 44:3;O | 0.44813 | -1.158 | 0.029684 | 1.5275 |
| MG 34:2 | 0.56847 | -0.81486 | 0.029744 | 1.5266 |
| PC 26:0_18:1 | 1.6997 | 0.76531 | 0.029753 | 1.5265 |
| TG 16:0_18:1_18:2 | 5.0801 | 2.3449 | 0.030098 | 1.5215 |
| PC 15:0_16:3 | 1.9887 | 0.99184 | 0.030174 | 1.5204 |
| Cer 44:4;O3 | 2.3491 | 1.2321 | 0.030207 | 1.5199 |
| CE 26:2 | 0.41841 | -1.257 | 0.030322 | 1.5182 |
| ST 30:2;O6;G | 0.32968 | -1.6009 | 0.030351 | 1.5178 |
| SM 40:2;O2 | 3.6278 | 1.8591 | 0.030353 | 1.5178 |
| TG O-57:3 | 1.6613 | 0.73233 | 0.030369 | 1.5176 |
| PIP 64:5;O | 0.50848 | -0.97573 | 0.030501 | 1.5157 |
| DG O-18:1_17:1 | 0.28876 | -1.7921 | 0.030798 | 1.5115 |
| TG O-55:2 | 1.9584 | 0.96969 | 0.030934 | 1.5096 |
| HBMP 18:0_20:3_18:0 | 0.59629 | -0.74592 | 0.0311 | 1.5072 |
| ST 26:1;O3;G | 0.39707 | -1.3325 | 0.031331 | 1.504 |
| PI 18:0_22:6 | 1.604 | 0.6817 | 0.031377 | 1.5034 |
| HexCer 44:6;O6 | 0.70713 | -0.49995 | 0.031472 | 1.5021 |
| PA 16:1_20:1 | 0.37809 | -1.4032 | 0.031477 | 1.502 |
| PC 18:1_20:1 | 0.61792 | -0.69452 | 0.031479 | 1.502 |
| LPC 32:1 | 0.68873 | -0.53799 | 0.031521 | 1.5014 |
| Cer 36:0;O2 | 1.7386 | 0.79795 | 0.031542 | 1.5011 |
| TG 12:0_12:3_20:6 | 1.6606 | 0.73172 | 0.031563 | 1.5008 |
| PE O-18:2_20:1 | 0.37647 | -1.4094 | 0.031649 | 1.4996 |
| DG 33:0_18:1 | 2.3491 | 1.2321 | 0.031981 | 1.4951 |
| GP 32:0 | 0.57346 | -0.80224 | 0.031981 | 1.4951 |
| DG 16:0_28:0 | 2.4833 | 1.3123 | 0.03228 | 1.4911 |
| HexCer 41:2;O4 | 0.52242 | -0.93673 | 0.032523 | 1.4878 |
| CerP 41:6;O4 | 1.5547 | 0.63667 | 0.032669 | 1.4859 |
| CPA 18:1 | 0.38132 | -1.3909 | 0.032697 | 1.4855 |
| TG 57:9 | 1.4962 | 0.58134 | 0.032776 | 1.4844 |
| PC O-42:3 | 0.42615 | -1.2306 | 0.032885 | 1.483 |
| PE O-16:0_20:5 | 0.56541 | -0.82262 | 0.03289 | 1.4829 |
| PE 38:0 | 1.5037 | 0.58851 | 0.032923 | 1.4825 |
| NA 40:2;O5;N | 0.55195 | -0.8574 | 0.033231 | 1.4785 |
| FA 20:0 | 0.67386 | -0.56947 | 0.033314 | 1.4774 |
| PG 54:6;O | 1.494 | 0.57918 | 0.033361 | 1.4768 |
| TG 58:7 | 2.9719 | 1.5714 | 0.033472 | 1.4753 |
| LPI 34:2 | 0.51012 | -0.97109 | 0.033552 | 1.4743 |
| MIPC 34:2;O2 | 2.039 | 1.0278 | 0.033753 | 1.4717 |
| DG 18:1_20:2 | 1.792 | 0.84156 | 0.034049 | 1.4679 |
| TG O-38:1 | 1.4007 | 0.48612 | 0.034076 | 1.4676 |
| PE 16:0_18:2 | 13.284 | 3.7316 | 0.034215 | 1.4658 |
| DG O-44:9 | 1.5684 | 0.64927 | 0.034303 | 1.4647 |
| DG 36:1;O2 | 0.62789 | -0.67141 | 0.034399 | 1.4635 |
| NAE 17:0 | 0.57625 | -0.79524 | 0.034765 | 1.4589 |
| SM 40:4;O2 | 0.33387 | -1.5826 | 0.034786 | 1.4586 |
| PC O-18:1_24:2 | 0.44687 | -1.1621 | 0.035093 | 1.4548 |
| DG O-48:3 | 2.1829 | 1.1262 | 0.03513 | 1.4543 |
| PIP2 21:5 | 0.58192 | -0.7811 | 0.035477 | 1.45 |
| Cer 45:2;O3 | 0.30205 | -1.7271 | 0.035683 | 1.4475 |
| DG 60:6 | 3.3278 | 1.7346 | 0.035713 | 1.4472 |
| HexCer 42:5;O4 | 1.5511 | 0.63329 | 0.036007 | 1.4436 |
| TG 56:3;O2 | 4.4591 | 2.1567 | 0.036147 | 1.4419 |
| FA 18:0;O | 0.51768 | -0.94987 | 0.036306 | 1.44 |
| PE O-18:2_22:1 | 0.44186 | -1.1784 | 0.036419 | 1.4387 |
| CL 74:4 | 0.64246 | -0.63833 | 0.036486 | 1.4379 |
| Cer 53:6;O | 0.49282 | -1.0209 | 0.036511 | 1.4376 |
| PG 56:3;O | 0.68607 | -0.54357 | 0.036523 | 1.4374 |
| PE 46:1 | 2.545 | 1.3477 | 0.036526 | 1.4374 |
| PI O-56:3 | 1.4854 | 0.57084 | 0.036588 | 1.4367 |
| HexCer 30:1;O6 | 1.416 | 0.50179 | 0.036622 | 1.4363 |
| ACer 56:0;O3 | 2.2694 | 1.1823 | 0.036761 | 1.4346 |
| LPC 20:0 | 1.8113 | 0.857 | 0.036815 | 1.434 |
| ST 33:6;O3 | 0.49022 | -1.0285 | 0.037132 | 1.4302 |
| PI 54:4;O | 0.48308 | -1.0497 | 0.03714 | 1.4302 |
| PI O-56:2 | 1.6615 | 0.73251 | 0.037269 | 1.4287 |
| ST 28:3;O3;Cl | 0.3683 | -1.441 | 0.037375 | 1.4274 |
| PG O-54:0;O | 0.64237 | -0.63853 | 0.037383 | 1.4273 |
| PA O-56:7 | 1.8618 | 0.89667 | 0.037482 | 1.4262 |
| PA 45:9 | 0.58617 | -0.7706 | 0.037619 | 1.4246 |
| PE-Cer 38:1;O2 | 0.69715 | -0.52046 | 0.037628 | 1.4245 |
| HexCer 54:6;O3 | 2.2407 | 1.1639 | 0.037781 | 1.4227 |
| DG dO-40:0 | 1.6469 | 0.71973 | 0.03792 | 1.4211 |
| Hex2Cer 30:0;O4 | 0.26399 | -1.9214 | 0.038194 | 1.418 |
| PI O-41:8 | 0.42003 | -1.2514 | 0.038204 | 1.4179 |
| DG 50:3 | 3.1024 | 1.6334 | 0.038525 | 1.4143 |
| TG 16:1_18:1_19:1 | 5.0963 | 2.3495 | 0.03855 | 1.414 |
| SM 20:0;O2/16:1 | 1.4452 | 0.53126 | 0.038682 | 1.4125 |
| DG 66:9;O | 0.69125 | -0.53272 | 0.038995 | 1.409 |
| Cer 44:1;O4 | 1.4904 | 0.57572 | 0.039267 | 1.406 |
| PE 54:6 | 3.3178 | 1.7302 | 0.039467 | 1.4038 |
| TG 16:0_18:2_20:2 | 2.5752 | 1.3647 | 0.03952 | 1.4032 |
| DG O-45:5 | 1.901 | 0.92678 | 0.039573 | 1.4026 |
| LPS O-26:0;O | 0.65845 | -0.60286 | 0.039778 | 1.4004 |
| HexCer 42:6;O3 | 0.67086 | -0.57592 | 0.03995 | 1.3985 |
| Cer 21:0;O3/24:1;(2OH) | 1.8158 | 0.86063 | 0.040037 | 1.3975 |
| TG 101:6 | 1.673 | 0.74243 | 0.040164 | 1.3962 |
| SPB 23:3;O3 | 0.36828 | -1.4411 | 0.040168 | 1.3961 |
| HexCer 42:3;O2 | 0.57273 | -0.80407 | 0.040245 | 1.3953 |
| LPC O-20:4 | 0.43178 | -1.2116 | 0.040343 | 1.3942 |
| LPC 33:2 | 3.238 | 1.6951 | 0.040368 | 1.394 |
| LPC O-29:0;O | 0.68482 | -0.5462 | 0.040512 | 1.3924 |
| TG 85:1;O2 | 0.34416 | -1.5389 | 0.040896 | 1.3883 |
| TG 15:0_21:2_22:6 | 3.6809 | 1.8801 | 0.041026 | 1.3869 |
| Cer 43:0;O2 | 0.67474 | -0.56759 | 0.041092 | 1.3862 |
| PC 18:0_22:3 | 0.64119 | -0.64119 | 0.041316 | 1.3839 |
| CerP 54:1;O3 | 0.64086 | -0.64192 | 0.041328 | 1.3838 |
| DG O-41:5 | 1.8897 | 0.91818 | 0.041379 | 1.3832 |
| SM 32:6;O5 | 1.5874 | 0.66664 | 0.041463 | 1.3823 |
| PE O-40:4 | 0.59586 | -0.74695 | 0.041529 | 1.3816 |
| SM 42:1;O2 | 0.46912 | -1.092 | 0.041604 | 1.3809 |
| TG 14:0_14:3_19:5 | 2.7075 | 1.4369 | 0.041814 | 1.3787 |
| PC O-20:3_16:0 | 0.63118 | -0.66388 | 0.042018 | 1.3766 |
| NAT 33:1;O | 1.5604 | 0.64191 | 0.042114 | 1.3756 |
| TG 73:1;O3 | 1.5727 | 0.65328 | 0.042158 | 1.3751 |
| DG O-56:6 | 2.2973 | 1.1999 | 0.042176 | 1.3749 |
| TG 16:1_18:0_20:2 | 5.1042 | 2.3517 | 0.042444 | 1.3722 |
| PIP 38:3 | 0.48311 | -1.0496 | 0.042662 | 1.37 |
| TG 14:0_15:0_16:0 | 1.413 | 0.49874 | 0.042721 | 1.3694 |
| DG 27:0_21:1 | 2.7603 | 1.4648 | 0.04282 | 1.3684 |
| Cer 45:6;O6 | 0.68952 | -0.53633 | 0.042926 | 1.3673 |
| Car 32:4;O | 1.8535 | 0.89025 | 0.04328 | 1.3637 |
| DG O-16:1_20:4 | 1.9815 | 0.98661 | 0.044069 | 1.3559 |
| TG 14:2_14:3_17:3 | 2.6453 | 1.4034 | 0.044451 | 1.3521 |
| ACer 58:4;O2 | 0.62833 | -0.67041 | 0.044478 | 1.3519 |
| Cer 48:6;O2 | 0.65636 | -0.60744 | 0.044494 | 1.3517 |
| PA 25:0_18:4 | 1.4608 | 0.54675 | 0.044502 | 1.3516 |
| SM 14:0;O2/18:2 | 0.38685 | -1.3702 | 0.044581 | 1.3509 |
| HC 15:1;O | 0.58874 | -0.76431 | 0.044782 | 1.3489 |
| PA O-39:1 | 0.57448 | -0.79966 | 0.044843 | 1.3483 |
| LPE 18:2 | 2.2334 | 1.1593 | 0.044969 | 1.3471 |
| PI-Cer 35:1;O2 | 1.7091 | 0.77325 | 0.045068 | 1.3461 |
| TG 72:12;O2 | 0.54101 | -0.88627 | 0.045439 | 1.3426 |
| PE 52:5;O | 1.7222 | 0.78422 | 0.04551 | 1.3419 |
| PE O-18:2_22:2 | 0.54309 | -0.88073 | 0.045632 | 1.3407 |
| LPC 26:0 | 1.4756 | 0.56131 | 0.045757 | 1.3395 |
| DG 68:10;O | 0.6893 | -0.5368 | 0.045778 | 1.3393 |
| DG 39:0;O2 | 1.6664 | 0.73677 | 0.045781 | 1.3393 |
| PE O-18:0_22:3 | 0.44898 | -1.1553 | 0.04597 | 1.3375 |
| DG 52:3 | 2.4541 | 1.2952 | 0.046037 | 1.3369 |
| TG 94:4;O2 | 2.1008 | 1.0709 | 0.046231 | 1.3351 |
| TG 97:7;O3 | 0.58336 | -0.77753 | 0.046301 | 1.3344 |
| LPC 32:4 | 0.69345 | -0.52814 | 0.046611 | 1.3315 |
| CerP 54:2;O3 | 0.57025 | -0.81034 | 0.046685 | 1.3308 |
| PE O-38:3 | 0.60367 | -0.72816 | 0.046779 | 1.3299 |
| PE 52:5 | 1.7686 | 0.8226 | 0.04708 | 1.3272 |
| PG 30:2 | 0.63594 | -0.65303 | 0.047237 | 1.3257 |
| TG 85:1;O3 | 1.6387 | 0.71253 | 0.047322 | 1.3249 |
| Cer 47:6;O2 | 0.69979 | -0.515 | 0.047373 | 1.3245 |
| DG 32:0_22:2 | 2.3101 | 1.208 | 0.047457 | 1.3237 |
| TG 60:3 | 4.3305 | 2.1145 | 0.047513 | 1.3232 |
| PG 28:2 | 1.5896 | 0.66863 | 0.047609 | 1.3223 |
| DG 33:3 | 1.4286 | 0.51462 | 0.048218 | 1.3168 |
| DG 62:6 | 2.6099 | 1.384 | 0.048246 | 1.3165 |
| PIP 68:8;O | 0.48315 | -1.0495 | 0.048617 | 1.3132 |
| DG 54:7 | 1.9146 | 0.93707 | 0.048676 | 1.3127 |
| PC 44:7 | 0.66212 | -0.59483 | 0.048825 | 1.3114 |
| PE-Cer 42:6;O3 | 1.993 | 0.99491 | 0.048936 | 1.3104 |
| PC 43:7 | 0.46591 | -1.1019 | 0.049275 | 1.3074 |
| Cer 35:1;O2 | 0.65243 | -0.6161 | 0.049337 | 1.3068 |
| HexCer 39:3;O5 | 0.38932 | -1.361 | 0.049459 | 1.3058 |

**Supplemental Table 4.**
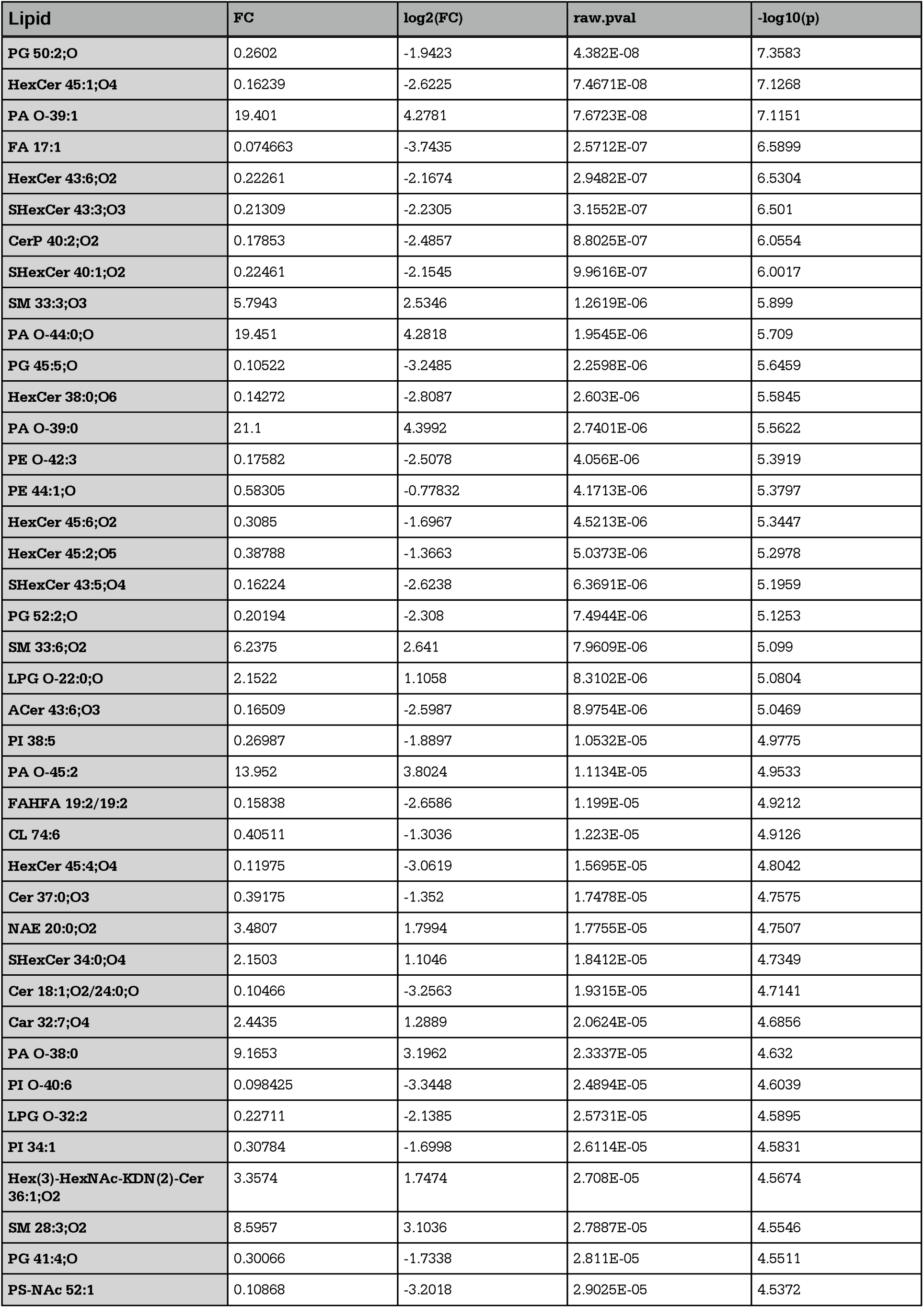
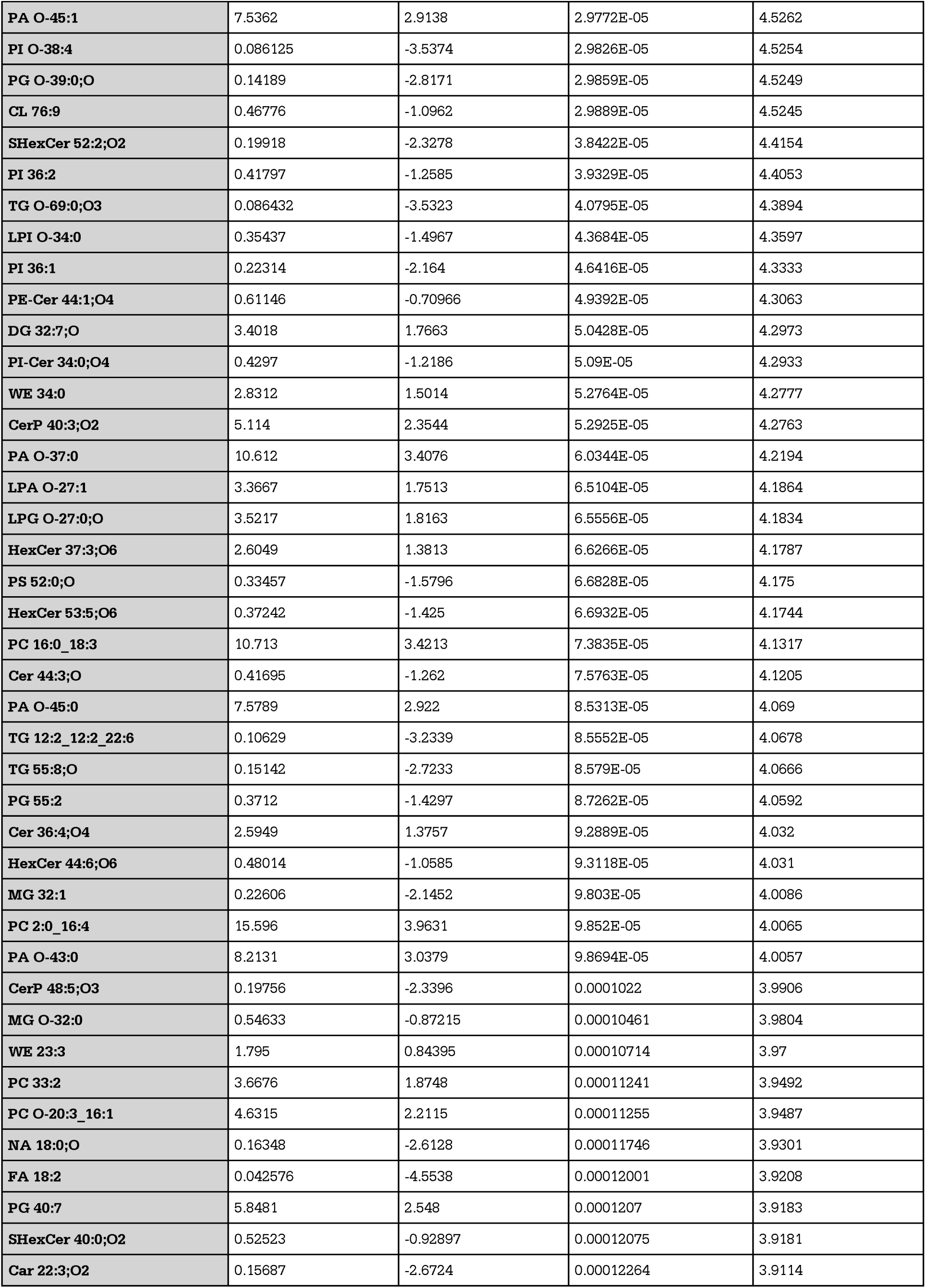
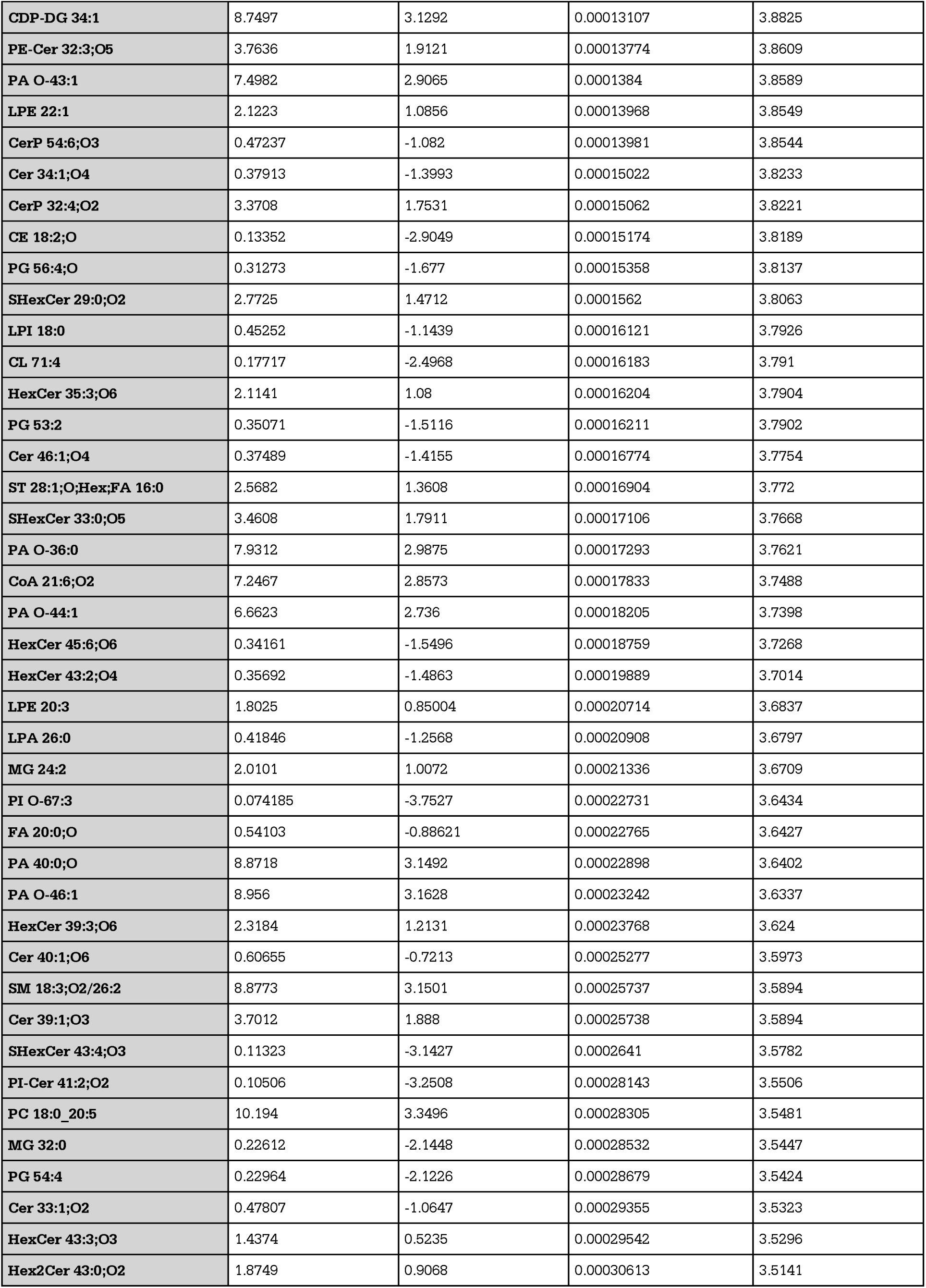
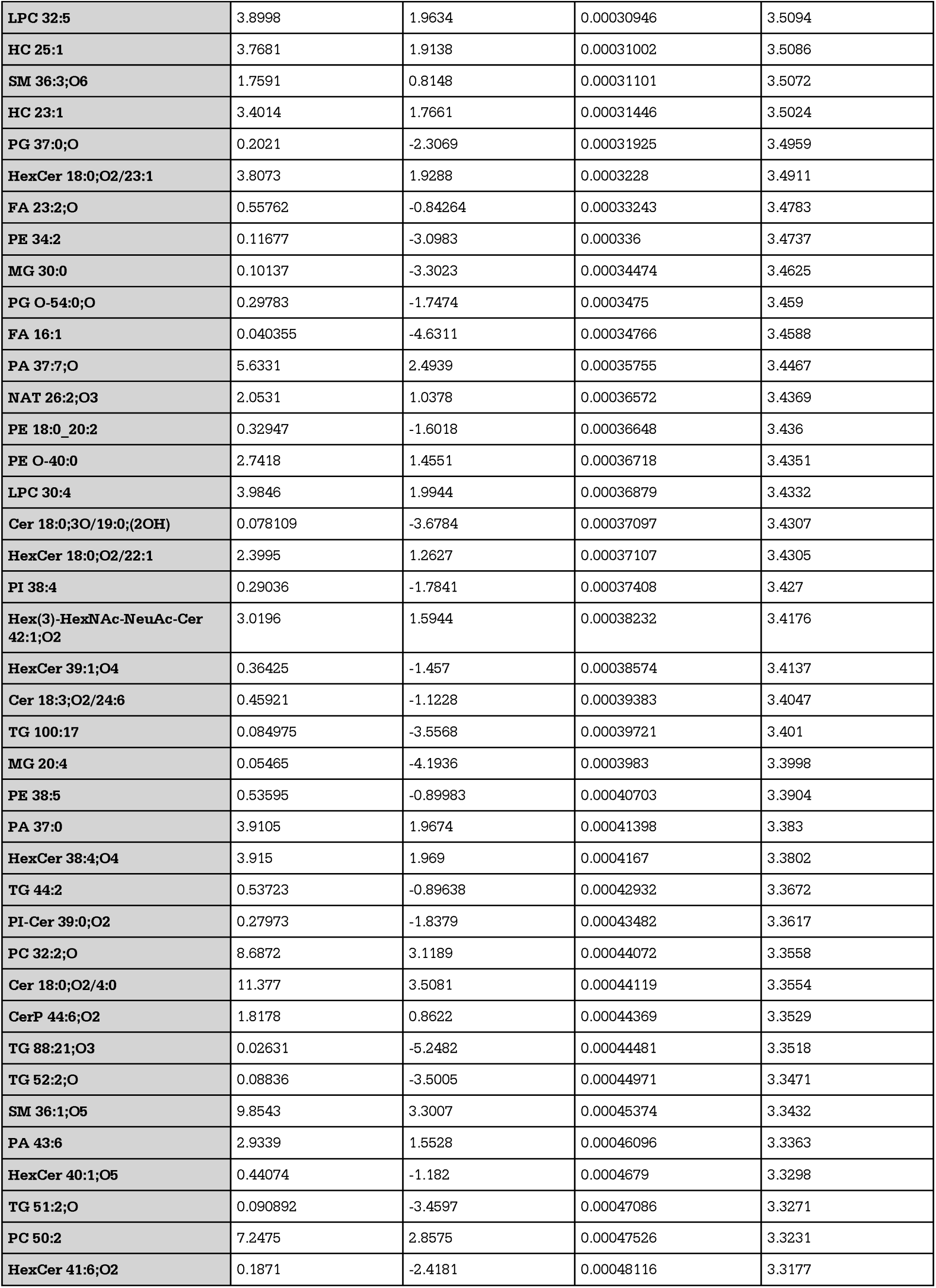
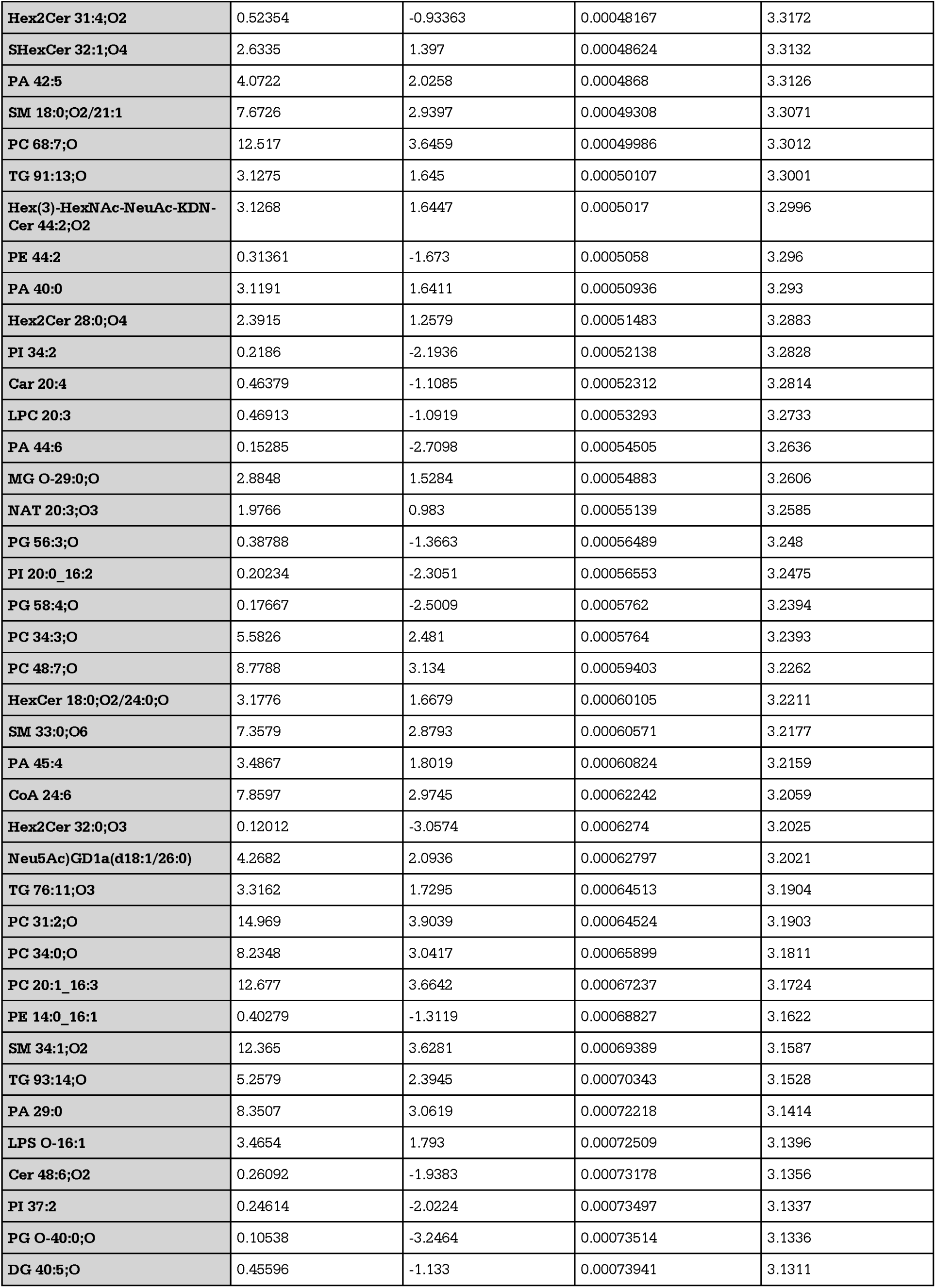
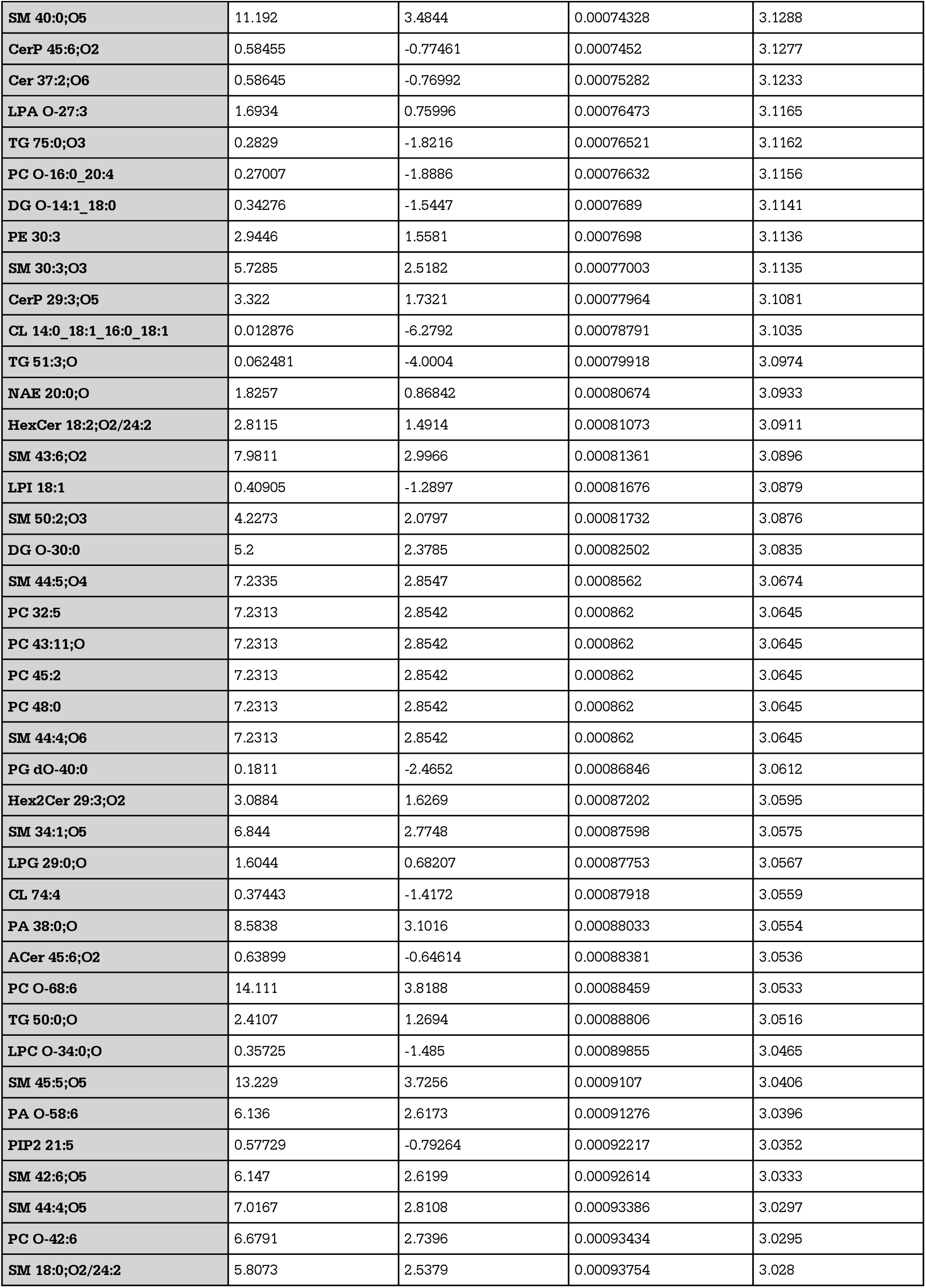
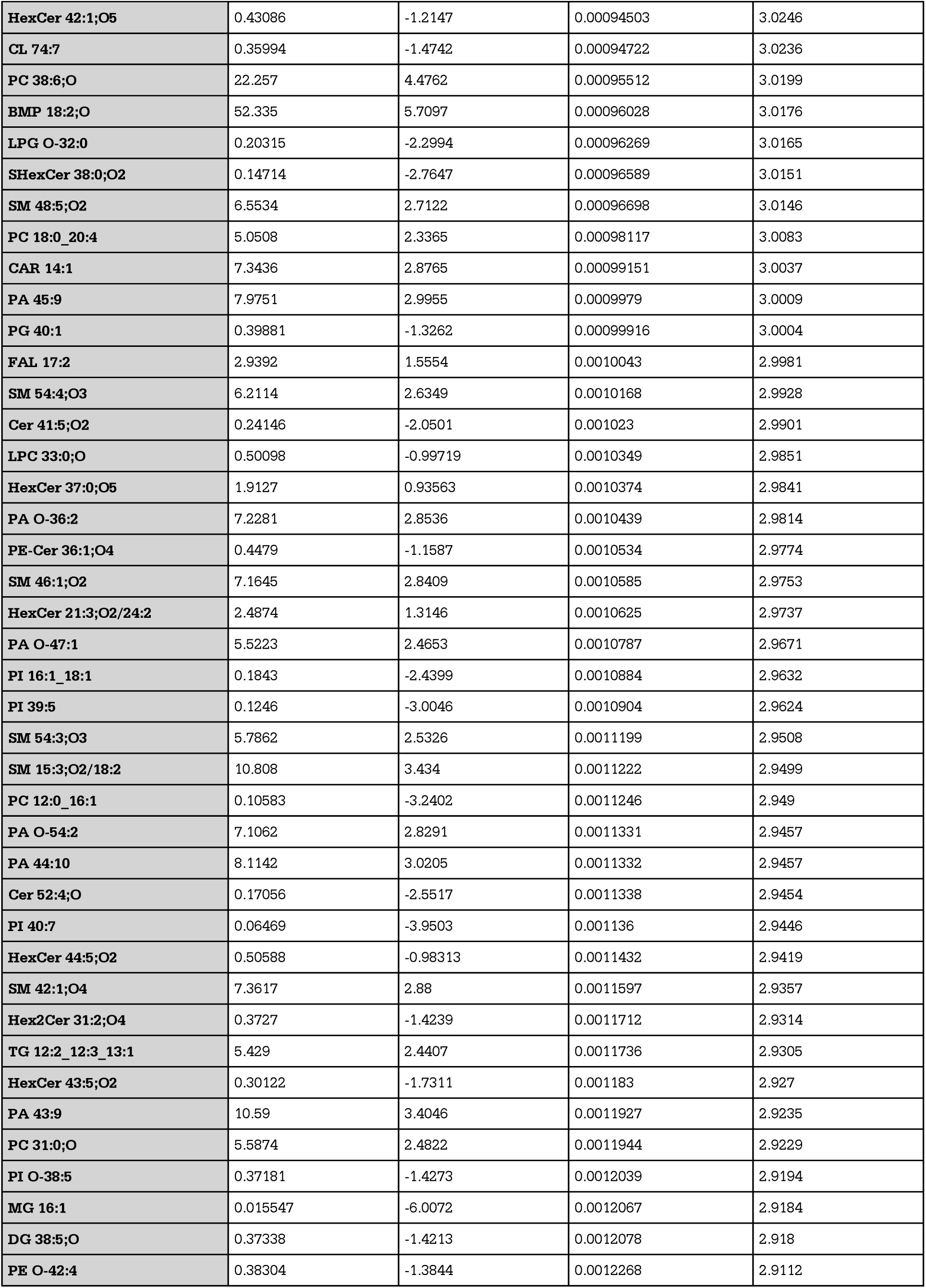
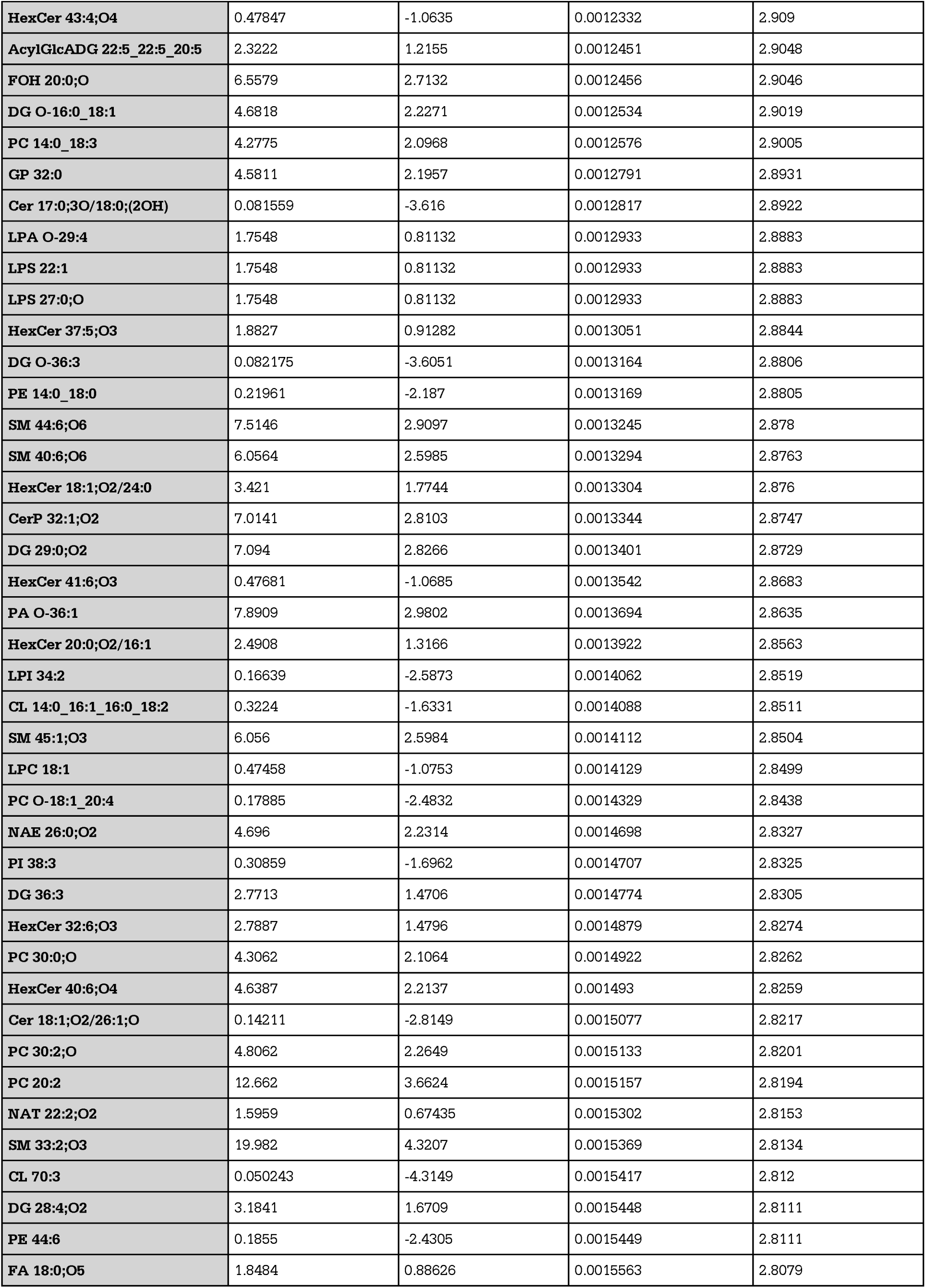
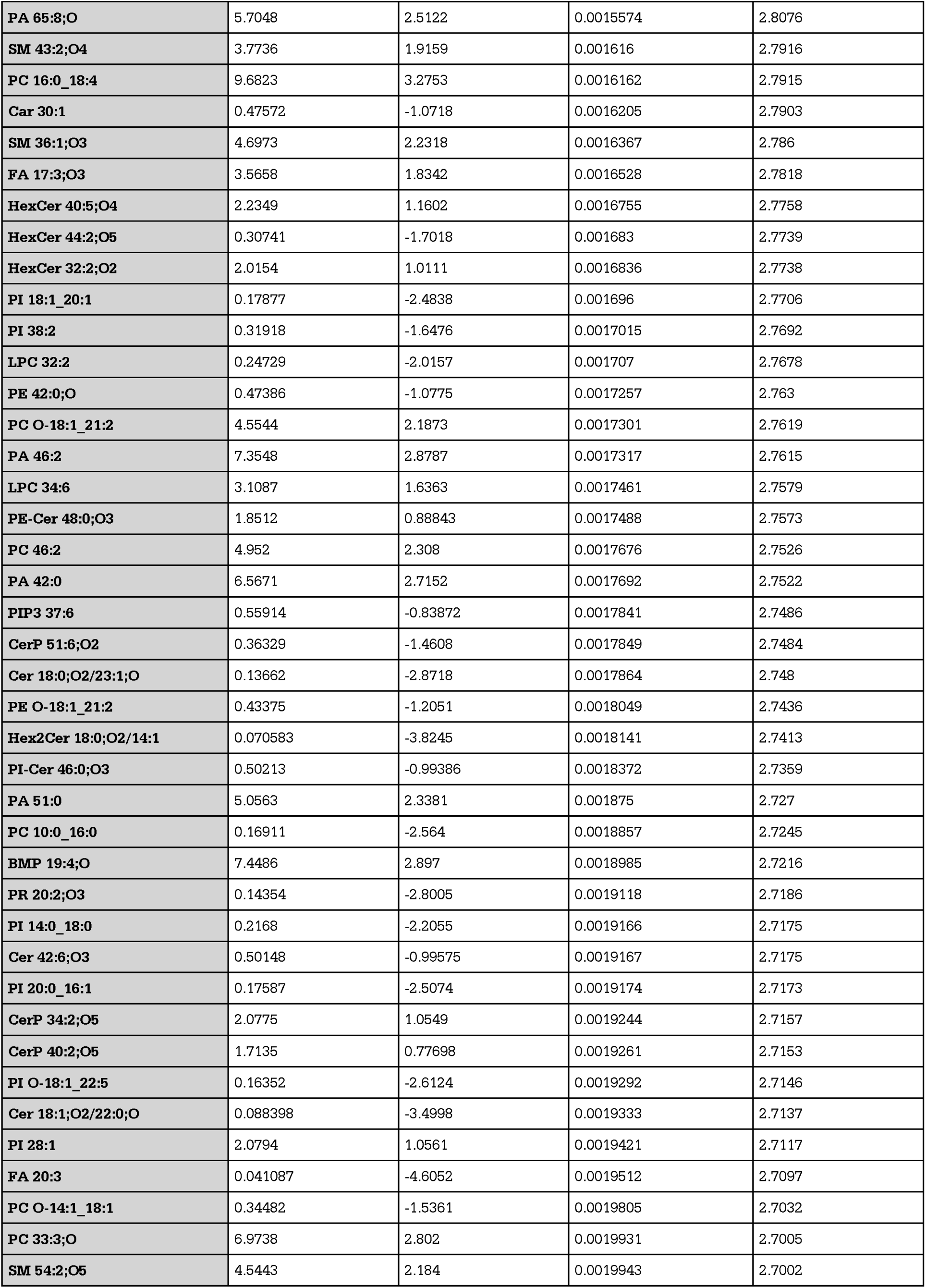
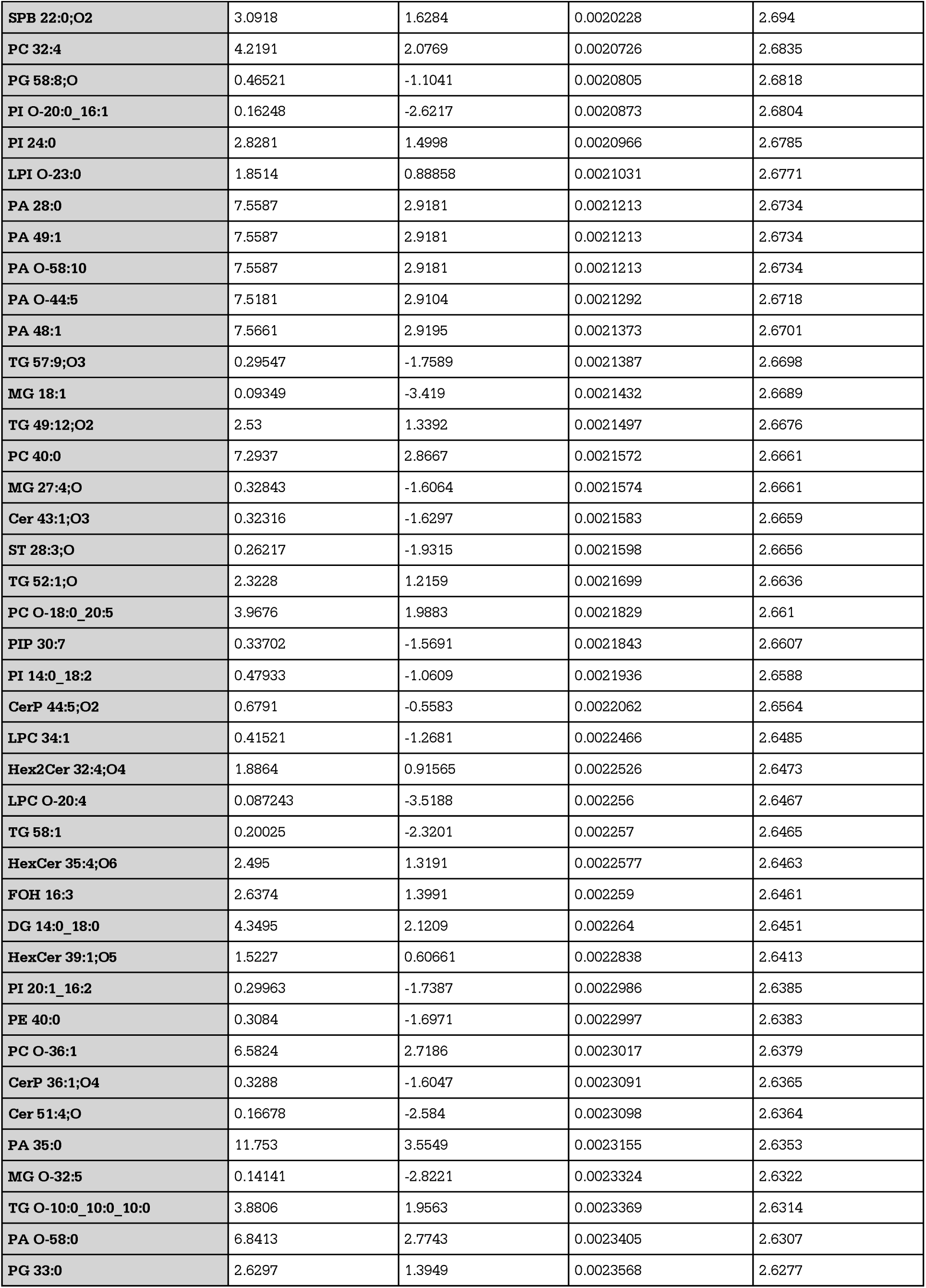
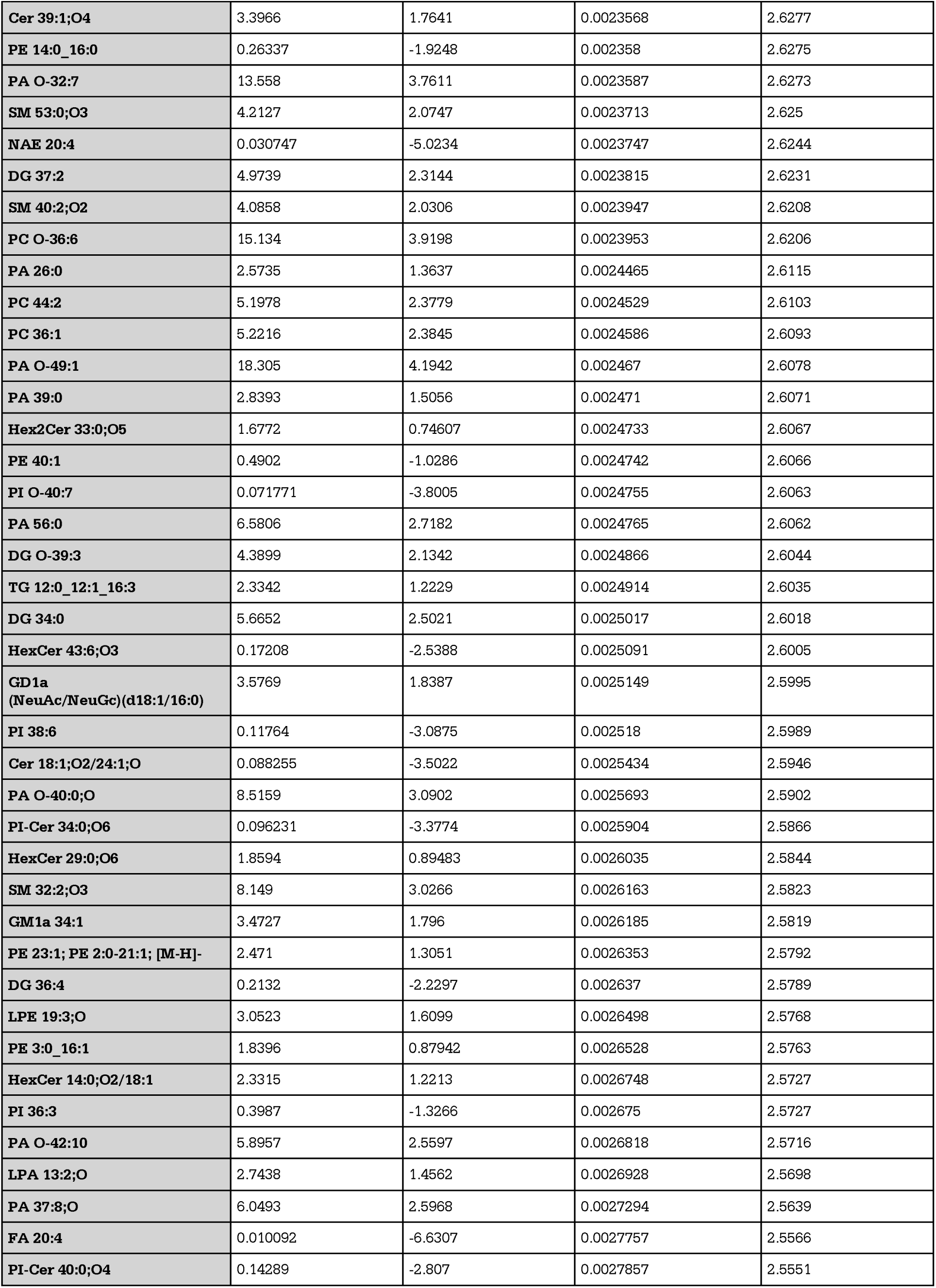
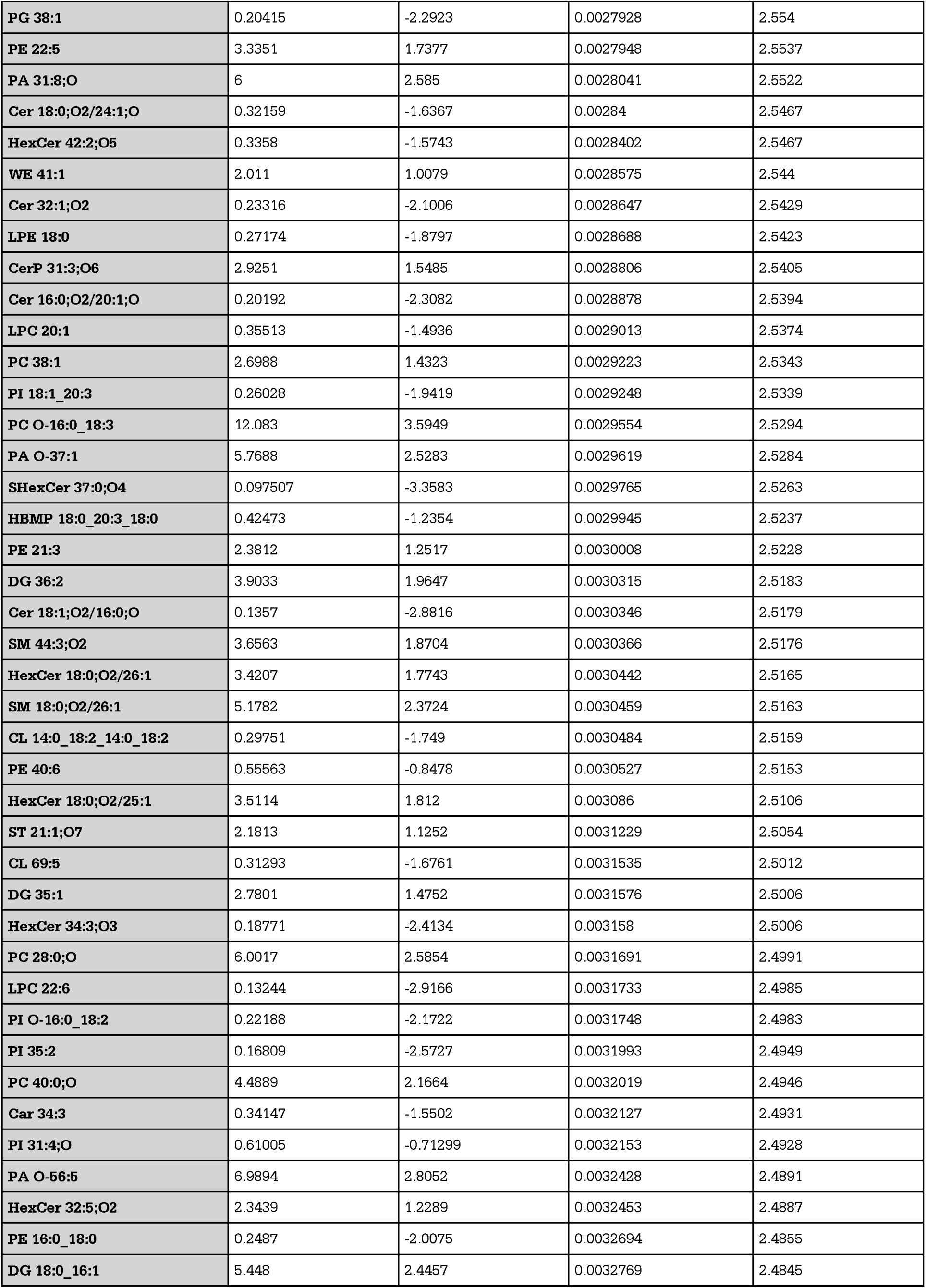
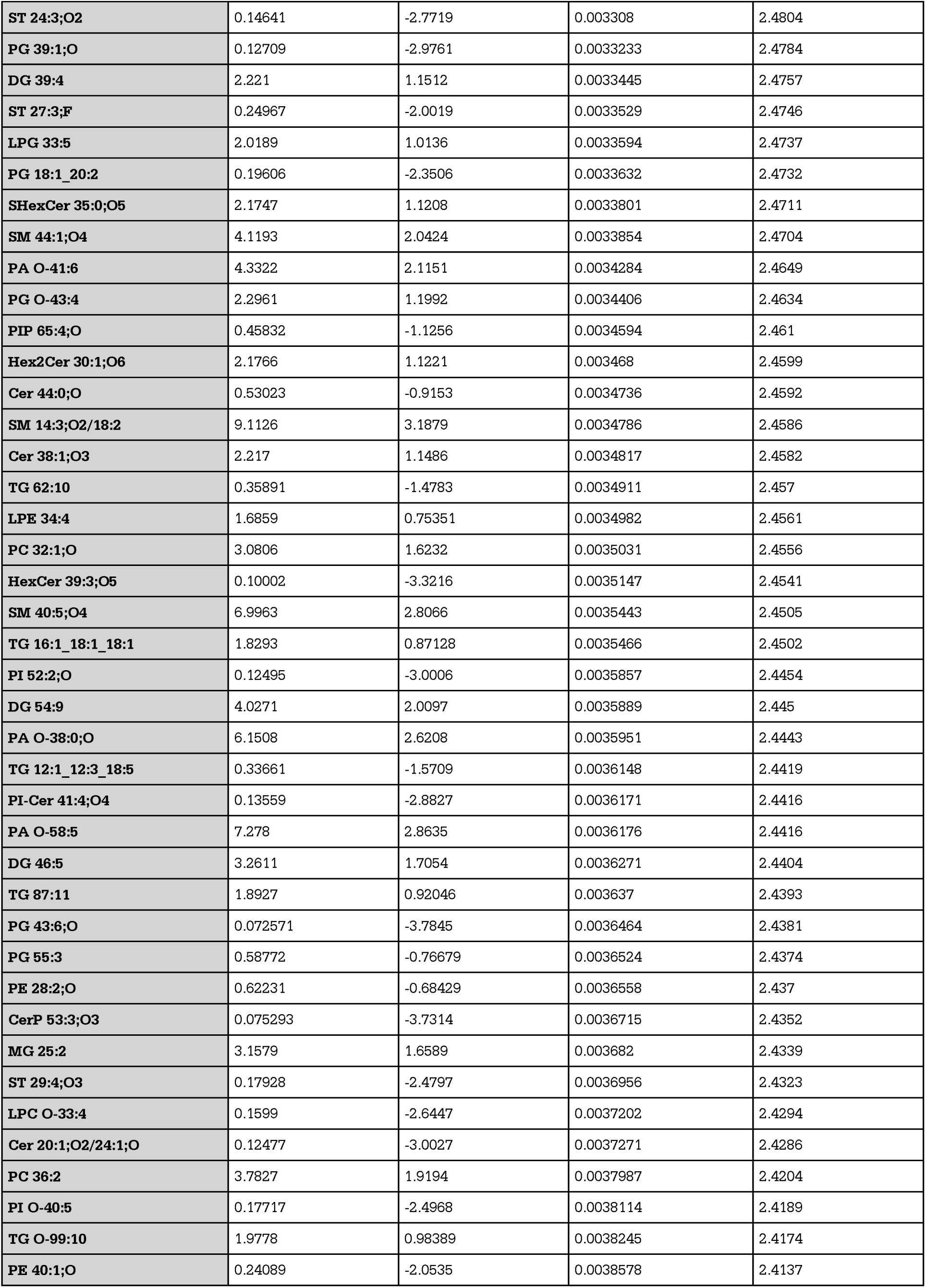
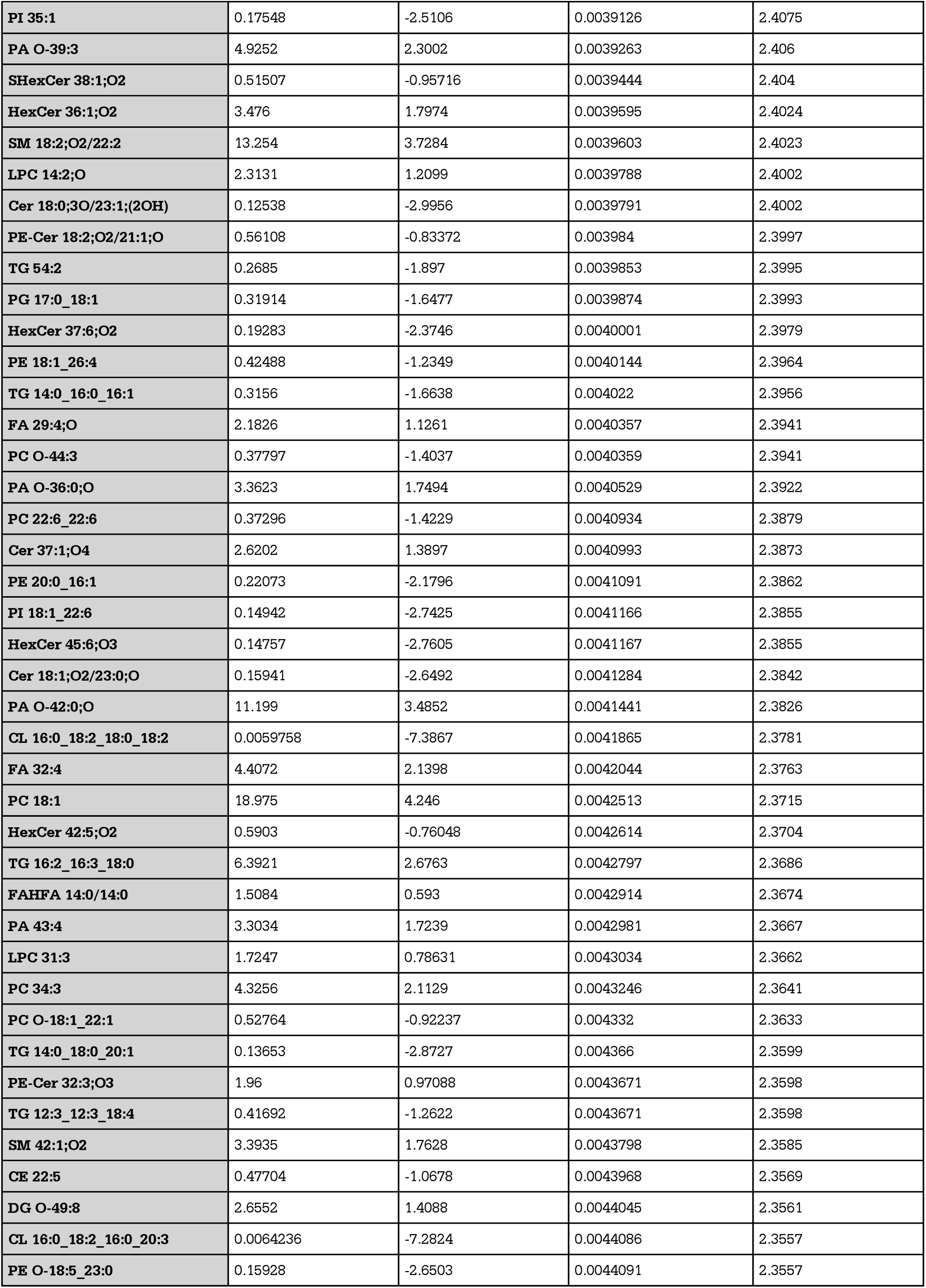
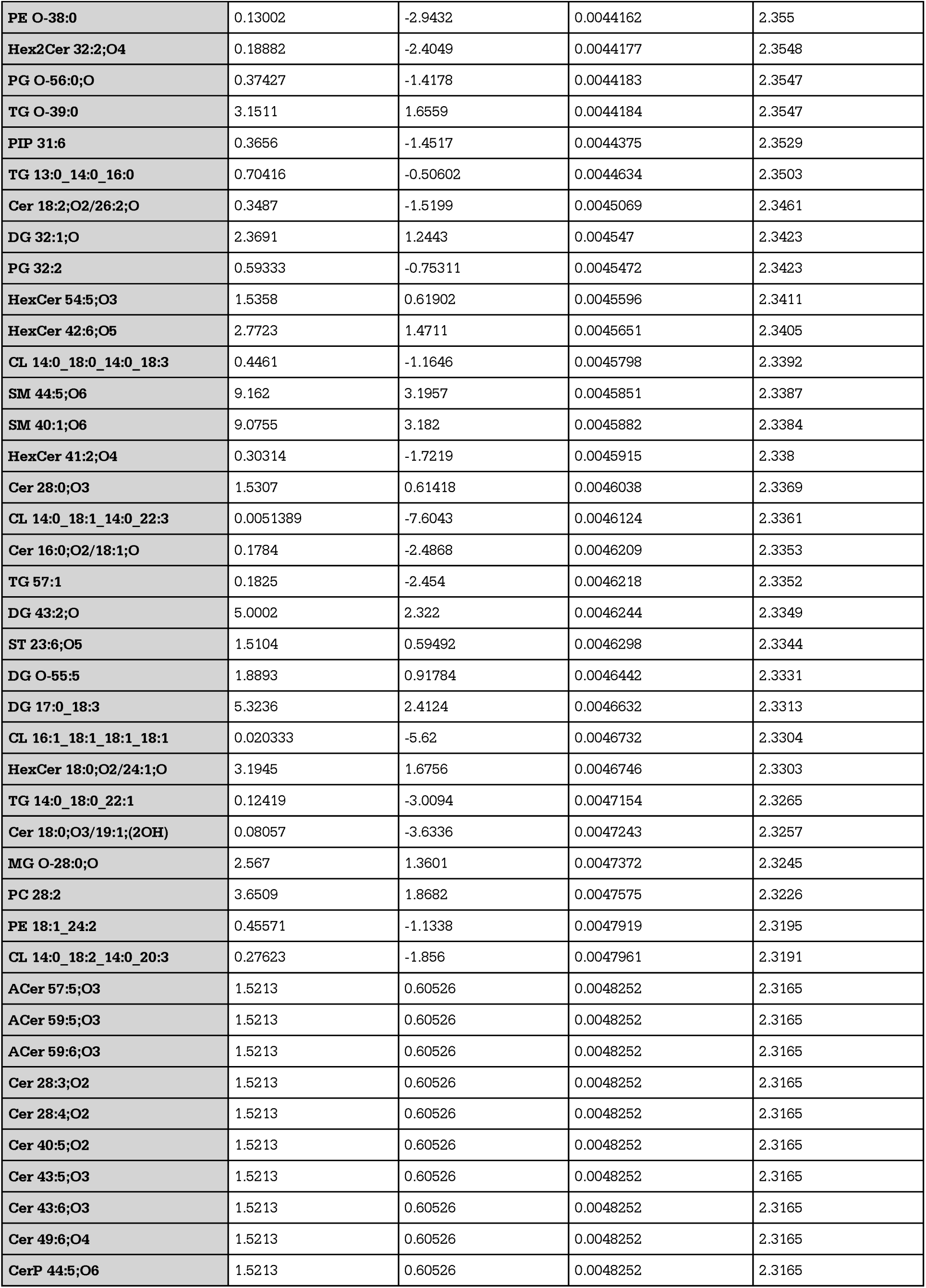
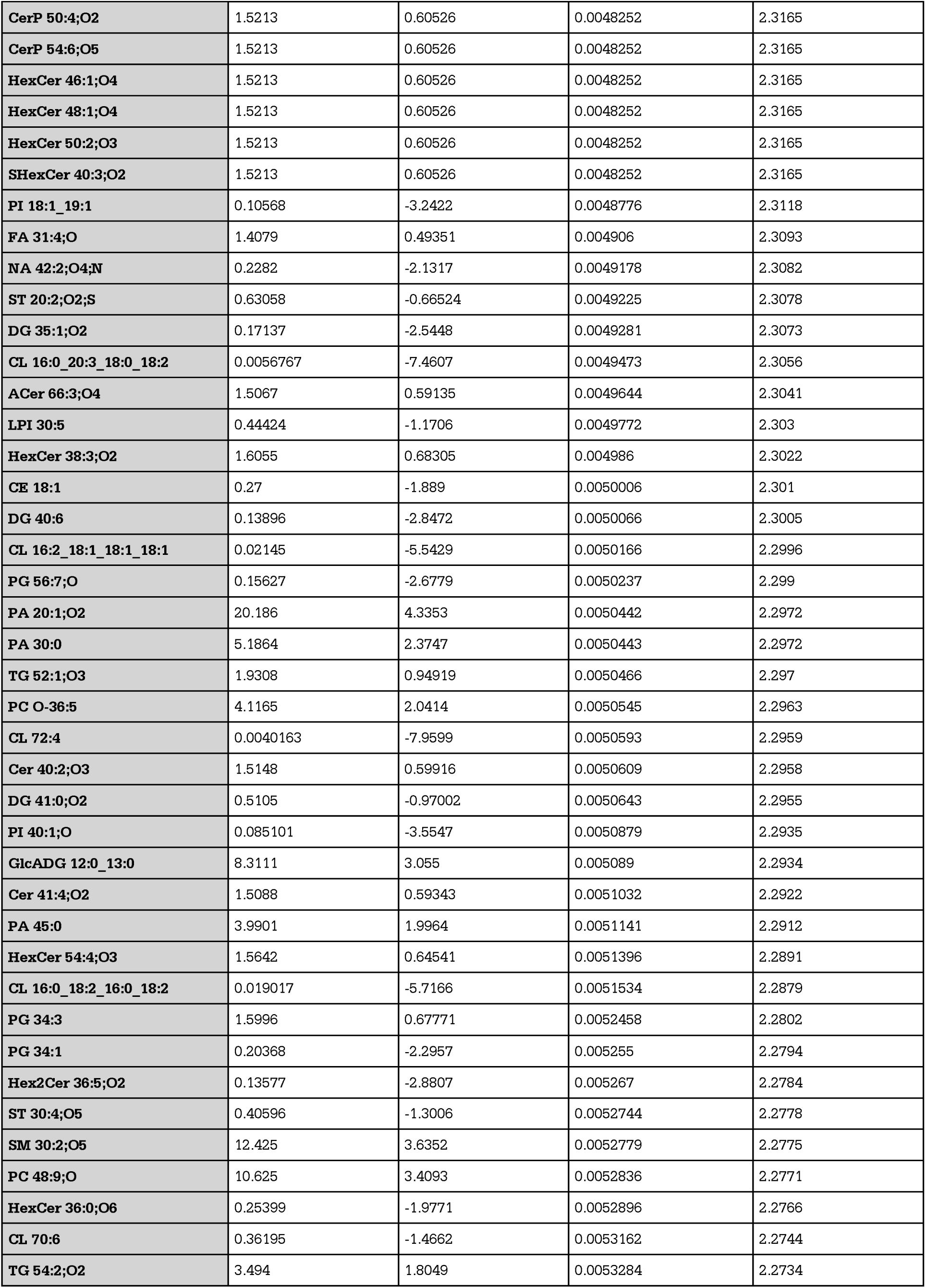
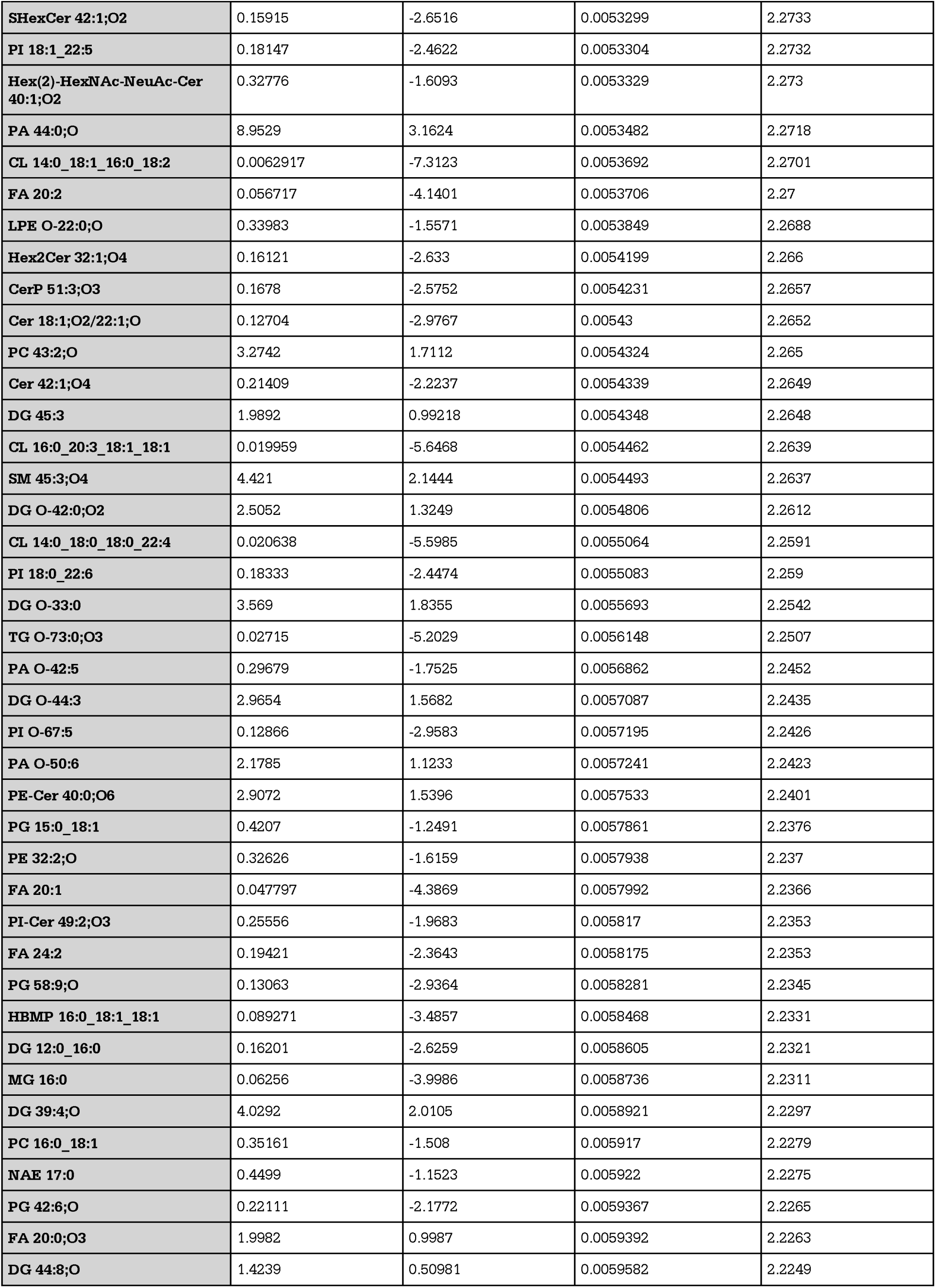
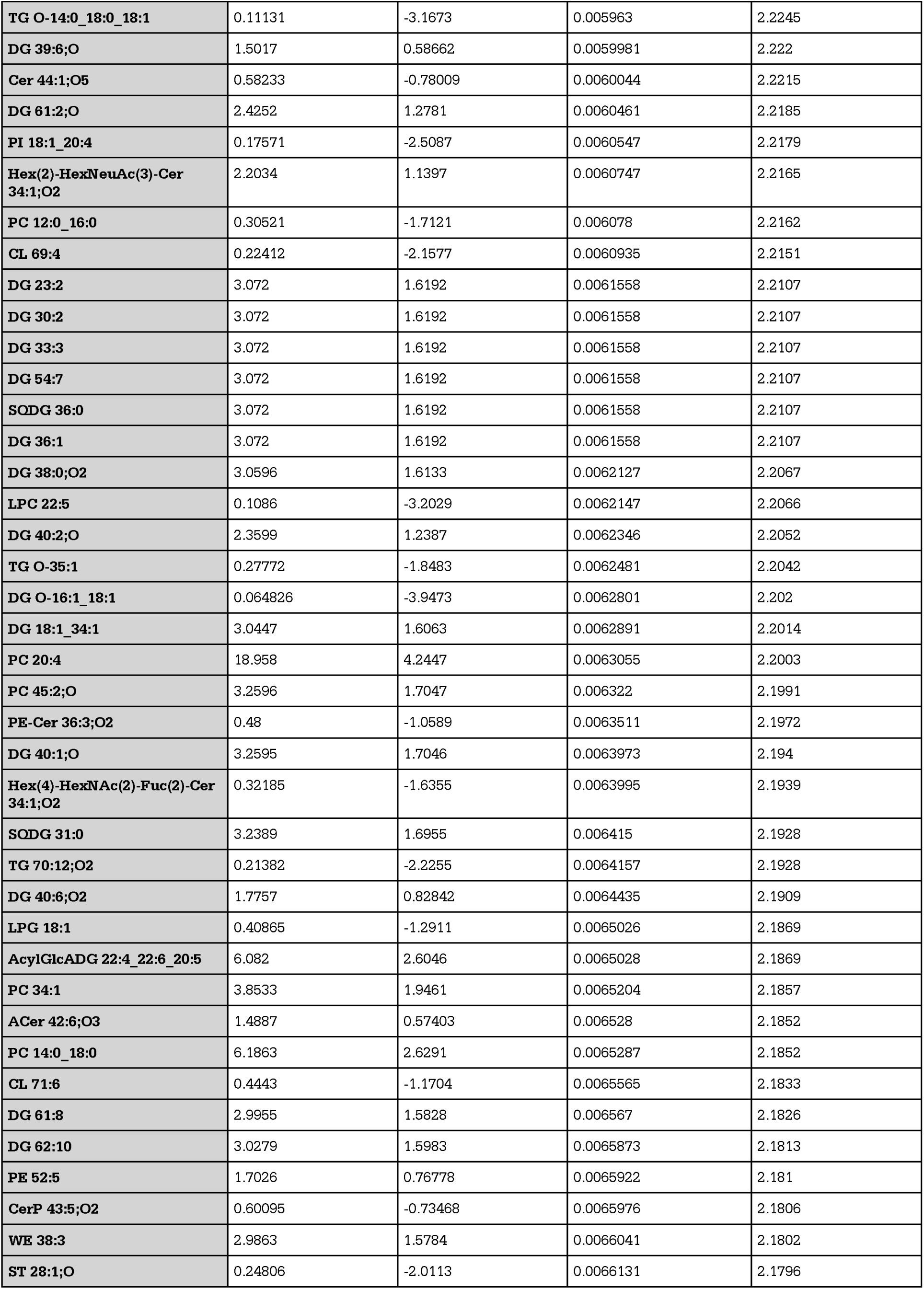
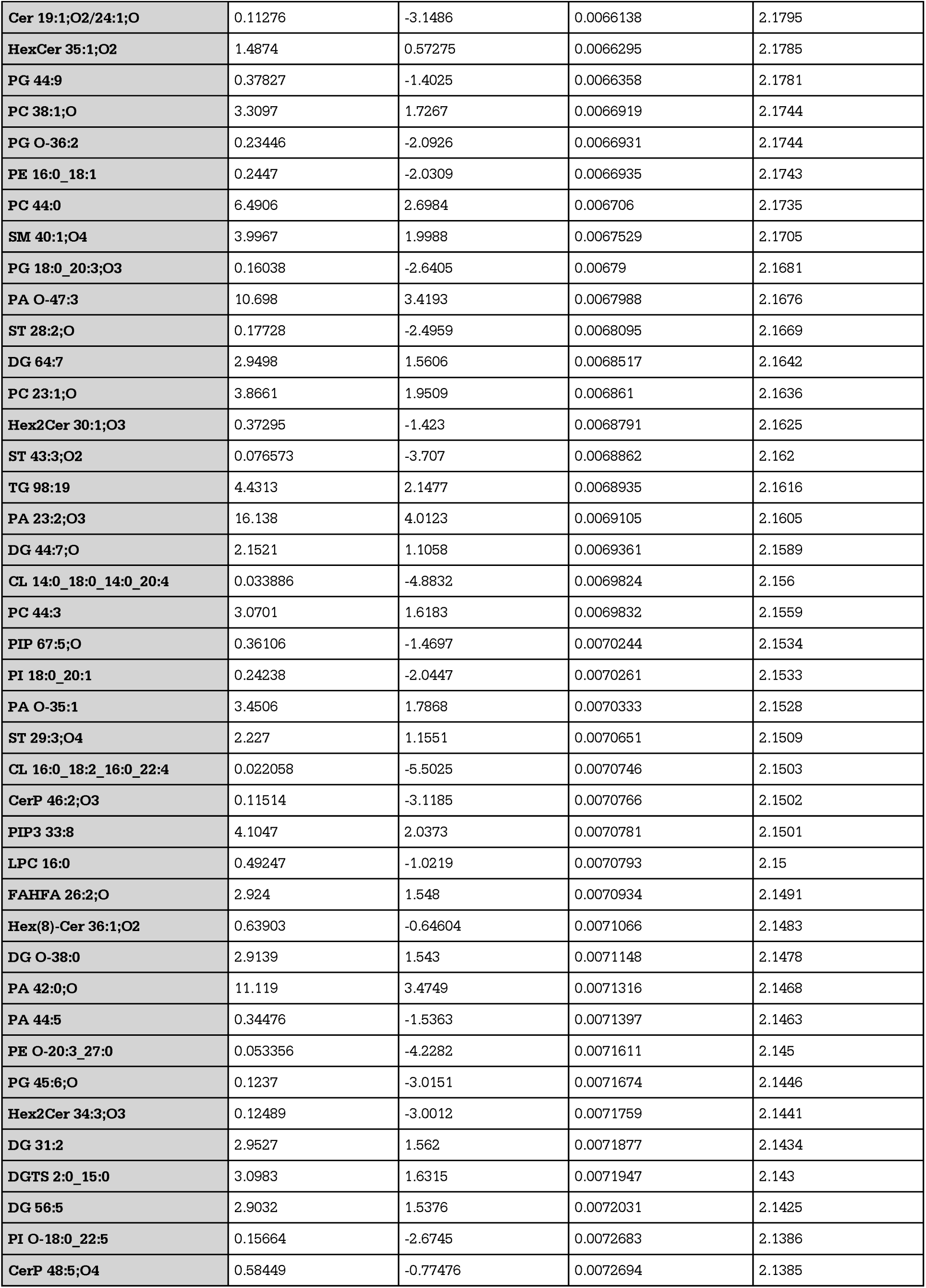
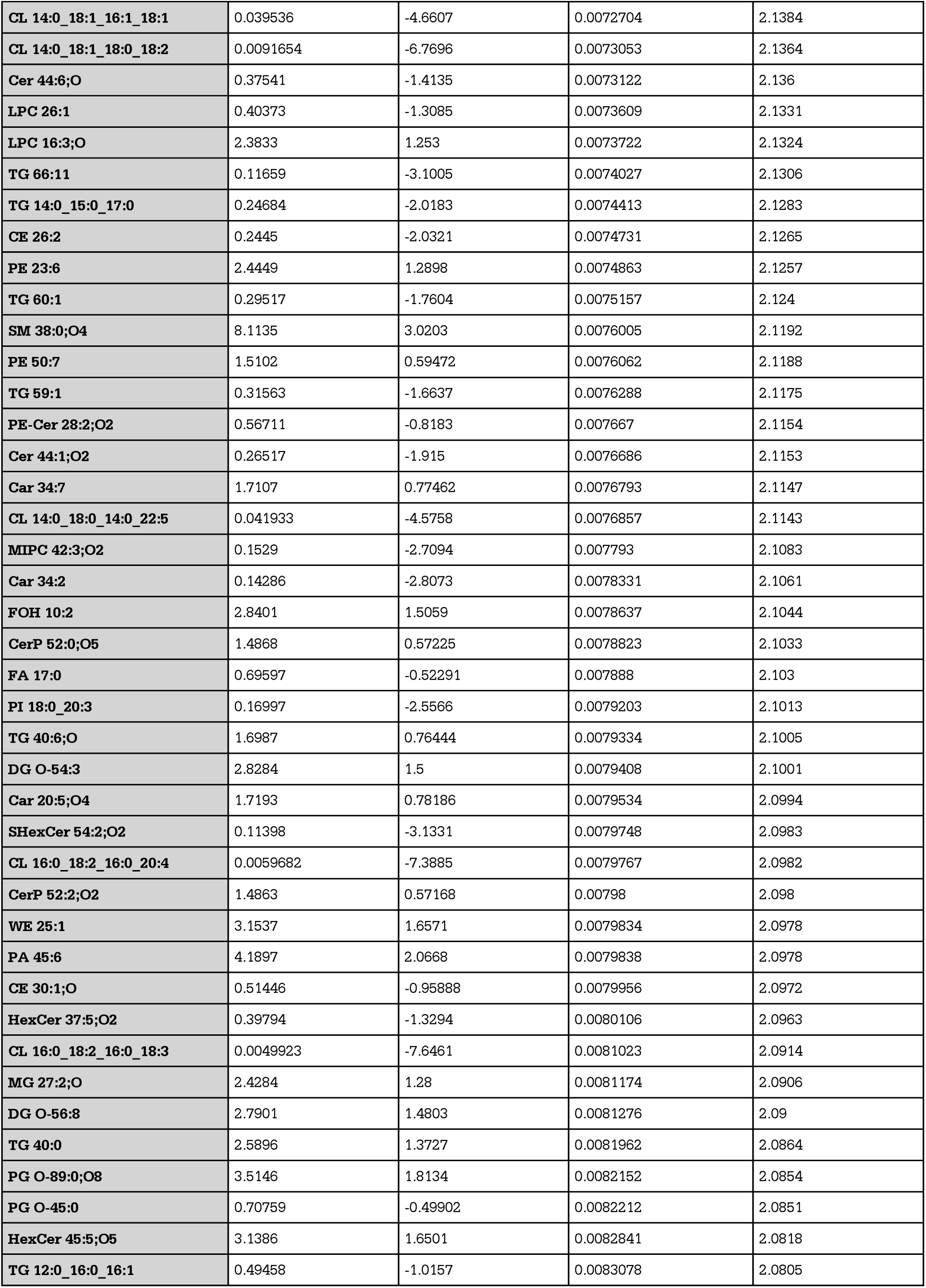
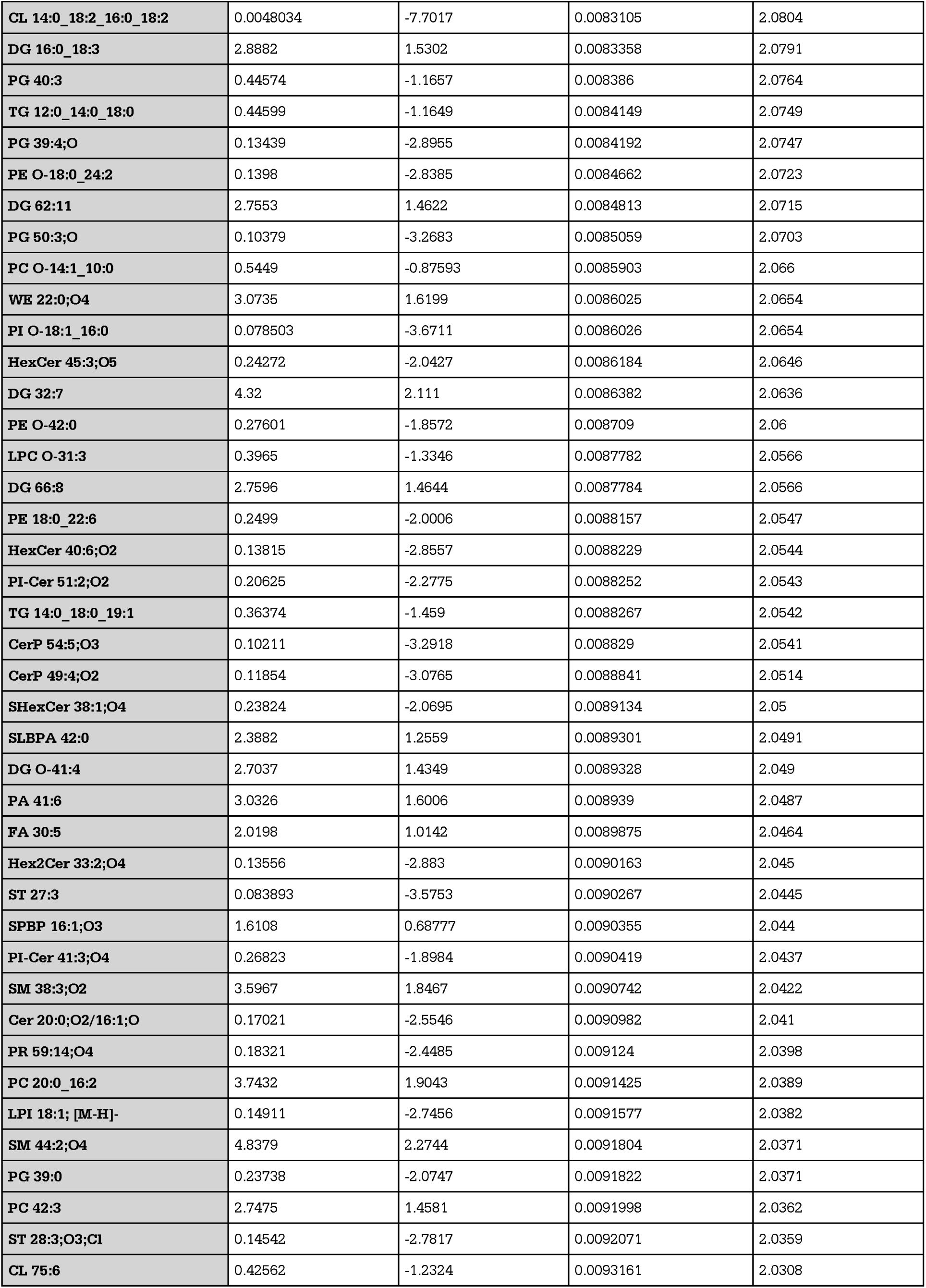
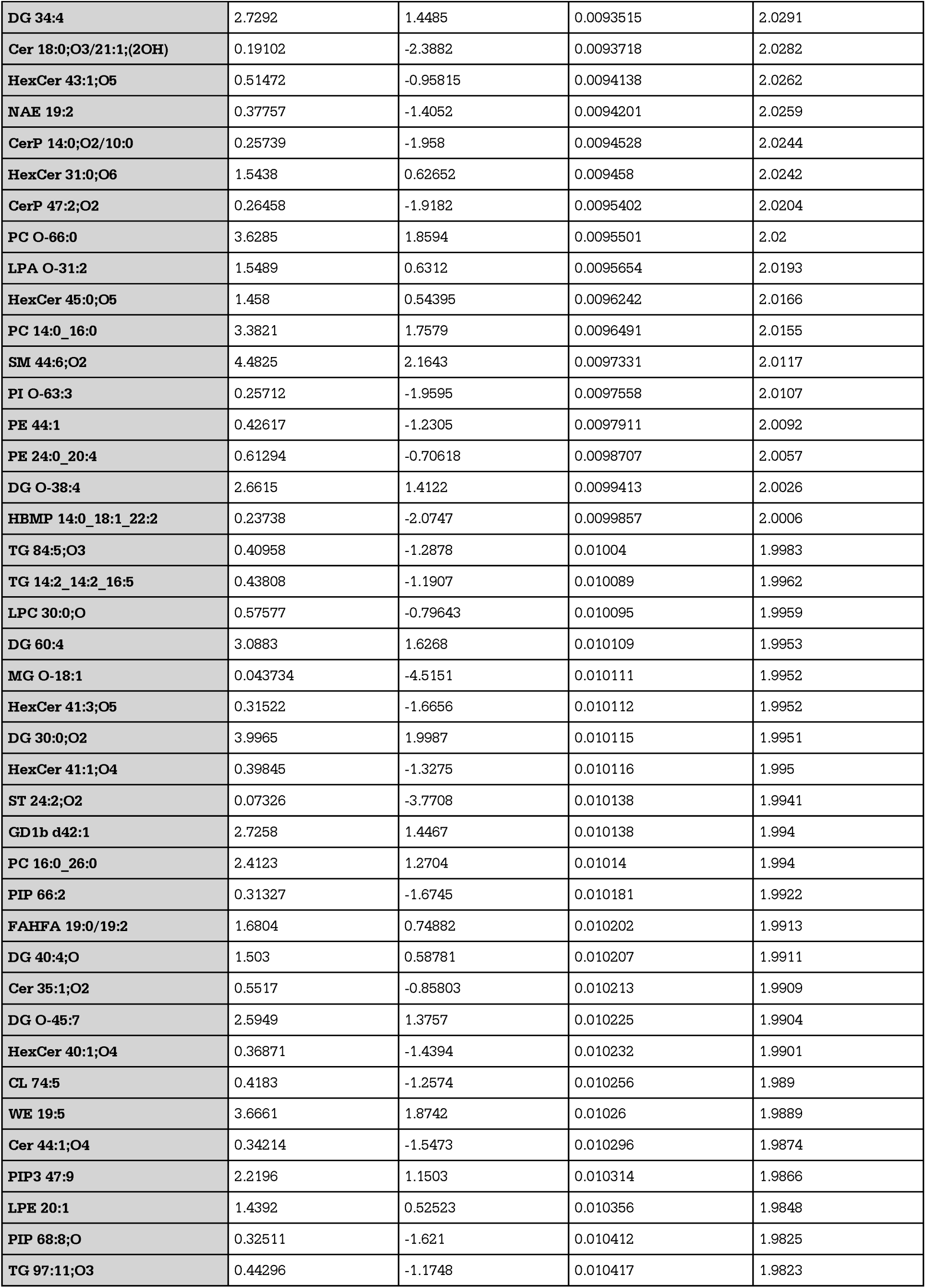
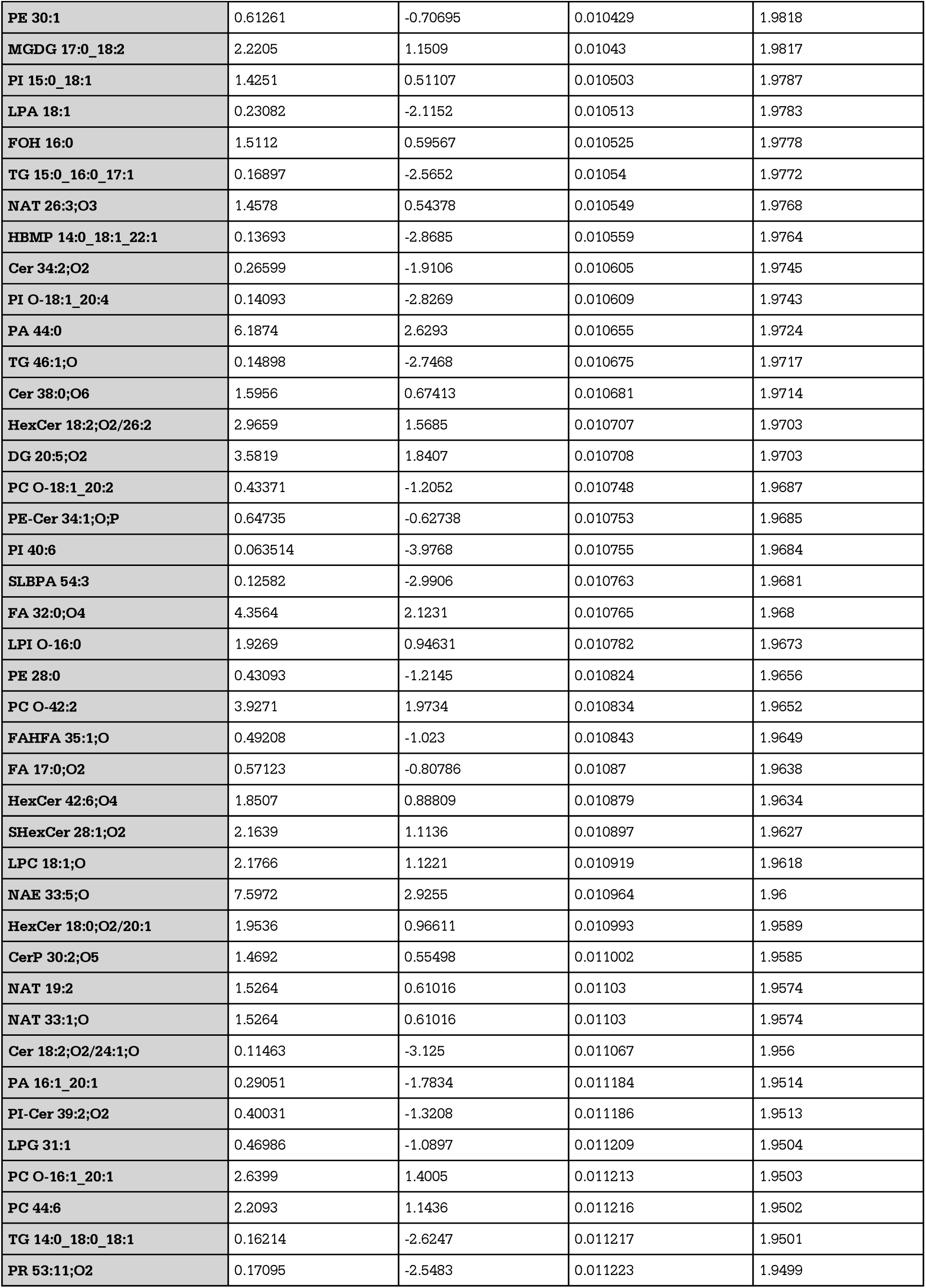
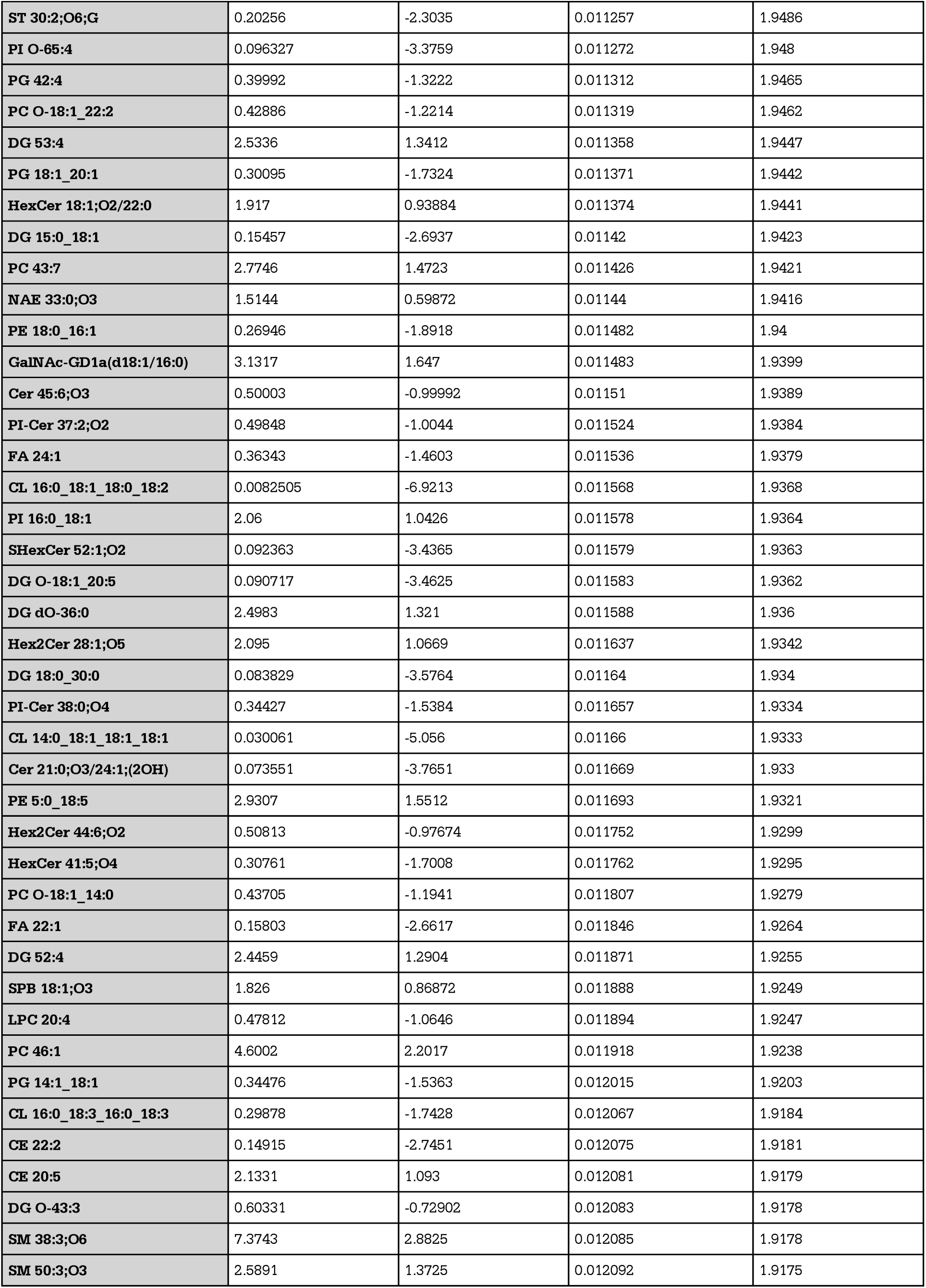
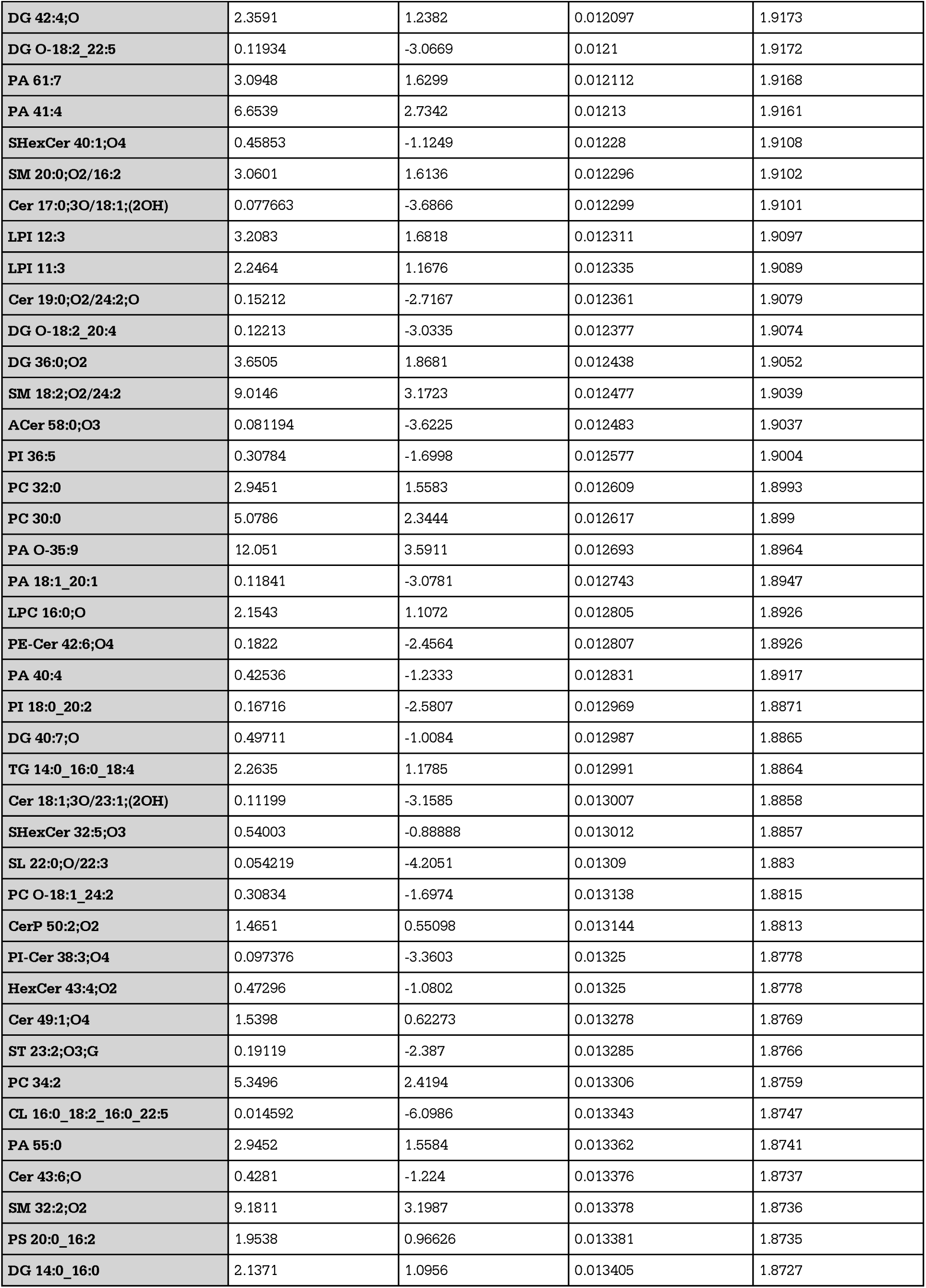
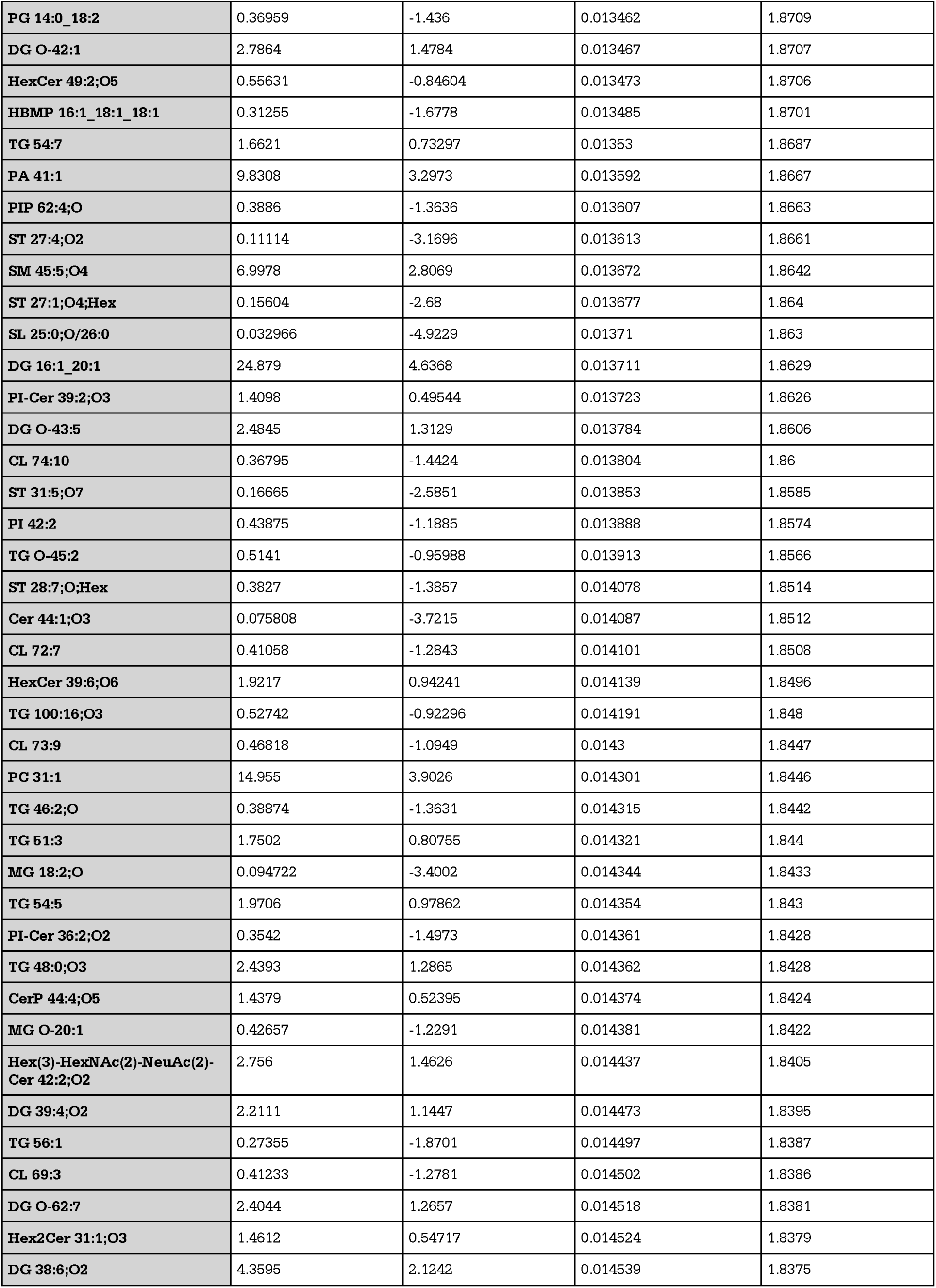
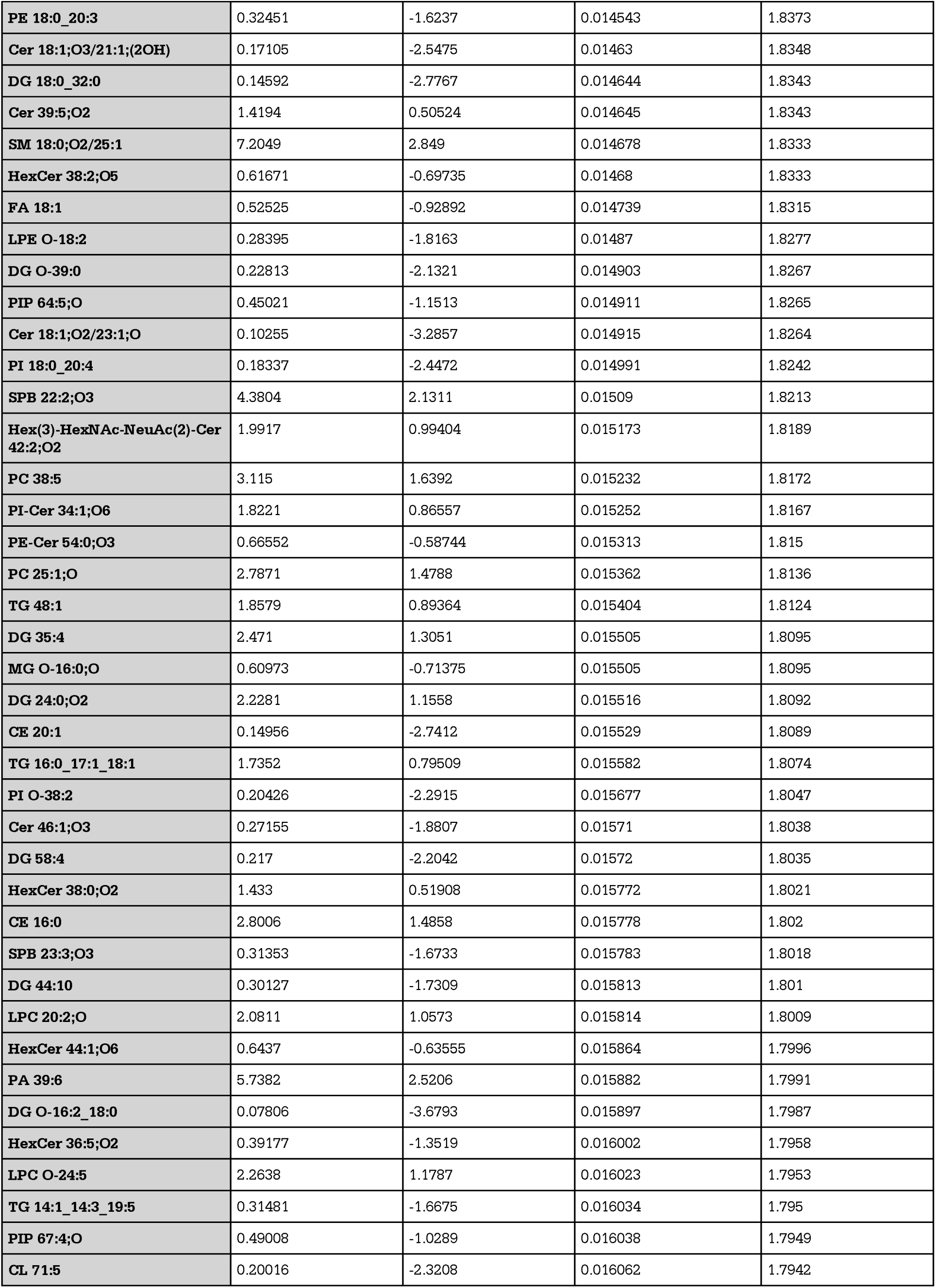
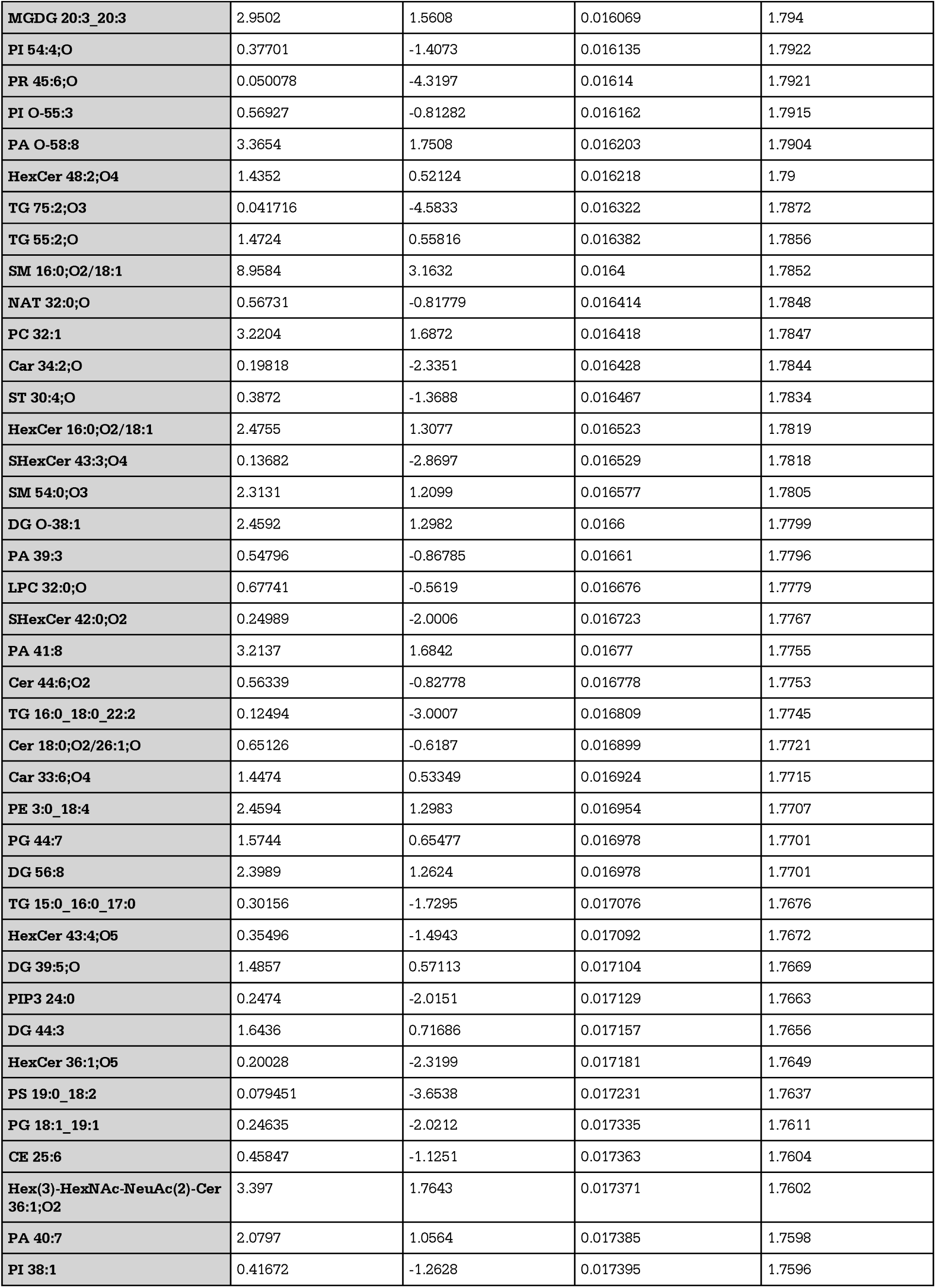
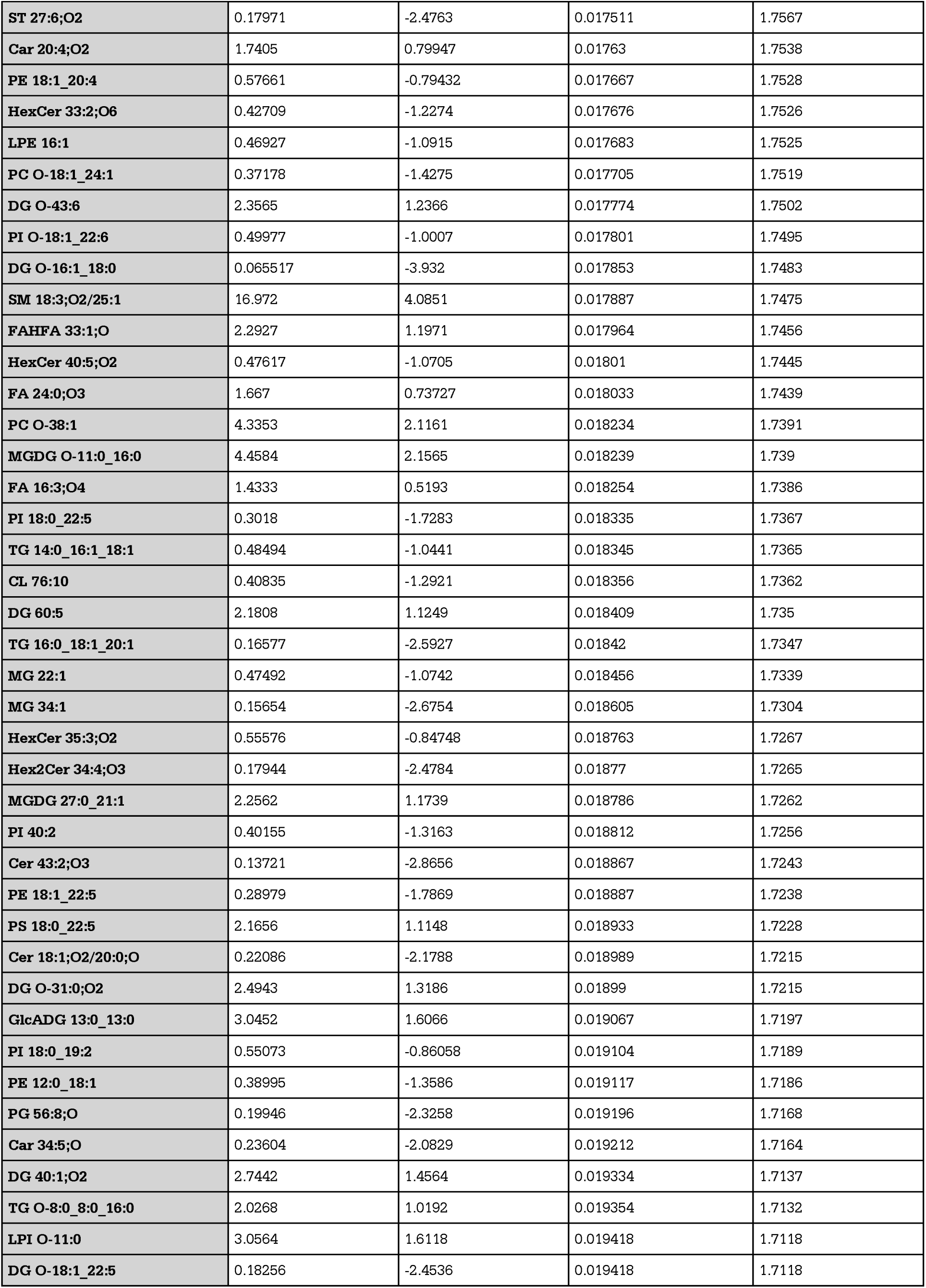
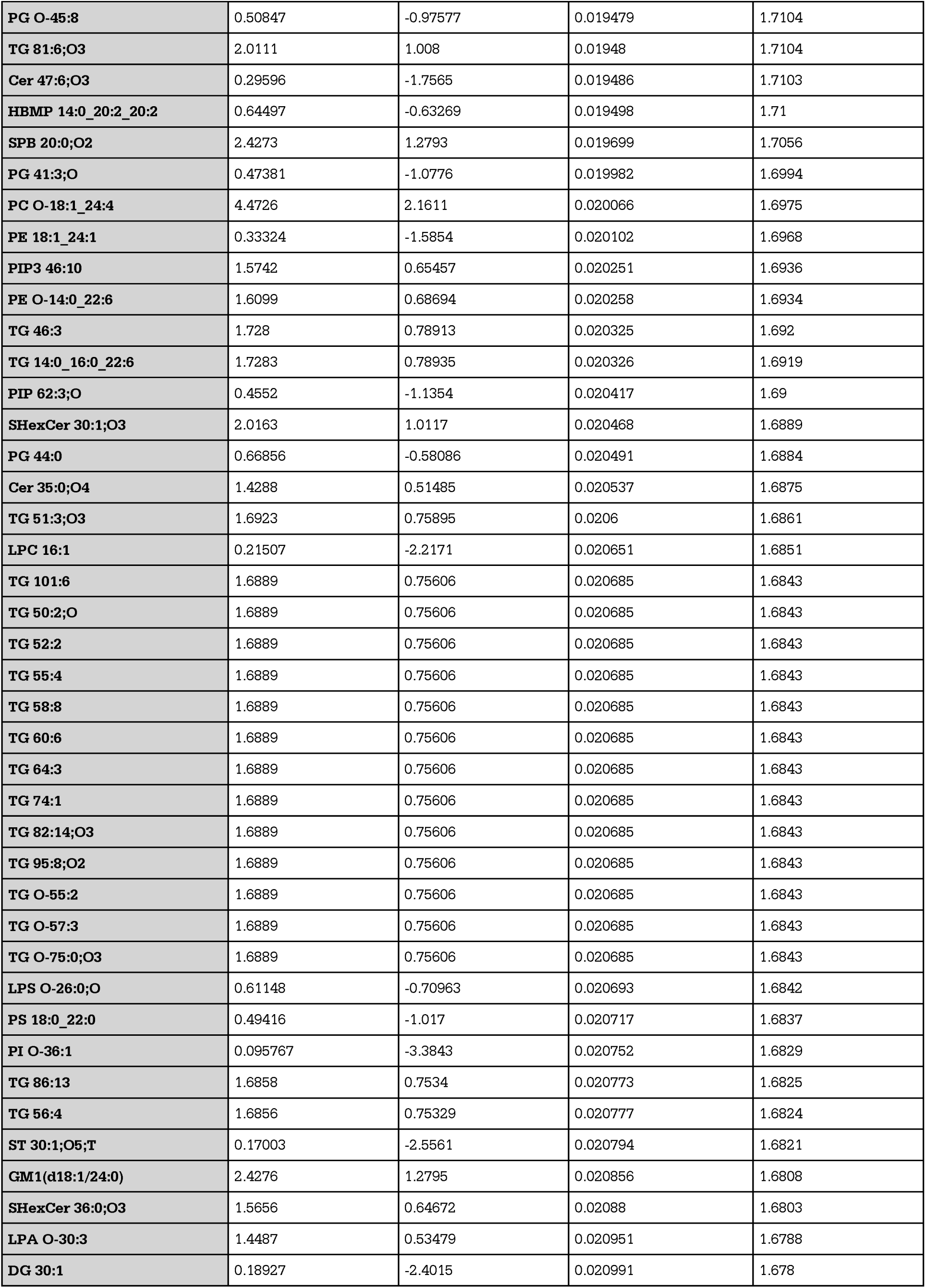
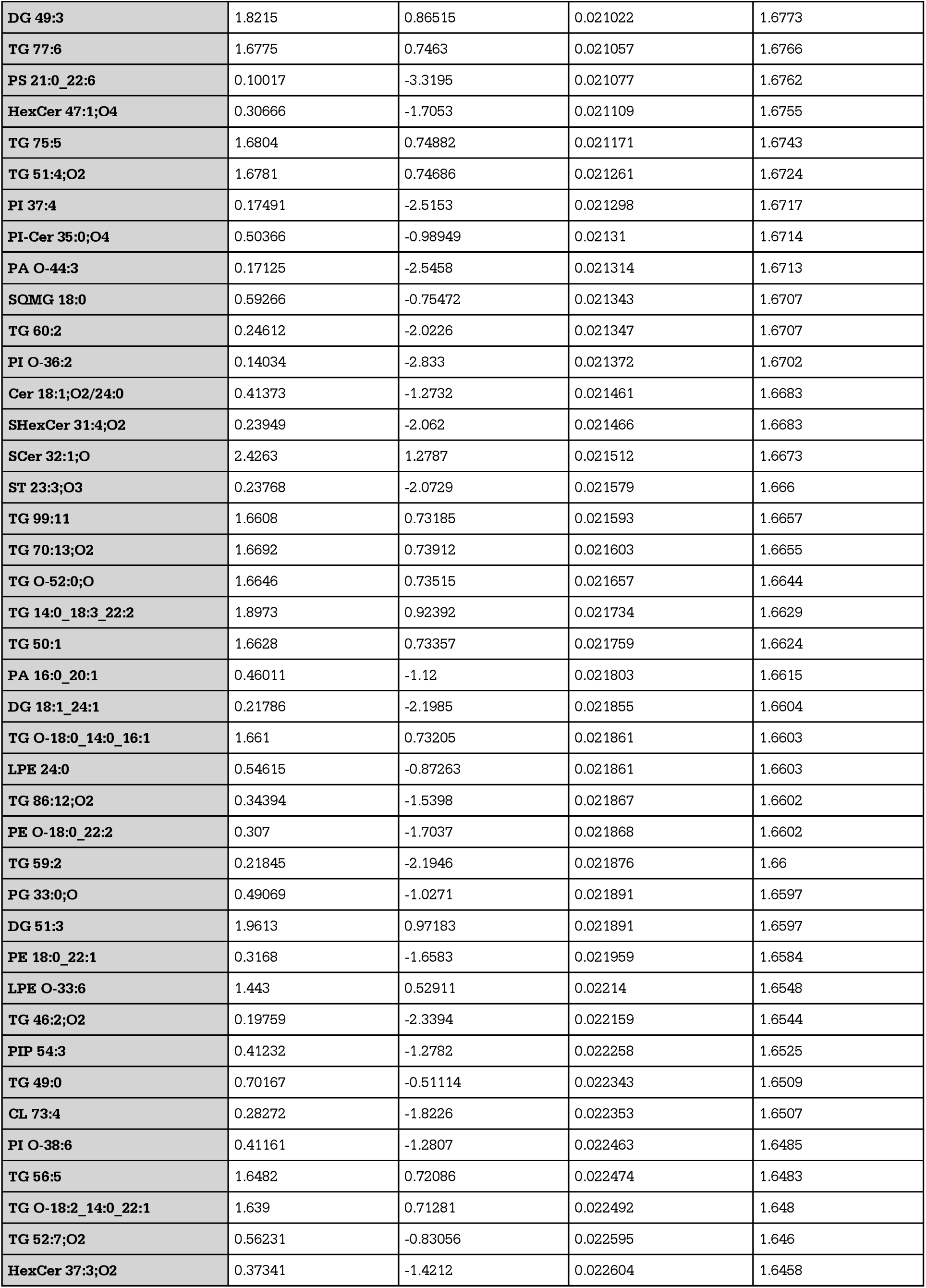
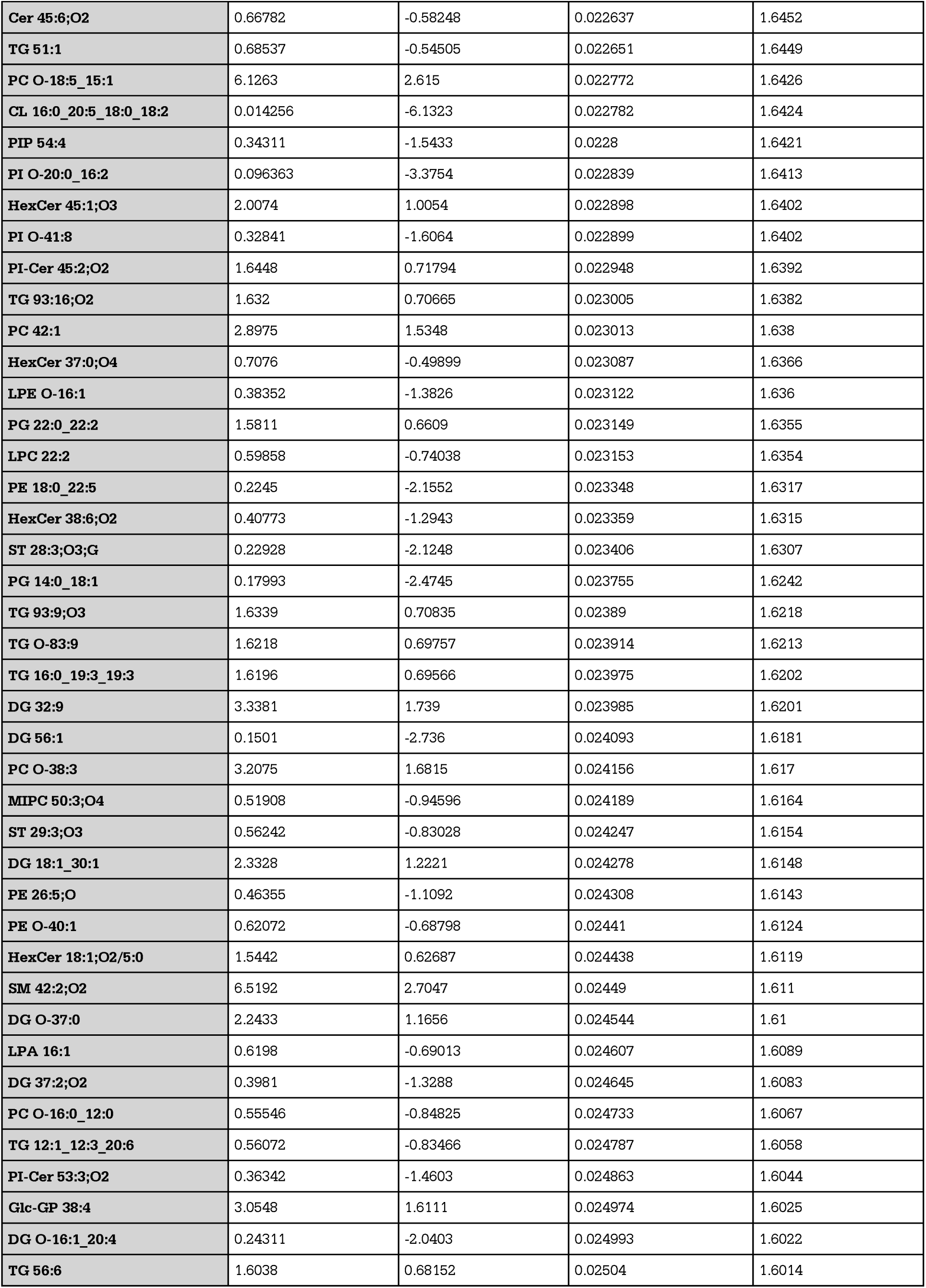
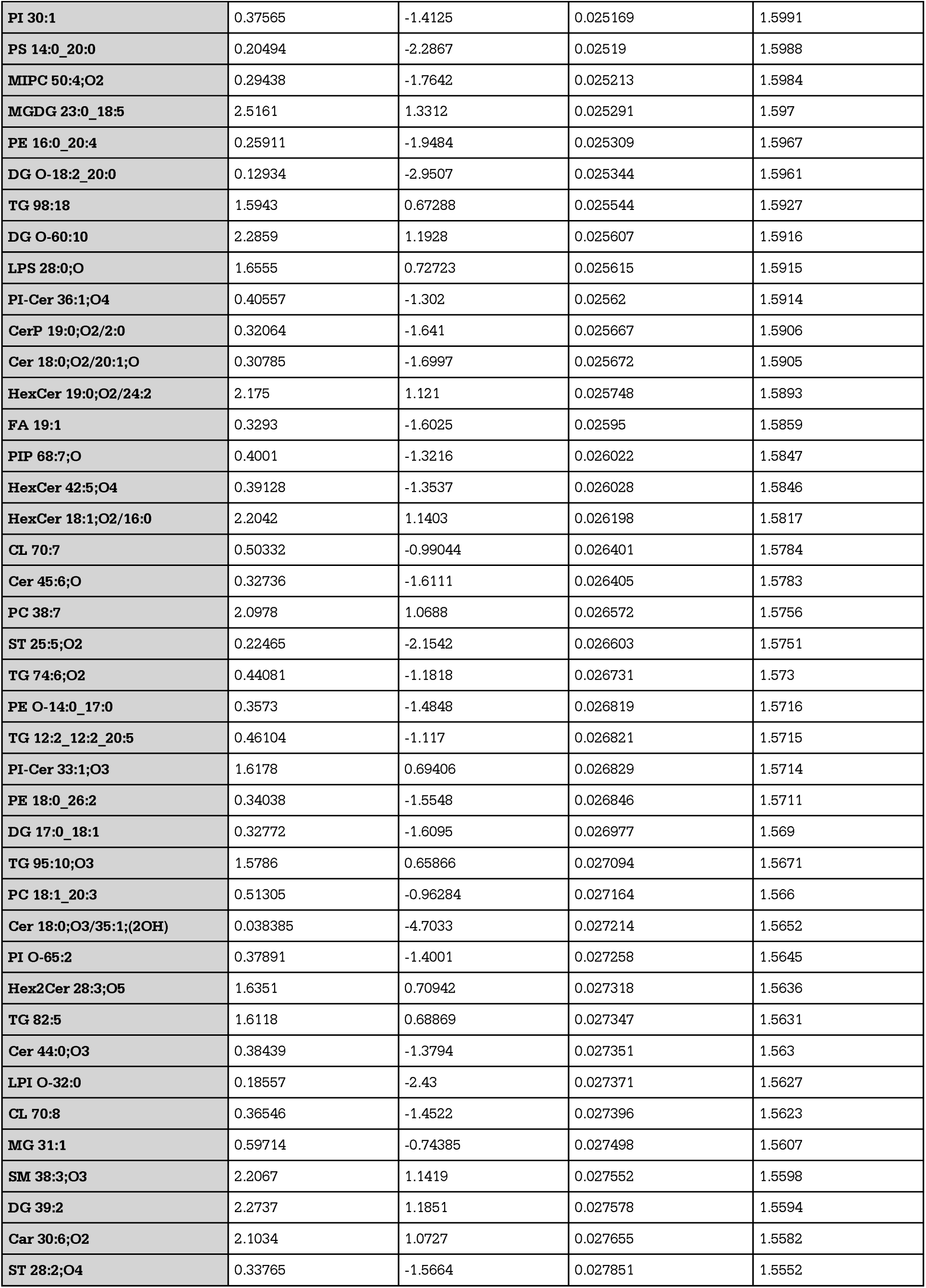
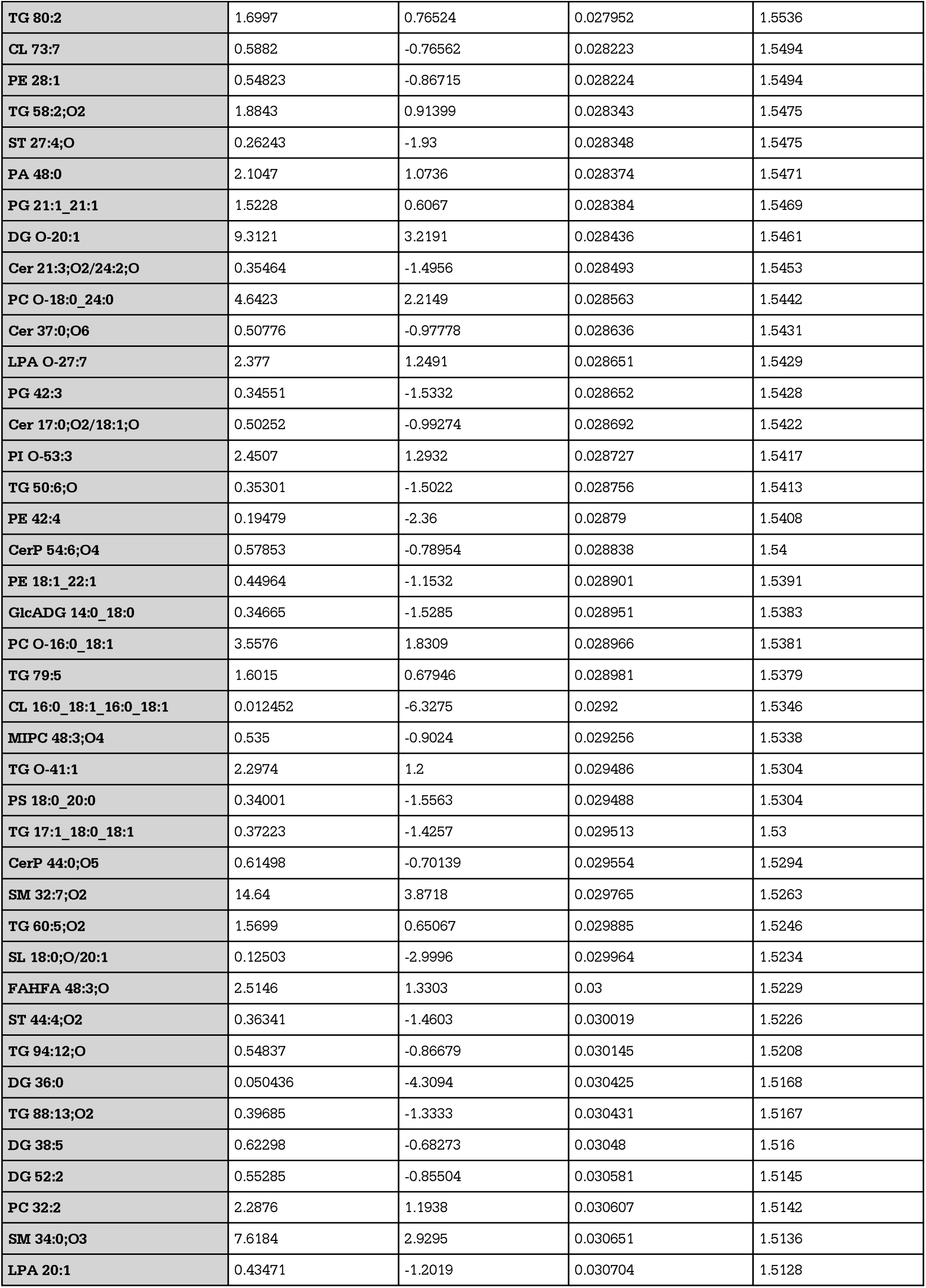
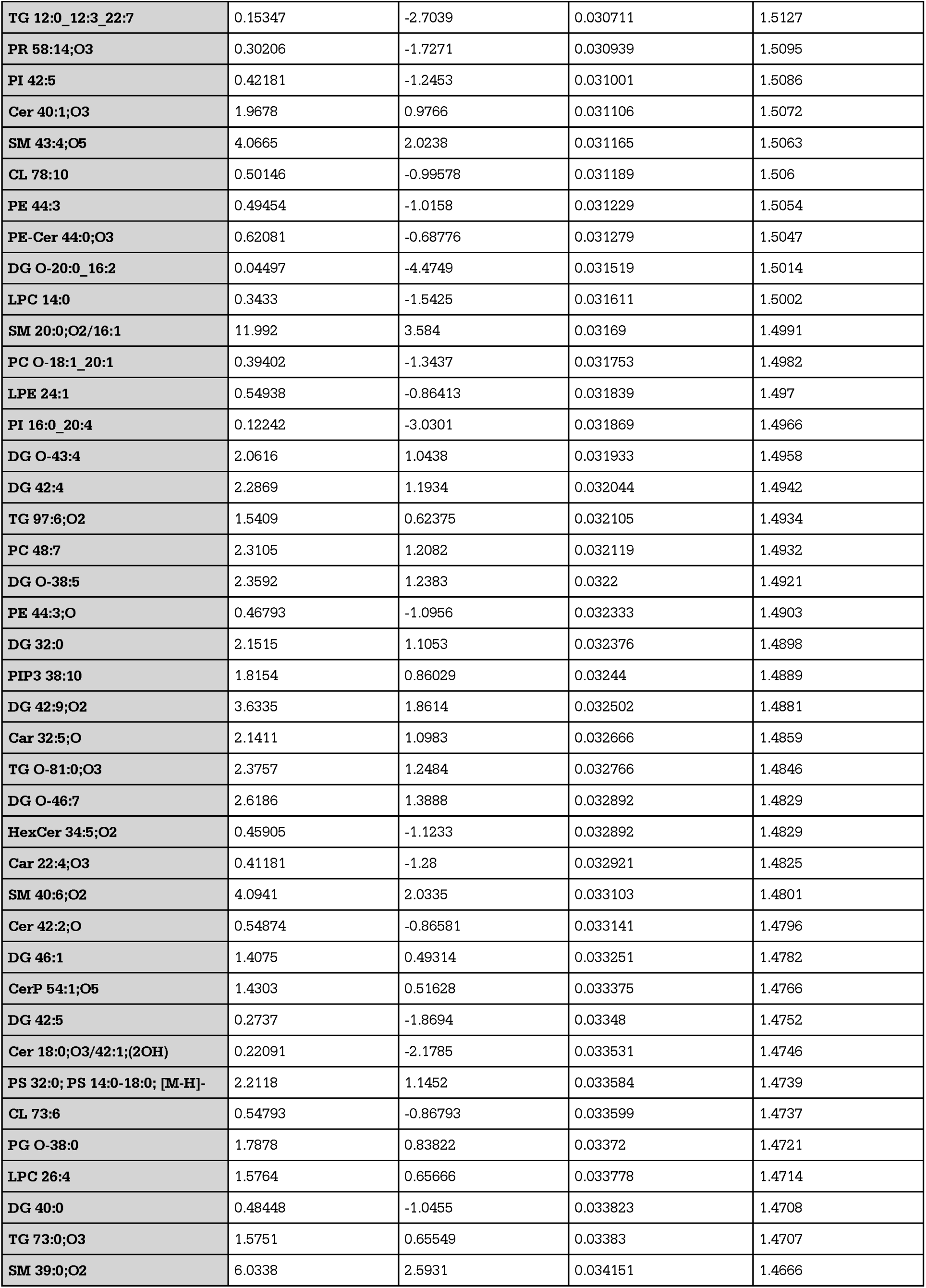
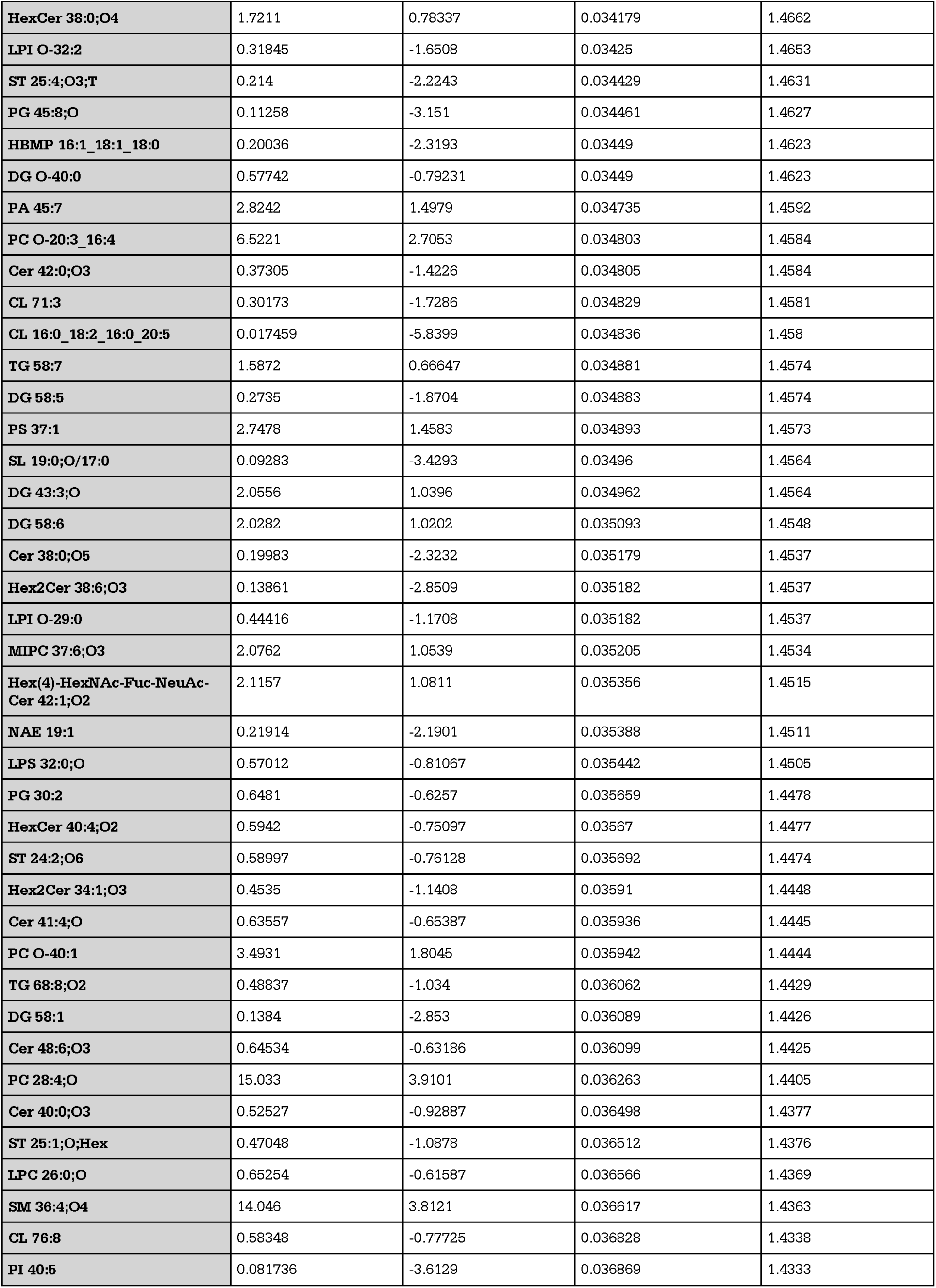
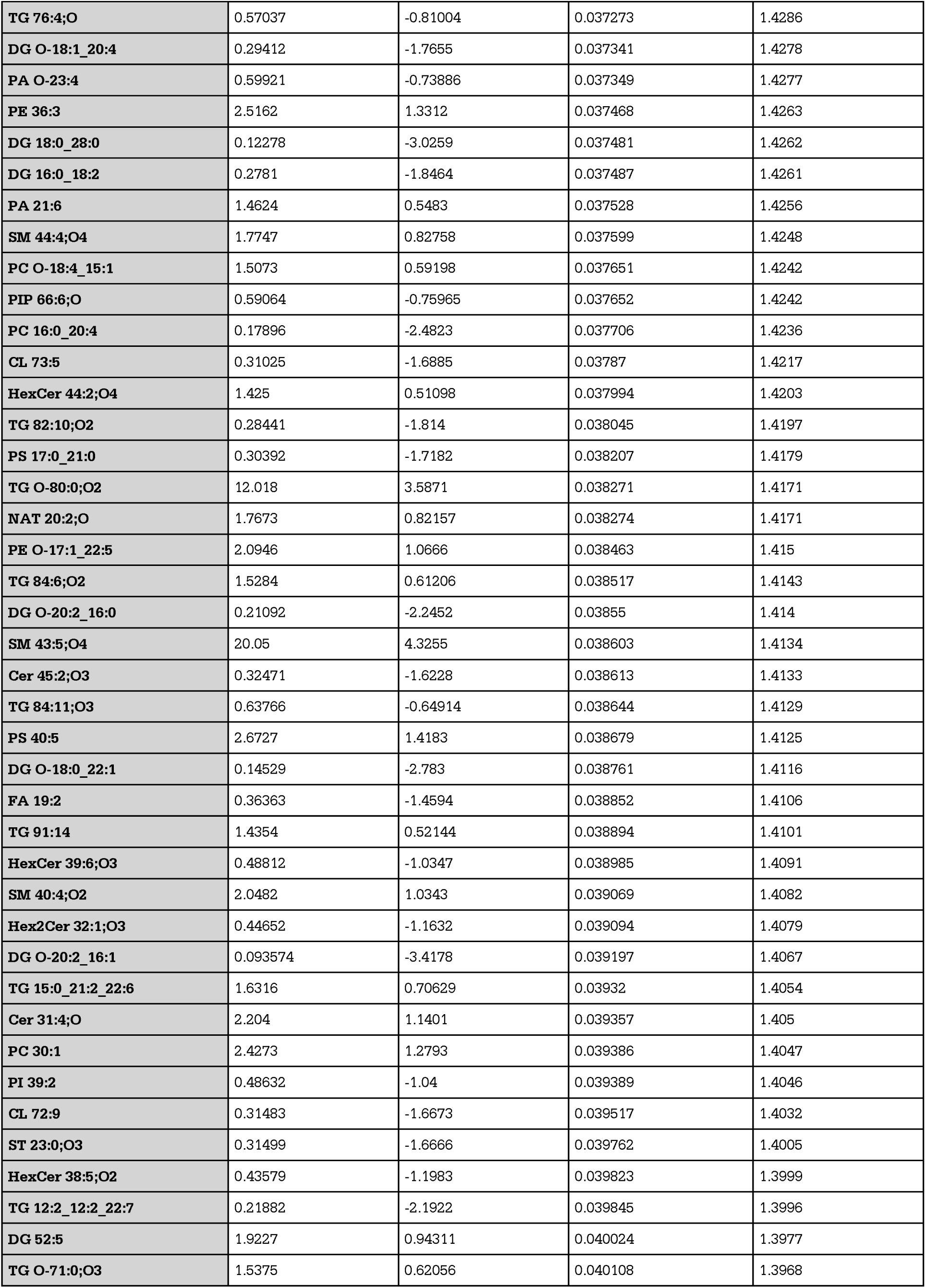
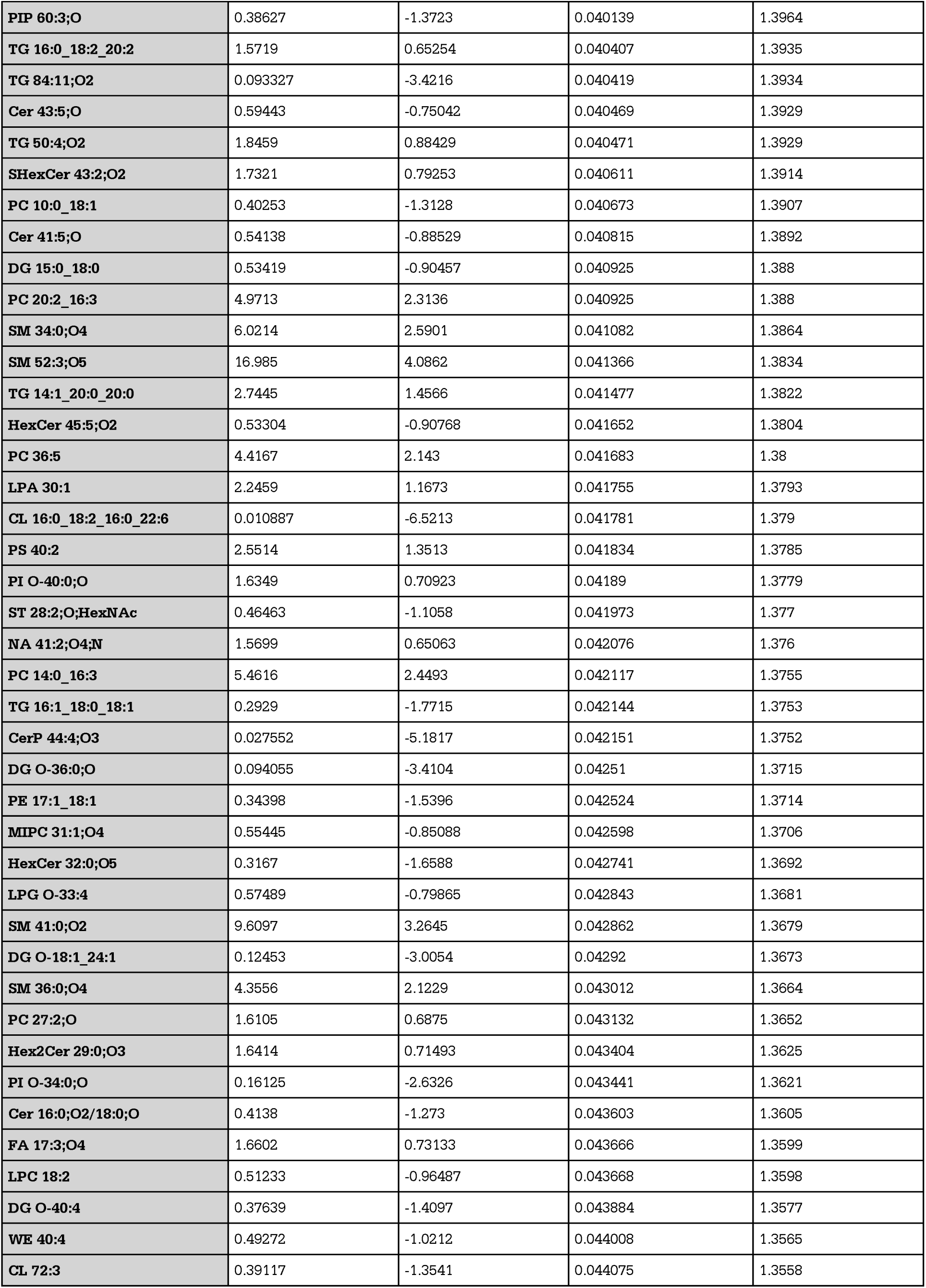
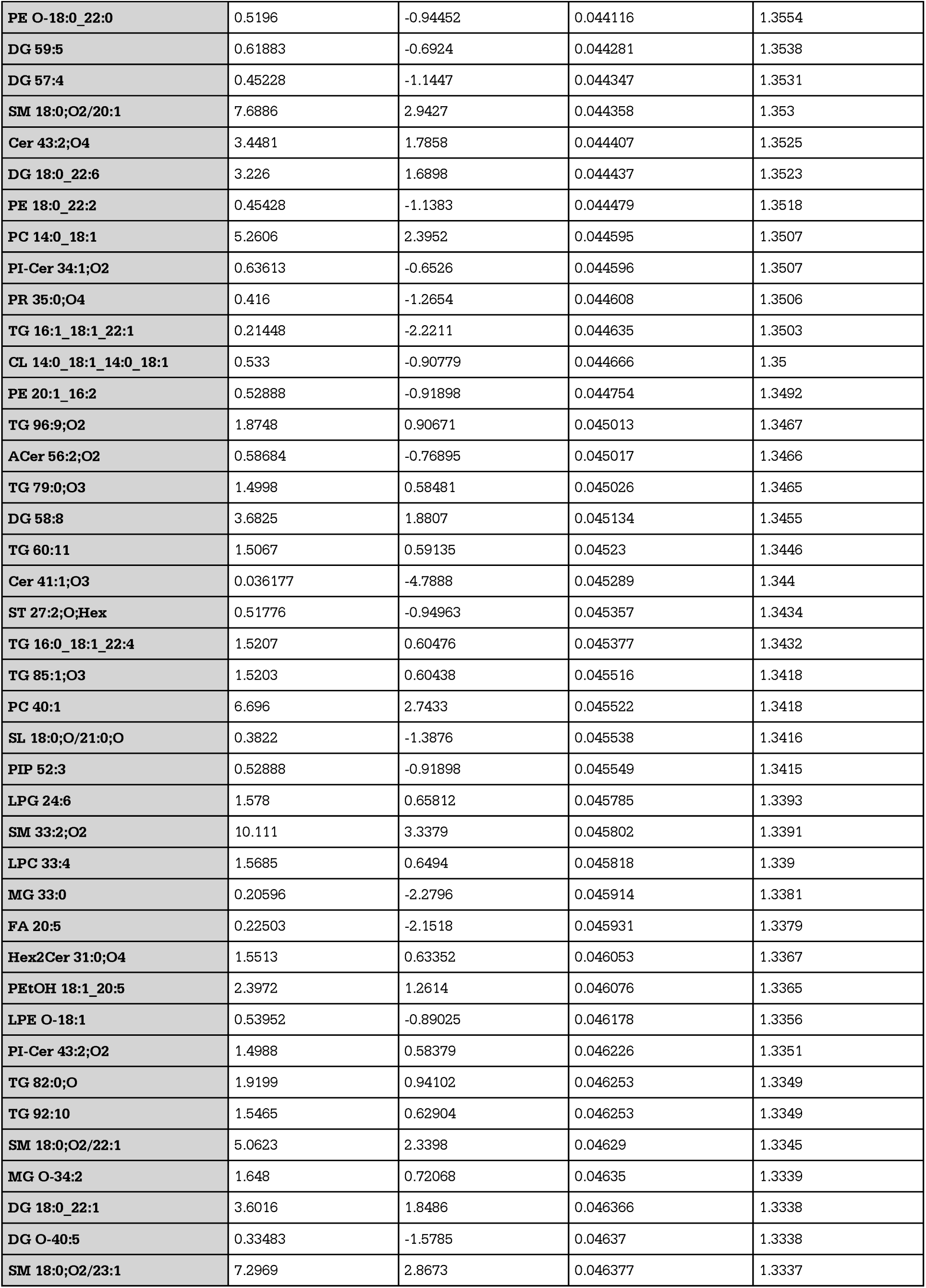
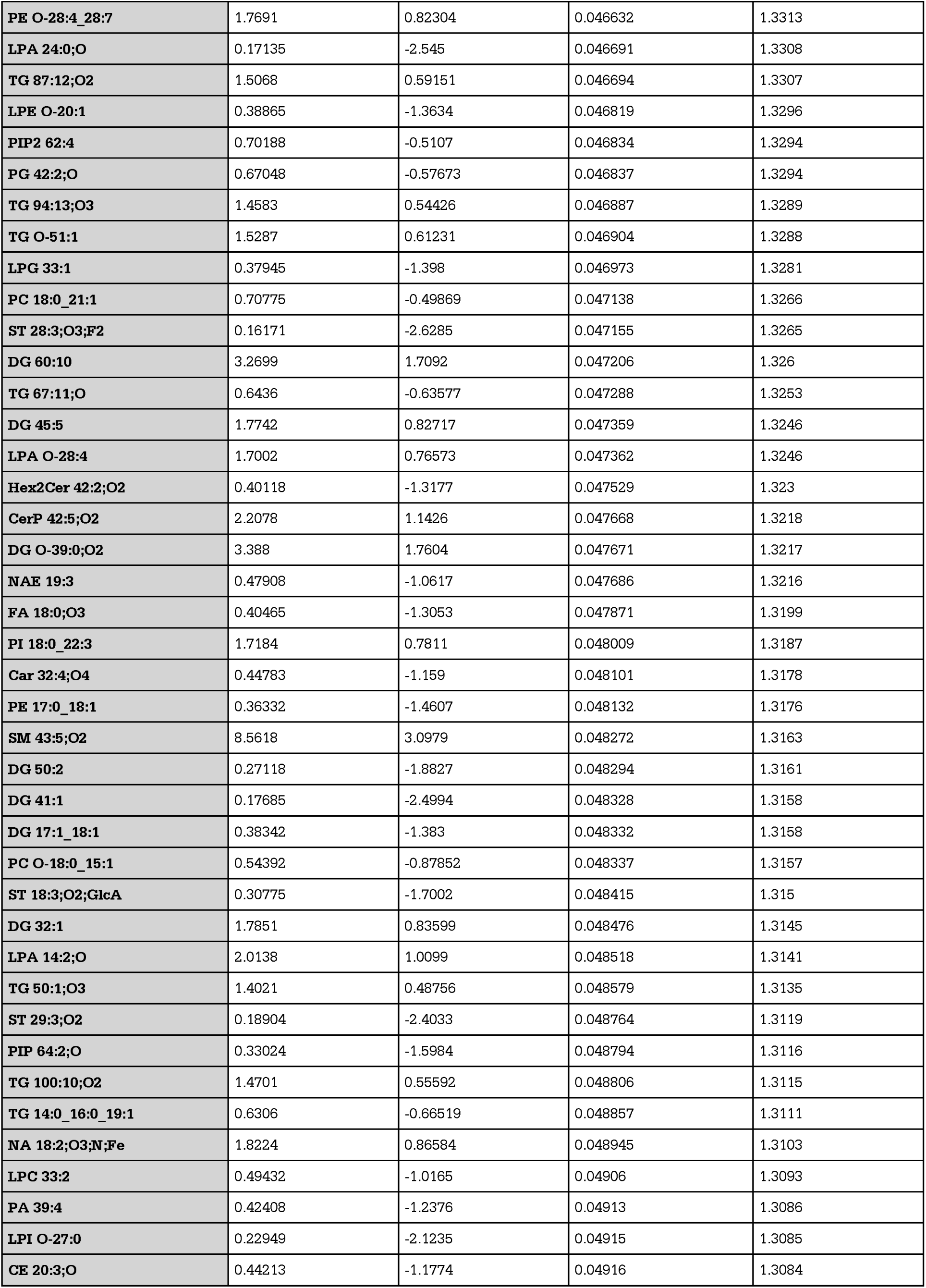
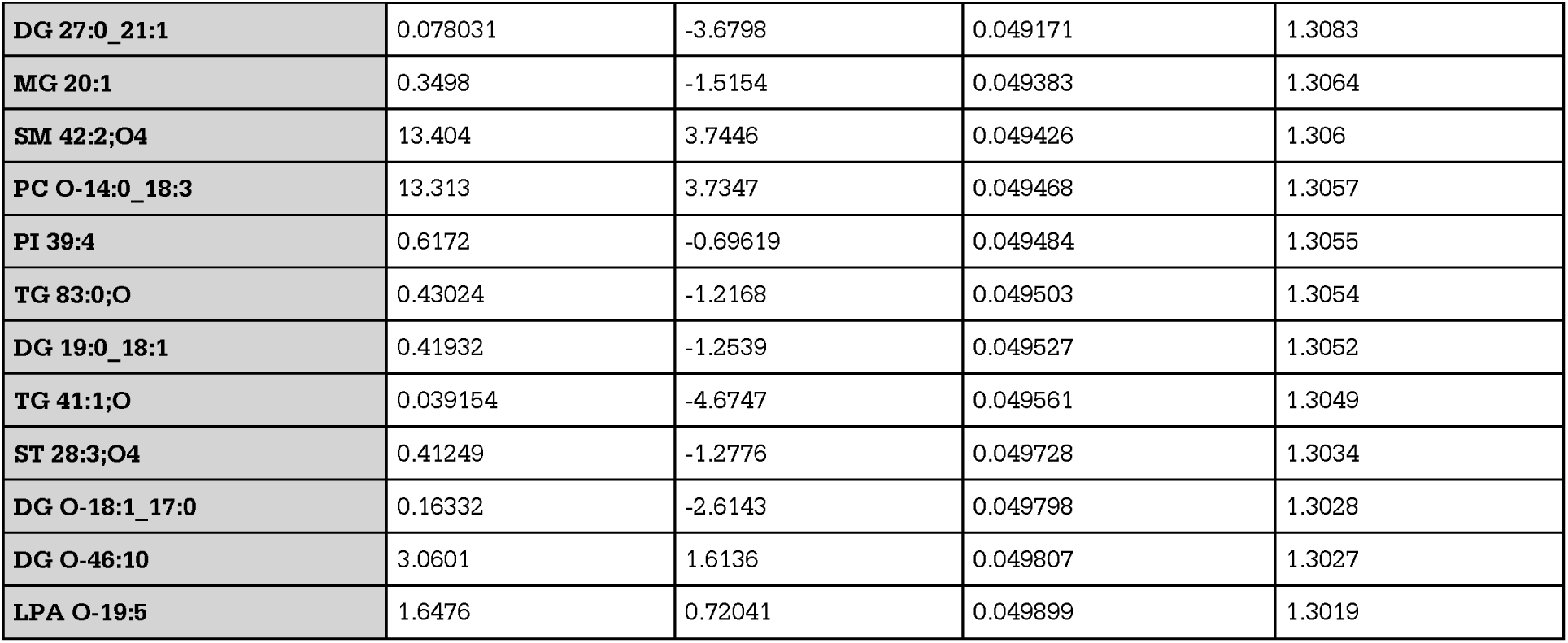
NT cells vs NT EVs volcano plot extended statistics. Extended statistics corresponding to Figure 3C in main text. Extended statistics include fold change (FC), log 2 fold change (log2(FC)), the raw p value (raw.pval) and the -log10 of the p-value (- log10(p)).

**Supplemental Table 5.** LPS cells vs LPS EVs volcano plot extended statistics. Extended statistics corresponding to Figure 3D in main text. Extended statistics include fold change (FC), log 2 fold change (log2(FC)), the raw p value (raw.pval) and the -log10 of the p-value (- log10(p)).

| Lipid | FC | log<sub>2</sub>(FC) | raw.pval | -log<sub>10</sub>(p) |
| --- | --- | --- | --- | --- |
| TG 48:1 | 0.066755 | -3.905 | 8.2173E-11 | 10.085 |
| PI-Cer 41:4;O4 | 0.25439 | -1.9749 | 1.5687E-09 | 8.8045 |
| PG 45:5;O | 0.11922 | -3.0682 | 2.9967E-09 | 8.5234 |
| PI 18:0_20:3 | 0.16457 | -2.6032 | 3.2657E-09 | 8.486 |
| HexCer 38:0;O6 | 0.16199 | -2.626 | 3.6587E-09 | 8.4367 |
| DG O-14:1_18:0 | 0.095247 | -3.3922 | 4.5692E-09 | 8.3402 |
| TG 75:0;O3 | 0.11978 | -3.0615 | 7.3047E-09 | 8.1364 |
| PG 38:1 | 0.085865 | -3.5418 | 1.4084E-08 | 7.8513 |
| PI-Cer 34:0;O6 | 0.1003 | -3.3177 | 1.7724E-08 | 7.7514 |
| Hex2Cer 36:5;O2 | 0.087015 | -3.5226 | 2.3324E-08 | 7.6322 |
| PG 37:0;O | 0.17074 | -2.5501 | 2.7375E-08 | 7.5626 |
| Hex2Cer 34:3;O3 | 0.11379 | -3.1355 | 2.9306E-08 | 7.533 |
| PI 36:1 | 0.21684 | -2.2053 | 4.4695E-08 | 7.3497 |
| PG 43:6;O | 0.069332 | -3.8503 | 4.7982E-08 | 7.3189 |
| PE 40:0 | 0.32609 | -1.6166 | 5.4012E-08 | 7.2675 |
| Hex2Cer 32:0;O3 | 0.12715 | -2.9754 | 5.9592E-08 | 7.2248 |
| HexCer 18:0;O2/23:1 | 4.1034 | 2.0368 | 7.6799E-08 | 7.1146 |
| LPC 33:0;O | 0.51249 | -0.96441 | 7.7315E-08 | 7.1117 |
| PI 20:1_16:2 | 0.3145 | -1.6689 | 8.8667E-08 | 7.0522 |
| PI-Cer 51:3;O4 | 0.6393 | -0.64543 | 8.8799E-08 | 7.0516 |
| HexCer 43:6;O2 | 0.20534 | -2.2839 | 9.0059E-08 | 7.0455 |
| **S**HexCer 40:1;O2 | 0.27641 | -1.8551 | 1.1535E-07 | 6.938 |
| **CL** 16:0_18:2_16:0_22:5 | 0.017185 | -5.8627 | 1.1564E-07 | 6.9369 |
| **Hex2**Cer 32:2;O4 | 0.15247 | -2.7134 | 1.2101E-07 | 6.9172 |
| **Cer** 37:2;O6 | 0.49223 | -1.0226 | 1.3333E-07 | 6.8751 |
| **Car** 34:7 | 2.4415 | 1.2877 | 1.3445E-07 | 6.8714 |
| **SL** 25:0;O/26:0 | 0.019286 | -5.6963 | 1.4272E-07 | 6.8455 |
| **PI** O-40:7 | 0.11472 | -3.1238 | 1.4358E-07 | 6.8429 |
| **Cer** 44:1;O4 | 0.2907 | -1.7824 | 1.5543E-07 | 6.8085 |
| **Hex**Cer 43:5;O2 | 0.27462 | -1.8645 | 1.9362E-07 | 6.7131 |
| **Hex**Cer 41:6;O2 | 0.21758 | -2.2004 | 2.188E-07 | 6.6599 |
| **S**HexCer 37:0;O4 | 0.10309 | -3.278 | 2.2194E-07 | 6.6538 |
| **Glc**ADG 14:0_18:0 | 0.060406 | -4.0492 | 2.244E-07 | 6.649 |
| **PI-Cer** 40:0;O4 | 0.23627 | -2.0815 | 2.3451E-07 | 6.6298 |
| **CL** 16:0_18:2_16:0_22:4 | 0.023627 | -5.4034 | 2.3774E-07 | 6.6239 |
| **TG** 16:2_16:3_18:0 | 7.0607 | 2.8198 | 2.451E-07 | 6.6107 |
| **TG** 51:2;O | 0.024661 | -5.3416 | 2.5419E-07 | 6.5948 |
| **Cer** 42:1;O4 | 0.18448 | -2.4385 | 2.5728E-07 | 6.5896 |
| **PI** O-38:4 | 0.08751 | -3.5144 | 2.7139E-07 | 6.5664 |
| **PI** 18:1_20:1 | 0.18783 | -2.4125 | 2.7337E-07 | 6.5632 |
| **TG** O-14:0_18:0_18:1 | 0.027998 | -5.1585 | 2.9852E-07 | 6.525 |
| **DG** 40:6 | 0.14364 | -2.7995 | 3.1359E-07 | 6.5036 |
| **CL** 16:0_18:2_16:0_20:3 | 0.0034004 | -8.2001 | 3.4782E-07 | 6.4586 |
| **DG** 36:4 | 0.17931 | -2.4794 | 3.5266E-07 | 6.4526 |
| **PI** 18:1_19:1 | 0.12968 | -2.947 | 3.9008E-07 | 6.4088 |
| **PI-Cer** 41:3;O4 | 0.34708 | -1.5267 | 4.0898E-07 | 6.3883 |
| **Hex**Cer 45:6;O3 | 0.16553 | -2.5948 | 4.2068E-07 | 6.376 |
| **CerP** 40:2;O2 | 0.17962 | -2.477 | 4.2709E-07 | 6.3695 |
| **PI** 38:6 | 0.18729 | -2.4166 | 4.4674E-07 | 6.3499 |
| **Hex**Cer 39:1;O4 | 0.19767 | -2.3388 | 4.5968E-07 | 6.3375 |
| **LPC** 32:2 | 0.19967 | -2.3243 | 4.6954E-07 | 6.3283 |
| **PI** 18:0_22:6 | 0.12247 | -3.0295 | 4.7028E-07 | 6.3276 |
| **CL** 16:0_18:2_18:0_18:2 | 0.0029123 | -8.4236 | 4.7548E-07 | 6.3229 |
| **TG** 73:1;O2 | 0.15186 | -2.7192 | 5.2105E-07 | 6.2831 |
| **PI** 20:0_16:2 | 0.2003 | -2.3197 | 5.218E-07 | 6.2825 |
| **CerP** 51:3;O3 | 0.17628 | -2.504 | 6.4515E-07 | 6.1903 |
| **CL** 16:0_20:3_18:0_18:2 | 0.0028487 | -8.4555 | 6.8399E-07 | 6.1649 |
| **Cer** 41:4;O | 0.28184 | -1.827 | 7.5228E-07 | 6.1236 |
| **TG** 12:1_12:3_18:5 | 0.16464 | -2.6026 | 7.5588E-07 | 6.1215 |
| **PI** O-40:6 | 0.13844 | -2.8527 | 7.9509E-07 | 6.0996 |
| **PC** 12:0_16:1 | 0.054758 | -4.1908 | 8.0222E-07 | 6.0957 |
| **PE** 24:0_20:4 | 0.61516 | -0.70097 | 8.3221E-07 | 6.0798 |
| HexCer 42:2;O5 | 0.33754 | -1.5669 | 8.4711E-07 | 6.0721 |
| Hex2Cer 18:0;O2/14:1 | 0.091396 | -3.4517 | 8.5101E-07 | 6.0701 |
| CerP 48:5;O3 | 0.051715 | -4.2733 | 8.5478E-07 | 6.0681 |
| PE 12:0_16:0 | 0.33739 | -1.5675 | 8.6885E-07 | 6.0611 |
| PR 52:2;O4 | 0.20453 | -2.2896 | 8.821E-07 | 6.0545 |
| Cer 48:6;O2 | 0.13091 | -2.9333 | 9.2688E-07 | 6.033 |
| HexCer 35:3;O2 | 0.41225 | -1.2784 | 9.7528E-07 | 6.0109 |
| TG 52:2;O | 0.025611 | -5.2871 | 9.757E-07 | 6.0107 |
| HexCer 43:2;O5 | 0.49077 | -1.0269 | 1.0021E-06 | 5.9991 |
| DG 40:0 | 0.16979 | -2.5582 | 1.0026E-06 | 5.9989 |
| HexCer 43:6;O3 | 0.081058 | -3.6249 | 1.055E-06 | 5.9768 |
| MG 28:1;O | 3.8968 | 1.9623 | 1.0591E-06 | 5.9751 |
| PI 38:3 | 0.37775 | -1.4045 | 1.0708E-06 | 5.9703 |
| HexCer 40:6;O2 | 0.11399 | -3.133 | 1.1412E-06 | 5.9426 |
| PE 14:0_16:0 | 0.17289 | -2.5321 | 1.2008E-06 | 5.9205 |
| PI 35:1 | 0.19079 | -2.3899 | 1.2696E-06 | 5.8963 |
| TG 83:4;O | 2.1609 | 1.1116 | 1.3298E-06 | 5.8762 |
| CL 74:7 | 0.45183 | -1.1461 | 1.3961E-06 | 5.8551 |
| PI 66:2 | 0.30317 | -1.7218 | 1.4745E-06 | 5.8313 |
| PI 16:1_18:1 | 0.17666 | -2.501 | 1.5555E-06 | 5.8081 |
| PE-Cer 44:5;O2 | 1.8762 | 0.9078 | 1.5601E-06 | 5.8069 |
| MG 32:0 | 0.2923 | -1.7745 | 1.6093E-06 | 5.7934 |
| PI 34:1 | 0.35124 | -1.5095 | 1.7925E-06 | 5.7465 |
| CL 16:0_20:3_18:1_18:1 | 0.01876 | -5.7362 | 1.8212E-06 | 5.7396 |
| PG O-39:0;O | 0.043856 | -4.5111 | 1.8531E-06 | 5.7321 |
| PI 16:0_20:4 | 0.11134 | -3.167 | 1.8747E-06 | 5.7271 |
| Hex2Cer 33:2;O4 | 0.13972 | -2.8394 | 1.9084E-06 | 5.7193 |
| TG 59:1 | 0.073165 | -3.7727 | 1.9969E-06 | 5.6997 |
| PG 34:1 | 0.23489 | -2.09 | 2.0682E-06 | 5.6844 |
| PG 50:2;O | 0.12524 | -2.9972 | 2.1764E-06 | 5.6623 |
| Hex2Cer 32:1;O4 | 0.14636 | -2.7724 | 2.1795E-06 | 5.6616 |
| CerP 49:4;O2 | 0.1321 | -2.9203 | 2.2301E-06 | 5.6517 |
| SL 18:0;O/20:1 | 0.12605 | -2.9879 | 2.283E-06 | 5.6415 |
| PI 36:3 | 0.5218 | -0.93843 | 2.4592E-06 | 5.6092 |
| HexCer 43:1;O5 | 0.3773 | -1.4062 | 2.5394E-06 | 5.5953 |
| PG O-40:0;O | 0.028008 | -5.158 | 2.6026E-06 | 5.5846 |
| PI 35:2 | 0.12268 | -3.027 | 2.6232E-06 | 5.5812 |
| DG O-18:1_20:5 | 0.072777 | -3.7804 | 2.7226E-06 | 5.565 |
| DG 40:1 | 0.23222 | -2.1064 | 2.7476E-06 | 5.5611 |
| CerP 45:6;O2 | 0.38258 | -1.3862 | 2.7915E-06 | 5.5542 |
| SHexCer 43:5;O4 | 0.12338 | -3.0188 | 2.8149E-06 | 5.5505 |
| PI 18:0_20:4 | 0.19475 | -2.3603 | 2.8691E-06 | 5.5423 |
| HexCer 42:5;O4 | 0.25343 | -1.9803 | 2.8703E-06 | 5.5421 |
| PI 18:0_22:5 | 0.2159 | -2.2116 | 2.9046E-06 | 5.5369 |
| HexCer 45:3;O5 | 0.17299 | -2.5313 | 3.0503E-06 | 5.5157 |
| CerP 53:3;O3 | 0.020863 | -5.5829 | 3.0544E-06 | 5.5151 |
| CL 16:2_18:1_18:1_18:1 | 0.016352 | -5.9344 | 3.6535E-06 | 5.4373 |
| PI 20:0_16:1 | 0.18491 | -2.4351 | 3.6836E-06 | 5.4337 |
| PC 10:0_16:0 | 0.067541 | -3.8881 | 3.7436E-06 | 5.4267 |
| HexCer 40:1;O5 | 0.3172 | -1.6565 | 3.749E-06 | 5.4261 |
| PI O-18:0_22:5 | 0.1072 | -3.2216 | 3.8585E-06 | 5.4136 |
| PE 32:2;O | 0.25167 | -1.9904 | 3.8597E-06 | 5.4134 |
| LPC 32:4 | 0.43288 | -1.2079 | 3.8939E-06 | 5.4096 |
| FA 31:4;O | 2.2986 | 1.2007 | 4.039E-06 | 5.3937 |
| Cer 20:1;O2/24:1;O | 0.12715 | -2.9754 | 4.3082E-06 | 5.3657 |
| HBMP 16:1_18:1_18:1 | 0.27678 | -1.8532 | 4.3343E-06 | 5.3631 |
| DG 18:0_28:0 | 0.037376 | -4.7417 | 4.513E-06 | 5.3455 |
| Cer 51:4;O | 0.11714 | -3.0937 | 4.5366E-06 | 5.3433 |
| PI O-40:5 | 0.13467 | -2.8925 | 4.6727E-06 | 5.3304 |
| Cer 18:0;O3/21:1;(2OH) | 0.24154 | -2.0497 | 4.8772E-06 | 5.3118 |
| TG 60:1 | 0.074661 | -3.7435 | 4.8783E-06 | 5.3117 |
| MG 30:0 | 0.1308 | -2.9346 | 4.9254E-06 | 5.3076 |
| PE 22:0_24:2 | 0.33573 | -1.5746 | 5.1058E-06 | 5.2919 |
| MG 34:0 | 0.064486 | -3.9549 | 5.3249E-06 | 5.2737 |
| PE 40:1;O | 0.65571 | -0.60886 | 5.5741E-06 | 5.2538 |
| PA 44:5 | 0.30338 | -1.7208 | 5.6698E-06 | 5.2464 |
| FAHFA 35:1;O | 0.23473 | -2.0909 | 6.2867E-06 | 5.2016 |
| SHexCer 42:0;O2 | 0.13279 | -2.9128 | 6.3383E-06 | 5.198 |
| SL 25:0;O/26:2;O | 0.065876 | -3.9241 | 6.5187E-06 | 5.1858 |
| Cer 52:4;O | 0.12629 | -2.9852 | 6.5383E-06 | 5.1845 |
| PC O-44:3 | 0.15581 | -2.6821 | 6.5456E-06 | 5.184 |
| Cer 17:0;3O/18:0;(2OH) | 0.095156 | -3.3936 | 6.6082E-06 | 5.1799 |
| HexCer 47:1;O5 | 0.41586 | -1.2658 | 7.1832E-06 | 5.1437 |
| CerP 29:3;O5 | 4.1993 | 2.0701 | 7.3093E-06 | 5.1361 |
| PS 37:1 | 0.35958 | -1.4756 | 7.3172E-06 | 5.1357 |
| LPG 34:1 | 0.1908 | -2.3899 | 7.3642E-06 | 5.1329 |
| MG O-18:1 | 0.06384 | -3.9694 | 7.4334E-06 | 5.1288 |
| DG O-43:3 | 0.15954 | -2.648 | 7.7875E-06 | 5.1086 |
| DG O-18:1_17:0 | 0.1289 | -2.9556 | 8.1503E-06 | 5.0888 |
| PE-Cer 48:0;O3 | 1.8802 | 0.91091 | 8.3527E-06 | 5.0782 |
| PS-NAc 52:1 | 0.11423 | -3.13 | 8.4344E-06 | 5.0739 |
| PI O-18:1_16:0 | 0.088449 | -3.499 | 8.5216E-06 | 5.0695 |
| PE O-42:3 | 0.15798 | -2.6622 | 8.5625E-06 | 5.0674 |
| DG O-18:0_22:1 | 0.095827 | -3.3834 | 8.6977E-06 | 5.0606 |
| CL 14:0_18:0_18:0_22:4 | 0.025484 | -5.2943 | 8.7038E-06 | 5.0603 |
| PI O-63:3 | 0.21117 | -2.2436 | 8.7809E-06 | 5.0565 |
| TG O-73:0;O3 | 0.013804 | -6.1787 | 8.8389E-06 | 5.0536 |
| PIP 34:5;O | 2.7109 | 1.4388 | 9.0509E-06 | 5.0433 |
| PI 18:0_20:1 | 0.19069 | -2.3907 | 9.1625E-06 | 5.038 |
| PG 42:6;O | 0.34769 | -1.5241 | 9.3776E-06 | 5.0279 |
| Cer 42:0;O3 | 0.33134 | -1.5936 | 9.3972E-06 | 5.027 |
| LPC 21:4 | 0.46091 | -1.1174 | 9.4287E-06 | 5.0256 |
| SHexCer 52:2;O2 | 0.2031 | -2.2997 | 9.5024E-06 | 5.0222 |
| TG 12:0_14:0_18:0 | 0.12504 | -2.9996 | 9.5049E-06 | 5.0221 |
| Cer 37:4;O3 | 1.7839 | 0.83505 | 9.6399E-06 | 5.0159 |
| TG 12:2_12:2_22:6 | 0.024197 | -5.369 | 9.7696E-06 | 5.0101 |
| HexCer 37:5;O2 | 0.31484 | -1.6673 | 9.8579E-06 | 5.0062 |
| PC 16:0_18:1 | 0.29763 | -1.7484 | 9.9398E-06 | 5.0026 |
| PG 39:0 | 0.11994 | -3.0596 | 9.9926E-06 | 5.0003 |
| TG 100:19 | 0.30056 | -1.7343 | 1.0015E-05 | 4.9994 |
| Cer 21:0;O3/24:1;(2OH) | 0.063549 | -3.976 | 1.0235E-05 | 4.9899 |
| DG 58:4 | 0.041053 | -4.6064 | 1.0281E-05 | 4.988 |
| Cer 18:3;O2/24:6 | 0.31775 | -1.654 | 1.0519E-05 | 4.978 |
| DG 18:0_30:0 | 0.058559 | -4.094 | 1.075E-05 | 4.9686 |
| PE 16:0_20:4 | 0.22006 | -2.1841 | 1.0752E-05 | 4.9685 |
| Cer 21:3;O2/24:2;O | 0.44312 | -1.1742 | 1.084E-05 | 4.965 |
| DG O-39:0 | 0.093018 | -3.4263 | 1.0864E-05 | 4.964 |
| HexCer 37:3;O2 | 0.26689 | -1.9057 | 1.1289E-05 | 4.9473 |
| Cer 36:4;O4 | 5.5835 | 2.4812 | 1.1324E-05 | 4.946 |
| PE 20:0_16:1 | 0.23705 | -2.0767 | 1.1535E-05 | 4.938 |
| TG 46:2;O | 0.15419 | -2.6972 | 1.1552E-05 | 4.9374 |
| Cer 16:0;O2/18:0;O | 0.14035 | -2.8329 | 1.1639E-05 | 4.9341 |
| PE 28:0 | 0.30177 | -1.7285 | 1.1666E-05 | 4.9331 |
| CL 16:1_18:1_18:1_18:1 | 0.014727 | -6.0854 | 1.1878E-05 | 4.9253 |
| LPC O-33:4 | 0.25662 | -1.9623 | 1.2011E-05 | 4.9204 |
| MG 34:1 | 0.15034 | -2.7337 | 1.2145E-05 | 4.9156 |
| MG 32:1 | 0.43397 | -1.2043 | 1.2488E-05 | 4.9035 |
| PS 21:0_22:6 | 0.1463 | -2.773 | 1.2551E-05 | 4.9013 |
| PI 35:3 | 2.0834 | 1.0589 | 1.2673E-05 | 4.8971 |
| DG 12:0_16:0 | 0.1409 | -2.8273 | 1.2826E-05 | 4.8919 |
| DG O-40:0 | 0.23845 | -2.0682 | 1.2916E-05 | 4.8889 |
| PI O-18:1_22:5 | 0.26017 | -1.9425 | 1.305E-05 | 4.8844 |
| TG 58:1 | 0.04259 | -4.5533 | 1.3078E-05 | 4.8835 |
| PG 18:0_20:3;O3 | 0.17361 | -2.5261 | 1.3451E-05 | 4.8713 |
| NAT 32:0;O | 0.34258 | -1.5455 | 1.3579E-05 | 4.8671 |
| PC O-16:0_20:4 | 0.18238 | -2.455 | 1.4184E-05 | 4.8482 |
| SHexCer 41:3;O3 | 0.27211 | -1.8778 | 1.4532E-05 | 4.8377 |
| Cer 18:1;O2/24:0;O | 0.18666 | -2.4215 | 1.4542E-05 | 4.8374 |
| PE 18:1_22:5 | 0.29679 | -1.7525 | 1.4849E-05 | 4.8283 |
| TG 12:2_12:2_20:5 | 0.22016 | -2.1834 | 1.5036E-05 | 4.8229 |
| HexCer 43:4;O5 | 0.16793 | -2.574 | 1.5489E-05 | 4.81 |
| MG 16:0 | 0.14321 | -2.8038 | 1.5548E-05 | 4.8083 |
| DG 26:0_18:1 | 0.15969 | -2.6467 | 1.5565E-05 | 4.8079 |
| PE 18:0_20:3 | 0.31443 | -1.6692 | 1.6016E-05 | 4.7955 |
| PI 37:2 | 0.39599 | -1.3365 | 1.6089E-05 | 4.7935 |
| Cer 18:0;O3/19:1;(2OH) | 0.1007 | -3.3118 | 1.7432E-05 | 4.7586 |
| LPC 31:3 | 1.9061 | 0.93066 | 1.762E-05 | 4.754 |
| PR 25:3;O | 0.30106 | -1.7319 | 1.7647E-05 | 4.7533 |
| LPE 19:3;O | 2.3922 | 1.2583 | 1.7673E-05 | 4.7527 |
| TG 14:0_15:0_17:0 | 0.16176 | -2.6281 | 1.787E-05 | 4.7479 |
| Car 34:0 | 1.9097 | 0.93332 | 1.7897E-05 | 4.7472 |
| SHexCer 40:0;O2 | 0.47862 | -1.063 | 1.8457E-05 | 4.7338 |
| LPI O-34:0 | 0.26348 | -1.9242 | 1.9003E-05 | 4.7212 |
| HexCer 43:4;O4 | 0.37857 | -1.4014 | 1.9043E-05 | 4.7203 |
| HexCer 41:3;O5 | 0.16081 | -2.6366 | 1.9322E-05 | 4.7139 |
| MG 18:1 | 0.18101 | -2.4659 | 1.9341E-05 | 4.7135 |
| FAHFA 19:2/19:2 | 0.2071 | -2.2716 | 1.9431E-05 | 4.7115 |
| HBMP 14:0_18:1_22:2 | 0.22178 | -2.1728 | 1.9475E-05 | 4.7105 |
| PE O-18:5_23:0 | 0.11078 | -3.1743 | 1.9569E-05 | 4.7084 |
| MG O-32:5 | 0.12979 | -2.9457 | 1.9744E-05 | 4.7046 |
| Cer 37:0;O3 | 0.29784 | -1.7474 | 1.9889E-05 | 4.7014 |
| PA 44:6 | 0.12082 | -3.049 | 1.9952E-05 | 4.7 |
| CL 72:4 | 0.0031083 | -8.3297 | 1.9989E-05 | 4.6992 |
| Cer 41:5;O | 0.39546 | -1.3384 | 2.004E-05 | 4.6981 |
| PI O-67:3 | 0.026925 | -5.2149 | 2.006E-05 | 4.6977 |
| PE 44:6 | 0.23257 | -2.1043 | 2.0158E-05 | 4.6956 |
| HexCer 45:2;O5 | 0.37446 | -1.4171 | 2.0329E-05 | 4.6919 |
| SHexCer 43:4;O4 | 0.125 | -3 | 2.0419E-05 | 4.69 |
| PI 39:5 | 0.19043 | -2.3927 | 2.125E-05 | 4.6726 |
| Cer 44:1;O3 | 0.20361 | -2.2961 | 2.2071E-05 | 4.6562 |
| ST 30:3;O | 0.15776 | -2.6642 | 2.2257E-05 | 4.6525 |
| LPC 32:0;O | 0.61089 | -0.71102 | 2.2302E-05 | 4.6517 |
| Cer 16:0;O2/18:1;O | 0.14244 | -2.8116 | 2.2408E-05 | 4.6496 |
| PG 56:7;O | 0.092175 | -3.4395 | 2.2597E-05 | 4.6459 |
| **TG 12:2_12:3_13:1** | 2.4445 | 1.2896 | 2.2764E-05 | 4.6427 |
| **TG 98:8** | 4.0904 | 2.0322 | 2.2927E-05 | 4.6397 |
| **PC O-18:1_20:4** | 0.15451 | -2.6942 | 2.3013E-05 | 4.638 |
| **PS 41:2** | 0.51319 | -0.96245 | 2.327E-05 | 4.6332 |
| **SHexCer 38:1;O4** | 0.21919 | -2.1898 | 2.3916E-05 | 4.6213 |
| **HexCer 15:3;O2/30:8** | 0.46977 | -1.09 | 2.3934E-05 | 4.621 |
| **CL 14:0_18:1_14:0_22:3** | 0.0032358 | -8.2717 | 2.3982E-05 | 4.6201 |
| **LPA O-27:1** | 1.6607 | 0.73175 | 2.4163E-05 | 4.6168 |
| **Cer 46:4;O3** | 0.21463 | -2.2201 | 2.4371E-05 | 4.6131 |
| **DG 42:1** | 0.37292 | -1.4231 | 2.4423E-05 | 4.6122 |
| **CE 18:2;O** | 0.18832 | -2.4087 | 2.4714E-05 | 4.607 |
| **SHexCer 33:0;O5** | 2.841 | 1.5064 | 2.4737E-05 | 4.6067 |
| **PI 36:2** | 0.59229 | -0.75563 | 2.4942E-05 | 4.6031 |
| **HexCer 45:5;O4** | 0.2353 | -2.0874 | 2.5221E-05 | 4.5982 |
| **SL 19:0;O/17:0** | 0.10814 | -3.209 | 2.5285E-05 | 4.5971 |
| **PIP3 47:9** | 2.7867 | 1.4786 | 2.5442E-05 | 4.5945 |
| **PE 42:1;O** | 0.2649 | -1.9165 | 2.5534E-05 | 4.5929 |
| **TG 14:0_18:0_22:1** | 0.033362 | -4.9056 | 2.5873E-05 | 4.5872 |
| **MG 20:4** | 0.074142 | -3.7536 | 2.6333E-05 | 4.5795 |
| **TG 57:1** | 0.039817 | -4.6505 | 2.6846E-05 | 4.5711 |
| **TG 66:11** | 0.0721 | -3.7939 | 2.7136E-05 | 4.5664 |
| **ST 27:3;O2;F2** | 0.25105 | -1.9939 | 2.7528E-05 | 4.5602 |
| **PE 16:0_18:1** | 0.25442 | -1.9747 | 2.8113E-05 | 4.5511 |
| **HBMP 14:0_18:1_22:1** | 0.11837 | -3.0786 | 2.8456E-05 | 4.5458 |
| **PG 52:2;O** | 0.16339 | -2.6136 | 2.8494E-05 | 4.5452 |
| **PC O-18:1_16:0** | 0.35508 | -1.4938 | 2.8962E-05 | 4.5382 |
| **DG O-37:3** | 0.2285 | -2.1297 | 2.9229E-05 | 4.5342 |
| **DG 15:0_18:0** | 0.11691 | -3.0966 | 2.9914E-05 | 4.5241 |
| **PI 38:4** | 0.40207 | -1.3145 | 3.05E-05 | 4.5157 |
| **PI 38:5** | 0.32436 | -1.6243 | 3.1022E-05 | 4.5083 |
| **DG 40:4;O** | 1.876 | 0.90764 | 3.1046E-05 | 4.508 |
| **DG 35:1;O2** | 0.12011 | -3.0575 | 3.1447E-05 | 4.5024 |
| **DG 44:10** | 0.19817 | -2.3352 | 3.1521E-05 | 4.5014 |
| **DG O-36:3** | 0.13615 | -2.8767 | 3.1807E-05 | 4.4975 |
| **PI 18:1_22:5** | 0.15192 | -2.7186 | 3.1959E-05 | 4.4954 |
| **Cer 18:1;O2/16:0;O** | 0.13176 | -2.924 | 3.3016E-05 | 4.4813 |
| **PE 34:2** | 0.17164 | -2.5426 | 3.3385E-05 | 4.4765 |
| **SHexCer 54:2;O2** | 0.096161 | -3.3784 | 3.3717E-05 | 4.4721 |
| **DG 38:5** | 0.48935 | -1.0311 | 3.6162E-05 | 4.4418 |
| **PE 14:0_18:0** | 0.21501 | -2.2175 | 3.7658E-05 | 4.4241 |
| **HexCer 33:2;O2** | 0.35218 | -1.5056 | 3.8852E-05 | 4.4106 |
| PI 18:1_20:3 | 0.17192 | -2.5402 | 3.8989E-05 | 4.4091 |
| PE 35:0 | 0.57243 | -0.80484 | 3.9847E-05 | 4.3996 |
| DG O-16:1_18:0 | 0.095753 | -3.3845 | 4.1545E-05 | 4.3815 |
| DG O-35:3 | 0.21038 | -2.2489 | 4.252E-05 | 4.3714 |
| PG 58:9;O | 0.091689 | -3.4471 | 4.2923E-05 | 4.3673 |
| Cer 47:6;O3 | 0.23135 | -2.1119 | 4.3339E-05 | 4.3631 |
| TG O-99:10 | 2.651 | 1.4065 | 4.3537E-05 | 4.3611 |
| PK 37:2;O8 | 0.27733 | -1.8503 | 4.3569E-05 | 4.3608 |
| DG O-18:2_22:5 | 0.11083 | -3.1735 | 4.4386E-05 | 4.3528 |
| Cer 18:1;O2/24:1;O | 0.096113 | -3.3791 | 4.4526E-05 | 4.3514 |
| DG 50:2 | 0.046642 | -4.4222 | 4.6074E-05 | 4.3365 |
| LPC 30:0;O | 0.57186 | -0.80628 | 4.6111E-05 | 4.3362 |
| CerP 47:2;O2 | 0.29118 | -1.78 | 4.6847E-05 | 4.3293 |
| CerP 31:3;O6 | 2.888 | 1.5301 | 4.7286E-05 | 4.3253 |
| Cer 20:0;O2/16:1;O | 0.12344 | -3.0181 | 4.7922E-05 | 4.3195 |
| Cer 17:0;3O/18:1;(2OH) | 0.10982 | -3.1868 | 4.8336E-05 | 4.3157 |
| DG O-18:1_24:0 | 0.24328 | -2.0393 | 5.066E-05 | 4.2953 |
| PC O-14:1_18:1 | 0.32448 | -1.6238 | 5.0709E-05 | 4.2949 |
| NAT 26:2;O3 | 3.2771 | 1.7124 | 5.0825E-05 | 4.2939 |
| PE 28:1 | 0.57838 | -0.78992 | 5.0964E-05 | 4.2927 |
| PI O-36:1 | 0.070142 | -3.8336 | 5.1735E-05 | 4.2862 |
| PC 18:1_26:1 | 0.35213 | -1.5058 | 5.2786E-05 | 4.2775 |
| PE 18:0_26:2 | 0.25907 | -1.9486 | 5.3755E-05 | 4.2696 |
| HexCer 41:6;O3 | 0.48808 | -1.0348 | 5.3874E-05 | 4.2686 |
| Cer 40:5;O3 | 0.35293 | -1.5026 | 5.4057E-05 | 4.2671 |
| HBMP 16:0_18:1_18:1 | 0.05993 | -4.0606 | 5.4118E-05 | 4.2667 |
| PI-Cer 41:2;O2 | 0.081512 | -3.6169 | 5.4129E-05 | 4.2666 |
| PE O-38:0 | 0.26339 | -1.9247 | 5.4587E-05 | 4.2629 |
| PE O-40:6 | 3.1629 | 1.6613 | 5.4833E-05 | 4.261 |
| PC 10:0_18:1 | 0.26397 | -1.9216 | 5.5772E-05 | 4.2536 |
| ACer 58:0;O3 | 0.02761 | -5.1787 | 5.6534E-05 | 4.2477 |
| DG 56:1 | 0.031484 | -4.9892 | 5.7349E-05 | 4.2415 |
| TG 84:11;O3 | 0.20547 | -2.283 | 5.8224E-05 | 4.2349 |
| HexCer 35:3;O3 | 0.52242 | -0.93672 | 5.8484E-05 | 4.233 |
| DG O-16:0_18:0 | 0.096865 | -3.3679 | 5.8791E-05 | 4.2307 |
| Cer 18:1;O2/26:1;O | 0.12569 | -2.992 | 5.9361E-05 | 4.2265 |
| PE 18:0_20:2 | 0.38522 | -1.3762 | 5.9585E-05 | 4.2249 |
| SHEXcer 43:3;O3 | 0.24542 | -2.0267 | 6.1308E-05 | 4.2125 |
| LPC 33:4 | 1.9825 | 0.98732 | 6.151E-05 | 4.2111 |
| TG 83:0;O3 | 0.27875 | -1.8429 | 6.2013E-05 | 4.2075 |
| TG O-84:9 | 0.24152 | -2.0498 | 6.3067E-05 | 4.2002 |
| **Car 32:4;O4** | 0.28373 | -1.8174 | 6.4302E-05 | 4.1918 |
| **FA 20:3** | 0.085388 | -3.5498 | 6.5875E-05 | 4.1813 |
| **HexCer 45:6;O2** | 0.32505 | -1.6213 | 6.7089E-05 | 4.1733 |
| **PG 34:1;O** | 0.27645 | -1.8549 | 6.7763E-05 | 4.169 |
| **SHexCer 52:1;O2** | 0.060563 | -4.0454 | 6.9437E-05 | 4.1584 |
| **PC 16:0_20:4** | 0.16161 | -2.6294 | 7.0297E-05 | 4.1531 |
| **Cer 18:0;3O/19:0;(2OH)** | 0.090636 | -3.4638 | 7.0424E-05 | 4.1523 |
| **DG 30:1** | 0.11869 | -3.0748 | 7.0593E-05 | 4.1512 |
| **Hex2Cer 38:6;O3** | 0.13675 | -2.8703 | 7.0785E-05 | 4.1501 |
| **PI 40:6** | 0.068739 | -3.8627 | 7.1268E-05 | 4.1471 |
| **HexCer 45:4;O4** | 0.18658 | -2.4222 | 7.1656E-05 | 4.1447 |
| **LPC 16:1** | 0.21439 | -2.2217 | 7.2787E-05 | 4.1379 |
| **PE O-18:0_15:0** | 0.28275 | -1.8224 | 7.3196E-05 | 4.1355 |
| **SHexCer 42:1;O2** | 0.063492 | -3.9773 | 7.4764E-05 | 4.1263 |
| **Cer 39:0;O3** | 0.39436 | -1.3424 | 7.5327E-05 | 4.1231 |
| **HexCer 41:2;O4** | 0.22749 | -2.1361 | 7.592E-05 | 4.1196 |
| **PE 18:0_24:1** | 0.34843 | -1.5211 | 7.6652E-05 | 4.1155 |
| **PE O-14:0_22:6** | 1.7091 | 0.77321 | 7.6922E-05 | 4.1139 |
| **HexCer 41:3;O4** | 0.40264 | -1.3124 | 7.7478E-05 | 4.1108 |
| **SM 44:4;O3** | 0.26278 | -1.928 | 7.7736E-05 | 4.1094 |
| **MG 24:2** | 1.9016 | 0.92719 | 7.8942E-05 | 4.1027 |
| **HexCer 32:0;O5** | 0.25284 | -1.9837 | 7.9943E-05 | 4.0972 |
| **PS 17:0_18:2** | 2.1088 | 1.0764 | 8.0219E-05 | 4.0957 |
| **PE-Cer 34:1;O;P** | 0.52274 | -0.93584 | 8.0224E-05 | 4.0957 |
| **TG O-8:0_8:0_16:0** | 7.0185 | 2.8112 | 8.0634E-05 | 4.0935 |
| **PA O-42:5** | 0.25615 | -1.9649 | 8.3961E-05 | 4.0759 |
| **TG 14:1_16:0_18:1** | 2.4224 | 1.2764 | 8.5309E-05 | 4.069 |
| **LPE O-19:2** | 1.6173 | 0.6936 | 8.557E-05 | 4.0677 |
| **LPI 20:4** | 1.6066 | 0.68405 | 8.5919E-05 | 4.0659 |
| **DG 14:0_18:0** | 6.2555 | 2.6451 | 8.6144E-05 | 4.0648 |
| **Hex2Cer 30:0;O3** | 0.45326 | -1.1416 | 8.6653E-05 | 4.0622 |
| **PI 18:1_20:4** | 0.15711 | -2.6701 | 8.885E-05 | 4.0513 |
| **PE 42:2** | 0.32087 | -1.6399 | 8.9194E-05 | 4.0497 |
| **MG 33:1** | 1.4467 | 0.53272 | 8.9891E-05 | 4.0463 |
| **DG 16:0_20:0** | 0.20113 | -2.3138 | 9.101E-05 | 4.0409 |
| **Cer 41:1;O3** | 0.064038 | -3.9649 | 9.1087E-05 | 4.0405 |
| **Cer 18:0;O2/22:0;O** | 0.28701 | -1.8008 | 9.1943E-05 | 4.0365 |
| **DG 19:0_18:1** | 0.24736 | -2.0153 | 9.3098E-05 | 4.0311 |
| **Cer 38:4;O3** | 2.8994 | 1.5358 | 9.3426E-05 | 4.0295 |
| **HexCer 39:3;O2** | 0.43669 | -1.1953 | 9.4203E-05 | 4.0259 |
| **PG O-43:4** | 2.4461 | 1.2905 | 9.5384E-05 | 4.0205 |
| HexCer 34:3;O3 | 0.27716 | -1.8512 | 9.5661E-05 | 4.0193 |
| HexCer 42:4;O4 | 0.44362 | -1.1726 | 9.6081E-05 | 4.0174 |
| DG O-18:1_22:5 | 0.095121 | -3.3941 | 9.8186E-05 | 4.008 |
| Cer 44:4;O3 | 0.17017 | -2.5549 | 9.8868E-05 | 4.0049 |
| Cer 32:1;O2 | 0.24143 | -2.0503 | 9.9577E-05 | 4.0018 |
| DG 15:0_18:1 | 0.084503 | -3.5649 | 9.9653E-05 | 4.0015 |
| LPG 33:5 | 1.5981 | 0.67634 | 0.00010011 | 3.9995 |
| ST 27:3 | 0.26041 | -1.9412 | 0.00010126 | 3.9946 |
| PS 18:1_20:3 | 1.9458 | 0.96039 | 0.00010455 | 3.9807 |
| PE 14:0_16:1 | 0.3181 | -1.6524 | 0.00010459 | 3.9805 |
| PC O-18:1_22:1 | 0.64366 | -0.63562 | 0.00010461 | 3.9804 |
| Cer 18:0;3O/23:1;(2OH) | 0.17709 | -2.4975 | 0.00010523 | 3.9779 |
| PE 42:2;O | 0.43243 | -1.2095 | 0.00010549 | 3.9768 |
| DG 40:7;O | 0.40415 | -1.307 | 0.00010561 | 3.9763 |
| TG 49:0 | 0.18389 | -2.4431 | 0.00010571 | 3.9759 |
| CL 16:0_18:2_16:0_20:4 | 0.0074214 | -7.0741 | 0.00010654 | 3.9725 |
| SLBPA 54:3 | 0.12699 | -2.9772 | 0.00010931 | 3.9614 |
| Hex2Cer 31:2;O4 | 0.40773 | -1.2943 | 0.0001099 | 3.959 |
| CL 16:0_18:2_16:0_18:2 | 0.014649 | -6.093 | 0.00011199 | 3.9508 |
| PI 52:2;O | 0.091186 | -3.455 | 0.00011336 | 3.9455 |
| Cer 16:0;O2/20:1;O | 0.14564 | -2.7795 | 0.00011491 | 3.9396 |
| DG O-35:0;O | 0.13831 | -2.8541 | 0.00011666 | 3.9331 |
| Cer 18:0;O3/42:1;(2OH) | 0.094011 | -3.411 | 0.00011701 | 3.9318 |
| PE O-14:0_17:0 | 0.30124 | -1.731 | 0.00012007 | 3.9206 |
| Cer 44:0;O3 | 0.47333 | -1.0791 | 0.00012009 | 3.9205 |
| TG 62:10 | 0.15097 | -2.7276 | 0.00012021 | 3.9201 |
| PE-Cer 44:0;O3 | 0.47467 | -1.075 | 0.00012031 | 3.9197 |
| PE-Cer 42:6;O4 | 0.19204 | -2.3805 | 0.00012296 | 3.9102 |
| PI-Cer 39:0;O2 | 0.15938 | -2.6495 | 0.00012454 | 3.9047 |
| SPB 20:0;O2 | 3.109 | 1.6365 | 0.0001246 | 3.9045 |
| Cer 18:1;O2/23:0;O | 0.17192 | -2.5402 | 0.00012533 | 3.9019 |
| LPC 22:6 | 0.092592 | -3.433 | 0.00012595 | 3.8998 |
| SM 36:3;O6 | 1.7816 | 0.83318 | 0.00013549 | 3.8681 |
| DG O-18:1_20:4 | 0.079487 | -3.6531 | 0.00013567 | 3.8675 |
| DG 58:1 | 0.036253 | -4.7858 | 0.00013594 | 3.8666 |
| MG O-32:6 | 2.2124 | 1.1456 | 0.0001372 | 3.8627 |
| PI O-34:0;O | 0.47873 | -1.0627 | 0.00013747 | 3.8618 |
| HexCer 40:5;O2 | 0.44413 | -1.1709 | 0.00013758 | 3.8614 |
| Cer 41:5;O2 | 0.33034 | -1.598 | 0.00013961 | 3.8551 |
| CL 14:0_18:1_18:0_18:2 | 0.0031372 | -8.3163 | 0.00014003 | 3.8538 |
| PE-NMe2 24:0 | 0.49346 | -1.019 | 0.00014027 | 3.853 |
| PE 18:0_16:1 | 0.27686 | -1.8528 | 0.00014055 | 3.8522 |
| PIP2 30:6;O | 2.0309 | 1.0221 | 0.00014064 | 3.8519 |
| TG 54:6;O2 | 2.0008 | 1.0006 | 0.00014154 | 3.8491 |
| Cer 42:6;O3 | 0.34771 | -1.524 | 0.00014321 | 3.844 |
| PI 38:1 | 0.49379 | -1.018 | 0.00014349 | 3.8432 |
| PG O-44:3 | 0.23527 | -2.0876 | 0.00014552 | 3.8371 |
| Cer 43:4;O2 | 2.055 | 1.0391 | 0.00014665 | 3.8337 |
| DG 17:0_18:1 | 0.20392 | -2.2939 | 0.00014732 | 3.8317 |
| PI O-20:0_16:1 | 0.21344 | -2.2281 | 0.00014786 | 3.8301 |
| MIPC 29:6;O5 | 2.6398 | 1.4004 | 0.00015161 | 3.8193 |
| TG 50:5;O2 | 0.35313 | -1.5017 | 0.0001535 | 3.8139 |
| HexCer 38:5;O2 | 0.37123 | -1.4296 | 0.00015351 | 3.8139 |
| SHexCer 38:0;O2 | 0.17345 | -2.5274 | 0.0001538 | 3.813 |
| LPI O-29:0 | 0.40132 | -1.3172 | 0.00015417 | 3.812 |
| HexCer 37:6;O2 | 0.27886 | -1.8424 | 0.00015673 | 3.8049 |
| Car 34:5;O | 0.19675 | -2.3456 | 0.00015684 | 3.8045 |
| PG 50:3;O | 0.096067 | -3.3798 | 0.00015811 | 3.801 |
| PC 12:0_16:0 | 0.23022 | -2.1189 | 0.00015827 | 3.8006 |
| PI-Cer 39:3;O6 | 2.1201 | 1.0842 | 0.00015887 | 3.7989 |
| DG O-18:2_20:0 | 0.10056 | -3.3138 | 0.00015892 | 3.7988 |
| TG 14:0_18:0_20:1 | 0.020035 | -5.6413 | 0.00015981 | 3.7964 |
| HexCer 18:0;O2/24:2 | 1.7543 | 0.81092 | 0.00016024 | 3.7952 |
| DG 57:6 | 0.16193 | -2.6266 | 0.0001616 | 3.7916 |
| PE 38:0;O | 1.5472 | 0.6297 | 0.00016284 | 3.7882 |
| SHexCer 43:3;O4 | 0.13045 | -2.9384 | 0.00016421 | 3.7846 |
| Car 34:2;O | 0.1726 | -2.5345 | 0.00016438 | 3.7842 |
| CL 14:0_18:0_14:0_22:5 | 0.052345 | -4.2558 | 0.00016724 | 3.7767 |
| LPC 17:1 | 0.54698 | -0.87044 | 0.00016807 | 3.7745 |
| NAT 20:3 | 1.7405 | 0.79946 | 0.00016927 | 3.7714 |
| DG O-16:2_18:0 | 0.085054 | -3.5555 | 0.0001696 | 3.7706 |
| PI-Cer 36:2;O3 | 2.5773 | 1.3659 | 0.00017042 | 3.7685 |
| TG 54:2 | 0.075942 | -3.7189 | 0.00017054 | 3.7682 |
| MG 33:0 | 0.14519 | -2.784 | 0.00017231 | 3.7637 |
| PE O-42:0 | 0.57268 | -0.8042 | 0.00017236 | 3.7636 |
| PG 14:0_18:0 | 0.10993 | -3.1854 | 0.00017257 | 3.763 |
| LPG 33:1 | 0.29226 | -1.7747 | 0.000173 | 3.762 |
| HexCer 45:6;O6 | 0.21167 | -2.2401 | 0.0001731 | 3.7617 |
| PE-Cer 44:6;O2 | 1.606 | 0.68344 | 0.00017439 | 3.7585 |
| PG O-45:0 | 0.70199 | -0.51049 | 0.00017778 | 3.7501 |
| TG 12:0_12:3_20:6 | 0.25618 | -1.9648 | 0.00017827 | 3.7489 |
| DG 55:4 | 0.098729 | -3.3404 | 0.00018027 | 3.7441 |
| DG 39:5;O | 0.55944 | -0.83794 | 0.00018269 | 3.7383 |
| PE 18:1_24:1 | 0.33207 | -1.5905 | 0.00018304 | 3.7375 |
| CE 28:1;O | 0.3406 | -1.5539 | 0.00018305 | 3.7374 |
| Cer 28:0;O3 | 3.4387 | 1.7819 | 0.0001845 | 3.734 |
| PE 18:0_20:1 | 0.39781 | -1.3298 | 0.00018812 | 3.7256 |
| DG 39:0;O2 | 0.27759 | -1.849 | 0.00018891 | 3.7237 |
| DG 32:0_22:2 | 0.016049 | -5.9614 | 0.00019145 | 3.7179 |
| HexCer 38:6;O2 | 0.40001 | -1.3219 | 0.00019245 | 3.7157 |
| Car 34:2 | 0.10855 | -3.2035 | 0.00019318 | 3.714 |
| TG O-8:0_8:0_18:2 | 0.51563 | -0.9556 | 0.00019535 | 3.7092 |
| Hex2Cer 30:1;O6 | 2.5047 | 1.3246 | 0.00019673 | 3.7061 |
| DG 41:2 | 0.16798 | -2.5736 | 0.00019704 | 3.7055 |
| PI 40:3 | 1.5643 | 0.64548 | 0.00019895 | 3.7013 |
| DG O-20:0_16:2 | 0.088489 | -3.4984 | 0.00019952 | 3.7 |
| DG 32:2 | 0.17588 | -2.5073 | 0.00019953 | 3.7 |
| CL 14:0_18:1_18:1_18:1 | 0.024659 | -5.3417 | 0.00019975 | 3.6995 |
| PI O-40:0;O | 1.9523 | 0.96517 | 0.00020023 | 3.6985 |
| TG 55:1 | 0.22014 | -2.1835 | 0.00020219 | 3.6942 |
| PS 18:0_22:0 | 0.49472 | -1.0153 | 0.00020485 | 3.6886 |
| DG 16:0_20:3 | 0.26964 | -1.8909 | 0.00020496 | 3.6883 |
| PC 20:4 | 22.35 | 4.4822 | 0.00020637 | 3.6854 |
| PG 45:8;O | 0.10318 | -3.2767 | 0.00020673 | 3.6846 |
| WE 40:4 | 0.20741 | -2.2695 | 0.00020689 | 3.6843 |
| Cer 38:0;O5 | 0.34796 | -1.523 | 0.00020894 | 3.68 |
| Car 22:4;O3 | 0.34917 | -1.518 | 0.00020917 | 3.6795 |
| HexCer 40:4;O3 | 0.36205 | -1.4658 | 0.00020945 | 3.6789 |
| PG 20:4;O | 0.51664 | -0.95276 | 0.00021258 | 3.6725 |
| PC O-36:6 | 17.137 | 4.0991 | 0.00021267 | 3.6723 |
| PG 39:4;O | 0.092721 | -3.431 | 0.00021451 | 3.6686 |
| CerP 43:5;O2 | 0.54495 | -0.87581 | 0.00021625 | 3.665 |
| DG 41:0;O2 | 0.3384 | -1.5632 | 0.00021665 | 3.6642 |
| PE 56:5 | 0.35187 | -1.5069 | 0.00021784 | 3.6619 |
| TG 100:17 | 0.15604 | -2.68 | 0.00021923 | 3.6591 |
| ACer 43:6;O3 | 0.2265 | -2.1424 | 0.00021927 | 3.659 |
| MG 31:0 | 0.51999 | -0.94345 | 0.00022027 | 3.657 |
| TG 12:3_12:3_18:4 | 0.15144 | -2.7232 | 0.00022048 | 3.6566 |
| TG 84:11;O2 | 0.1169 | -3.0967 | 0.00022409 | 3.6496 |
| PS 32:1 | 2.6534 | 1.4078 | 0.0002242 | 3.6494 |
| PE 18:0_22:6 | 0.26037 | -1.9414 | 0.00022738 | 3.6432 |
| HexCer 45:1;O3 | 2.265 | 1.1795 | 0.00022801 | 3.642 |
| TG 75:0;O2 | 0.19224 | -2.379 | 0.00023111 | 3.6362 |
| TG 51:3;O | 0.015166 | -6.043 | 0.00023331 | 3.6321 |
| PC 20:1_16:3 | 23.717 | 4.5679 | 0.00023436 | 3.6301 |
| LPS 33:0;O | 0.38138 | -1.3907 | 0.00023461 | 3.6297 |
| HexCer 38:2;O5 | 0.44689 | -1.162 | 0.00023509 | 3.6288 |
| Cer 18:0;O2/24:1;O | 0.24608 | -2.0228 | 0.00023572 | 3.6276 |
| TG 12:1_12:3_20:6 | 0.39557 | -1.338 | 0.000236 | 3.6271 |
| LPI 18:1; [M-H]- | 0.21741 | -2.2015 | 0.0002406 | 3.6187 |
| PG 18:1_20:2 | 0.45527 | -1.1352 | 0.00024126 | 3.6175 |
| FA 17:0;O2 | 0.30921 | -1.6934 | 0.00024188 | 3.6164 |
| CL 14:0_18:1_16:0_18:2 | 0.0044223 | -7.821 | 0.00024304 | 3.6143 |
| TG O-83:5 | 0.20887 | -2.2593 | 0.00024405 | 3.6125 |
| HexCer 40:4;O2 | 0.49903 | -1.0028 | 0.00024482 | 3.6112 |
| PE 44:1 | 0.48927 | -1.0313 | 0.00024635 | 3.6085 |
| NAT 28:0 | 1.9397 | 0.9558 | 0.00024869 | 3.6043 |
| DG 36:0 | 0.096705 | -3.3703 | 0.00024941 | 3.6031 |
| PR 59:14;O4 | 0.19767 | -2.3388 | 0.00024971 | 3.6026 |
| CL 73:7 | 0.68481 | -0.54622 | 0.00025361 | 3.5958 |
| PG 39:1;O | 0.10556 | -3.2439 | 0.00025543 | 3.5927 |
| DG 42:5 | 0.41586 | -1.2658 | 0.00025755 | 3.5891 |
| PE O-18:0_22:2 | 0.33083 | -1.5958 | 0.00025794 | 3.5885 |
| DG 33:0;O2 | 0.34752 | -1.5248 | 0.00026588 | 3.5753 |
| PC 18:0_21:1 | 0.56432 | -0.82542 | 0.00026667 | 3.574 |
| TG 52:6;O2 | 0.31785 | -1.6536 | 0.00026784 | 3.5721 |
| HexCer 48:6;O4 | 0.38659 | -1.3711 | 0.00026856 | 3.571 |
| PI O-20:0_16:2 | 0.095706 | -3.3852 | 0.00026906 | 3.5702 |
| PS O-15:0_20:4 | 2.4267 | 1.279 | 0.00026984 | 3.5689 |
| TG 85:1;O | 0.14515 | -2.7844 | 0.00027187 | 3.5656 |
| PI 18:1_22:6 | 0.13565 | -2.882 | 0.00028224 | 3.5494 |
| PS 18:0_22:2 | 1.8586 | 0.89418 | 0.0002849 | 3.5453 |
| DG 41:1 | 0.10801 | -3.2107 | 0.0002866 | 3.5427 |
| PIP3 33:8 | 5.8449 | 2.5472 | 0.00028908 | 3.539 |
| PS 18:0_20:0 | 0.46565 | -1.1027 | 0.00029023 | 3.5373 |
| CL 14:0_18:1_16:1_18:1 | 0.041133 | -4.6035 | 0.00029118 | 3.5358 |
| PS 19:0_18:2 | 0.15301 | -2.7083 | 0.00029513 | 3.53 |
| PS 18:0_20:3 | 2.231 | 1.1577 | 0.00029586 | 3.5289 |
| DG O-18:1_24:1 | 0.10939 | -3.1924 | 0.00030012 | 3.5227 |
| PIP3 40:11 | 1.8163 | 0.86101 | 0.00030095 | 3.5215 |
| FA 20:1 | 0.12448 | -3.006 | 0.00030468 | 3.5162 |
| Hex2Cer 28:1;O5 | 3.9129 | 1.9682 | 0.00030906 | 3.51 |
| PE 30:3 | 3.2547 | 1.7025 | 0.0003126 | 3.505 |
| TG 50:6;O2 | 0.30234 | -1.7257 | 0.00031323 | 3.5041 |
| DG O-39:4 | 0.22294 | -2.1653 | 0.00031737 | 3.4984 |
| DG dO-40:2 | 0.33244 | -1.5888 | 0.00031746 | 3.4983 |
| Hex2Cer 30:0;O4 | 0.34583 | -1.5319 | 0.00031893 | 3.4963 |
| LPC 34:3 | 0.37467 | -1.4163 | 0.00032189 | 3.4923 |
| MG 33:2;O | 0.54125 | -0.88564 | 0.0003259 | 3.4869 |
| DG O-36:0;O | 0.089795 | -3.4772 | 0.00032629 | 3.4864 |
| TG 14:0_18:0_19:1 | 0.069769 | -3.8413 | 0.00032804 | 3.4841 |
| ST 27:3;F | 0.55724 | -0.84363 | 0.00033298 | 3.4776 |
| PE 50:7 | 1.493 | 0.57823 | 0.00033345 | 3.477 |
| TG 55:8;O | 0.06891 | -3.8591 | 0.00033601 | 3.4736 |
| PC O-18:0_15:1 | 0.44616 | -1.1644 | 0.00033714 | 3.4722 |
| DG 38:6;O | 0.38111 | -1.3917 | 0.00033715 | 3.4722 |
| PG O-44:8 | 0.59481 | -0.74949 | 0.00034355 | 3.464 |
| DG O-40:6 | 0.30004 | -1.7368 | 0.00034414 | 3.4633 |
| PE 32:3;O | 2.9284 | 1.5501 | 0.00034724 | 3.4594 |
| LPG O-27:0;O | 2.0044 | 1.0032 | 0.00035617 | 3.4483 |
| Cer 18:1;O2/20:0;O | 0.13388 | -2.901 | 0.00035746 | 3.4468 |
| TG 14:1_14:3_19:5 | 0.199 | -2.3292 | 0.00036757 | 3.4347 |
| LPA 18:1 | 0.34448 | -1.5375 | 0.0003682 | 3.4339 |
| PI O-38:6 | 0.65855 | -0.60264 | 0.00037166 | 3.4299 |
| PG 18:1_19:1 | 0.48348 | -1.0485 | 0.00037167 | 3.4298 |
| PE 56:5;O | 0.38355 | -1.3825 | 0.00037638 | 3.4244 |
| Cer 18:1;O2/22:1;O | 0.17463 | -2.5176 | 0.00037716 | 3.4235 |
| TG O-97:5 | 0.13964 | -2.8403 | 0.00037874 | 3.4217 |
| DG O-16:1_20:4 | 0.085423 | -3.5492 | 0.00038494 | 3.4146 |
| ST 23:6;O5 | 1.4609 | 0.54687 | 0.0003939 | 3.4046 |
| CL 71:5 | 0.24504 | -2.0289 | 0.00039601 | 3.4023 |
| PC O-18:1_14:0 | 0.37515 | -1.4144 | 0.00039766 | 3.4005 |
| DG O-18:0_20:4 | 0.29292 | -1.7714 | 0.00039893 | 3.3991 |
| DG O-20:1_16:0 | 0.093729 | -3.4154 | 0.00039973 | 3.3982 |
| PE 18:0_22:5 | 0.20883 | -2.2596 | 0.00040063 | 3.3973 |
| Cer 18:0;O3/35:1;(2OH) | 0.0071637 | -7.1251 | 0.00040238 | 3.3954 |
| CL 70:6 | 0.42056 | -1.2496 | 0.00040446 | 3.3931 |
| Cer 43:0;O3 | 0.36521 | -1.4532 | 0.00040667 | 3.3908 |
| PE 58:6 | 0.41216 | -1.2787 | 0.00040694 | 3.3905 |
| GlcADG 12:0_13:0 | 2.6399 | 1.4005 | 0.00040709 | 3.3903 |
| LPA 30:1 | 3.9975 | 1.9991 | 0.00040862 | 3.3887 |
| PI O-47:9 | 1.6023 | 0.68016 | 0.00040948 | 3.3878 |
| DG O-45:5 | 0.17355 | -2.5266 | 0.00041516 | 3.3818 |
| PA 32:0 | 0.41355 | -1.2739 | 0.00041666 | 3.3802 |
| PG 42:4 | 0.41504 | -1.2687 | 0.0004189 | 3.3779 |
| TG O-69:0;O3 | 0.21594 | -2.2113 | 0.00042327 | 3.3734 |
| PA 18:1_20:1 | 0.26283 | -1.9278 | 0.00042535 | 3.3713 |
| PI-Cer 51:2;O2 | 0.23112 | -2.1133 | 0.00043264 | 3.3639 |
| Hex2Cer 34:4;O3 | 0.25292 | -1.9832 | 0.00043725 | 3.3593 |
| LPA O-30:3 | 1.932 | 0.95009 | 0.00044259 | 3.354 |
| DG O-41:5 | 0.18928 | -2.4014 | 0.00044455 | 3.3521 |
| Car 27:3;O | 1.736 | 0.79574 | 0.0004452 | 3.3514 |
| MG O-19:1 | 1.527 | 0.61071 | 0.00044748 | 3.3492 |
| PIP3 46:11 | 2.6086 | 1.3833 | 0.00045016 | 3.3466 |
| PI O-36:2 | 0.15808 | -2.6612 | 0.00045063 | 3.3462 |
| TG 44:2 | 0.34347 | -1.5418 | 0.00045961 | 3.3376 |
| Cer 43:2;O | 1.7442 | 0.80258 | 0.00046895 | 3.3289 |
| PI O-65:4 | 0.12293 | -3.024 | 0.0004767 | 3.3218 |
| ST 29:3;O2 | 0.50232 | -0.99332 | 0.00047736 | 3.3212 |
| DG O-40:4 | 0.24377 | -2.0364 | 0.00047859 | 3.32 |
| LPE 16:0 | 0.40106 | -1.3181 | 0.00048246 | 3.3165 |
| PG 17:0_18:1 | 0.41588 | -1.2658 | 0.00048302 | 3.316 |
| PE 18:0_20:0 | 0.30887 | -1.6949 | 0.00048531 | 3.314 |
| HexCer 16:0;O2/18:1 | 2.3776 | 1.2495 | 0.00049158 | 3.3084 |
| Cer 18:0;O2/4:0 | 5.6487 | 2.4979 | 0.00049577 | 3.3047 |
| TG 95:5;O | 0.50476 | -0.98634 | 0.00049669 | 3.3039 |
| SL 17:0;O_18:1 | 0.21099 | -2.2448 | 0.00049672 | 3.3039 |
| SM 36:0;O3 | 11.82 | 3.5631 | 0.00049694 | 3.3037 |
| LPA 14:2;O | 2.0633 | 1.045 | 0.00050105 | 3.3001 |
| SM 52:3;O5 | 22.647 | 4.5013 | 0.00050285 | 3.2986 |
| Cer 18:2;O2/24:1;O | 0.18529 | -2.4322 | 0.00050392 | 3.2976 |
| HexCer 18:0;O2/24:0;O | 2.6863 | 1.4256 | 0.00050639 | 3.2955 |
| ST 28:1;O4 | 0.20684 | -2.2734 | 0.00051966 | 3.2843 |
| PIP3 33:7;O | 2.8513 | 1.5116 | 0.00052294 | 3.2815 |
| TG 12:0_12:2_22:7 | 0.40482 | -1.3046 | 0.00052765 | 3.2777 |
| LPI O-11:0 | 3.5134 | 1.8128 | 0.00053112 | 3.2748 |
| DG O-18:1_22:6 | 0.21457 | -2.2204 | 0.00054006 | 3.2676 |
| LPC 33:6 | 1.7006 | 0.76607 | 0.00054307 | 3.2651 |
| CerP 41:4;O2 | 0.70209 | -0.51026 | 0.00054872 | 3.2607 |
| LPI O-32:2 | 0.42782 | -1.2249 | 0.00055312 | 3.2572 |
| Car 34:3 | 0.30399 | -1.7179 | 0.00055412 | 3.2564 |
| PC O-16:1_24:1 | 0.51892 | -0.94643 | 0.00055458 | 3.256 |
| PE 18:1_22:1 | 0.43958 | -1.1858 | 0.0005548 | 3.2559 |
| DG 22:3_22:6 | 0.40124 | -1.3175 | 0.00055484 | 3.2558 |
| PS 18:0_20:2 | 2.4444 | 1.2895 | 0.00055731 | 3.2539 |
| LPC 18:1;O | 2.0214 | 1.0154 | 0.00056402 | 3.2487 |
| **TG 76:2** | 0.27308 | -1.8726 | 0.00056489 | 3.248 |
| **TG 48:0;O3** | 3.1022 | 1.6333 | 0.00056502 | 3.2479 |
| **Cer 18:0;O2/23:1;O** | 0.17157 | -2.5431 | 0.00056631 | 3.2469 |
| **TG 16:0_16:1_20:5** | 2.5299 | 1.3391 | 0.0005711 | 3.2433 |
| **ST 20:2;O** | 0.35717 | -1.4853 | 0.00057322 | 3.2417 |
| **CL 16:0_18:2_16:0_18:3** | 0.0052588 | -7.5711 | 0.000574 | 3.2411 |
| **Cer 46:1;O4** | 0.32664 | -1.6142 | 0.00057475 | 3.2405 |
| **DG 28:1** | 0.17083 | -2.5494 | 0.00057866 | 3.2376 |
| **PI 64:1** | 0.4361 | -1.1973 | 0.00059758 | 3.2236 |
| **LPE O-22:0;O** | 0.44503 | -1.168 | 0.00060506 | 3.2182 |
| **ST 28:2;O** | 0.53374 | -0.90579 | 0.00061114 | 3.2139 |
| **FA 32:4** | 2.8734 | 1.5228 | 0.00061421 | 3.2117 |
| **LPC 34:1** | 0.40974 | -1.2872 | 0.00061566 | 3.2107 |
| **PC 15:0_18:3** | 0.39364 | -1.345 | 0.00062115 | 3.2068 |
| **Cer 42:5;O** | 2.6272 | 1.3935 | 0.00062519 | 3.204 |
| **Hex(5)-HexNAc(3)-Cer 34:1;O2** | 0.52928 | -0.91788 | 0.00062552 | 3.2038 |
| **DG O-16:1_18:1** | 0.083726 | -3.5782 | 0.00062762 | 3.2023 |
| **PR 20:2;O** | 0.51315 | -0.96256 | 0.0006281 | 3.202 |
| **PI 40:7** | 0.08345 | -3.5829 | 0.00063183 | 3.1994 |
| **TG 99:5** | 0.20664 | -2.2748 | 0.00063331 | 3.1984 |
| **TG 51:3;O3** | 0.40637 | -1.2991 | 0.00063473 | 3.1974 |
| **PC O-18:1_24:2** | 0.64939 | -0.62284 | 0.00063728 | 3.1957 |
| **PI 40:4** | 0.65311 | -0.61461 | 0.00064583 | 3.1899 |
| **HexCer 44:5;O2** | 0.4403 | -1.1835 | 0.00064734 | 3.1889 |
| **DG 34:0** | 10.216 | 3.3528 | 0.00065219 | 3.1856 |
| **PE 46:1** | 0.4105 | -1.2846 | 0.00065473 | 3.1839 |
| **DG O-18:2_20:4** | 0.064503 | -3.9545 | 0.00065788 | 3.1819 |
| **ST 30:4;O** | 0.56366 | -0.8271 | 0.00066192 | 3.1792 |
| **NAE 20:1;O2** | 1.5697 | 0.65052 | 0.00066924 | 3.1744 |
| **PC 26:0_18:1** | 0.42034 | -1.2504 | 0.00067192 | 3.1727 |
| **TG 70:3;O3** | 4.0146 | 2.0053 | 0.00067509 | 3.1706 |
| **CerP 52:2;O2** | 0.38623 | -1.3725 | 0.00068335 | 3.1654 |
| **NA 18:2;O3;N;Fe** | 1.6102 | 0.6872 | 0.00069609 | 3.1573 |
| **NAT 15:3** | 1.6102 | 0.6872 | 0.00069609 | 3.1573 |
| **Cer 18:1;O2/22:0;O** | 0.076622 | -3.7061 | 0.00069706 | 3.1567 |
| **CerP 44:0;O5** | 0.39612 | -1.336 | 0.00070075 | 3.1544 |
| **TG 56:1** | 0.02509 | -5.3168 | 0.00070532 | 3.1516 |
| **HexCer 14:0;O2/18:1** | 2.024 | 1.0172 | 0.00070817 | 3.1499 |
| **HexCer 18:0;O2/25:1** | 3.4763 | 1.7975 | 0.00070922 | 3.1492 |
| **HexCer 49:3;O5** | 0.41579 | -1.2661 | 0.00071008 | 3.1487 |
| PE 18:0_22:1 | 0.30927 | -1.6931 | 0.00071097 | 3.1482 |
| CL 70:3 | 0.06009 | -4.0567 | 0.00071294 | 3.1469 |
| HexCer 40:2;O5 | 0.43641 | -1.1963 | 0.00071458 | 3.1459 |
| PG 42:3 | 0.30238 | -1.7256 | 0.00071936 | 3.1431 |
| LPA O-30:2 | 3.6722 | 1.8766 | 0.00072566 | 3.1393 |
| ST 26:0;O8;G | 0.35611 | -1.4896 | 0.0007281 | 3.1378 |
| HexCer 32:2;O2 | 1.8362 | 0.87673 | 0.00073395 | 3.1343 |
| ACer 56:0;O3 | 0.1259 | -2.9896 | 0.00074226 | 3.1294 |
| CL 74:6 | 0.57629 | -0.79512 | 0.00074354 | 3.1287 |
| DG 33:0_18:1 | 0.10671 | -3.2282 | 0.00074391 | 3.1285 |
| PI O-18:1_20:4 | 0.14473 | -2.7885 | 0.00075719 | 3.1208 |
| HexCer 36:5;O2 | 0.31821 | -1.652 | 0.00075922 | 3.1196 |
| PI 14:0_18:0 | 0.17929 | -2.4797 | 0.00076358 | 3.1171 |
| FA 24:0;O3 | 1.511 | 0.59551 | 0.00076836 | 3.1144 |
| TG 14:0_18:0_18:1 | 0.015466 | -6.0148 | 0.00077546 | 3.1104 |
| CL 14:0_18:0_14:0_20:4 | 0.041567 | -4.5884 | 0.0007758 | 3.1103 |
| HexCer 18:0;O2/24:1;O | 2.7267 | 1.4471 | 0.0007761 | 3.1101 |
| HexCer 40:1;O4 | 0.38529 | -1.376 | 0.00078195 | 3.1068 |
| MG O-22:3 | 1.4477 | 0.53378 | 0.00078352 | 3.1059 |
| TG 13:0_14:0_16:0 | 0.48353 | -1.0483 | 0.0007846 | 3.1054 |
| DG 40:6;O | 0.57739 | -0.79239 | 0.0007857 | 3.1047 |
| DG 37:0;O2 | 0.46138 | -1.116 | 0.00078949 | 3.1027 |
| HBMP 16:1_18:0_18:0 | 0.3649 | -1.4544 | 0.00079005 | 3.1023 |
| SL 22:0;O/22:3 | 0.11684 | -3.0974 | 0.00079251 | 3.101 |
| DG 44:9 | 0.41263 | -1.2771 | 0.00079444 | 3.0999 |
| LPA O-27:3 | 0.43168 | -1.212 | 0.00079809 | 3.0979 |
| PG 27:2;O | 1.9351 | 0.9524 | 0.00081212 | 3.0904 |
| ST 28:1;O;Hex;FA 16:0 | 3.0294 | 1.599 | 0.00082082 | 3.0858 |
| SPB 22:0;O2 | 2.7517 | 1.4603 | 0.0008225 | 3.0849 |
| PI-Cer 39:1;O4 | 2.2131 | 1.1461 | 0.00084104 | 3.0752 |
| TG 12:0_12:1_16:3 | 1.6087 | 0.68585 | 0.00084268 | 3.0743 |
| DG 53:3 | 0.26678 | -1.9063 | 0.00084511 | 3.0731 |
| ACer 58:4;O2 | 2.1645 | 1.114 | 0.00084775 | 3.0717 |
| SQMG 18:0 | 1.4389 | 0.52494 | 0.00084863 | 3.0713 |
| PG 20:0_16:1 | 0.46285 | -1.1114 | 0.00084938 | 3.0709 |
| TG 16:0_18:1_18:1 | 0.18075 | -2.4679 | 0.00085043 | 3.0704 |
| HexCer 35:0;O4 | 0.574 | -0.80089 | 0.0008553 | 3.0679 |
| CL 16:0_18:1_18:0_18:2 | 0.0057163 | -7.4507 | 0.00085604 | 3.0675 |
| CerP 51:6;O2 | 0.54591 | -0.87327 | 0.00086074 | 3.0651 |
| CAR 14:1 | 4.2994 | 2.1041 | 0.00086729 | 3.0618 |
| FA 24:1;O | 8.6772 | 3.1172 | 0.00086805 | 3.0615 |
| DG O-35:0 | 0.37816 | -1.4029 | 0.00089178 | 3.0497 |
| CL 71:4 | 0.26531 | -1.9142 | 0.00090903 | 3.0414 |
| DG O-20:2_16:0 | 0.10379 | -3.2683 | 0.00091051 | 3.0407 |
| HexCer 18:2;O2/21:1 | 3.6557 | 1.8702 | 0.00091843 | 3.037 |
| PIP 52:2 | 0.6461 | -0.63018 | 0.00091959 | 3.0364 |
| NAT 17:2;O | 1.5789 | 0.65893 | 0.00092364 | 3.0345 |
| ST 24:3;O3;G | 0.42685 | -1.2282 | 0.00092555 | 3.0336 |
| PE 44:3 | 0.55714 | -0.84389 | 0.00093216 | 3.0305 |
| SM 18:2;O2/24:2 | 9.3633 | 3.227 | 0.00093872 | 3.0275 |
| PS 17:0_21:0 | 0.4864 | -1.0398 | 0.00094103 | 3.0264 |
| Hex2Cer 32:0;O2 | 0.16494 | -2.6 | 0.00094295 | 3.0255 |
| HexCer 42:6;O4 | 0.55737 | -0.84328 | 0.00095108 | 3.0218 |
| PE O-18:0_19:2 | 0.46219 | -1.1134 | 0.00095169 | 3.0215 |
| Cer 44:2;O | 1.525 | 0.60882 | 0.00095437 | 3.0203 |
| PR 45:6;O | 0.086879 | -3.5249 | 0.00095632 | 3.0194 |
| LPC 20:3 | 0.53039 | -0.91489 | 0.00095886 | 3.0182 |
| LPA O-27:4 | 0.22503 | -2.1518 | 0.00097186 | 3.0124 |
| SM 42:2;O2 | 11.726 | 3.5516 | 0.0009738 | 3.0115 |
| PI-Cer 48:2;O3 | 1.4464 | 0.5325 | 0.00097399 | 3.0114 |
| HexCer 18:2;O2/24:2 | 2.2895 | 1.195 | 0.00098253 | 3.0077 |
| PE-Cer 42:6;O3 | 0.3794 | -1.3982 | 0.00099003 | 3.0044 |
| CerP 46:3;O3 | 0.4602 | -1.1197 | 0.0010023 | 2.999 |
| TG 82:10;O2 | 0.15814 | -2.6607 | 0.0010112 | 2.9952 |
| PIP3 45:8;O | 0.67133 | -0.57492 | 0.001015 | 2.9935 |
| HexCer 36:1;O5 | 0.1572 | -2.6694 | 0.0010175 | 2.9925 |
| TG 49:5;O2 | 4.3372 | 2.1168 | 0.0010352 | 2.985 |
| ST 28:2;O;HexNAc | 0.43487 | -1.2014 | 0.0010382 | 2.9837 |
| LPC O-31:3 | 0.40695 | -1.2971 | 0.0010403 | 2.9828 |
| PI-Cer 34:6;O5 | 1.8621 | 0.89689 | 0.001046 | 2.9805 |
| MG O-34:1 | 1.6578 | 0.72924 | 0.0010616 | 2.9741 |
| MG O-31:0;O | 1.4194 | 0.50525 | 0.0010628 | 2.9736 |
| LPA 24:0;O | 0.15497 | -2.69 | 0.0010647 | 2.9728 |
| PE O-34:3 | 1.582 | 0.66177 | 0.0010668 | 2.9719 |
| DG O-46:10 | 3.0114 | 1.5904 | 0.0010686 | 2.9712 |
| DG O-33:0 | 2.2443 | 1.1663 | 0.0010686 | 2.9712 |
| DG 37:2 | 4.6221 | 2.2086 | 0.0010858 | 2.9643 |
| CE 20:5 | 1.4854 | 0.57088 | 0.0010947 | 2.9607 |
| PC 34:3 | 11.176 | 3.4823 | 0.0010961 | 2.9602 |
| DG O-44:9 | 0.34105 | -1.5519 | 0.0010985 | 2.9592 |
| PI 40:5 | 0.075976 | -3.7183 | 0.0011009 | 2.9582 |
| Cer 44:1;O5 | 0.56927 | -0.81282 | 0.0011049 | 2.9567 |
| **Cer 18:0;O2/20:1;O** | 0.18626 | -2.4246 | 0.0011081 | 2.9554 |
| **SHexCer 29:0;O2** | 1.7756 | 0.82833 | 0.0011158 | 2.9524 |
| **Cer 44:6;O2** | 0.28904 | -1.7907 | 0.0011203 | 2.9507 |
| **PG 58:8;O** | 0.47564 | -1.0721 | 0.0011291 | 2.9473 |
| **FA 19:0;O2** | 0.52409 | -0.93211 | 0.0011363 | 2.9445 |
| **TG 88:21;O3** | 0.021649 | -5.5295 | 0.0011403 | 2.943 |
| **DG 43:1;O2** | 0.39841 | -1.3277 | 0.0011431 | 2.9419 |
| **PE 46:0;O** | 0.49145 | -1.0249 | 0.0011471 | 2.9404 |
| **PE-Cer 37:2;O3** | 0.54268 | -0.88184 | 0.0011536 | 2.9379 |
| **MG 34:1;O** | 4.0419 | 2.015 | 0.0011556 | 2.9372 |
| **PA 42:4** | 0.55676 | -0.84488 | 0.0011583 | 2.9362 |
| **PnC 34:1** | 5.4043 | 2.4341 | 0.0011641 | 2.934 |
| **Cer 44:1;O2** | 0.16966 | -2.5593 | 0.0011714 | 2.9313 |
| **PIP3 46:10** | 1.8836 | 0.91353 | 0.0011937 | 2.9231 |
| **TG 48:5;O** | 0.3058 | -1.7094 | 0.0011969 | 2.9219 |
| **HexCer 18:1;O2/16:0** | 2.248 | 1.1686 | 0.0012045 | 2.9192 |
| **PI-Cer 40:1;O2** | 2.0861 | 1.0608 | 0.0012048 | 2.9191 |
| **DG 39:4;O2** | 2.221 | 1.1512 | 0.0012068 | 2.9184 |
| **DG 18:0_32:0** | 0.12929 | -2.9513 | 0.0012376 | 2.9074 |
| **PS 36:2** | 2.4427 | 1.2885 | 0.0012382 | 2.9072 |
| **Car 25:0;O4** | 3.4527 | 1.7877 | 0.0012451 | 2.9048 |
| **LPA 32:1** | 0.27844 | -1.8446 | 0.0012461 | 2.9044 |
| **PS 52:0;O** | 0.37876 | -1.4006 | 0.0012796 | 2.8929 |
| **Car 32:7;O4** | 1.9658 | 0.97511 | 0.0012874 | 2.8903 |
| **HexCer 43:1;O4** | 0.27226 | -1.877 | 0.0012969 | 2.8871 |
| **TG 76:11;O3** | 2.6257 | 1.3927 | 0.0013016 | 2.8855 |
| **PG O-36:2** | 0.33487 | -1.5783 | 0.0013294 | 2.8764 |
| **TG 75:2;O3** | 0.079215 | -3.6581 | 0.0013294 | 2.8764 |
| **PE O-18:1_21:2** | 0.56919 | -0.81302 | 0.0013334 | 2.875 |
| **PC O-20:3_16:4** | 4.9284 | 2.3011 | 0.0013405 | 2.8727 |
| **GP 30:0** | 4.8018 | 2.2636 | 0.0013418 | 2.8723 |
| **Car 32:6;O3** | 1.5297 | 0.61326 | 0.0013563 | 2.8676 |
| **PC 30:0** | 6.6438 | 2.732 | 0.0013584 | 2.867 |
| **DG 16:0_32:2** | 0.29212 | -1.7754 | 0.0013766 | 2.8612 |
| **SM 20:0;O2/16:2** | 2.1835 | 1.1267 | 0.0013858 | 2.8583 |
| **PE 30:1;O** | 0.37517 | -1.4144 | 0.0013865 | 2.8581 |
| **PI O-67:5** | 0.2043 | -2.2912 | 0.0013918 | 2.8564 |
| **PS 24:4_18:5** | 3.7142 | 1.8931 | 0.0013929 | 2.8561 |
| **PG O-39:0** | 0.62446 | -0.67931 | 0.0013944 | 2.8556 |
| **CL 16:0_20:5_18:0_18:2** | 0.015307 | -6.0297 | 0.0013948 | 2.8555 |
| **PC O-16:0_12:0** | 0.38981 | -1.3592 | 0.0013968 | 2.8549 |
| DG O-19:1_18:1 | 0.29339 | -1.7691 | 0.0014246 | 2.8463 |
| PI-Cer 34:1;O2 | 0.40691 | -1.2972 | 0.0014385 | 2.8421 |
| HexCer 18:1;O2/4:0 | 4.0358 | 2.0128 | 0.001452 | 2.838 |
| SM 34:1;O2 | 33.927 | 5.0843 | 0.001456 | 2.8368 |
| TG 16:1_18:1_18:1 | 0.25707 | -1.9598 | 0.0014634 | 2.8346 |
| LPS O-16:1 | 2.4592 | 1.2982 | 0.0014714 | 2.8323 |
| PC O-68:6 | 22.661 | 4.5022 | 0.0014773 | 2.8305 |
| ACer 71:3;O6 | 0.50826 | -0.97637 | 0.0014853 | 2.8282 |
| DG 18:1_24:1 | 0.13436 | -2.8958 | 0.0014941 | 2.8256 |
| LPC 34:0 | 0.53391 | -0.90534 | 0.0014997 | 2.824 |
| PC 16:0_18:4 | 16.973 | 4.0852 | 0.0015059 | 2.8222 |
| LPE 12:0 | 1.5835 | 0.6631 | 0.0015072 | 2.8218 |
| PG 45:6;O | 0.21604 | -2.2106 | 0.0015185 | 2.8186 |
| SM 44:5;O6 | 13.188 | 3.7211 | 0.0015229 | 2.8173 |
| LPG 25:0;O | 2.6346 | 1.3976 | 0.0015283 | 2.8158 |
| PS 20:0_16:2 | 2.273 | 1.1846 | 0.0015313 | 2.8149 |
| DG 16:0_18:2 | 0.11378 | -3.1356 | 0.0015328 | 2.8145 |
| PC 34:0;O | 16.734 | 4.0647 | 0.0015355 | 2.8138 |
| TG O-10:0_10:0_10:0 | 6.414 | 2.6812 | 0.0015403 | 2.8124 |
| TG 50:1;O2 | 0.50946 | -0.97297 | 0.0015479 | 2.8102 |
| SM 18:0;O2/20:1 | 7.401 | 2.8877 | 0.0015493 | 2.8099 |
| PG 37:0 | 0.14841 | -2.7523 | 0.0015547 | 2.8084 |
| HexCer 18:1;O2/24:0 | 3.4782 | 1.7983 | 0.0015559 | 2.808 |
| LPC 28:0 | 0.46631 | -1.1006 | 0.0015595 | 2.807 |
| DG 43:2;O | 3.3363 | 1.7382 | 0.0015639 | 2.8058 |
| LPC 16:3;O | 2.4262 | 1.2787 | 0.00158 | 2.8013 |
| PG 18:0_20:2 | 0.36253 | -1.4638 | 0.0015898 | 2.7987 |
| TG 76:4;O | 0.39613 | -1.3359 | 0.0015934 | 2.7977 |
| PI 30:1 | 0.53945 | -0.89043 | 0.0016078 | 2.7938 |
| HexCer 42:5;O2 | 0.47652 | -1.0694 | 0.0016085 | 2.7936 |
| SL 18:0;O/19:1 | 0.37157 | -1.4283 | 0.0016093 | 2.7934 |
| PG 38:0;O | 0.13197 | -2.9218 | 0.0016102 | 2.7931 |
| ST 28:1;O3 | 0.46438 | -1.1066 | 0.0016238 | 2.7895 |
| SM 43:6;O2 | 11.322 | 3.501 | 0.0016285 | 2.7882 |
| DG O-46:7 | 2.2676 | 1.1812 | 0.0016319 | 2.7873 |
| DG 16:1_20:1 | 12.289 | 3.6193 | 0.001634 | 2.7867 |
| Car 32:2;O2 | 1.5665 | 0.64754 | 0.0016362 | 2.7862 |
| LPI 18:0 | 0.46865 | -1.0934 | 0.0016371 | 2.7859 |
| PE-Cer 36:3;O2 | 0.5735 | -0.80214 | 0.0016518 | 2.782 |
| ST 28:3;O | 0.4602 | -1.1197 | 0.0016633 | 2.779 |
| CL 69:4 | 0.28721 | -1.7998 | 0.0016671 | 2.778 |
| DG 59:5 | 0.10505 | -3.2508 | 0.0016725 | 2.7766 |
| CerP 14:0;O2/10:0 | 0.33391 | -1.5825 | 0.00168 | 2.7747 |
| DG O-44:8 | 0.44595 | -1.165 | 0.0016924 | 2.7715 |
| PE 17:1_18:1 | 0.37548 | -1.4132 | 0.0017 | 2.7696 |
| LPI 34:2 | 0.23261 | -2.104 | 0.0017012 | 2.7692 |
| PE 3:0_16:1 | 1.6068 | 0.6842 | 0.0017043 | 2.7685 |
| DG 38:5;O | 0.49941 | -1.0017 | 0.0017103 | 2.7669 |
| DG 57:4 | 0.049568 | -4.3344 | 0.0017406 | 2.7593 |
| LPC 16:0;O | 1.7796 | 0.83154 | 0.0017475 | 2.7576 |
| NA 25:4;O2 | 5.3369 | 2.416 | 0.0017672 | 2.7527 |
| PG O-36:0 | 1.506 | 0.5907 | 0.0017716 | 2.7516 |
| TG 81:1;O3 | 0.56434 | -0.82536 | 0.0017902 | 2.7471 |
| PE-Cer 36:1;O4 | 0.54581 | -0.87353 | 0.0017994 | 2.7449 |
| SM 18:0;O2/26:1 | 6.3477 | 2.6662 | 0.0018007 | 2.7445 |
| Cer 51:6;O2 | 0.39744 | -1.3312 | 0.0018044 | 2.7437 |
| Cer 18:1;3O/23:1;(2OH) | 0.11983 | -3.0609 | 0.0018066 | 2.7431 |
| PG 56:8;O | 0.12502 | -2.9998 | 0.0018101 | 2.7423 |
| SM 18:3;O2/26:2 | 17.313 | 4.1138 | 0.0018103 | 2.7422 |
| SM 30:2;O5 | 14.625 | 3.8703 | 0.0018108 | 2.7421 |
| Cer 40:6;O3 | 0.40557 | -1.302 | 0.0018109 | 2.7421 |
| PC O-18:1_20:1 | 0.39921 | -1.3248 | 0.0018253 | 2.7387 |
| SM 33:6;O2 | 9.3328 | 3.2223 | 0.0018286 | 2.7379 |
| DG O-20:2_16:1 | 0.051446 | -4.2808 | 0.0018484 | 2.7332 |
| ST 27:6;O | 0.27906 | -1.8414 | 0.0018488 | 2.7331 |
| Cer 18:0;O2/26:1;O | 0.3667 | -1.4473 | 0.0018555 | 2.7315 |
| NAE 19:0;O | 1.5929 | 0.67169 | 0.001859 | 2.7307 |
| ST 29:7;O | 0.043335 | -4.5283 | 0.0018606 | 2.7303 |
| NA 18:2;O3 | 1.5542 | 0.63615 | 0.0018729 | 2.7275 |
| LPC 30:3 | 1.6365 | 0.71063 | 0.0018828 | 2.7252 |
| PC 40:1 | 19.999 | 4.3218 | 0.0018863 | 2.7244 |
| PS O-40:7 | 2.6133 | 1.3859 | 0.001887 | 2.7242 |
| Cer 18:1;O3/21:1;(2OH) | 0.11408 | -3.1318 | 0.0018943 | 2.7225 |
| PI O-67:6 | 0.64219 | -0.63893 | 0.0019008 | 2.7211 |
| Car 30:6;O2 | 1.5702 | 0.65094 | 0.0019118 | 2.7186 |
| FA 29:4;O | 1.7269 | 0.78815 | 0.0019279 | 2.7149 |
| PG 21:1_21:1 | 1.6322 | 0.70685 | 0.0019434 | 2.7114 |
| DG 41:6;O | 0.65158 | -0.61798 | 0.0019516 | 2.7096 |
| LPC 19:1 | 0.4222 | -1.244 | 0.0019518 | 2.7096 |
| DG 18:1_19:1 | 0.1725 | -2.5353 | 0.0019566 | 2.7085 |
| CE 18:1 | 0.27606 | -1.8569 | 0.0019751 | 2.7044 |
| NAT 22:2;O2 | 1.4689 | 0.55472 | 0.0019751 | 2.7044 |
| HexCer 20:0;O2/16:1 | 1.8454 | 0.88392 | 0.0019762 | 2.7042 |
| TG 44:6;O2 | 0.08066 | -3.632 | 0.0019789 | 2.7036 |
| SM 18:0;O2/25:1 | 9.8 | 3.2928 | 0.0019818 | 2.7029 |
| PA O-50:6 | 3.5481 | 1.827 | 0.0019863 | 2.702 |
| SM 40:0;O5 | 16.671 | 4.0593 | 0.0020082 | 2.6972 |
| PI 28:1 | 1.8554 | 0.89171 | 0.0020124 | 2.6963 |
| PR 58:14;O3 | 0.075688 | -3.7238 | 0.002014 | 2.6959 |
| SM 32:2;O4 | 18.416 | 4.2029 | 0.0020309 | 2.6923 |
| DG O-33:0;O2 | 2.8544 | 1.5132 | 0.0020341 | 2.6916 |
| PE 42:5 | 0.15892 | -2.6536 | 0.0020343 | 2.6916 |
| PI 18:0_22:3 | 2.0973 | 1.0685 | 0.0020461 | 2.6891 |
| TG 53:1 | 0.3305 | -1.5973 | 0.0020489 | 2.6885 |
| SM 38:3;O6 | 8.6724 | 3.1164 | 0.0020628 | 2.6855 |
| PE-Cer 32:1;O;P | 0.60158 | -0.73317 | 0.0020708 | 2.6839 |
| FA 17:1 | 0.18565 | -2.4294 | 0.0020727 | 2.6835 |
| PE-Cer 32:3;O5 | 2.0776 | 1.0549 | 0.0020797 | 2.682 |
| LPE 18:0 | 0.46252 | -1.1124 | 0.0020859 | 2.6807 |
| TG 12:0_12:3_22:7 | 0.1168 | -3.0979 | 0.0020881 | 2.6802 |
| PC 16:0_18:3 | 25.662 | 4.6816 | 0.0020957 | 2.6787 |
| Cer 45:1;O3 | 0.62596 | -0.67587 | 0.0021049 | 2.6768 |
| NAT 33:0 | 3.069 | 1.6178 | 0.0021302 | 2.6716 |
| PG 48:2;O | 0.37682 | -1.408 | 0.0021405 | 2.6695 |
| DG 27:0_21:1 | 0.0099618 | -6.6494 | 0.0021434 | 2.6689 |
| PC 14:0_18:3 | 7.4208 | 2.8916 | 0.0021573 | 2.6661 |
| PC 14:0_16:1 | 0.39731 | -1.3317 | 0.0021686 | 2.6638 |
| SM 45:5;O5 | 30.194 | 4.9162 | 0.0021741 | 2.6627 |
| Cer 44:3;O | 0.48675 | -1.0387 | 0.0021813 | 2.6613 |
| PC 31:0;O | 11.513 | 3.5252 | 0.0022003 | 2.6575 |
| SM 54:4;O3 | 11.513 | 3.5252 | 0.0022003 | 2.6575 |
| PI 15:0_18:1 | 0.57721 | -0.79282 | 0.0022196 | 2.6537 |
| HexCer 18:2;O2/22:6 | 1.7372 | 0.79679 | 0.002228 | 2.6521 |
| PG 14:0_16:0 | 0.21154 | -2.241 | 0.0022301 | 2.6517 |
| SM 16:0;O2/18:1 | 10.238 | 3.3559 | 0.0022415 | 2.6495 |
| MG O-29:0;O | 2.0427 | 1.0305 | 0.0022529 | 2.6473 |
| Cer 18:2;O2/26:7;O | 0.61291 | -0.70625 | 0.002265 | 2.6449 |
| MG O-32:0;O | 0.3941 | -1.3434 | 0.0022804 | 2.642 |
| TG 83:0;O | 0.21661 | -2.2068 | 0.0022819 | 2.6417 |
| DG 26:0 | 0.29605 | -1.7561 | 0.0022844 | 2.6412 |
| PG 40:7 | 5.8101 | 2.5386 | 0.0023076 | 2.6368 |
| SM 36:1;O3 | 10.55 | 3.3992 | 0.0023192 | 2.6347 |
| SM 54:3;O3 | 10.552 | 3.3994 | 0.0023268 | 2.6332 |
| MG 25:2 | 3.2446 | 1.6981 | 0.0023324 | 2.6322 |
| HexCer 44:2;O5 | 0.56521 | -0.82314 | 0.0023346 | 2.6318 |
| TG 16:3_22:7_22:7 | 5.523 | 2.4655 | 0.0023492 | 2.6291 |
| Cer 50:3;O | 0.36554 | -1.4519 | 0.0023581 | 2.6274 |
| SM 53:0;O3 | 10.328 | 3.3684 | 0.0023642 | 2.6263 |
| Car 22:3;O2 | 0.20354 | -2.2966 | 0.0023704 | 2.6252 |
| ST 28:0;O3 | 0.15557 | -2.6844 | 0.0023724 | 2.6248 |
| PS 18:0_17:1 | 2.94 | 1.5558 | 0.002373 | 2.6247 |
| HexCer 41:4;O4 | 0.55374 | -0.85271 | 0.0023798 | 2.6235 |
| DG O-58:5 | 0.035445 | -4.8183 | 0.0023857 | 2.6224 |
| PS 40:5 | 4.7743 | 2.2553 | 0.0023908 | 2.6215 |
| SM 40:1;O2 | 46.38 | 5.5354 | 0.0023928 | 2.6211 |
| ST 44:4;O2 | 0.34488 | -1.5358 | 0.0024031 | 2.6192 |
| PC 14:0_18:0 | 11.374 | 3.5077 | 0.0024196 | 2.6163 |
| DG 16:0_28:0 | 0.17566 | -2.5092 | 0.002436 | 2.6133 |
| PI O-18:1_22:6 | 0.69865 | -0.51735 | 0.002437 | 2.6132 |
| SM 42:1;O4 | 10.626 | 3.4095 | 0.0024454 | 2.6117 |
| PC O-18:1_21:2 | 9.8817 | 3.3048 | 0.0024505 | 2.6107 |
| SM 42:1;O2 | 9.6264 | 3.267 | 0.002463 | 2.6085 |
| SM 54:2;O5 | 9.8043 | 3.2934 | 0.0024673 | 2.6078 |
| ST 28:2;O4;G | 0.5855 | -0.77225 | 0.0024842 | 2.6048 |
| TG 14:0_16:0_19:1 | 0.097524 | -3.3581 | 0.0024878 | 2.6042 |
| SM 18:2;O2/22:2 | 31.956 | 4.998 | 0.0024969 | 2.6026 |
| HexCer 39:5;O2 | 0.56294 | -0.82895 | 0.0025105 | 2.6002 |
| FA 18:2 | 0.13801 | -2.8572 | 0.0025265 | 2.5975 |
| PC O-52:9 | 5.8322 | 2.544 | 0.0025315 | 2.5966 |
| LPC 18:0 | 0.39657 | -1.3343 | 0.0025585 | 2.592 |
| SM 40:1;O5 | 18.577 | 4.2154 | 0.0025606 | 2.5917 |
| PE 40:4 | 0.15862 | -2.6563 | 0.0025641 | 2.5911 |
| PS 32:0; PS 14:0-18:0; [M-H]- | 2.366 | 1.2424 | 0.0025758 | 2.5891 |
| SHexCer 31:4;O2 | 0.28994 | -1.7862 | 0.0025784 | 2.5886 |
| SM 18:3;O2/25:1 | 66.017 | 6.0448 | 0.0026024 | 2.5846 |
| PC 32:5 | 8.8464 | 3.1451 | 0.0026092 | 2.5835 |
| PC 45:2 | 8.9284 | 3.1584 | 0.002614 | 2.5827 |
| NA 41:2;O4;N | 1.7373 | 0.79682 | 0.0026166 | 2.5823 |
| FA 18:0;O;[cy] | 0.24742 | -2.0149 | 0.0026184 | 2.582 |
| SM 32:7;O2 | 33.634 | 5.0718 | 0.0026191 | 2.5819 |
| PC 38:6;O | 13.526 | 3.7577 | 0.0026253 | 2.5808 |
| HexCer 42:1;O5 | 0.57879 | -0.7889 | 0.0026491 | 2.5769 |
| NAT 20:3;O3 | 1.6797 | 0.74822 | 0.0026496 | 2.5768 |
| PE 12:0_18:1 | 0.39925 | -1.3246 | 0.002651 | 2.5766 |
| TG 98:9;O2 | 0.4435 | -1.173 | 0.002652 | 2.5764 |
| PS 20:0_18:1 | 1.5637 | 0.64496 | 0.0026534 | 2.5762 |
| ST 29:4;O3 | 0.28586 | -1.8066 | 0.0026603 | 2.5751 |
| TG 8:0_8:0_17:2 | 1.6344 | 0.70875 | 0.0026666 | 2.574 |
| LPC 16:0 | 0.55911 | -0.8388 | 0.0026708 | 2.5734 |
| TG 14:0_16:0_18:4 | 1.9373 | 0.95404 | 0.0026725 | 2.5731 |
| SM 46:1;O2 | 8.5835 | 3.1016 | 0.0026738 | 2.5729 |
| PC O-18:5_15:1 | 8.8672 | 3.1485 | 0.0026741 | 2.5728 |
| PC 45:2;O | 8.5706 | 3.0994 | 0.0026794 | 2.572 |
| SM 41:1;O2 | 45.977 | 5.5228 | 0.0026812 | 2.5717 |
| PC O-18:1_24:1 | 0.31632 | -1.6606 | 0.0027039 | 2.568 |
| PE 24:0_18:1 | 0.45097 | -1.1489 | 0.0027296 | 2.5639 |
| Cer 18:3;O2/30:7;O | 2.3454 | 1.2298 | 0.0027607 | 2.559 |
| PS 14:0_20:0 | 0.33846 | -1.5629 | 0.0027622 | 2.5587 |
| SM 32:2;O2 | 18.472 | 4.2072 | 0.002764 | 2.5585 |
| NA 42:2;O4;N | 0.19898 | -2.3293 | 0.002768 | 2.5578 |
| PE 22:0_18:1 | 0.39893 | -1.3258 | 0.0027717 | 2.5572 |
| Cer 19:1;O2/24:1;O | 0.14136 | -2.8225 | 0.0027761 | 2.5566 |
| PA O-47:3 | 16.784 | 4.069 | 0.0027763 | 2.5565 |
| LPC 26:0;O | 0.66661 | -0.58508 | 0.0027782 | 2.5562 |
| SM 18:0;O2/21:1 | 13.143 | 3.7162 | 0.0027785 | 2.5562 |
| SM 33:3;O3 | 9.8263 | 3.2966 | 0.002786 | 2.555 |
| SM 44:4;O6 | 8.6126 | 3.1065 | 0.0027877 | 2.5548 |
| HexCer 37:0;O4 | 0.50083 | -0.99759 | 0.0027938 | 2.5538 |
| PC 43:2;O | 8.0618 | 3.0111 | 0.0027958 | 2.5535 |
| TG 14:0_18:0_18:2 | 0.072845 | -3.779 | 0.0027983 | 2.5531 |
| SM 34:1;O5 | 11.01 | 3.4608 | 0.0028173 | 2.5502 |
| PC 34:3;O | 10.872 | 3.4425 | 0.0028233 | 2.5492 |
| TG 79:0;O2 | 0.22252 | -2.168 | 0.0028421 | 2.5464 |
| SM 50:3;O3 | 8.2174 | 3.0387 | 0.0028449 | 2.5459 |
| SM 33:2;O3 | 19.779 | 4.3059 | 0.0028461 | 2.5457 |
| PE 40:7 | 1.8093 | 0.85543 | 0.0028475 | 2.5455 |
| TG 55:2;O | 2.0012 | 1.0009 | 0.0028486 | 2.5454 |
| SM 41:0;O2 | 10.75 | 3.4262 | 0.0028666 | 2.5426 |
| SM 40:6;O6 | 5.8781 | 2.5553 | 0.0028702 | 2.5421 |
| HBMP 16:1_18:1_18:0 | 0.27182 | -1.8793 | 0.002894 | 2.5385 |
| Cer 32:1;O6 | 3.1836 | 1.6707 | 0.0028951 | 2.5383 |
| SM 42:6;O5 | 7.9484 | 2.9907 | 0.0028991 | 2.5377 |
| PS 42:3;O | 1.5475 | 0.62996 | 0.002903 | 2.5372 |
| PIP2 62:4 | 1.466 | 0.55189 | 0.0029047 | 2.5369 |
| PC 43:7 | 7.8754 | 2.9774 | 0.0029146 | 2.5354 |
| PE 18:1_24:2 | 0.60882 | -0.71592 | 0.0029294 | 2.5332 |
| PC 31:2;O | 17.239 | 4.1076 | 0.0029317 | 2.5329 |
| SM 36:1;O5 | 13.423 | 3.7467 | 0.0029541 | 2.5296 |
| DG O-42:5 | 0.24206 | -2.0465 | 0.0029638 | 2.5282 |
| SM 32:2;O3 | 13.994 | 3.8068 | 0.0029647 | 2.528 |
| PG 40:3 | 0.3982 | -1.3284 | 0.0029773 | 2.5262 |
| SM 18:0;O2/24:2 | 8.1704 | 3.0304 | 0.0029818 | 2.5255 |
| CerP 44:0;O6 | 2.167 | 1.1157 | 0.0029879 | 2.5246 |
| PC 48:0 | 7.6928 | 2.9435 | 0.0029894 | 2.5244 |
| SHexCer 43:2;O2 | 1.7992 | 0.84736 | 0.0029963 | 2.5234 |
| SM 40:4;O2 | 8.1534 | 3.0274 | 0.0030058 | 2.522 |
| PE-Cer 32:3;O3 | 1.7814 | 0.833 | 0.0030133 | 2.521 |
| MIPC 48:2;O4 | 0.65133 | -0.61854 | 0.0030221 | 2.5197 |
| ST 18:2;O3;G | 0.5341 | -0.90481 | 0.003026 | 2.5191 |
| LPC O-34:0;O | 0.31369 | -1.6726 | 0.0030299 | 2.5186 |
| PC 18:0_20:5 | 20.874 | 4.3837 | 0.0030315 | 2.5183 |
| Cer 39:6;O2 | 0.55154 | -0.85848 | 0.0030466 | 2.5162 |
| PE O-18:0_20:1 | 0.55556 | -0.84799 | 0.0030559 | 2.5149 |
| ST 33:5;O6 | 0.28841 | -1.7938 | 0.0030578 | 2.5146 |
| Hex(4)-HexNAc-Fuc-Cer 42:1;O2 | 1.7205 | 0.78287 | 0.0030635 | 2.5138 |
| PC O-42:6 | 7.2382 | 2.8556 | 0.0030827 | 2.5111 |
| HexCer 42:3;O3 | 0.49077 | -1.0269 | 0.0030835 | 2.511 |
| SM 40:6;O2 | 3.3911 | 1.7618 | 0.0030862 | 2.5106 |
| LPC 18:4;O | 2.8123 | 1.4918 | 0.0030944 | 2.5094 |
| DG 37:2;O2 | 0.23232 | -2.1058 | 0.0030998 | 2.5087 |
| SM 45:3;O4 | 8.3745 | 3.066 | 0.003103 | 2.5082 |
| Cer 9:0;O2/16:4 | 1.9293 | 0.94807 | 0.0031091 | 2.5074 |
| HexCer 40:0;O4 | 0.37426 | -1.4179 | 0.003113 | 2.5068 |
| DG O-44:2 | 1.8767 | 0.90817 | 0.0031132 | 2.5068 |
| SM 45:2;O3 | 12.727 | 3.6699 | 0.0031207 | 2.5058 |
| ST 28:3;O3;Cl | 0.31076 | -1.6861 | 0.0031282 | 2.5047 |
| DG 50:4 | 0.053121 | -4.2346 | 0.0031642 | 2.4997 |
| PE-Cer 44:1;O4 | 0.58139 | -0.78243 | 0.0031706 | 2.4989 |
| PS 23:0_18:5 | 0.32168 | -1.6363 | 0.003175 | 2.4983 |
| SM 45:5;O4 | 19.915 | 4.3158 | 0.0032001 | 2.4948 |
| FA 24:2 | 0.43267 | -1.2087 | 0.003201 | 2.4947 |
| SM 34:0;O3 | 5.8463 | 2.5475 | 0.0032035 | 2.4944 |
| SM 14:3;O2/18:2 | 13.672 | 3.7731 | 0.0032128 | 2.4931 |
| Car 33:6;O4 | 1.5584 | 0.64007 | 0.0032218 | 2.4919 |
| DG 17:1_18:1 | 0.15199 | -2.718 | 0.0032352 | 2.4901 |
| DG 42:7 | 0.65751 | -0.60491 | 0.0032412 | 2.4893 |
| TG 15:1_18:1_18:1 | 3.2843 | 1.7156 | 0.0032493 | 2.4882 |
| PC 36:2 | 7.4555 | 2.8983 | 0.003274 | 2.4849 |
| PI O-55:3 | 0.43897 | -1.1878 | 0.0032891 | 2.4829 |
| HexCer 38:4;O4 | 5.5969 | 2.4846 | 0.0032895 | 2.4829 |
| DG 56:6 | 0.25572 | -1.9674 | 0.0032968 | 2.4819 |
| HexCer 18:0;O2/26:1 | 3.2271 | 1.6902 | 0.003311 | 2.48 |
| HexCer 41:3;O2 | 0.28101 | -1.8313 | 0.0033241 | 2.4783 |
| PE 44:1;O | 0.55483 | -0.84988 | 0.0033438 | 2.4758 |
| ST 43:3;O2 | 0.15841 | -2.6583 | 0.0033595 | 2.4737 |
| PS 40:6 | 2.9948 | 1.5825 | 0.0033641 | 2.4731 |
| PE 54:5;O | 0.40599 | -1.3005 | 0.0033694 | 2.4725 |
| PC O-18:1_17:0 | 0.37611 | -1.4108 | 0.0033818 | 2.4709 |
| DG O-56:5 | 0.011953 | -6.3865 | 0.0033844 | 2.4705 |
| PC O-38:3 | 6.7904 | 2.7635 | 0.0033884 | 2.47 |
| PA 23:2;O3 | 16.631 | 4.0558 | 0.0033909 | 2.4697 |
| ST 18:3;O2;GlcA | 0.28197 | -1.8264 | 0.0034046 | 2.4679 |
| SM 50:2;O3 | 5.3281 | 2.4136 | 0.0034127 | 2.4669 |
| PC 28:4;O | 29.362 | 4.8759 | 0.0034403 | 2.4634 |
| HexCer 32:6;O3 | 1.5442 | 0.62686 | 0.0034408 | 2.4633 |
| TG 15:0_16:0_17:1 | 0.015938 | -5.9714 | 0.0034659 | 2.4602 |
| SM 44:6;O6 | 13.522 | 3.7573 | 0.0034662 | 2.4601 |
| PC 36:1 | 6.0544 | 2.598 | 0.0034741 | 2.4592 |
| LPA O-27:6 | 0.36238 | -1.4644 | 0.0034841 | 2.4579 |
| PC 42:3 | 6.6782 | 2.7395 | 0.0035052 | 2.4553 |
| Glc-GP 38:4 | 7.4362 | 2.8946 | 0.0035066 | 2.4551 |
| SM 44:1;O4 | 6.4239 | 2.6834 | 0.0035118 | 2.4545 |
| LPC 26:0 | 0.50085 | -0.99756 | 0.0035239 | 2.453 |
| NAE 27:1;O | 2.116 | 1.0814 | 0.0035322 | 2.452 |
| PI-Cer 34:1;O6 | 2.3651 | 1.2419 | 0.0035413 | 2.4508 |
| TG 48:2 | 0.14129 | -2.8233 | 0.0035418 | 2.4508 |
| TG 60:5;O2 | 0.43427 | -1.2033 | 0.0035565 | 2.449 |
| PI-Cer 34:0;O4 | 0.62359 | -0.68134 | 0.0035825 | 2.4458 |
| TG 86:4;O | 0.44272 | -1.1755 | 0.0036002 | 2.4437 |
| PE 18:0_22:2 | 0.43281 | -1.2082 | 0.0036518 | 2.4375 |
| PC 2:0_16:4 | 19.456 | 4.2821 | 0.0036587 | 2.4367 |
| PI-Cer 36:1;O4 | 0.47364 | -1.0781 | 0.0036877 | 2.4332 |
| SM 21:3;O2/24:2 | 17.054 | 4.092 | 0.0036915 | 2.4328 |
| PC 36:5 | 6.6253 | 2.728 | 0.0036997 | 2.4318 |
| CL 14:0_18:1_16:0_18:1 | 0.014024 | -6.156 | 0.003712 | 2.4304 |
| GP 32:0 | 7.5521 | 2.9169 | 0.0037165 | 2.4299 |
| PIM1 35:3 | 2.9455 | 1.5585 | 0.0037319 | 2.4281 |
| TG 59:2 | 0.031734 | -4.9778 | 0.0037522 | 2.4257 |
| PE-Cer 33:2;O2 | 1.7931 | 0.84242 | 0.0037639 | 2.4244 |
| CL 14:0_18:2_16:0_18:2 | 0.0059326 | -7.3971 | 0.0037641 | 2.4243 |
| Car 26:6 | 2.7668 | 1.4682 | 0.0037677 | 2.4239 |
| SM 45:1;O3 | 9.0433 | 3.1769 | 0.0037776 | 2.4228 |
| PC 46:2 | 6.0359 | 2.5936 | 0.0038112 | 2.4189 |
| CerP 45:0;O6 | 1.8901 | 0.9185 | 0.0038261 | 2.4172 |
| FA 20:0 | 1.5136 | 0.59796 | 0.0038425 | 2.4154 |
| SM 28:3;O2 | 11.19 | 3.4841 | 0.0038565 | 2.4138 |
| ST 27:1;O;S | 4.5993 | 2.2014 | 0.0038827 | 2.4109 |
| HexCer 34:5;O2 | 0.43237 | -1.2096 | 0.0038829 | 2.4108 |
| DG 57:5 | 0.079968 | -3.6444 | 0.0038897 | 2.4101 |
| HexCer 39:3;O3 | 0.54749 | -0.8691 | 0.0039045 | 2.4084 |
| TG 96:9;O2 | 2.346 | 1.2302 | 0.0039085 | 2.408 |
| SM 54:0;O3 | 5.9592 | 2.5751 | 0.0039098 | 2.4078 |
| CerP 40:3;O2 | 4.2103 | 2.0739 | 0.0039102 | 2.4078 |
| TG 60:2 | 0.040602 | -4.6223 | 0.0039123 | 2.4076 |
| TG 51:3;O2 | 0.51073 | -0.96938 | 0.0039144 | 2.4073 |
| PA 44:4 | 0.56495 | -0.8238 | 0.0039192 | 2.4068 |
| SM 15:3;O2/18:2 | 13.864 | 3.7932 | 0.0039206 | 2.4066 |
| DG 42:0;O4;N | 0.41937 | -1.2537 | 0.0039356 | 2.405 |
| TG 79:0;O3 | 0.44646 | -1.1634 | 0.0039477 | 2.4037 |
| Cer 42:4;O2 | 1.5418 | 0.62466 | 0.003986 | 2.3995 |
| Cer 18:1;O2/23:1;O | 0.13671 | -2.8708 | 0.0039973 | 2.3982 |
| PG 15:0_18:1 | 0.54156 | -0.8848 | 0.0040037 | 2.3975 |
| SM 20:0;O2/16:1 | 9.6276 | 3.2672 | 0.004031 | 2.3946 |
| PC 50:2 | 9.4552 | 3.2411 | 0.0040336 | 2.3943 |
| DG O-43:2 | 1.8848 | 0.91443 | 0.0040442 | 2.3932 |
| PG O-38:0 | 0.34576 | -1.5322 | 0.0040457 | 2.393 |
| DG O-54:5 | 0.027218 | -5.1993 | 0.0040963 | 2.3876 |
| PIP3 37:6 | 0.47651 | -1.0694 | 0.0041243 | 2.3847 |
| PC 16:0_18:0 | 4.9174 | 2.2979 | 0.0041637 | 2.3805 |
| PC 25:1;O | 5.5019 | 2.4599 | 0.0041784 | 2.379 |
| HexCer 21:3;O2/24:2 | 2.1754 | 1.1213 | 0.0042116 | 2.3755 |
| PC 23:1;O | 4.9247 | 2.3 | 0.0042139 | 2.3753 |
| TG 97:7;O2 | 1.8551 | 0.89147 | 0.0042295 | 2.3737 |
| PC 43:11;O | 54.087 | 5.7572 | 0.0042391 | 2.3727 |
| SM 44:3;O2 | 5.4458 | 2.4451 | 0.0042446 | 2.3722 |
| CerP 32:4;O2 | 2.099 | 1.0697 | 0.0042467 | 2.3719 |
| DG O-40:5 | 0.12708 | -2.9762 | 0.0042568 | 2.3709 |
| MGDG 23:0_18:5 | 7.5934 | 2.9247 | 0.0043165 | 2.3649 |
| PA O-36:2 | 12.015 | 3.5867 | 0.0043295 | 2.3636 |
| SM 36:0;O2 | 19.01 | 4.2487 | 0.0043676 | 2.3598 |
| TG 16:0_18:0_22:2 | 0.019988 | -5.6447 | 0.0043983 | 2.3567 |
| HexCer 47:2;O5 | 0.58589 | -0.7713 | 0.0044019 | 2.3564 |
| SM 30:3;O3 | 13.411 | 3.7454 | 0.0044089 | 2.3557 |
| PC 28:0 | 4.9984 | 2.3215 | 0.0044117 | 2.3554 |
| Car 31:4;O | 2.6475 | 1.4046 | 0.004414 | 2.3552 |
| PE 44:2 | 0.49954 | -1.0013 | 0.0044143 | 2.3551 |
| SM 18:0;O2/23:1 | 9.6547 | 3.2712 | 0.0044151 | 2.3551 |
| PC 44:2 | 5.1718 | 2.3707 | 0.0044234 | 2.3542 |
| SM 43:2;O4 | 5.3262 | 2.4131 | 0.0044477 | 2.3519 |
| DG 28:0_18:1 | 0.1111 | -3.17 | 0.0044787 | 2.3488 |
| DG 18:1_20:2 | 0.46095 | -1.1173 | 0.0044938 | 2.3474 |
| CerP 44:6;O2 | 1.7794 | 0.8314 | 0.0045053 | 2.3463 |
| HexCer 43:0;O4 | 0.62423 | -0.67984 | 0.0045184 | 2.345 |
| CDP-DG 34:1 | 13.784 | 3.7849 | 0.0045263 | 2.3443 |
| Hex(3)-Cer 44:1;O2 | 0.59745 | -0.74312 | 0.0045399 | 2.343 |
| MG O-32:1 | 1.5373 | 0.6204 | 0.0045756 | 2.3395 |
| PA O-39:4 | 1.608 | 0.68524 | 0.0045846 | 2.3387 |
| TG 9:0_11:0_14:0 | 1.6977 | 0.76362 | 0.0045877 | 2.3384 |
| Hex(3)-HexNAc-KDN(2)-Cer 36:1;O2 | 3.223 | 1.6884 | 0.0046092 | 2.3364 |
| Cer 18:3;O2/35:7;O | 1.4408 | 0.52689 | 0.0046128 | 2.336 |
| PS O-18:3_22:5 | 4.3504 | 2.1211 | 0.004618 | 2.3355 |
| ST 29:0;O2;GlcA | 0.62588 | -0.67605 | 0.0046223 | 2.3351 |
| PC O-36:1 | 7.5046 | 2.9078 | 0.0046302 | 2.3344 |
| SM 46:1;O6 | 15.535 | 3.9575 | 0.0046324 | 2.3342 |
| ST 30:4;O2 | 0.30144 | -1.7301 | 0.0046383 | 2.3336 |
| DG 58:5 | 0.023089 | -5.4366 | 0.0046581 | 2.3318 |
| DG 44:2;O2 | 2.2596 | 1.1761 | 0.0046617 | 2.3315 |
| FAHFA 19:0/19:2 | 1.7552 | 0.81167 | 0.0046922 | 2.3286 |
| PC 40:0;O | 5.0157 | 2.3264 | 0.0046941 | 2.3284 |
| PC O-16:1_18:1 | 0.30375 | -1.7191 | 0.0047067 | 2.3273 |
| BMP 18:2;O | 66.005 | 6.0445 | 0.0047262 | 2.3255 |
| PE 28:2;O | 0.67849 | -0.55961 | 0.0047268 | 2.3254 |
| DG dO-40:0 | 0.38971 | -1.3595 | 0.0047335 | 2.3248 |
| FA 33:3;O2 | 0.38115 | -1.3916 | 0.004738 | 2.3244 |
| NAE 26:0;O2 | 2.7274 | 1.4475 | 0.004761 | 2.3223 |
| HexCer 38:0;O4 | 1.8492 | 0.88693 | 0.004792 | 2.3195 |
| LPC 20:4 | 0.58452 | -0.77468 | 0.0048024 | 2.3185 |
| PA O-58:6 | 10.822 | 3.4359 | 0.0048204 | 2.3169 |
| PC 40:0 | 7.0311 | 2.8138 | 0.004833 | 2.3158 |
| PE O-38:2 | 0.32596 | -1.6172 | 0.0048497 | 2.3143 |
| NAT 17:0 | 1.4526 | 0.5386 | 0.0048515 | 2.3141 |
| ACer 62:1;O3 | 0.11509 | -3.1191 | 0.004866 | 2.3128 |
| HexCer 45:6;O5 | 0.36857 | -1.44 | 0.0048717 | 2.3123 |
| NA 18:0 | 1.9759 | 0.98249 | 0.0048834 | 2.3113 |
| TG 71:0;O3 | 0.5087 | -0.97512 | 0.0049207 | 2.308 |
| PG O-33:7 | 1.5439 | 0.62656 | 0.0049413 | 2.3062 |
| Cer 43:5;O | 0.62338 | -0.68182 | 0.0049446 | 2.3059 |
| SM 42:4;O2 | 25.939 | 4.6971 | 0.0049897 | 2.3019 |
| Cer 15:2;2O/38:10 | 0.0049014 | -7.6726 | 0.0050154 | 2.2997 |
| FAHFA 48:3;O | 1.9536 | 0.96611 | 0.0050169 | 2.2996 |
| PI-Cer 35:0;O2 | 0.50119 | -0.99658 | 0.0050255 | 2.2988 |
| PE 54:2;O | 0.48323 | -1.0492 | 0.0050311 | 2.2983 |
| CL 72:7 | 0.46857 | -1.0937 | 0.0050448 | 2.2972 |
| ACer 64:2;O4 | 0.25071 | -1.9959 | 0.0051242 | 2.2904 |
| TG O-14:0_18:0_20:2 | 0.40673 | -1.2979 | 0.0051474 | 2.2884 |
| ST 28:7;O;HexNAc | 0.29121 | -1.7799 | 0.0051621 | 2.2872 |
| DG 42:6 | 0.53774 | -0.89501 | 0.0051743 | 2.2861 |
| Cer 43:6;O | 0.58869 | -0.76443 | 0.0051948 | 2.2844 |
| HexCer 49:2;O5 | 0.48454 | -1.0453 | 0.0051957 | 2.2844 |
| HexCer 39:6;O5 | 3.3163 | 1.7296 | 0.0052531 | 2.2796 |
| SM 18:1;O2/26:2 | 6.8155 | 2.7688 | 0.0052716 | 2.2781 |
| PG 41:4;O | 0.51099 | -0.96863 | 0.0052959 | 2.2761 |
| PC 33:2 | 5.6653 | 2.5021 | 0.0053053 | 2.2753 |
| SM 38:3;O3 | 4.1206 | 2.0429 | 0.0053085 | 2.275 |
| SM 38:0;O4 | 8.0702 | 3.0126 | 0.0053281 | 2.2734 |
| DG 51:4 | 0.32155 | -1.6369 | 0.0053764 | 2.2695 |
| TG 92:1 | 1.4891 | 0.57446 | 0.0053807 | 2.2692 |
| LPC 30:1 | 4.2032 | 2.0715 | 0.0054024 | 2.2674 |
| TG 12:0_16:0_16:1 | 0.10156 | -3.2996 | 0.0054123 | 2.2666 |
| Cer 40:1;O | 2.005 | 1.0036 | 0.0054288 | 2.2653 |
| PI-Cer 53:3;O2 | 0.54987 | -0.86285 | 0.0054289 | 2.2653 |
| PI-Cer 45:2;O2 | 1.5551 | 0.63702 | 0.0054653 | 2.2624 |
| LPC O-23:5 | 2.7096 | 1.4381 | 0.005476 | 2.2615 |
| DG 24:0;O2 | 3.2683 | 1.7085 | 0.0054802 | 2.2612 |
| FA 20:2 | 0.1234 | -3.0186 | 0.0054939 | 2.2601 |
| PA 48:0 | 5.7673 | 2.5279 | 0.0055086 | 2.259 |
| TG 56:6 | 0.30246 | -1.7252 | 0.0055578 | 2.2551 |
| SM 48:4;O3 | 4.9059 | 2.2945 | 0.0055768 | 2.2536 |
| MGDG 20:3_20:3 | 2.6344 | 1.3975 | 0.005596 | 2.2521 |
| ST 29:2;O2 | 0.30626 | -1.7071 | 0.0056237 | 2.25 |
| PG 22:6_22:6 | 0.47039 | -1.0881 | 0.0056342 | 2.2492 |
| FA 18:0;O5 | 1.8256 | 0.86838 | 0.0056376 | 2.2489 |
| PC 46:1 | 5.2536 | 2.3933 | 0.0056459 | 2.2483 |
| PC 44:0 | 5.8106 | 2.5387 | 0.0056489 | 2.248 |
| LPC 14:0 | 0.45254 | -1.1439 | 0.0056573 | 2.2474 |
| PE 18:1_20:2 | 0.5352 | -0.90185 | 0.0056931 | 2.2446 |
| SM 18:1;O2/25:1 | 7.0934 | 2.8265 | 0.0057032 | 2.2439 |
| HexCer 38:2;O2 | 0.58878 | -0.7642 | 0.0057273 | 2.242 |
| HexCer 35:2;O5 | 0.1964 | -2.3481 | 0.0057295 | 2.2419 |
| HexCer 18:0;O2/22:1 | 1.8858 | 0.9152 | 0.0057364 | 2.2414 |
| PC 44:1 | 5.1139 | 2.3544 | 0.0057425 | 2.2409 |
| NAT 26:3;O3 | 1.721 | 0.78324 | 0.0057654 | 2.2392 |
| PC 32:4 | 4.3948 | 2.1358 | 0.0057663 | 2.2391 |
| LPA O-27:7 | 2.5358 | 1.3424 | 0.0057677 | 2.239 |
| LPS 28:0;O | 0.61461 | -0.70225 | 0.0057706 | 2.2388 |
| TG 54:5 | 0.30552 | -1.7107 | 0.0057753 | 2.2384 |
| PC O-16:0_18:1 | 5.9752 | 2.579 | 0.0058009 | 2.2365 |
| PIP3 26:6 | 2.9023 | 1.5372 | 0.0058101 | 2.2358 |
| PC 34:1 | 5.7016 | 2.5114 | 0.0058356 | 2.2339 |
| HexCer 42:4;O2 | 0.60323 | -0.72922 | 0.0058359 | 2.2339 |
| DG 36:0;O2 | 4.8656 | 2.2826 | 0.0059254 | 2.2273 |
| TG 46:4;O | 0.38509 | -1.3767 | 0.0059667 | 2.2243 |
| Cer 37:6;O2 | 0.45255 | -1.1438 | 0.0060219 | 2.2203 |
| TG 49:12;O2 | 1.7746 | 0.8275 | 0.0060282 | 2.2198 |
| MG 19:0;O | 0.32088 | -1.6399 | 0.006049 | 2.2183 |
| PC 32:0 | 4.294 | 2.1023 | 0.0060573 | 2.2177 |
| LPC 17:0 | 0.41685 | -1.2624 | 0.0061076 | 2.2141 |
| SM 18:2;O2/26:2 | 15.907 | 3.9916 | 0.0061101 | 2.214 |
| DG O-50:4 | 0.16505 | -2.599 | 0.0061276 | 2.2127 |
| DG 60:6 | 0.039087 | -4.6772 | 0.0061448 | 2.2115 |
| SM 40:1;O4 | 4.6276 | 2.2103 | 0.0061475 | 2.2113 |
| PC 30:1 | 4.247 | 2.0865 | 0.006159 | 2.2105 |
| TG O-18:3_8:0_12:0 | 0.54939 | -0.86409 | 0.0061826 | 2.2088 |
| LPE 32:1 | 0.58641 | -0.77003 | 0.0061966 | 2.2078 |
| HexCer 37:0;O5 | 2.1397 | 1.0974 | 0.0062177 | 2.2064 |
| PE 42:7 | 0.42378 | -1.2386 | 0.006226 | 2.2058 |
| TG 15:0_16:0_16:1 | 0.24873 | -2.0074 | 0.0062281 | 2.2056 |
| DG 18:1_21:1 | 0.21215 | -2.2368 | 0.006247 | 2.2043 |
| PC 48:7 | 4.2717 | 2.0948 | 0.0062756 | 2.2023 |
| PA 40:0;O | 17.309 | 4.1135 | 0.00628 | 2.202 |
| PA 26:0 | 9.6941 | 3.2771 | 0.0062903 | 2.2013 |
| PI-Cer 43:2;O2 | 1.4468 | 0.53288 | 0.0062927 | 2.2012 |
| PC 28:0;O | 4.4995 | 2.1698 | 0.0063175 | 2.1995 |
| CL 76:9 | 0.50975 | -0.97213 | 0.0063198 | 2.1993 |
| CL 16:0_18:1_16:0_18:1 | 0.010815 | -6.5308 | 0.0063206 | 2.1992 |
| HexCer 40:4;O4 | 0.58837 | -0.7652 | 0.0063556 | 2.1968 |
| TG 48:8;O3 | 1.9552 | 0.96729 | 0.0064032 | 2.1936 |
| HexCer 37:3;O6 | 1.8388 | 0.8788 | 0.0064044 | 2.1935 |
| TG O-36:1 | 2.2542 | 1.1726 | 0.0064243 | 2.1922 |
| DG O-58:6 | 0.19291 | -2.374 | 0.006447 | 2.1906 |
| CE 32:3;O | 0.66698 | -0.58428 | 0.0064604 | 2.1897 |
| SM 44:5;O4 | 4.1843 | 2.065 | 0.0064673 | 2.1893 |
| Cer 21:3;O2/24:6;O | 0.52821 | -0.92082 | 0.0064779 | 2.1886 |
| PA O-56:5 | 14.804 | 3.8879 | 0.0064916 | 2.1876 |
| SM 17:0;O2/18:1 | 7.4946 | 2.9058 | 0.0065017 | 2.187 |
| PC O-38:1 | 6.5521 | 2.712 | 0.0065226 | 2.1856 |
| PC 28:2 | 4.2707 | 2.0945 | 0.0065348 | 2.1848 |
| DG 55:5 | 0.12105 | -3.0463 | 0.0065553 | 2.1834 |
| PC O-16:0_18:3 | 13.021 | 3.7028 | 0.0065648 | 2.1828 |
| PA 38:0;O | 16.15 | 4.0134 | 0.0066441 | 2.1776 |
| DG 24:0_26:2 | 0.022411 | -5.4797 | 0.0066814 | 2.1751 |
| FA 22:1 | 0.37887 | -1.4002 | 0.0066929 | 2.1744 |
| PE O-16:0_20:2 | 0.63825 | -0.64781 | 0.0067004 | 2.1739 |
| CE 20:1 | 0.1713 | -2.5454 | 0.0067117 | 2.1732 |
| LPC 18:2 | 0.55178 | -0.85784 | 0.0067161 | 2.1729 |
| Hex2Cer 42:2;O2 | 0.4506 | -1.1501 | 0.0067335 | 2.1718 |
| TG 14:0_16:0_16:1 | 0.039462 | -4.6634 | 0.0067403 | 2.1713 |
| PC O-20:3_16:1 | 6.7237 | 2.7493 | 0.0067516 | 2.1706 |
| PE O-38:4 | 1.5812 | 0.66099 | 0.0067583 | 2.1702 |
| HexCer 38:3;O3 | 0.53532 | -0.90153 | 0.0067615 | 2.17 |
| PC 44:6 | 4.0571 | 2.0205 | 0.0067671 | 2.1696 |
| PC 34:2 | 4.5283 | 2.179 | 0.006817 | 2.1664 |
| PA 43:9 | 12.538 | 3.6482 | 0.006859 | 2.1637 |
| PA O-37:1 | 12.538 | 3.6482 | 0.006859 | 2.1637 |
| MG O-16:0;O | 0.62395 | -0.68049 | 0.006867 | 2.1632 |
| PC 20:2 | 15.064 | 3.913 | 0.0068859 | 2.162 |
| TG 15:0_16:0_17:0 | 0.46504 | -1.1046 | 0.0068883 | 2.1619 |
| DG 16:0_18:3 | 4.4572 | 2.1561 | 0.0069275 | 2.1594 |
| TG 83:21;O3 | 1.6141 | 0.6907 | 0.0069413 | 2.1586 |
| HBMP 16:0_20:2_18:0 | 0.45238 | -1.1444 | 0.006978 | 2.1563 |
| PA 39:2 | 0.56666 | -0.81945 | 0.0070289 | 2.1531 |
| LPE 24:1 | 1.5625 | 0.64384 | 0.0070588 | 2.1513 |
| LPC O-34:3 | 0.27193 | -1.8787 | 0.0070635 | 2.151 |
| TG 16:0_18:1_20:1 | 0.017655 | -5.8238 | 0.0070674 | 2.1507 |
| PC O-18:0_19:1 | 5.4449 | 2.4449 | 0.0070688 | 2.1507 |
| SM 36:2;O2 | 23.52 | 4.5558 | 0.0071049 | 2.1484 |
| PA 37:8;O | 11.547 | 3.5294 | 0.0071249 | 2.1472 |
| PR 53:11;O2 | 0.59938 | -0.73845 | 0.0071274 | 2.1471 |
| ST 23:0;O3 | 0.59938 | -0.73845 | 0.0071274 | 2.1471 |
| ST 25:3;O | 0.59938 | -0.73845 | 0.0071274 | 2.1471 |
| ST 27:7;O | 0.59938 | -0.73845 | 0.0071274 | 2.1471 |
| ST 29:0;O7 | 0.59938 | -0.73845 | 0.0071274 | 2.1471 |
| PC 32:1;O | 3.8342 | 1.9389 | 0.0071481 | 2.1458 |
| Cer 38:1;O3 | 2.2689 | 1.182 | 0.0071611 | 2.145 |
| PE 52:3 | 2.0055 | 1.004 | 0.007204 | 2.1424 |
| FA 16:3;O4 | 1.5607 | 0.64219 | 0.0072057 | 2.1423 |
| DG 42:7;O | 0.68909 | -0.53724 | 0.0072129 | 2.1419 |
| PA 28:0 | 11.28 | 3.4956 | 0.0072158 | 2.1417 |
| FOH 16:0 | 1.4018 | 0.48732 | 0.0072184 | 2.1416 |
| MIPC 37:6;O4 | 2.2446 | 1.1665 | 0.0072503 | 2.1396 |
| PA 55:0 | 7.3186 | 2.8716 | 0.0072508 | 2.1396 |
| HexCer 42:6;O5 | 2.7993 | 1.4851 | 0.0072815 | 2.1378 |
| TG 12:0_14:0_16:0 | 0.30271 | -1.724 | 0.0072867 | 2.1375 |
| Cer 40:1;O3 | 2.7936 | 1.4821 | 0.0073286 | 2.135 |
| PG 14:0_18:1 | 0.18052 | -2.4697 | 0.0074021 | 2.1306 |
| TG 48:5;O2 | 0.52652 | -0.92543 | 0.0074138 | 2.13 |
| CerP 40:2;O5 | 1.602 | 0.67985 | 0.0074312 | 2.1289 |
| HexCer 36:1;O2 | 2.9079 | 1.54 | 0.007447 | 2.128 |
| CerP 51:3;O2 | 0.18133 | -2.4633 | 0.0074809 | 2.126 |
| SM 18:0;O2/22:1 | 5.8383 | 2.5455 | 0.0074957 | 2.1252 |
| PE 16:0_18:0 | 0.42505 | -1.2343 | 0.0075029 | 2.1248 |
| HexCer 31:0;O6 | 1.7513 | 0.80843 | 0.0075114 | 2.1243 |
| SM 18:1;O2/23:1 | 7.0934 | 2.8265 | 0.0075145 | 2.1241 |
| PE-Cer 28:2;O2 | 0.22306 | -2.1645 | 0.0075339 | 2.123 |
| PI-Cer 29:1;O3 | 2.0936 | 1.066 | 0.0075828 | 2.1202 |
| PG 56:3;O | 0.62798 | -0.67121 | 0.0075855 | 2.12 |
| NAE 20:0;O2 | 1.7053 | 0.77001 | 0.0075863 | 2.12 |
| PA 48:1 | 10.115 | 3.3384 | 0.0075902 | 2.1197 |
| TG 75:5 | 0.65268 | -0.61556 | 0.0075924 | 2.1196 |
| PA O-42:4 | 0.5495 | -0.86381 | 0.0075969 | 2.1194 |
| PA 49:1 | 10.088 | 3.3346 | 0.0076802 | 2.1146 |
| PC 17:0_18:0 | 0.71139 | -0.49129 | 0.0076834 | 2.1144 |
| DG 59:6 | 0.15624 | -2.6782 | 0.0077039 | 2.1133 |
| SLBPA 42:0 | 4.4233 | 2.1451 | 0.0077105 | 2.1129 |
| SCer 32:1;O | 2.0784 | 1.0555 | 0.0077165 | 2.1126 |
| PC 42:4 | 4.3482 | 2.1204 | 0.0077395 | 2.1113 |
| TG 51:1 | 0.081298 | -3.6206 | 0.0077724 | 2.1094 |
| PG 56:4;O | 0.58796 | -0.7662 | 0.0077963 | 2.1081 |
| LPC O-20:4 | 0.074975 | -3.7375 | 0.0078091 | 2.1074 |
| PI 14:0_16:0 | 0.48092 | -1.0561 | 0.007839 | 2.1057 |
| NAT 33:0;O4 | 0.67994 | -0.55653 | 0.0078479 | 2.1052 |
| DG 38:6;O2 | 3.4786 | 1.7985 | 0.0078496 | 2.1052 |
| PR 38:7;O2 | 0.66299 | -0.59295 | 0.0078632 | 2.1044 |
| SM 18:0;O2/26:2 | 5.2699 | 2.3978 | 0.0078637 | 2.1044 |
| LPC 32:0 | 0.32288 | -1.6309 | 0.0078675 | 2.1042 |
| PA 61:7 | 6.9229 | 2.7914 | 0.0078686 | 2.1041 |
| SM 43:5;O4 | 19.667 | 4.2977 | 0.0079256 | 2.101 |
| CerP 54:6;O3 | 0.5891 | -0.76341 | 0.007927 | 2.1009 |
| ST 28:1;O | 0.47913 | -1.0615 | 0.0079567 | 2.0993 |
| LPE O-20:2 | 1.4786 | 0.56427 | 0.007964 | 2.0989 |
| PE O-18:0_17:1 | 0.34723 | -1.526 | 0.0079806 | 2.098 |
| DG O-18:0_22:4 | 0.26637 | -1.9085 | 0.0079914 | 2.0974 |
| SHexCer 28:1;O2 | 2.7495 | 1.4592 | 0.0080226 | 2.0957 |
| PA O-58:10 | 9.3101 | 3.2188 | 0.0080245 | 2.0956 |
| HBMP 60:2; HBMP 20:0-22:1-18:1; [M+NH4]<sup>+</sup> | 7.3189 | 2.8716 | 0.0080568 | 2.0938 |
| PG 34:3 | 1.5939 | 0.67258 | 0.0080642 | 2.0934 |
| PC 38:7 | 3.7435 | 1.9044 | 0.0080953 | 2.0918 |
| PA 45:9 | 10.8 | 3.433 | 0.0081012 | 2.0914 |
| TG 58:2 | 0.048455 | -4.3672 | 0.0081025 | 2.0914 |
| PC 16:0_17:0 | 0.65953 | -0.6005 | 0.0081053 | 2.0912 |
| FAHFA 20:1/20:0 | 1.4372 | 0.52325 | 0.0081382 | 2.0895 |
| ST 19:0;O3;Hex | 1.4372 | 0.52325 | 0.0081382 | 2.0895 |
| ST 28:1;O;S | 1.4372 | 0.52325 | 0.0081382 | 2.0895 |
| ST 44:5;O2 | 1.4372 | 0.52325 | 0.0081382 | 2.0895 |
| Cer 19:3;O2/36:7;O | 0.46505 | -1.1046 | 0.0081786 | 2.0873 |
| PG O-54:0;O | 0.51539 | -0.95625 | 0.0082146 | 2.0854 |
| Cer 18:3;O2/32:7;O | 2.9487 | 1.5601 | 0.008234 | 2.0844 |
| Car 12:2;O2 | 2.2486 | 1.169 | 0.0082409 | 2.084 |
| CoA 24:6 | 8.9255 | 3.1579 | 0.008281 | 2.0819 |
| PC 68:7;O | 24.333 | 4.6049 | 0.0082891 | 2.0815 |
| PC O-14:0_4:0 | 0.45445 | -1.1378 | 0.0083172 | 2.08 |
| HexCer 35:4;O6 | 3.4342 | 1.78 | 0.0083326 | 2.0792 |
| PR 42:1;O8;N3 | 0.56081 | -0.83443 | 0.0084013 | 2.0757 |
| PE O-42:4 | 0.48919 | -1.0315 | 0.0084209 | 2.0746 |
| TG 50:6;O | 0.36953 | -1.4362 | 0.0084492 | 2.0732 |
| PA 31:8;O | 8.6961 | 3.1204 | 0.0084599 | 2.0726 |
| TG 45:2 | 0.0062555 | -7.3207 | 0.0084624 | 2.0725 |
| TG 57:2 | 0.043132 | -4.5351 | 0.0084705 | 2.0721 |
| TG 46:2 | 0.13325 | -2.9078 | 0.0085325 | 2.0689 |
| PC 30:2;O | 5.4278 | 2.4404 | 0.0085415 | 2.0685 |
| PC 38:0;O | 3.5682 | 1.8352 | 0.0085436 | 2.0684 |
| PA O-35:1 | 8.3939 | 3.0693 | 0.0085439 | 2.0683 |
| Cer 35:3;O3 | 1.5498 | 0.63211 | 0.0085712 | 2.067 |
| SM 42:1;O3 | 5.5735 | 2.4786 | 0.0085929 | 2.0659 |
| PA O-39:1 | 37.27 | 5.2199 | 0.0086002 | 2.0655 |
| Cer 19:0;O2/24:2;O | 0.21332 | -2.2289 | 0.0086011 | 2.0654 |
| PA 56:0 | 8.6565 | 3.1138 | 0.0086131 | 2.0648 |
| PA 41:4 | 16.058 | 4.0052 | 0.0086162 | 2.0647 |
| PE 40:0;O | 0.33852 | -1.5627 | 0.0086541 | 2.0628 |
| Car 19:2;O2 | 1.7622 | 0.81736 | 0.0086816 | 2.0614 |
| HexCer 45:3;O4 | 0.41558 | -1.2668 | 0.008689 | 2.061 |
| HexCer 19:0;O2/24:2 | 2.5162 | 1.3313 | 0.0086918 | 2.0609 |
| PA O-58:0 | 8.4058 | 3.0714 | 0.0087053 | 2.0602 |
| PI O-63:1 | 0.51336 | -0.96195 | 0.008709 | 2.06 |
| Cer 39:4;O | 0.61389 | -0.70396 | 0.0087311 | 2.0589 |
| LPC 20:2;O | 1.9465 | 0.96088 | 0.0087445 | 2.0583 |
| PC 18:0_20:4 | 5.4616 | 2.4493 | 0.0087505 | 2.058 |
| PC O-16:1_20:4 | 0.55274 | -0.85532 | 0.0087637 | 2.0573 |
| ST 22:0;O7 | 0.55063 | -0.86086 | 0.0087816 | 2.0564 |
| LPA 26:4 | 0.39166 | -1.3523 | 0.0087966 | 2.0557 |
| TG 42:1 | 0.405 | -1.304 | 0.0088746 | 2.0519 |
| PA O-58:5 | 18.913 | 4.2413 | 0.0088786 | 2.0517 |
| DG 18:0_16:1 | 4.3576 | 2.1235 | 0.0088899 | 2.0511 |
| CL 16:0_18:2_16:0_22:6 | 0.014132 | -6.1449 | 0.0088916 | 2.051 |
| LPA 20:1 | 0.48986 | -1.0296 | 0.00895 | 2.0482 |
| TG 14:2_14:3_17:3 | 0.2318 | -2.109 | 0.0089575 | 2.0478 |
| PA 65:8;O | 11.993 | 3.5841 | 0.008983 | 2.0466 |
| SM 44:4;O5 | 3.7934 | 1.9235 | 0.0089927 | 2.0461 |
| CerP 34:2;O5 | 1.5782 | 0.65832 | 0.0090044 | 2.0455 |
| TG 41:1;O | 0.046392 | -4.43 | 0.0090122 | 2.0452 |
| Hex2Cer 29:0;O3 | 1.8481 | 0.88605 | 0.0090435 | 2.0437 |
| PE-Cer 33:2;O3 | 3.159 | 1.6595 | 0.0090633 | 2.0427 |
| ST 28:3;O3;F2 | 0.34935 | -1.5172 | 0.0090757 | 2.0421 |
| DG 41:6 | 1.4152 | 0.50105 | 0.0090861 | 2.0416 |
| CerP 41:2;O2 | 0.093264 | -3.4225 | 0.0091198 | 2.04 |
| DG 29:0;O2 | 2.8873 | 1.5297 | 0.0091226 | 2.0399 |
| PC 14:0_16:0 | 4.7054 | 2.2343 | 0.0091304 | 2.0395 |
| ST 25:0;O | 0.42509 | -1.2341 | 0.009136 | 2.0392 |
| TG 16:1_18:0_18:1 | 0.031037 | -5.0099 | 0.0091368 | 2.0392 |
| ST 20:2;O2;S | 0.67191 | -0.57366 | 0.0091412 | 2.039 |
| TG 82:1;O | 2.0523 | 1.0372 | 0.0091962 | 2.0364 |
| HexCer 38:4;O2 | 0.60582 | -0.72304 | 0.0092478 | 2.034 |
| TG 18:1_19:0_21:4 | 0.22754 | -2.1358 | 0.0092561 | 2.0336 |
| CL 73:5 | 0.47003 | -1.0892 | 0.0093647 | 2.0285 |
| Car 30:1;O2 | 1.6258 | 0.70117 | 0.0093823 | 2.0277 |
| PC 18:0_24:0 | 7.3008 | 2.8681 | 0.0093918 | 2.0273 |
| Cer 34:2;O2 | 0.41973 | -1.2525 | 0.0093959 | 2.0271 |
| SM 18:1;O2/24:2 | 8.2531 | 3.0449 | 0.0094037 | 2.0267 |
| HexCer 37:2;O6 | 2.5407 | 1.3452 | 0.0094772 | 2.0233 |
| PC O-40:1 | 4.423 | 2.145 | 0.0094845 | 2.023 |
| HexCer 32:5;O2 | 2.0521 | 1.0371 | 0.0095303 | 2.0209 |
| TG 77:1;O3 | 0.50455 | -0.98693 | 0.009534 | 2.0207 |
| PE-Cer 52:6;O2 | 0.60273 | -0.73041 | 0.0095582 | 2.0196 |
| FA 19:2;O4 | 2.7569 | 1.463 | 0.0097201 | 2.0123 |
| ST 27:3;O (Cholest-4,6-Dien-3-One) | 0.21826 | -2.1959 | 0.0097661 | 2.0103 |
| SPB 28:1;O4;N | 1.595 | 0.67359 | 0.0097922 | 2.0091 |
| HexCer 45:0;O4 | 0.26522 | -1.9147 | 0.0098487 | 2.0066 |
| TG 97:11;O3 | 0.62537 | -0.67722 | 0.0098795 | 2.0053 |
| PI-Cer 37:1;O2 | 0.62175 | -0.6856 | 0.0099258 | 2.0032 |
| PC 44:3 | 3.4969 | 1.8061 | 0.0099617 | 2.0017 |
| TG 55:7;O3 | 1.9291 | 0.94791 | 0.0099643 | 2.0016 |
| PC O-42:0 | 5.9834 | 2.581 | 0.0099994 | 2 |
| PC 32:2;O | 6.8137 | 2.7684 | 0.010102 | 1.9956 |
| TG 93:14;O | 1.6619 | 0.73282 | 0.010122 | 1.9947 |
| PA O-39:3 | 6.8487 | 2.7758 | 0.010138 | 1.9941 |
| PA 37:7;O | 6.7135 | 2.7471 | 0.010145 | 1.9937 |
| SM 42:5;O4 | 11.618 | 3.5383 | 0.010157 | 1.9932 |
| SM 34:2;O5 | 4.2271 | 2.0797 | 0.01016 | 1.9931 |
| PE 40:1 | 0.626 | -0.67577 | 0.010191 | 1.9918 |
| DG 56:2 | 2.581 | 1.3679 | 0.010202 | 1.9913 |
| DG 40:5;O | 0.62784 | -0.67154 | 0.010232 | 1.99 |
| LPA O-30:4 | 0.49189 | -1.0236 | 0.01027 | 1.9884 |
| TG 17:1_18:0_18:1 | 0.056946 | -4.1343 | 0.010279 | 1.9881 |
| HexCer 48:3;O3 | 2.1825 | 1.126 | 0.010331 | 1.9859 |
| TG 16:0_16:5_17:3 | 0.21105 | -2.2443 | 0.010379 | 1.9838 |
| Cer 43:2;O3 | 0.35443 | -1.4964 | 0.010379 | 1.9838 |
| SHexCer 43:4;O3 | 0.25705 | -1.9599 | 0.010485 | 1.9794 |
| HexCer 18:0;O2/26:2 | 1.6831 | 0.75109 | 0.010512 | 1.9783 |
| PC O-14:0_18:3 | 22.263 | 4.4766 | 0.010589 | 1.9752 |
| PE O-20:3_27:0 | 0.15314 | -2.7071 | 0.010598 | 1.9748 |
| PA O-41:6 | 10.048 | 3.3288 | 0.010621 | 1.9738 |
| DG 18:1_36:2 | 0.065045 | -3.9424 | 0.010622 | 1.9738 |
| PE 36:0 | 0.54051 | -0.8876 | 0.010634 | 1.9733 |
| PE 5:0_18:5 | 2.3967 | 1.2611 | 0.010687 | 1.9711 |
| TG 56:5 | 0.45585 | -1.1334 | 0.01071 | 1.9702 |
| TG 14:0_16:1_18:1 | 0.067925 | -3.8799 | 0.010722 | 1.9697 |
| MG 33:0;O | 2.1149 | 1.0806 | 0.010741 | 1.9689 |
| PIP 41:3 | 2.1277 | 1.0893 | 0.010761 | 1.9681 |
| TG 97:7;O3 | 2.008 | 1.0058 | 0.01077 | 1.9678 |
| SM 18:0;O2/22:2 | 8.2673 | 3.0474 | 0.010789 | 1.967 |
| PC 14:0_18:1 | 6.9006 | 2.7867 | 0.010805 | 1.9664 |
| SHexCer 30:1;O3 | 2.2848 | 1.1921 | 0.010825 | 1.9656 |
| PIP 54:4 | 0.70675 | -0.50072 | 0.010843 | 1.9649 |
| DG 50:5 | 0.23553 | -2.086 | 0.010855 | 1.9644 |
| PC 18:0_22:0 | 7.4355 | 2.8944 | 0.010855 | 1.9644 |
| CerP 36:1;O4 | 0.41227 | -1.2783 | 0.010933 | 1.9613 |
| SM 45:5;O2 | 7.6767 | 2.9405 | 0.010938 | 1.9611 |
| PC 18:1 | 22.075 | 4.4643 | 0.010946 | 1.9608 |
| TG 91:14 | 2.1827 | 1.1261 | 0.010952 | 1.9605 |
| PI O-61:3 | 2.0834 | 1.0589 | 0.010966 | 1.96 |
| PK 33:8;O18 | 0.57648 | -0.79466 | 0.010967 | 1.9599 |
| TG 16:0_19:3_19:3 | 0.2339 | -2.096 | 0.010996 | 1.9588 |
| PG 22:0_22:2 | 1.4341 | 0.52012 | 0.011003 | 1.9585 |
| PA O-36:1 | 17.742 | 4.1491 | 0.011067 | 1.956 |
| MG 16:1 | 0.024164 | -5.371 | 0.011068 | 1.9559 |
| PC 20:0_16:2 | 5.5565 | 2.4742 | 0.011072 | 1.9558 |
| TG 55:2 | 0.3062 | -1.7075 | 0.011088 | 1.9551 |
| CL 74:5 | 0.56503 | -0.82361 | 0.0111 | 1.9547 |
| PIP3 28:8 | 2.4942 | 1.3186 | 0.01113 | 1.9535 |
| HC 23:1 | 3.2586 | 1.7042 | 0.011137 | 1.9532 |
| Cer 41:0;O3 | 0.6885 | -0.53847 | 0.011158 | 1.9524 |
| MG O-32:0 | 0.58162 | -0.78185 | 0.01119 | 1.9512 |
| PC 32:1 | 3.9938 | 1.9978 | 0.011209 | 1.9504 |
| Car 16:2;O4 | 1.4881 | 0.5735 | 0.011225 | 1.9498 |
| PE 18:0_26:1 | 0.62265 | -0.6835 | 0.011244 | 1.9491 |
| PC O-18:4_15:1 | 1.415 | 0.50079 | 0.011255 | 1.9487 |
| ST 21:3;O3;G | 0.57943 | -0.78729 | 0.011326 | 1.9459 |
| HexCer 31:2;O4 | 1.4933 | 0.57851 | 0.011334 | 1.9456 |
| HexCer 42:1;O4 | 0.64145 | -0.6406 | 0.011366 | 1.9444 |
| Hex2Cer 32:1;O3 | 0.56754 | -0.81722 | 0.01138 | 1.9439 |
| ST 28:4;O3 | 0.58032 | -0.78507 | 0.011397 | 1.9432 |
| PC 20:1_16:2 | 4.2865 | 2.0998 | 0.011428 | 1.942 |
| PC O-38:0;O | 3.2272 | 1.6903 | 0.01146 | 1.9408 |
| LPE O-20:1 | 0.60484 | -0.72538 | 0.011467 | 1.9405 |
| ST 29:3;O4 | 2.7874 | 1.4789 | 0.011516 | 1.9387 |
| PE O-18:2_24:1 | 0.52902 | -0.9186 | 0.011521 | 1.9385 |
| PA 45:0 | 5.5776 | 2.4796 | 0.011523 | 1.9384 |
| PS 17:0_18:1 | 2.4054 | 1.2663 | 0.011555 | 1.9372 |
| PI 52:0;O | 0.66642 | -0.5855 | 0.011577 | 1.9364 |
| HexCer 18:0;O2/20:1 | 1.6624 | 0.73327 | 0.01162 | 1.9348 |
| PA 45:6 | 4.9939 | 2.3202 | 0.011647 | 1.9338 |
| HexCer 42:0;O6 | 0.56351 | -0.82748 | 0.011711 | 1.9314 |
| SM 33:1;O6 | 13.694 | 3.7755 | 0.011738 | 1.9304 |
| PC 34:4 | 2.6018 | 1.3795 | 0.011757 | 1.9297 |
| PE O-18:0_22:3 | 0.49536 | -1.0134 | 0.011777 | 1.929 |
| PA O-38:0;O | 10.103 | 3.3367 | 0.011801 | 1.9281 |
| DG 35:3 | 3.3562 | 1.7468 | 0.011819 | 1.9274 |
| FA 16:1 | 0.075988 | -3.7181 | 0.011859 | 1.926 |
| Cer 40:0;O | 0.47169 | -1.0841 | 0.011864 | 1.9258 |
| PA 42:0 | 12.276 | 3.6178 | 0.011928 | 1.9234 |
| TG 51:3 | 1.7565 | 0.8127 | 0.011933 | 1.9232 |
| PC 47:2 | 17.37 | 4.1186 | 0.011979 | 1.9216 |
| CerP 29:6;O2 | 2.6928 | 1.4291 | 0.012 | 1.9208 |
| PA O-44:0;O | 17.011 | 4.0884 | 0.012053 | 1.9189 |
| PC 31:1 | 9.2604 | 3.2111 | 0.012108 | 1.9169 |
| PA O-50:5 | 0.49238 | -1.0222 | 0.012119 | 1.9165 |
| PC 44:4 | 3.0314 | 1.6 | 0.012158 | 1.9151 |
| LPC 26:4 | 1.544 | 0.62671 | 0.012168 | 1.9148 |
| PA O-45:0 | 11.111 | 3.4739 | 0.01235 | 1.9083 |
| HexCer 40:5;O4 | 1.8709 | 0.90373 | 0.012407 | 1.9063 |
| PC 14:0_16:3 | 5.7667 | 2.5277 | 0.012419 | 1.9059 |
| PI 18:0_20:2 | 0.36323 | -1.461 | 0.012454 | 1.9047 |
| CerP 44:4;O3 | 0.028807 | -5.1174 | 0.012467 | 1.9042 |
| PE 14:0_18:2 | 0.36269 | -1.4632 | 0.012497 | 1.9032 |
| PI 18:0_22:4 | 0.64111 | -0.64137 | 0.012518 | 1.9025 |
| TG 62:2 | 0.29435 | -1.7644 | 0.012595 | 1.8998 |
| LPA 13:2;O | 1.9533 | 0.96593 | 0.012621 | 1.8989 |
| DG O-30:0 | 3.7236 | 1.8967 | 0.012715 | 1.8957 |
| DG 28:1;O | 2.4098 | 1.2689 | 0.012719 | 1.8956 |
| PIP3 35:5 | 0.45527 | -1.1352 | 0.012757 | 1.8942 |
| PC O-18:1_24:4 | 4.1779 | 2.0628 | 0.012899 | 1.8894 |
| TG 14:0_15:1_16:0 | 0.4293 | -1.22 | 0.012949 | 1.8878 |
| SHexCer 36:0;O2 | 0.66355 | -0.59173 | 0.012982 | 1.8867 |
| PC 18:0_21:2 | 3.5782 | 1.8392 | 0.013085 | 1.8832 |
| PG 54:6;O | 0.61811 | -0.69407 | 0.013098 | 1.8828 |
| HexCer 18:1;O2/22:0 | 1.8551 | 0.89149 | 0.013126 | 1.8819 |
| PC O-42:2 | 5.1791 | 2.3727 | 0.013169 | 1.8805 |
| LPC 27:3 | 1.783 | 0.83434 | 0.013212 | 1.879 |
| PA O-37:0 | 16.437 | 4.0389 | 0.013328 | 1.8752 |
| SM 15:0;O2/16:1 | 19.438 | 4.2808 | 0.013329 | 1.8752 |
| NAT 18:2 | 3.5561 | 1.8303 | 0.013345 | 1.8747 |
| TG 16:0_18:1_22:4 | 0.29111 | -1.7804 | 0.013348 | 1.8746 |
| PC 33:5 | 3.0271 | 1.5979 | 0.01335 | 1.8745 |
| Car 32:6;O2 | 1.5831 | 0.66276 | 0.013385 | 1.8734 |
| PC 36:0 | 7.8065 | 2.9647 | 0.013386 | 1.8733 |
| HexCer 41:0;O4 | 0.46294 | -1.1111 | 0.013389 | 1.8733 |
| Car 30:0;O | 0.42027 | -1.2506 | 0.013403 | 1.8728 |
| PG 28:2 | 0.55823 | -0.84106 | 0.013512 | 1.8693 |
| FA 20:0;O3 | 1.9923 | 0.99441 | 0.013551 | 1.868 |
| MG 27:2;O | 3.1116 | 1.6376 | 0.013641 | 1.8652 |
| PC O-36:5 | 2.9719 | 1.5714 | 0.013722 | 1.8626 |
| HexCer 36:4;O2 | 0.63794 | -0.64851 | 0.013741 | 1.862 |
| TG 16:0_18:1_22:6 | 0.17869 | -2.4845 | 0.013813 | 1.8597 |
| HC 25:1 | 3.1746 | 1.6666 | 0.013816 | 1.8596 |
| DG 17:0_18:3 | 5.1485 | 2.3642 | 0.013823 | 1.8594 |
| PA 25:0_18:4 | 0.66575 | -0.58694 | 0.013902 | 1.8569 |
| PC O-22:3_24:0 | 0.43298 | -1.2076 | 0.013974 | 1.8547 |
| FOH 16:3 | 3.0206 | 1.5949 | 0.014075 | 1.8516 |
| SM 38:3;O2 | 3.012 | 1.5907 | 0.014091 | 1.8511 |
| SM 43:5;O2 | 9.2163 | 3.2042 | 0.014105 | 1.8506 |
| DG 54:5 | 0.31711 | -1.6569 | 0.014109 | 1.8505 |
| DG O-42:6 | 0.57239 | -0.80494 | 0.014125 | 1.85 |
| HexCer 23:0;O2/18:2 | 1.9676 | 0.97644 | 0.014137 | 1.8497 |
| PC 33:3;O | 3.003 | 1.5864 | 0.014171 | 1.8486 |
| ST 25:4;O4 | 0.3845 | -1.3789 | 0.014309 | 1.8444 |
| HexCer 18:2;O2/26:2 | 3.9052 | 1.9654 | 0.014347 | 1.8432 |
| PI 18:1_22:4 | 0.18361 | -2.4453 | 0.014358 | 1.8429 |
| TG 16:1_18:1_22:1 | 0.033507 | -4.8994 | 0.014369 | 1.8426 |
| Cer 38:0;O | 0.50757 | -0.97833 | 0.014414 | 1.8412 |
| SM 34:0;O4 | 28.437 | 4.8297 | 0.014416 | 1.8412 |
| FA 20:4 | 0.015909 | -5.974 | 0.014424 | 1.8409 |
| LPC 14:2;O | 1.7889 | 0.83911 | 0.014541 | 1.8374 |
| SM 19:3;O2/24:2 | 14.098 | 3.8175 | 0.014564 | 1.8367 |
| PI-Cer 45:2;O3 | 1.4687 | 0.55458 | 0.01464 | 1.8345 |
| DG 39:4;O | 3.6655 | 1.874 | 0.014704 | 1.8326 |
| PA O-49:1 | 33.132 | 5.0502 | 0.014749 | 1.8312 |
| PIP 54:3 | 0.62975 | -0.66715 | 0.014825 | 1.829 |
| Hex(3)-HexNAc-NeuAc-Cer 42:1;O2 | 3.0735 | 1.6199 | 0.014871 | 1.8277 |
| TG 33:1;O3 | 0.65333 | -0.61413 | 0.014939 | 1.8257 |
| PG 54:4 | 0.41094 | -1.283 | 0.01497 | 1.8248 |
| DG 46:3 | 0.38648 | -1.3715 | 0.014996 | 1.824 |
| PC O-40:3 | 3.9095 | 1.967 | 0.015004 | 1.8238 |
| TG 53:0 | 0.59493 | -0.74921 | 0.015016 | 1.8235 |
| CE 27:7 | 1.9664 | 0.97554 | 0.015045 | 1.8226 |
| SM 39:0;O2 | 15.696 | 3.9724 | 0.015046 | 1.8226 |
| DG 35:1 | 3.8154 | 1.9318 | 0.015064 | 1.8221 |
| PR 35:0;O4 | 0.62702 | -0.67342 | 0.015118 | 1.8205 |
| PC 43:1;O | 3.474 | 1.7966 | 0.015156 | 1.8194 |
| TG 48:4;O | 0.40715 | -1.2964 | 0.015169 | 1.819 |
| ST 23:3;O4;G | 0.63848 | -0.64728 | 0.015183 | 1.8187 |
| PI 66:0 | 1.7249 | 0.78649 | 0.015213 | 1.8178 |
| PC 38:1 | 2.417 | 1.2732 | 0.015232 | 1.8172 |
| LPG 31:1 | 0.6921 | -0.53095 | 0.015264 | 1.8163 |
| DG 32:9 | 2.037 | 1.0264 | 0.015276 | 1.816 |
| Cer 22:2;O2/24:6 | 1.7352 | 0.7951 | 0.015374 | 1.8132 |
| PA 46:2 | 5.5278 | 2.4667 | 0.015376 | 1.8132 |
| SM 18:0;O2/24:1 | 5.9142 | 2.5642 | 0.015406 | 1.8123 |
| HexCer 40:3;O4 | 1.7754 | 0.82814 | 0.015409 | 1.8122 |
| DG 40:1;O | 2.8229 | 1.4972 | 0.015431 | 1.8116 |
| PC 30:3;O2 | 9.5715 | 3.2587 | 0.015469 | 1.8105 |
| ST 25:1;O2 | 0.40428 | -1.3066 | 0.015494 | 1.8098 |
| PE O-28:4_28:7 | 1.5292 | 0.6128 | 0.015518 | 1.8092 |
| LPI O-16:0 | 1.5726 | 0.65316 | 0.015552 | 1.8082 |
| Hex(5)-HexNAc(2)-Fuc-Cer 36:1;O2 | 0.51944 | -0.94496 | 0.015597 | 1.8069 |
| PI 16:0_18:0 | 0.44321 | -1.1739 | 0.015608 | 1.8067 |
| PC O-18:0_14:0 | 0.54966 | -0.86338 | 0.015619 | 1.8063 |
| ST 28:2;O4 | 0.61458 | -0.70234 | 0.015666 | 1.805 |
| TG 16:0_17:1_18:1 | 1.6449 | 0.71797 | 0.015674 | 1.8048 |
| PE O-18:1_16:0 | 1.948 | 0.962 | 0.015709 | 1.8038 |
| PI O-65:2 | 0.49393 | -1.0176 | 0.015727 | 1.8033 |
| Car 22:4;O4 | 0.42032 | -1.2504 | 0.015765 | 1.8023 |
| PE 23:6 | 2.0525 | 1.0374 | 0.01578 | 1.8019 |
| PIP3 44:9 | 1.4075 | 0.49318 | 0.015878 | 1.7992 |
| MG O-18:0 | 0.41042 | -1.2848 | 0.015942 | 1.7974 |
| DG 44:7;O | 1.6832 | 0.75117 | 0.016008 | 1.7957 |
| CL 74:4 | 0.63306 | -0.65959 | 0.016016 | 1.7954 |
| PA 44:0;O | 9.1073 | 3.187 | 0.016082 | 1.7937 |
| MG 23:2;O | 0.25627 | -1.9642 | 0.016195 | 1.7906 |
| FA 17:3;O3 | 2.9624 | 1.5668 | 0.016265 | 1.7887 |
| PE-Cer 40:0;O6 | 2.6664 | 1.4149 | 0.016285 | 1.7882 |
| DG O-35:0;O2 | 4.6538 | 2.2184 | 0.016311 | 1.7875 |
| Car 30:2;O2 | 1.5475 | 0.62996 | 0.016346 | 1.7866 |
| PC O-18:0_20:5 | 13.129 | 3.7146 | 0.0164 | 1.7851 |
| SM 32:0;O4 | 18.162 | 4.1829 | 0.01667 | 1.7781 |
| TG 57:10 | 0.59681 | -0.74466 | 0.016825 | 1.774 |
| PE O-18:0_20:2 | 0.5613 | -0.83316 | 0.01687 | 1.7729 |
| CerP 14:0;O2/12:0 | 0.21864 | -2.1934 | 0.016895 | 1.7722 |
| CL 16:0_18:2_16:0_20:5 | 0.022223 | -5.4918 | 0.016953 | 1.7707 |
| PC 15:0_18:0 | 4.314 | 2.109 | 0.016959 | 1.7706 |
| DG 52:4 | 0.40845 | -1.2918 | 0.017021 | 1.769 |
| DG O-56:6 | 0.30217 | -1.7266 | 0.01704 | 1.7685 |
| TG 14:0_18:1_19:1 | 0.10614 | -3.236 | 0.017136 | 1.7661 |
| PA 43:6 | 3.9185 | 1.9703 | 0.0172 | 1.7645 |
| CL 74:9 | 0.30524 | -1.712 | 0.017222 | 1.7639 |
| PC 20:2_16:3 | 3.1378 | 1.6498 | 0.017417 | 1.759 |
| SM 48:5;O2 | 2.8719 | 1.522 | 0.01751 | 1.7567 |
| NAE 33:5;O | 4.5113 | 2.1736 | 0.017729 | 1.7513 |
| AcylGlcADG<br>22:5_22:5_20:5 | 2.1033 | 1.0726 | 0.01782 | 1.7491 |
| ST 28:0;O8;T | 0.51519 | -0.95682 | 0.017826 | 1.7489 |
| Car 22:2;O4 | 0.49547 | -1.0131 | 0.017843 | 1.7485 |
| DG 60:7 | 0.29539 | -1.7593 | 0.017939 | 1.7462 |
| ST 24:3;O3 | 0.66148 | -0.59624 | 0.017992 | 1.7449 |
| PA 45:4 | 10.806 | 3.4337 | 0.018017 | 1.7443 |
| PA O-39:0 | 20.949 | 4.3888 | 0.018046 | 1.7436 |
| PA O-44:5 | 4.1451 | 2.0514 | 0.018178 | 1.7405 |
| NAT 34:2;O4 | 1.6707 | 0.74045 | 0.018188 | 1.7402 |
| PG O-43:0 | 1.5826 | 0.6623 | 0.01827 | 1.7383 |
| DG 59:2 | 1.6869 | 0.75441 | 0.018329 | 1.7369 |
| PG O-44:5 | 0.69708 | -0.5206 | 0.018352 | 1.7363 |
| PE O-16:0_18:1 | 0.33354 | -1.5841 | 0.018427 | 1.7345 |
| PE O-18:0_17:0 | 0.5576 | -0.84271 | 0.018478 | 1.7333 |
| ST 23:3;O3 | 0.19041 | -2.3929 | 0.018511 | 1.7326 |
| HexCer 18:1;O2/24:1 | 1.6807 | 0.74908 | 0.018524 | 1.7323 |
| TG 96:8;O | 0.6687 | -0.58057 | 0.018532 | 1.7321 |
| ST 30:2;O | 0.54681 | -0.8709 | 0.018597 | 1.7306 |
| DG 35:4 | 2.8303 | 1.501 | 0.018605 | 1.7304 |
| LPC 26:1 | 0.53204 | -0.9104 | 0.018634 | 1.7297 |
| PA 42:5 | 3.2744 | 1.7112 | 0.018731 | 1.7274 |
| Car 32:5;O | 1.8563 | 0.89245 | 0.018939 | 1.7226 |
| LPC 30:0 | 0.48439 | -1.0458 | 0.019054 | 1.72 |
| LPC 33:2 | 0.16139 | -2.6314 | 0.019066 | 1.7197 |
| TG 14:0_14:3_19:5 | 0.28392 | -1.8164 | 0.019079 | 1.7194 |
| DG 39:2 | 0.56244 | -0.83023 | 0.019114 | 1.7186 |
| PE O-16:2_23:0 | 0.66752 | -0.58311 | 0.019136 | 1.7182 |
| DG 52:3 | 0.2728 | -1.8741 | 0.019156 | 1.7177 |
| DG 60:10 | 0.24563 | -2.0254 | 0.019235 | 1.7159 |
| ST 21:1;O7 | 1.655 | 0.72681 | 0.019337 | 1.7136 |
| FA 28:0;O3 | 5.0982 | 2.35 | 0.019377 | 1.7127 |
| SM 34:1;O2(FA 24:1) | 2.6623 | 1.4127 | 0.019593 | 1.7079 |
| PG 23:1;O | 1.7264 | 0.78778 | 0.019721 | 1.7051 |
| PA O-40:0;O | 10.297 | 3.3642 | 0.019729 | 1.7049 |
| SM 53:6;O2 | 8.5352 | 3.0934 | 0.019786 | 1.7036 |
| PEtOH 18:1_20:5 | 3.6055 | 1.8502 | 0.019942 | 1.7002 |
| Cer 42:1;O3 | 1.4781 | 0.56373 | 0.019987 | 1.6992 |
| PC 18:1_20:3 | 0.48018 | -1.0584 | 0.02012 | 1.6964 |
| DG O-32:0 | 2.6515 | 1.4068 | 0.020171 | 1.6953 |
| Car 30:2;O3 | 1.898 | 0.92448 | 0.020208 | 1.6945 |
| PA 42:0;O | 10.923 | 3.4493 | 0.020247 | 1.6936 |
| PA 41:8 | 3.8529 | 1.9459 | 0.020276 | 1.693 |
| PE 52:5;O | 0.57329 | -0.80266 | 0.020333 | 1.6918 |
| ST 27:1;O | 1.9732 | 0.98051 | 0.020372 | 1.691 |
| DG O-48:3 | 0.42321 | -1.2405 | 0.020493 | 1.6884 |
| PG 32:0;O | 0.28648 | -1.8035 | 0.020503 | 1.6882 |
| PE 52:4;O | 0.47339 | -1.0789 | 0.020509 | 1.6881 |
| FA 24:1 | 0.54002 | -0.88893 | 0.020541 | 1.6874 |
| ST 28:3;O4 | 0.43866 | -1.1888 | 0.020571 | 1.6867 |
| TG 94:4;O2 | 0.42837 | -1.2231 | 0.020624 | 1.6856 |
| Cer 31:4;O | 1.4991 | 0.58412 | 0.020639 | 1.6853 |
| Cer 43:5;O2 | 1.5913 | 0.67016 | 0.020641 | 1.6853 |
| SHEX Cer 35:4;O3 | 1.4749 | 0.56066 | 0.020642 | 1.6853 |
| PIP 66:2 | 0.50981 | -0.97197 | 0.020782 | 1.6823 |
| PC 38:5 | 5.6232 | 2.4914 | 0.020917 | 1.6795 |
| CoA 21:6;O2 | 13.066 | 3.7078 | 0.020955 | 1.6787 |
| WE 23:3 | 1.5753 | 0.65567 | 0.020992 | 1.6779 |
| TG 45:1 | 2.1118 | 1.0784 | 0.021099 | 1.6757 |
| DG 47:1 | 3.7872 | 1.9211 | 0.021286 | 1.6719 |
| PG 32:0 | 0.64539 | -0.63175 | 0.02137 | 1.6702 |
| CL 73:4 | 0.48215 | -1.0524 | 0.021403 | 1.6695 |
| PE 17:0_18:1 | 0.51076 | -0.96929 | 0.021553 | 1.6665 |
| DG 56:8 | 0.42704 | -1.2276 | 0.021714 | 1.6633 |
| PI-Cer 38:3;O4 | 0.11978 | -3.0616 | 0.021865 | 1.6603 |
| PA 51:0 | 6.5181 | 2.7044 | 0.021925 | 1.6591 |
| PC 48:7;O | 23.289 | 4.5416 | 0.021968 | 1.6582 |
| PR 30:7 | 0.51392 | -0.96039 | 0.022052 | 1.6566 |
| NAE 20:4 | 0.055943 | -4.1599 | 0.022111 | 1.6554 |
| PC O-22:3_26:1 | 0.36107 | -1.4697 | 0.022161 | 1.6544 |
| Cer 38:0;O4 | 0.60089 | -0.73482 | 0.022196 | 1.6537 |
| PC 18:0_20:0 | 6.2055 | 2.6335 | 0.022236 | 1.653 |
| PA O-32:7 | 20.03 | 4.3241 | 0.022293 | 1.6518 |
| ST 29:6;O8;Hex | 0.33091 | -1.5955 | 0.022297 | 1.6517 |
| PI O-32:5 | 1.5399 | 0.6228 | 0.022363 | 1.6505 |
| CerP 32:1;O2 | 3.4811 | 1.7996 | 0.022377 | 1.6502 |
| HexCer 35:3;O6 | 2.2187 | 1.1497 | 0.022518 | 1.6475 |
| DG 40:2 | 0.4701 | -1.089 | 0.022527 | 1.6473 |
| PIP 52:4 | 0.60841 | -0.71689 | 0.022539 | 1.6471 |
| MIPC 37:0;O6 | 2.2992 | 1.2011 | 0.022549 | 1.6469 |
| DG 40:4 | 0.67256 | -0.57226 | 0.022552 | 1.6468 |
| PA 35:0 | 39.751 | 5.3129 | 0.022929 | 1.6396 |
| PE 18:1_19:1 | 0.55875 | -0.83972 | 0.022983 | 1.6386 |
| Cer 22:1;O2/24:6 | 1.4094 | 0.49513 | 0.02321 | 1.6343 |
| CL 75:6 | 0.59951 | -0.73815 | 0.023276 | 1.6331 |
| SM 14:0;O2/18:1 | 6.915 | 2.7897 | 0.023276 | 1.6331 |
| DG 60:5 | 0.50013 | -0.99964 | 0.023325 | 1.6322 |
| PG 38:5 | 2.2153 | 1.1475 | 0.023539 | 1.6282 |
| NAE 31:1;O | 2.6385 | 1.3997 | 0.023563 | 1.6278 |
| DG dO-36:0 | 0.61002 | -0.71306 | 0.023586 | 1.6273 |
| PS 21:2_18:3 | 2.4617 | 1.2997 | 0.023608 | 1.6269 |
| PS 14:0_18:0 | 1.8453 | 0.88387 | 0.023679 | 1.6256 |
| BMP 19:4;O | 3.9116 | 1.9677 | 0.023752 | 1.6243 |
| GD1b d42:1 | 2.2599 | 1.1762 | 0.023759 | 1.6242 |
| TG 100:16;O3 | 0.6798 | -0.55681 | 0.02378 | 1.6238 |
| TG 16:0_18:1_18:2 | 0.15426 | -2.6965 | 0.023783 | 1.6237 |
| Hex2Cer 34:2;O2 | 2.3045 | 1.2045 | 0.023879 | 1.622 |
| HexCer 54:6;O3 | 0.49872 | -1.0037 | 0.024002 | 1.6197 |
| TG O-42:1 | 0.56186 | -0.83173 | 0.024444 | 1.6118 |
| DG 38:4 | 0.56862 | -0.81445 | 0.024548 | 1.61 |
| ST 28:3;O3;G | 0.38039 | -1.3945 | 0.024602 | 1.609 |
| ST 30:0;O8 | 0.67682 | -0.56316 | 0.024705 | 1.6072 |
| TG 50:3 | 0.32993 | -1.5998 | 0.024739 | 1.6066 |
| ST 27:2;O;Hex | 0.6542 | -0.6122 | 0.024752 | 1.6064 |
| ST 25:1;O4;Hex | 1.7192 | 0.7817 | 0.02478 | 1.6059 |
| Car 32:1;O | 0.66857 | -0.58085 | 0.024866 | 1.6044 |
| HexCer 37:3;O4 | 1.7398 | 0.79895 | 0.025027 | 1.6016 |
| DG 58:6 | 0.30902 | -1.6942 | 0.025044 | 1.6013 |
| Hex2Cer 41:5;O6 | 1.75 | 0.80736 | 0.0251 | 1.6003 |
| TG 16:1_18:0_20:2 | 0.095369 | -3.3903 | 0.025266 | 1.5975 |
| PC 48:9;O | 36.779 | 5.2008 | 0.025416 | 1.5949 |
| PEtOH 16:1_20:3 | 3.4312 | 1.7787 | 0.025444 | 1.5944 |
| PG O-56:0;O | 0.66208 | -0.59492 | 0.025478 | 1.5938 |
| Cer 37:1;O4 | 2.3964 | 1.2608 | 0.025554 | 1.5925 |
| ST 28:3;O7;T | 0.67719 | -0.56236 | 0.025622 | 1.5914 |
| Cer 29:0;O2 | 1.8269 | 0.86939 | 0.025647 | 1.591 |
| PA O-36:0 | 18.037 | 4.1729 | 0.025896 | 1.5868 |
| SHexCer 43:3;O2 | 1.6301 | 0.70499 | 0.026081 | 1.5837 |
| WE 34:0 | 3.1083 | 1.6361 | 0.026109 | 1.5832 |
| DG 32:0 | 0.48474 | -1.0447 | 0.02615 | 1.5825 |
| TG 97:6;O2 | 0.60239 | -0.73122 | 0.026195 | 1.5818 |
| FAL 17:2 | 2.1999 | 1.1375 | 0.026288 | 1.5802 |
| PS 20:1;O | 2.9039 | 1.538 | 0.026291 | 1.5802 |
| Car 32:0;O4 | 2.2708 | 1.1832 | 0.026418 | 1.5781 |
| PA 20:1;O2 | 15.126 | 3.919 | 0.026609 | 1.575 |
| PA 43:4 | 7.6422 | 2.934 | 0.026698 | 1.5735 |
| Car 34:6;O | 0.51421 | -0.95958 | 0.026837 | 1.5713 |
| HexCer 44:6;O6 | 0.57622 | -0.79531 | 0.026846 | 1.5711 |
| LPI 11:3 | 1.597 | 0.6754 | 0.026848 | 1.5711 |
| PE 38:0 | 0.70328 | -0.50782 | 0.026866 | 1.5708 |
| LPG 22:6 | 2.8608 | 1.5164 | 0.026887 | 1.5705 |
| DG 54:0 | 0.60264 | -0.73063 | 0.026956 | 1.5694 |
| PE 23:1; PE 2:0-21:1; [M-H]- | 1.7263 | 0.78765 | 0.027154 | 1.5662 |
| PE-Cer 15:0;2O/16:4 | 1.7043 | 0.76919 | 0.027189 | 1.5656 |
| PE 3:0_18:4 | 1.9298 | 0.94848 | 0.027192 | 1.5656 |
| FOH 20:0;O | 2.7893 | 1.4799 | 0.027205 | 1.5654 |
| TG 82:3;O2 | 3.6887 | 1.8831 | 0.027231 | 1.5649 |
| HexCer 18:3;O2/26:2 | 2.2642 | 1.179 | 0.027252 | 1.5646 |
| DG 61:2;O | 2.2597 | 1.1761 | 0.02727 | 1.5643 |
| TG 14:0_18:1_18:2 | 0.2652 | -1.9148 | 0.027287 | 1.564 |
| PEtOH 16:0_18:1 | 3.2994 | 1.7222 | 0.027477 | 1.561 |
| SM 16:0;O2/18:0 | 4.3231 | 2.1121 | 0.027496 | 1.5607 |
| Cer 39:1;O4 | 3.7829 | 1.9195 | 0.027551 | 1.5599 |
| FA 32:0;O4 | 2.8321 | 1.5019 | 0.027764 | 1.5565 |
| TG 67:11;O | 0.60906 | -0.71534 | 0.027782 | 1.5562 |
| PG 25:1;O2 | 1.7543 | 0.81088 | 0.02787 | 1.5549 |
| CerP 54:5;O3 | 0.25896 | -1.9492 | 0.027926 | 1.554 |
| HexCer 37:2;O4 | 1.415 | 0.50078 | 0.027997 | 1.5529 |
| CL 71:6 | 0.70353 | -0.50732 | 0.028003 | 1.5528 |
| PA O-58:8 | 4.555 | 2.1874 | 0.028176 | 1.5501 |
| ST 29:3;O6 | 0.66487 | -0.58885 | 0.028373 | 1.5471 |
| PA O-47:1 | 10.743 | 3.4254 | 0.028385 | 1.5469 |
| LPI 12:3 | 2.0655 | 1.0465 | 0.028418 | 1.5464 |
| DG O-36:1 | 1.5278 | 0.61144 | 0.028476 | 1.5455 |
| MG O-20:0 | 0.63513 | -0.65488 | 0.028575 | 1.544 |
| HexCer 41:0;O3 | 1.6775 | 0.74635 | 0.028858 | 1.5397 |
| SM 15:0;O2/18:1 | 12.786 | 3.6765 | 0.028903 | 1.5391 |
| PA O-45:1 | 14.603 | 3.8682 | 0.029013 | 1.5374 |
| HexCer 32:5;O3 | 1.7632 | 0.81822 | 0.029176 | 1.535 |
| MGDG 18:0_20:1 | 0.62308 | -0.6825 | 0.029259 | 1.5337 |
| DG 52:1 | 2.7687 | 1.4692 | 0.029537 | 1.5296 |
| PIP3 38:9 | 0.52692 | -0.92433 | 0.029752 | 1.5265 |
| TG 54:3 | 0.38147 | -1.3904 | 0.030103 | 1.5214 |
| TG 16:1_18:1_19:1 | 0.14718 | -2.7644 | 0.030109 | 1.5213 |
| PC O-36:0 | 4.8612 | 2.2813 | 0.030157 | 1.5206 |
| PE 56:3 | 0.46801 | -1.0954 | 0.030306 | 1.5185 |
| DG 58:2 | 0.68938 | -0.53662 | 0.030317 | 1.5183 |
| ST 23:2;O3;G | 0.2075 | -2.2688 | 0.030525 | 1.5153 |
| PA O-56:7 | 0.57395 | -0.801 | 0.030673 | 1.5132 |
| HexCer 40:0;O5 | 0.43259 | -1.2089 | 0.030691 | 1.513 |
| Hex2Cer 33:0;O5 | 1.4753 | 0.56097 | 0.030719 | 1.5126 |
| TG 60:3 | 0.14974 | -2.7395 | 0.030781 | 1.5117 |
| NAE 20:0 | 0.6996 | -0.5154 | 0.030814 | 1.5112 |
| ST 30:4;O5 | 0.7055 | -0.50328 | 0.030835 | 1.511 |
| PC 16:0_24:1 | 0.69923 | -0.51616 | 0.030877 | 1.5104 |
| CL 72:3 | 0.47212 | -1.0828 | 0.030971 | 1.509 |
| Hex(2)-HexNeuAc(3)-Cer 34:1;O2 | 2.1313 | 1.0917 | 0.031003 | 1.5086 |
| NAE 20:2 | 1.5334 | 0.61672 | 0.031065 | 1.5077 |
| ST 30:0;O2;GlcA | 0.65323 | -0.61433 | 0.03114 | 1.5067 |
| HexCer 15:3;O2/28:7 | 0.68403 | -0.54786 | 0.031162 | 1.5064 |
| PE O-38:5 | 0.67247 | -0.57246 | 0.031286 | 1.5047 |
| PE 14:1_18:1 | 0.31269 | -1.6772 | 0.031337 | 1.5039 |
| NA 22:0 | 1.539 | 0.62197 | 0.031398 | 1.5031 |
| CE 22:2 | 0.36647 | -1.4482 | 0.031503 | 1.5017 |
| PC O-18:1_22:6 | 0.44671 | -1.1626 | 0.031565 | 1.5008 |
| ST 21:1;O6 | 1.5921 | 0.67096 | 0.031606 | 1.5002 |
| PA O-43:1 | 17.591 | 4.1368 | 0.031679 | 1.4992 |
| PR 20:2;O2 | 0.42797 | -1.2244 | 0.031788 | 1.4977 |
| DG 54:4 | 0.41677 | -1.2627 | 0.032088 | 1.4937 |
| PG 40:4 | 0.61795 | -0.69443 | 0.032094 | 1.4936 |
| TG 52:5 | 0.38777 | -1.3667 | 0.032205 | 1.4921 |
| DG 42:9 | 0.5865 | -0.76981 | 0.032217 | 1.4919 |
| SHexCer 32:0;O3 | 2.9669 | 1.5689 | 0.032557 | 1.4874 |
| LPG 32:1 | 0.44242 | -1.1765 | 0.032796 | 1.4842 |
| PS 18:0_22:1 | 1.5917 | 0.67059 | 0.032929 | 1.4824 |
| PE 25:1;O | 0.71333 | -0.48736 | 0.033019 | 1.4812 |
| PA 37:0 | 8.3609 | 3.0637 | 0.033308 | 1.4774 |
| PC 36:0;O | 2.8729 | 1.5225 | 0.033355 | 1.4768 |
| HexCer 41:1;O4 | 0.55302 | -0.85461 | 0.033362 | 1.4767 |
| HexCer 39:6;O3 | 0.62996 | -0.66666 | 0.033538 | 1.4745 |
| Cer 28:6;O | 1.5477 | 0.63011 | 0.033625 | 1.4733 |
| Cer 40:1;O6 | 1.5477 | 0.63011 | 0.033625 | 1.4733 |
| Cer 44:5;O | 1.5477 | 0.63011 | 0.033625 | 1.4733 |
| Cer 45:4;O3 | 1.5477 | 0.63011 | 0.033625 | 1.4733 |
| Cer 47:1;O3 | 1.5477 | 0.63011 | 0.033625 | 1.4733 |
| Cer 47:6;O2 | 1.5477 | 0.63011 | 0.033625 | 1.4733 |
| Cer 52:6;O | 1.5477 | 0.63011 | 0.033625 | 1.4733 |
| CerP 44:5;O5 | 1.5477 | 0.63011 | 0.033625 | 1.4733 |
| Hex2Cer 40:6;O2 | 1.5477 | 0.63011 | 0.033625 | 1.4733 |
| HexCer 32:5;O5 | 1.5477 | 0.63011 | 0.033625 | 1.4733 |
| SPB 18:1;O3 | 1.5477 | 0.63011 | 0.033625 | 1.4733 |
| SPB 20:2;O6 | 1.5477 | 0.63011 | 0.033625 | 1.4733 |
| HexCer 28:0;O4 | 0.67445 | -0.56823 | 0.033636 | 1.4732 |
| PC 30:0;O | 2.1457 | 1.1015 | 0.033639 | 1.4732 |
| PA 35:1 | 2.145 | 1.101 | 0.033659 | 1.4729 |
| DG 52:2 | 0.55227 | -0.85657 | 0.033667 | 1.4728 |
| TG 14:0_16:0_22:6 | 0.36766 | -1.4436 | 0.033751 | 1.4717 |
| MIPC 34:2;O2 | 0.55358 | -0.85314 | 0.033855 | 1.4704 |
| PC O-20:3_16:0 | 3.6031 | 1.8492 | 0.033858 | 1.4703 |
| SM 34:2;O2 | 26.885 | 4.7487 | 0.033882 | 1.47 |
| PA O-35:9 | 12.711 | 3.668 | 0.033938 | 1.4693 |
| PG 55:3 | 0.66346 | -0.59192 | 0.03403 | 1.4681 |
| DG 32:7 | 2.4245 | 1.2777 | 0.034243 | 1.4654 |
| PR 50:10;O | 0.70807 | -0.49803 | 0.034297 | 1.4647 |
| TG O-44:0;O2 | 1.5334 | 0.61676 | 0.034339 | 1.4642 |
| CL 71:3 | 0.44666 | -1.1628 | 0.034378 | 1.4637 |
| ST 23:5;O3 | 1.426 | 0.51194 | 0.034473 | 1.4625 |
| PA 40:3;O | 22.906 | 4.5177 | 0.034579 | 1.4612 |
| PA O-43:0 | 13.909 | 3.7979 | 0.034668 | 1.4601 |
| MG 21:1 | 2.0768 | 1.0544 | 0.034724 | 1.4594 |
| SHexCer 40:3;O2 | 1.5392 | 0.62216 | 0.034735 | 1.4592 |
| TG 58:3;O2 | 0.29685 | -1.7522 | 0.034787 | 1.4586 |
| LPI O-32:0 | 0.45535 | -1.135 | 0.034889 | 1.4573 |
| LPG O-31:5 | 1.8335 | 0.87458 | 0.034932 | 1.4568 |
| PIP 60:3;O | 0.50581 | -0.98334 | 0.034946 | 1.4566 |
| SM 33:0;O6 | 12.266 | 3.6166 | 0.034971 | 1.4563 |
| PE O-18:2_24:2 | 0.62723 | -0.67294 | 0.035021 | 1.4557 |
| HBMP 14:0_20:2_20:2 | 0.70754 | -0.49911 | 0.035081 | 1.4549 |
| TG O-52:0;O2 | 2.0845 | 1.0597 | 0.035212 | 1.4533 |
| DG 50:3 | 0.32287 | -1.631 | 0.035229 | 1.4531 |
| CerP 54:1;O3 | 1.5362 | 0.61932 | 0.035497 | 1.4498 |
| ST 29:0;O8;HexNAc | 1.5627 | 0.64406 | 0.03564 | 1.4481 |
| PC 44:5 | 2.115 | 1.0807 | 0.03586 | 1.4454 |
| PA 40:0 | 4.7609 | 2.2512 | 0.035873 | 1.4452 |
| PC O-18:0_24:0 | 6.2633 | 2.6469 | 0.035907 | 1.4448 |
| DG 30:0;O2 | 3.4322 | 1.7791 | 0.035949 | 1.4443 |
| PG 55:2 | 0.52717 | -0.92367 | 0.035993 | 1.4438 |
| Cer 42:1;O | 0.48178 | -1.0536 | 0.036298 | 1.4401 |
| TG 60:15 | 2.3928 | 1.2587 | 0.036302 | 1.4401 |
| SM 48:1;3O | 2.2985 | 1.2007 | 0.036571 | 1.4369 |
| PI O-16:0_18:2 | 0.50963 | -0.97247 | 0.036643 | 1.436 |
| PC 16:0_20:3 | 0.53396 | -0.9052 | 0.036718 | 1.4351 |
| Hex2Cer 29:3;O2 | 3.5013 | 1.8079 | 0.036761 | 1.4346 |
| **TG 15:0_16:1_18:1** | 0.45388 | -1.1396 | 0.037141 | 1.4301 |
| **PS O-18:3_22:4** | 2.0828 | 1.0585 | 0.037187 | 1.4296 |
| **CerP 19:0;O2/2:0** | 0.56791 | -0.81627 | 0.037201 | 1.4294 |
| **NAE 20:2;O** | 1.4684 | 0.55425 | 0.037211 | 1.4293 |
| **DG O-42:1** | 1.7744 | 0.8273 | 0.037287 | 1.4284 |
| **TG 46:3** | 0.7029 | -0.50861 | 0.037291 | 1.4284 |
| **Hex(3)-HexNAc-Fuc-NeuAc(2)-Cer 36:1;O2** | 0.70244 | -0.50956 | 0.037324 | 1.428 |
| **PA O-36:0;O** | 8.9319 | 3.159 | 0.037469 | 1.4263 |
| **SM 36:0;O4** | 10.414 | 3.3805 | 0.037698 | 1.4237 |
| **CL 76:10** | 0.56429 | -0.82549 | 0.037836 | 1.4221 |
| **PA O-42:0;O** | 7.2916 | 2.8662 | 0.038015 | 1.42 |
| **LPA 34:3** | 1.7649 | 0.81958 | 0.038019 | 1.42 |
| **PC 14:0_15:1** | 2.0032 | 1.0023 | 0.038294 | 1.4169 |
| **LPE O-18:2** | 0.5284 | -0.92029 | 0.03853 | 1.4142 |
| **Cer 18:0;O3/25:** | 0.39778 | -1.33 | 0.038554 | 1.4139 |
| **TG 54:7** | 0.41662 | -1.2632 | 0.039248 | 1.4062 |
| **SM 14:0;O2/18:0** | 5.8009 | 2.5363 | 0.039249 | 1.4062 |
| **DG 53:4** | 0.52313 | -0.93475 | 0.039328 | 1.4053 |
| **TG 8:0_8:0_16:2** | 0.58066 | -0.78424 | 0.039534 | 1.403 |
| **DG O-39:3** | 1.719 | 0.78155 | 0.039616 | 1.4021 |
| **CerP 52:5;O2** | 1.5834 | 0.66307 | 0.039812 | 1.4 |
| **SM 14:0;O2/18:2** | 24.099 | 4.5909 | 0.039815 | 1.4 |
| **GM1(d18:1/24:0)** | 2.6721 | 1.418 | 0.039893 | 1.3991 |
| **CerP 51:5;O2** | 2.9516 | 1.5615 | 0.039946 | 1.3985 |
| **PE-Cer 30:0;O5** | 2.2594 | 1.1759 | 0.039956 | 1.3984 |
| **CL 69:5** | 0.53401 | -0.90506 | 0.040072 | 1.3972 |
| **PC 12:0_14:0** | 0.51219 | -0.96524 | 0.040134 | 1.3965 |
| **PI O-56:2** | 0.66924 | -0.57941 | 0.040138 | 1.3964 |
| **PA 52:4** | 1.5695 | 0.65028 | 0.040209 | 1.3957 |
| **PA 39:6** | 2.5726 | 1.3632 | 0.04022 | 1.3956 |
| **DG 53:5** | 0.51106 | -0.96845 | 0.040236 | 1.3954 |
| **HexCer 40:6;O4** | 1.7221 | 0.78416 | 0.040236 | 1.3954 |
| **PE O-18:0_24:2** | 0.18893 | -2.4041 | 0.040343 | 1.3942 |
| **TG 75:1;O3** | 0.50463 | -0.98669 | 0.040354 | 1.3941 |
| **DG O-38:4** | 0.64644 | -0.62941 | 0.040597 | 1.3915 |
| **PI 36:4** | 0.059635 | -4.0677 | 0.040689 | 1.3905 |
| **PR 20:1;O4** | 0.6609 | -0.5975 | 0.040922 | 1.388 |
| **HexCer 16:3;O2/12:1;O** | 1.8313 | 0.8729 | 0.041067 | 1.3865 |
| **DG 60:2** | 0.69936 | -0.51589 | 0.041381 | 1.3832 |
| **Cer 18:3;O2/26:2;O** | 1.5412 | 0.62403 | 0.041514 | 1.3818 |
| PA O-54:2 | 3.4645 | 1.7926 | 0.041558 | 1.3813 |
| Cer 43:4;O3 | 3.1501 | 1.6554 | 0.041667 | 1.3802 |
| SM 33:2;O2 | 15.136 | 3.9199 | 0.04169 | 1.38 |
| DG 52:5 | 0.43934 | -1.1866 | 0.041958 | 1.3772 |
| Car 31:1;O2 | 1.9982 | 0.99871 | 0.04221 | 1.3746 |
| FA 21:2;O | 0.64444 | -0.63388 | 0.042211 | 1.3746 |
| CL 74:10 | 0.60937 | -0.7146 | 0.042231 | 1.3744 |
| PC 16:0_20:0 | 3.2156 | 1.6851 | 0.042285 | 1.3738 |
| TG O-81:0;O3 | 3.6149 | 1.854 | 0.042296 | 1.3737 |
| PC O-38:2 | 3.0198 | 1.5945 | 0.042329 | 1.3734 |
| CL 69:3 | 0.51345 | -0.9617 | 0.042754 | 1.369 |
| Cer 53:6;O | 1.9721 | 0.97973 | 0.042764 | 1.3689 |
| DG 62:6 | 0.35425 | -1.4971 | 0.042813 | 1.3684 |
| SM 45:2;O4 | 1.9866 | 0.9903 | 0.042911 | 1.3674 |
| Cer 40:2;O3 | 0.64483 | -0.63301 | 0.043106 | 1.3655 |
| PG 53:2 | 0.51721 | -0.95118 | 0.043224 | 1.3643 |
| PA 18:1_26:2 | 1.7704 | 0.82404 | 0.043434 | 1.3622 |
| PE 38:5 | 0.55779 | -0.84222 | 0.04402 | 1.3564 |
| PA 29:0 | 8.6217 | 3.108 | 0.044052 | 1.356 |
| GD1a<br>(NeuAc/NeuGc)(d18:1/16:0) | 3.5988 | 1.8475 | 0.044161 | 1.355 |
| LPC O-24:5 | 1.6494 | 0.72196 | 0.044272 | 1.3539 |
| FAHFA 14:0/14:0 | 3.7328 | 1.9002 | 0.044403 | 1.3526 |
| PC 15:0_16:0 | 2.8129 | 1.4921 | 0.044726 | 1.3494 |
| TG 52:7;O2 | 1.5233 | 0.60715 | 0.044807 | 1.3487 |
| TG 70:13;O2 | 1.5233 | 0.60715 | 0.044807 | 1.3487 |
| TG 72:12;O2 | 1.5233 | 0.60715 | 0.044807 | 1.3487 |
| TG 81:6;O3 | 1.5233 | 0.60715 | 0.044807 | 1.3487 |
| TG 82:1;O3 | 1.5233 | 0.60715 | 0.044807 | 1.3487 |
| TG 82:4;O3 | 1.5233 | 0.60715 | 0.044807 | 1.3487 |
| TG 82:5 | 1.5233 | 0.60715 | 0.044807 | 1.3487 |
| TG 91:13;O | 1.5233 | 0.60715 | 0.044807 | 1.3487 |
| TG 93:16;O2 | 1.5233 | 0.60715 | 0.044807 | 1.3487 |
| TG O-45:2 | 1.5233 | 0.60715 | 0.044807 | 1.3487 |
| TG O-61:0;O3 | 1.5233 | 0.60715 | 0.044807 | 1.3487 |
| PEtOH 18:1_20:4 | 1.9404 | 0.95633 | 0.044997 | 1.3468 |
| SM 44:2;O4 | 4.7789 | 2.2567 | 0.045074 | 1.3461 |
| Car 16:1;O | 1.439 | 0.52509 | 0.045415 | 1.3428 |
| PE O-18:2_22:3 | 0.64259 | -0.63804 | 0.045445 | 1.3425 |
| SHexCer 32:1;O4 | 2.3597 | 1.2386 | 0.045541 | 1.3416 |
| SM 44:6;O2 | 3.6846 | 1.8815 | 0.045979 | 1.3374 |
| Cer 18:1;O2/24:0 | 0.65252 | -0.61591 | 0.046283 | 1.3346 |
| DG 23:2 | 1.9629 | 0.97298 | 0.046516 | 1.3324 |
| CerP 54:6;O5 | 0.63053 | -0.66536 | 0.046595 | 1.3317 |
| PC 15:1_16:1 | 3.5892 | 1.8437 | 0.04681 | 1.3297 |
| PA 41:6 | 3.5436 | 1.8252 | 0.047027 | 1.3276 |
| TG 12:2_12:2_22:7 | 0.41442 | -1.2708 | 0.047147 | 1.3265 |
| SM 45:6;O2 | 7.5985 | 2.9257 | 0.047163 | 1.3264 |
| TG 42:0 | 0.55379 | -0.85258 | 0.047171 | 1.3263 |
| SM 40:4;O5 | 2.0298 | 1.0213 | 0.047312 | 1.325 |
| PC O-18:0_22:0 | 6.1841 | 2.6286 | 0.047323 | 1.3249 |
| ST 24:2;O2 | 0.17192 | -2.5402 | 0.047365 | 1.3245 |
| HexCer 18:0;O2/15:0;O | 1.4845 | 0.56994 | 0.047719 | 1.3213 |
| Cer 42:2;O | 0.65172 | -0.61769 | 0.047865 | 1.32 |
| SL 21:1;O_21:1;O | 0.70371 | -0.50694 | 0.048093 | 1.3179 |
| DG 18:0_22:4 | 0.43676 | -1.1951 | 0.048315 | 1.3159 |
| SPB 17:0;O3 | 1.5026 | 0.58744 | 0.048415 | 1.315 |
| DG 42:3 | 0.32929 | -1.6026 | 0.048645 | 1.313 |
| TG 90:10 | 1.9769 | 0.98325 | 0.04879 | 1.3117 |
| SHexCer 35:0;O5 | 1.4515 | 0.53757 | 0.048836 | 1.3113 |
| FOH 19:2;O3 | 0.30912 | -1.6938 | 0.048868 | 1.311 |
| HexCer 45:2;O4 | 1.4713 | 0.55707 | 0.049027 | 1.3096 |
| HexCer 46:1;O4 | 1.4792 | 0.56484 | 0.049033 | 1.3095 |
| PC O-18:0_17:0 | 8.1945 | 3.0347 | 0.049129 | 1.3087 |
| PE 36:5 | 2.3071 | 1.2061 | 0.049176 | 1.3082 |
| Cer 18:0;O2/30:7 | 1.4284 | 0.51435 | 0.049434 | 1.306 |
| Hex2Cer 44:6;O2 | 1.4253 | 0.51131 | 0.049724 | 1.3034 |
| Cer 43:5;O3 | 1.4902 | 0.57547 | 0.049728 | 1.3034 |
| PE-Cer 16:0;O2/18:1 | 1.9808 | 0.98607 | 0.049742 | 1.3033 |

**Supplemental Table 6.**
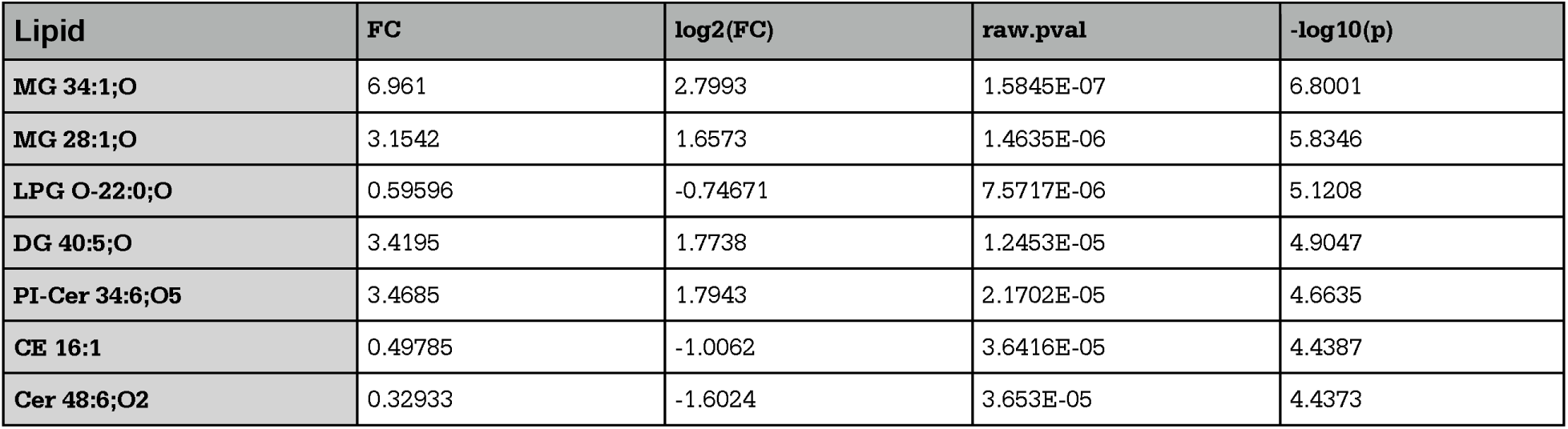
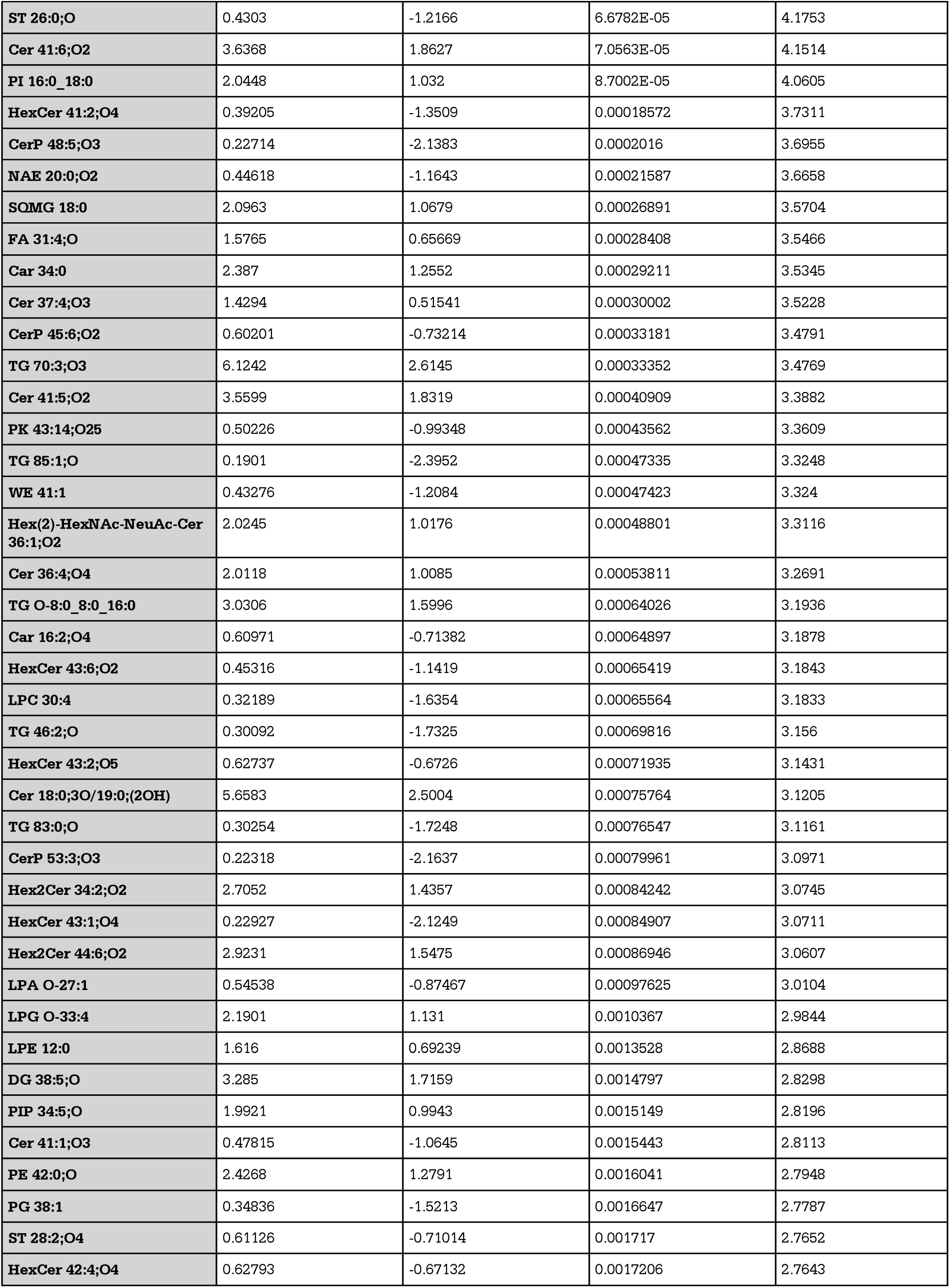
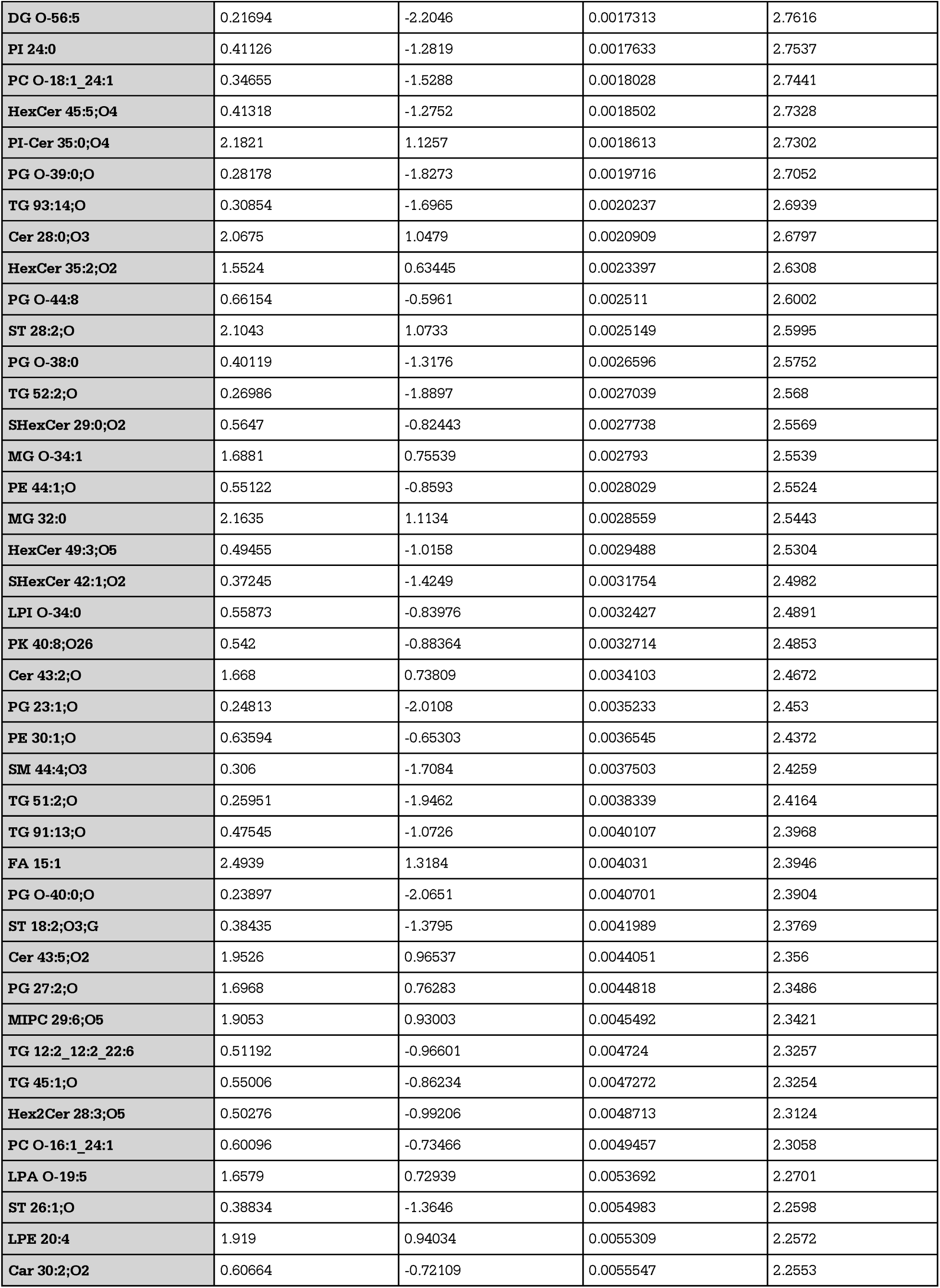
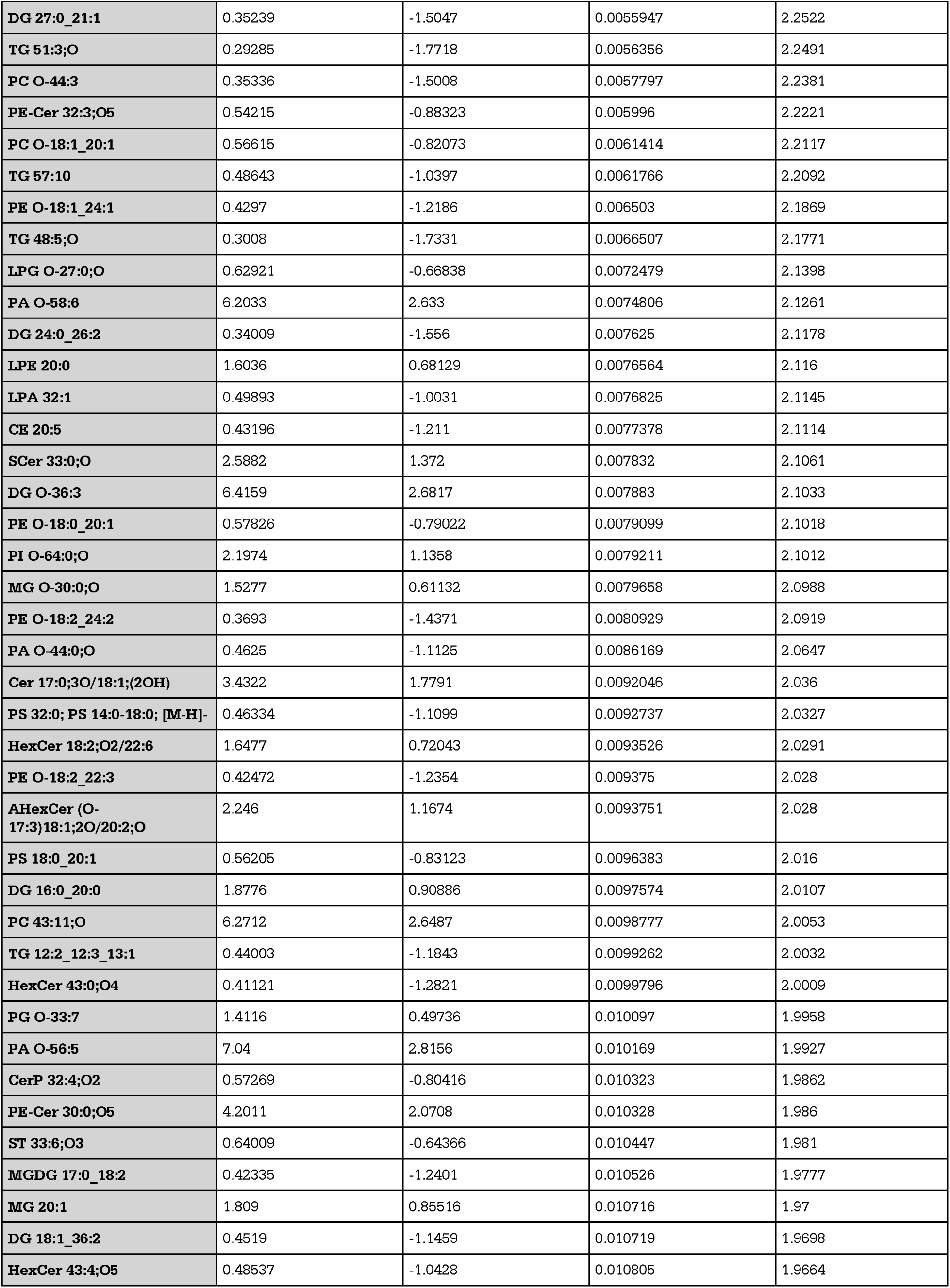
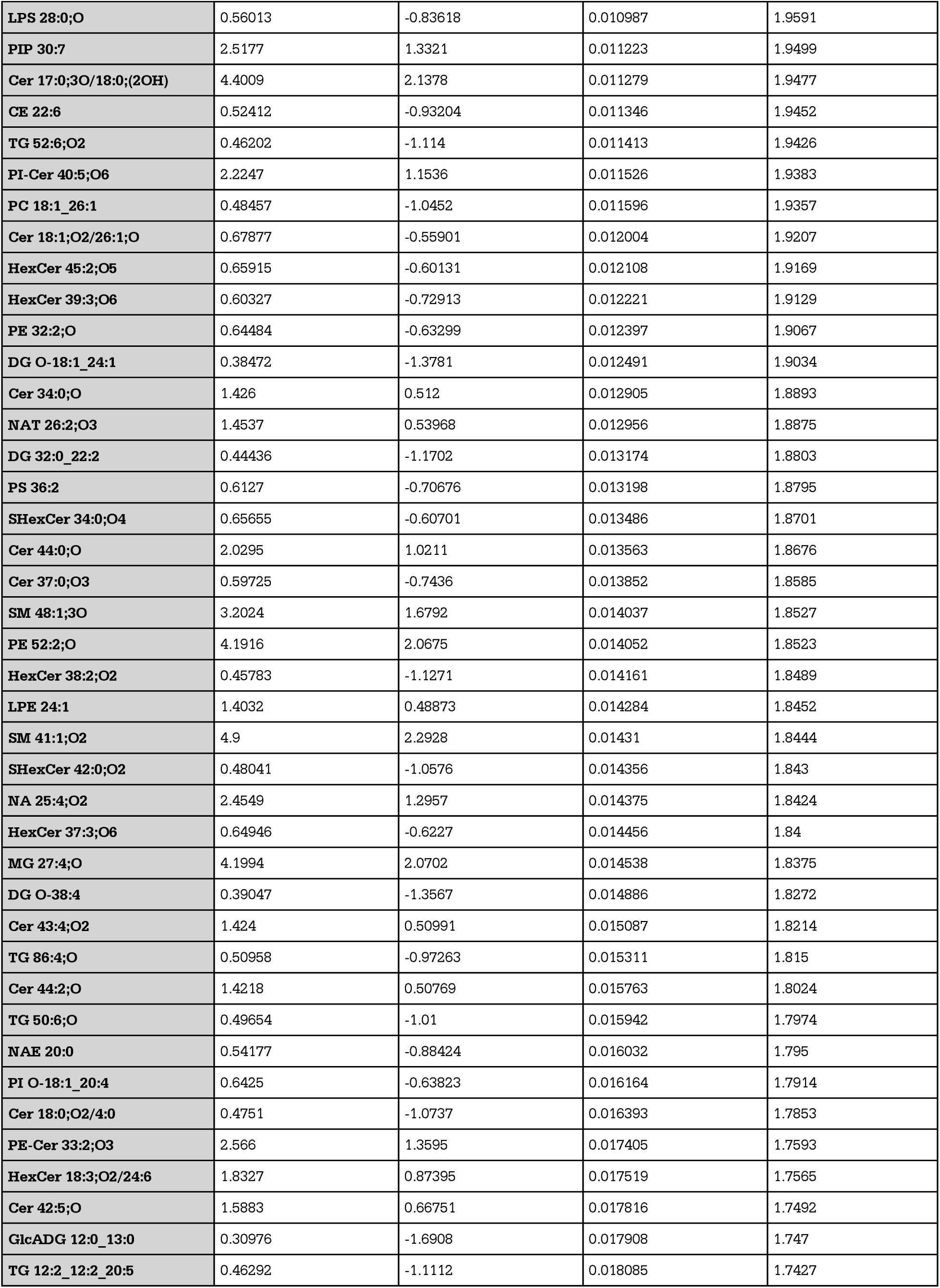
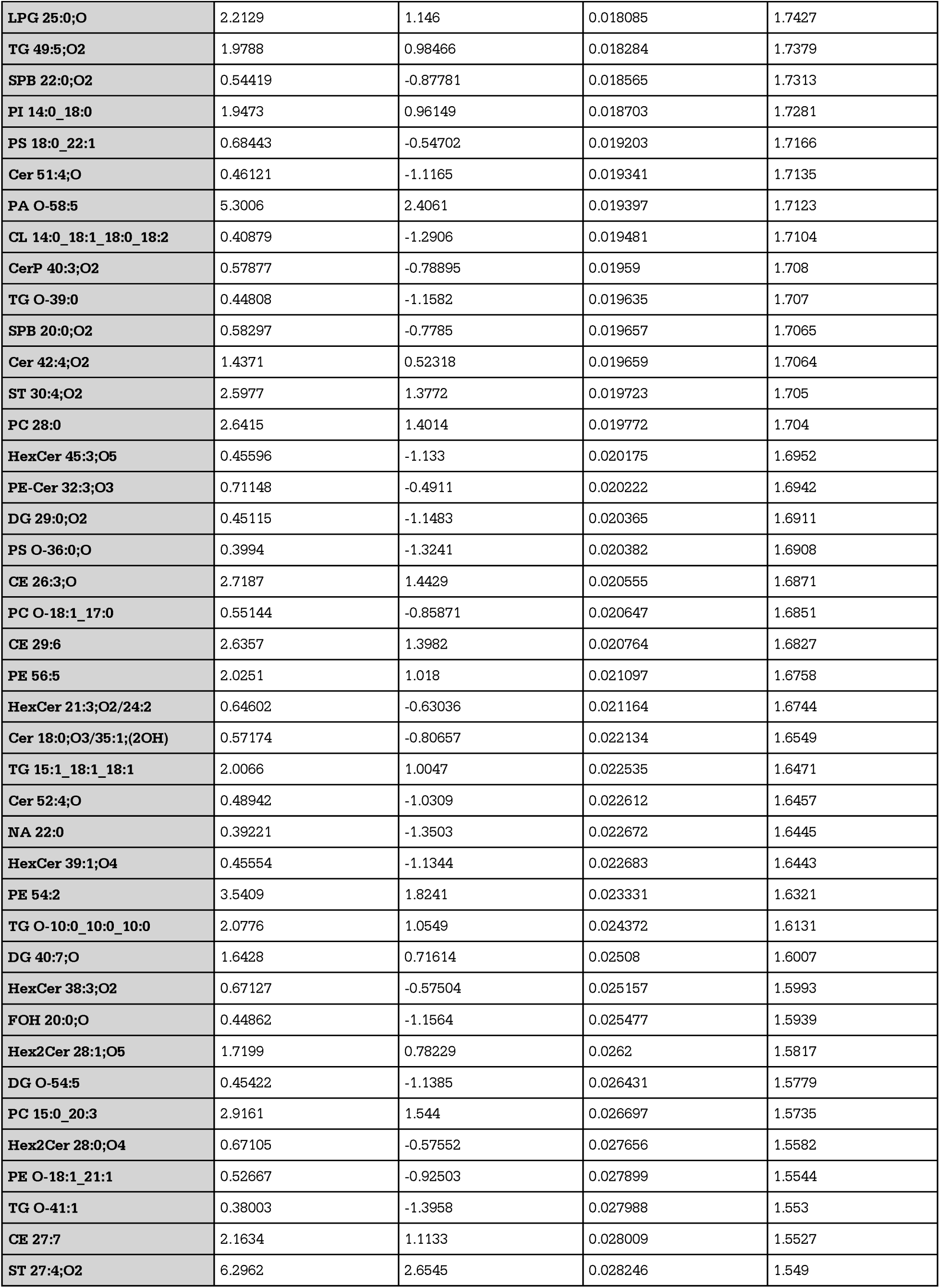
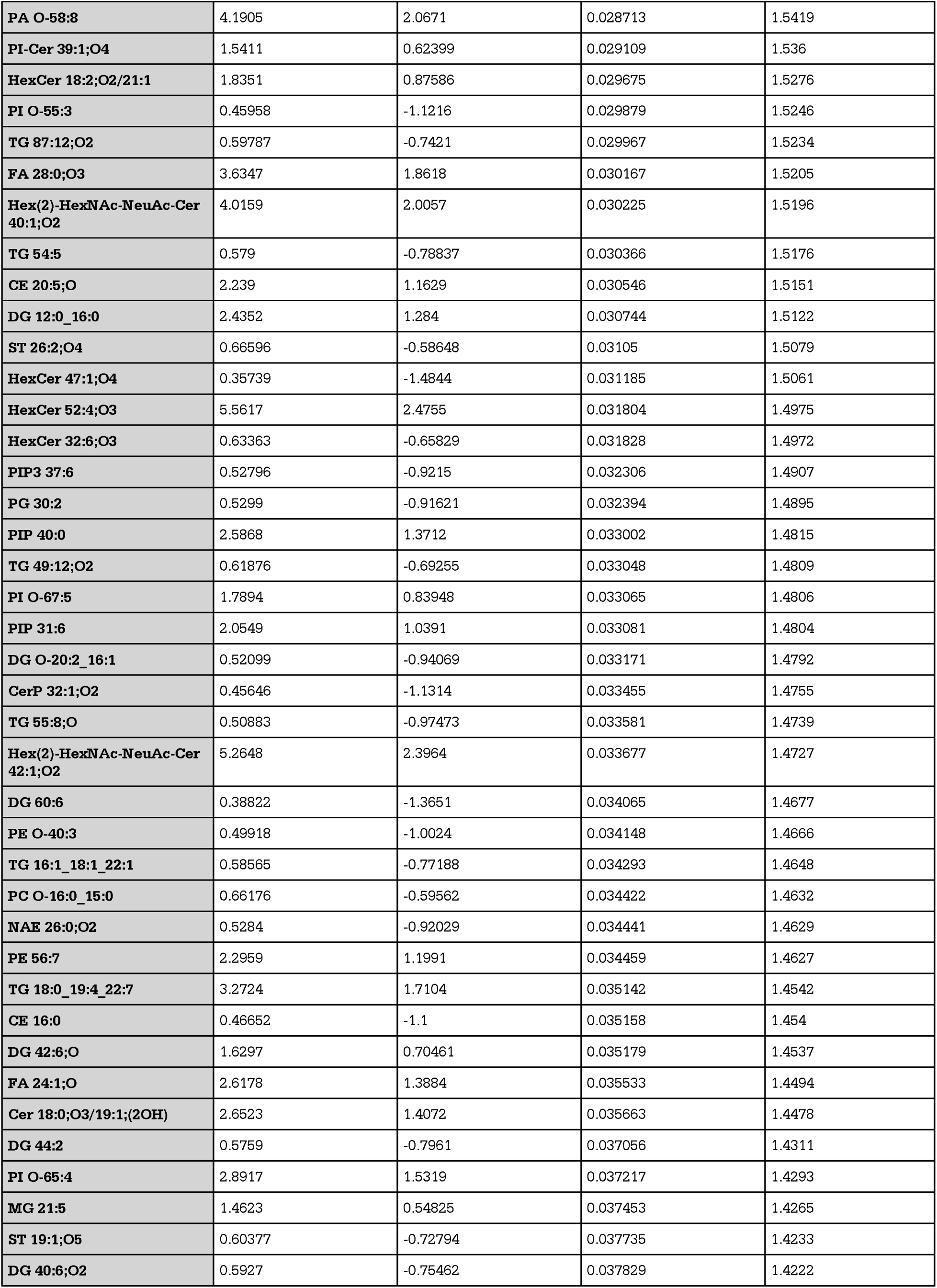
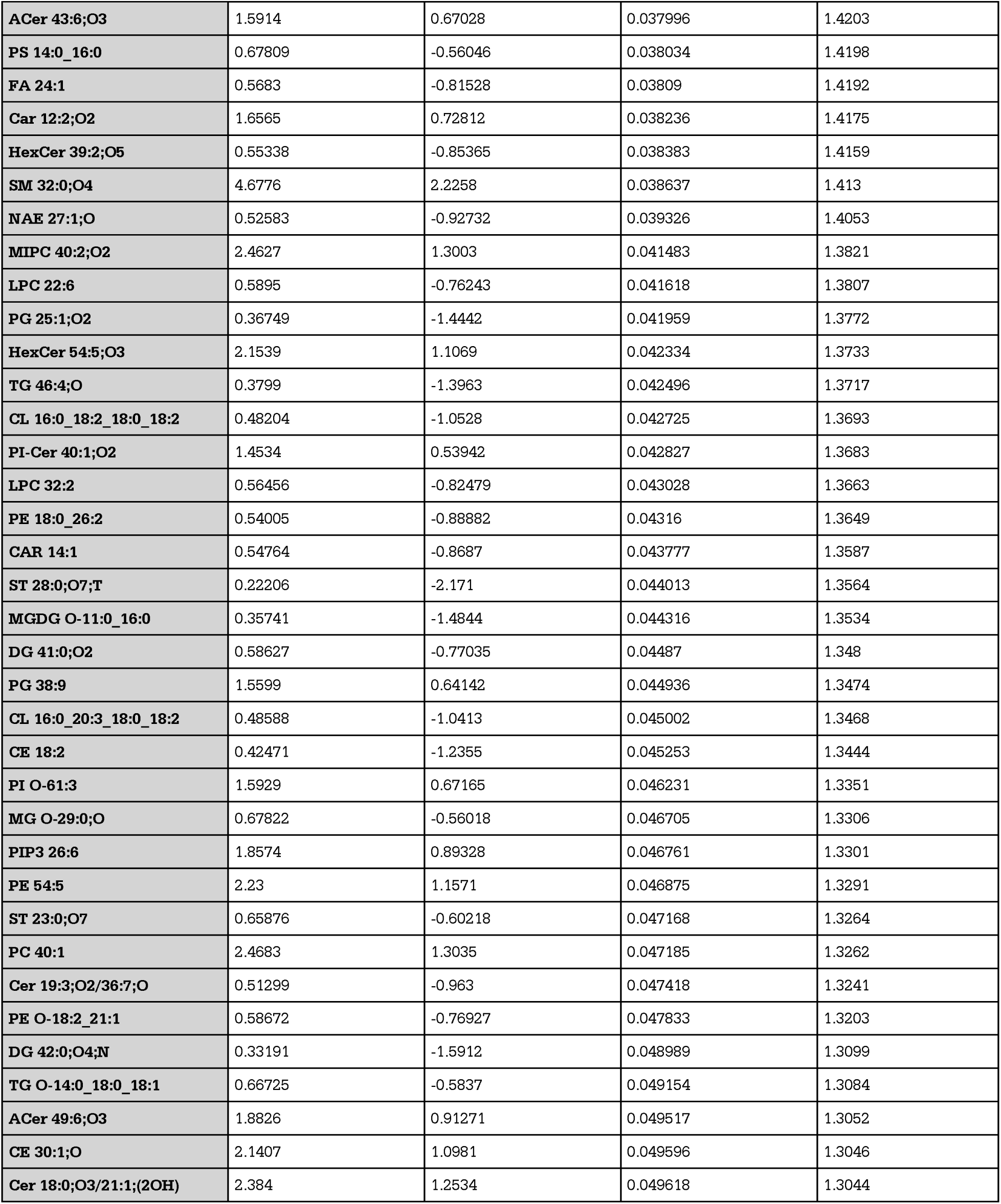
NT EVs vs LPS EVs volcano plot extended statistics. Extended statistics corresponding to Figure 3F in main text. Extended statistics include fold change (FC), log 2 fold change (log2(FC)), the raw p value (raw.pval) and the -log10 of the p-value (- log10(p)).

**Supplemental Table 7.**
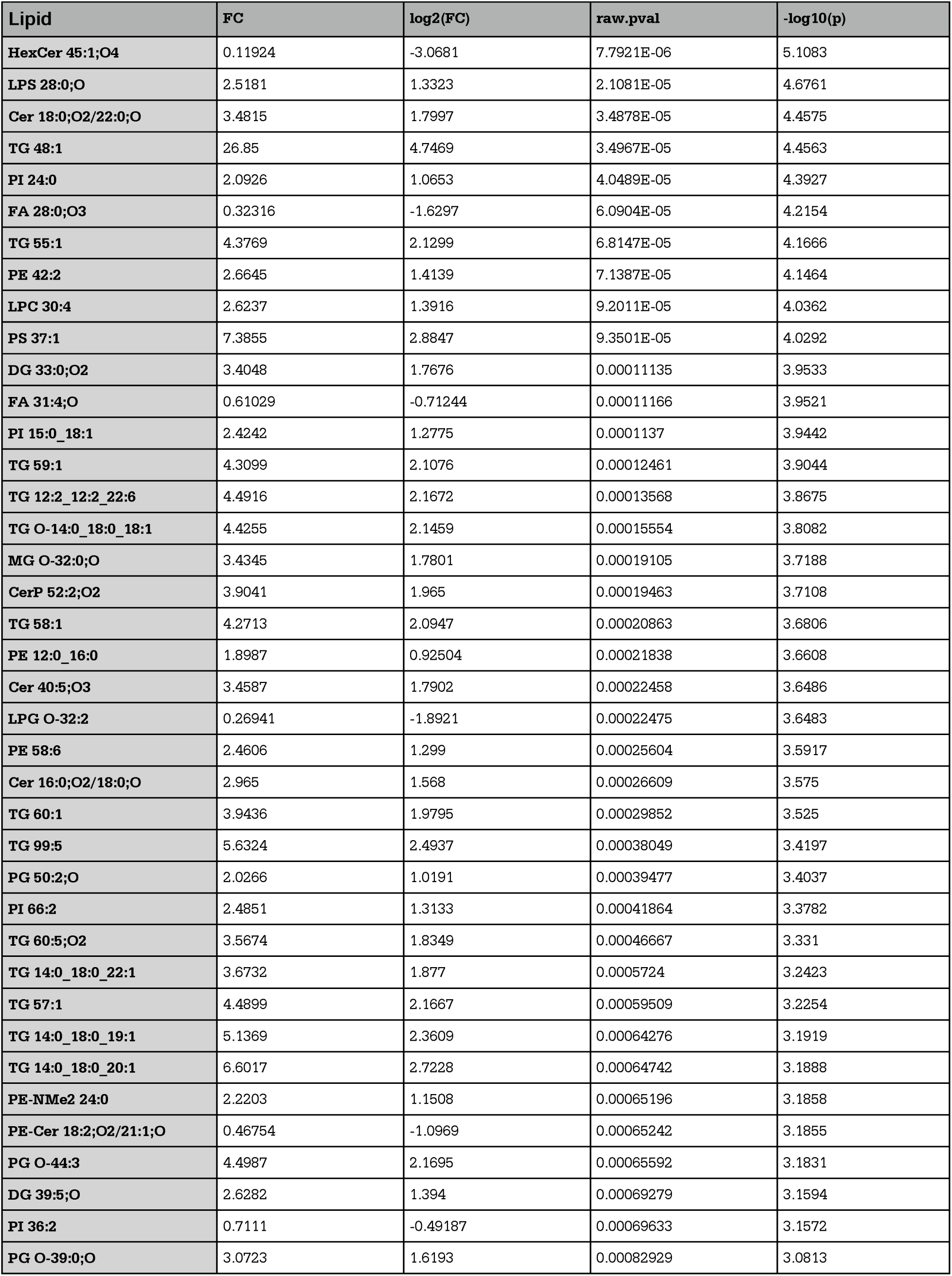
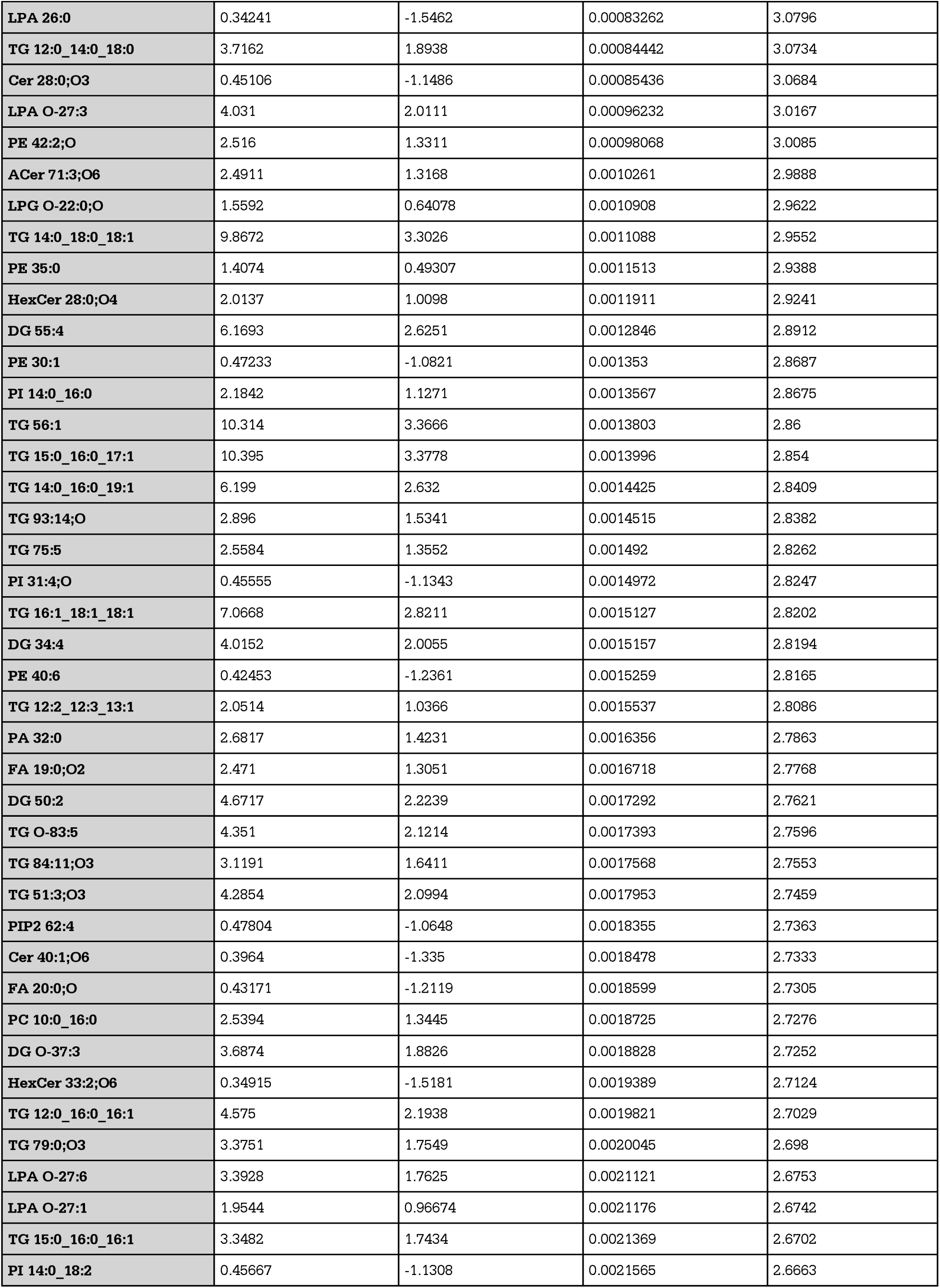
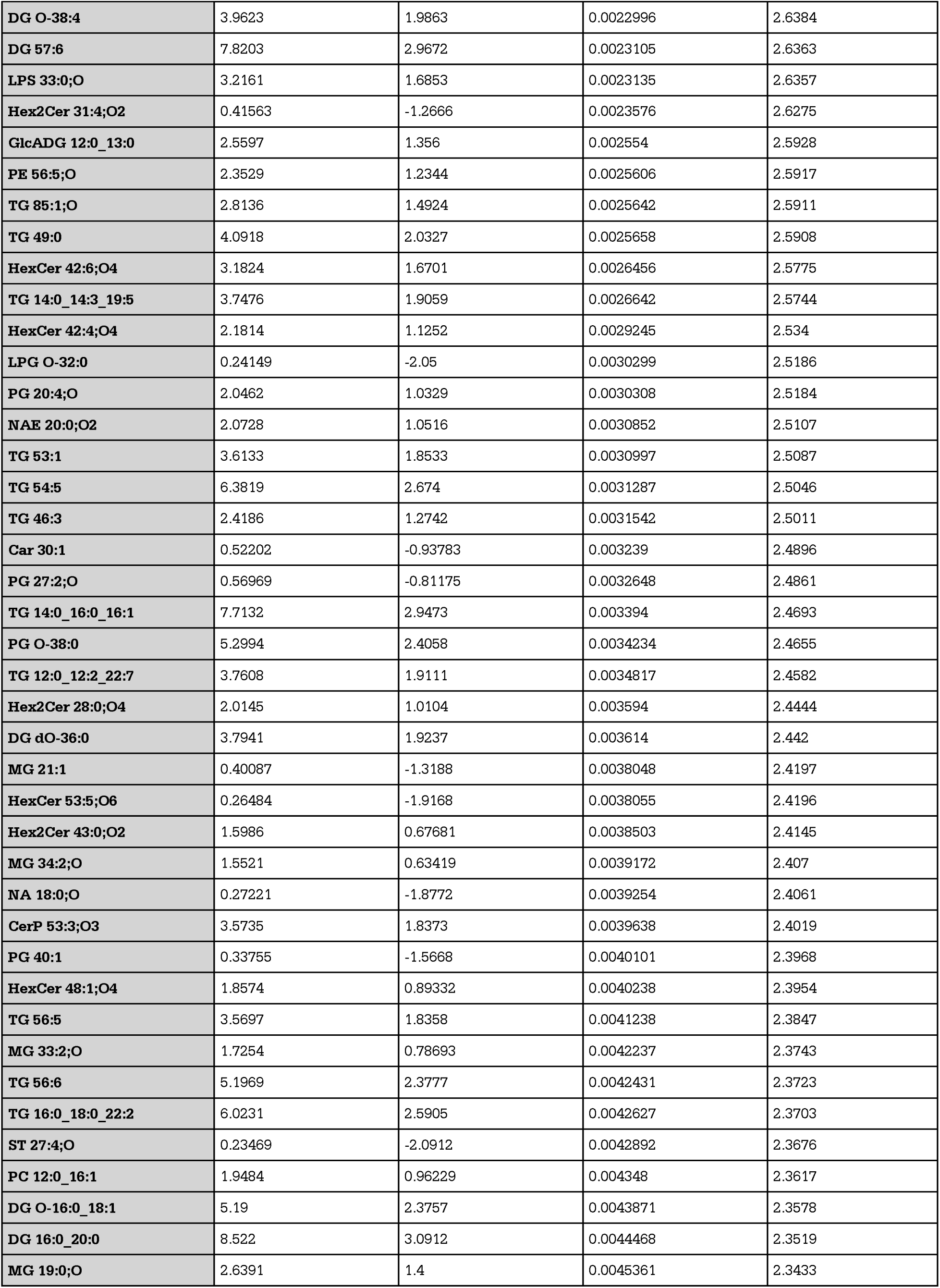
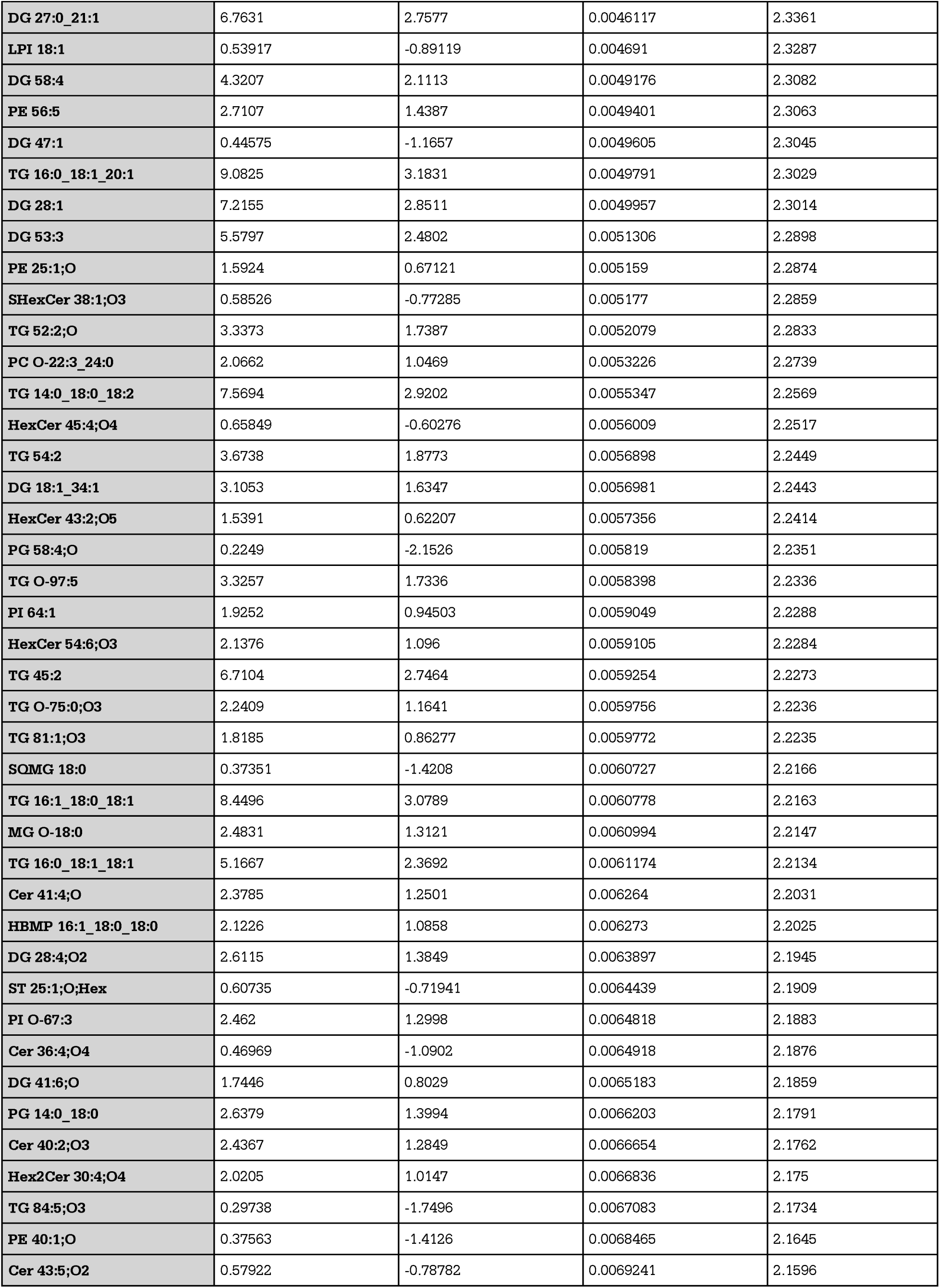
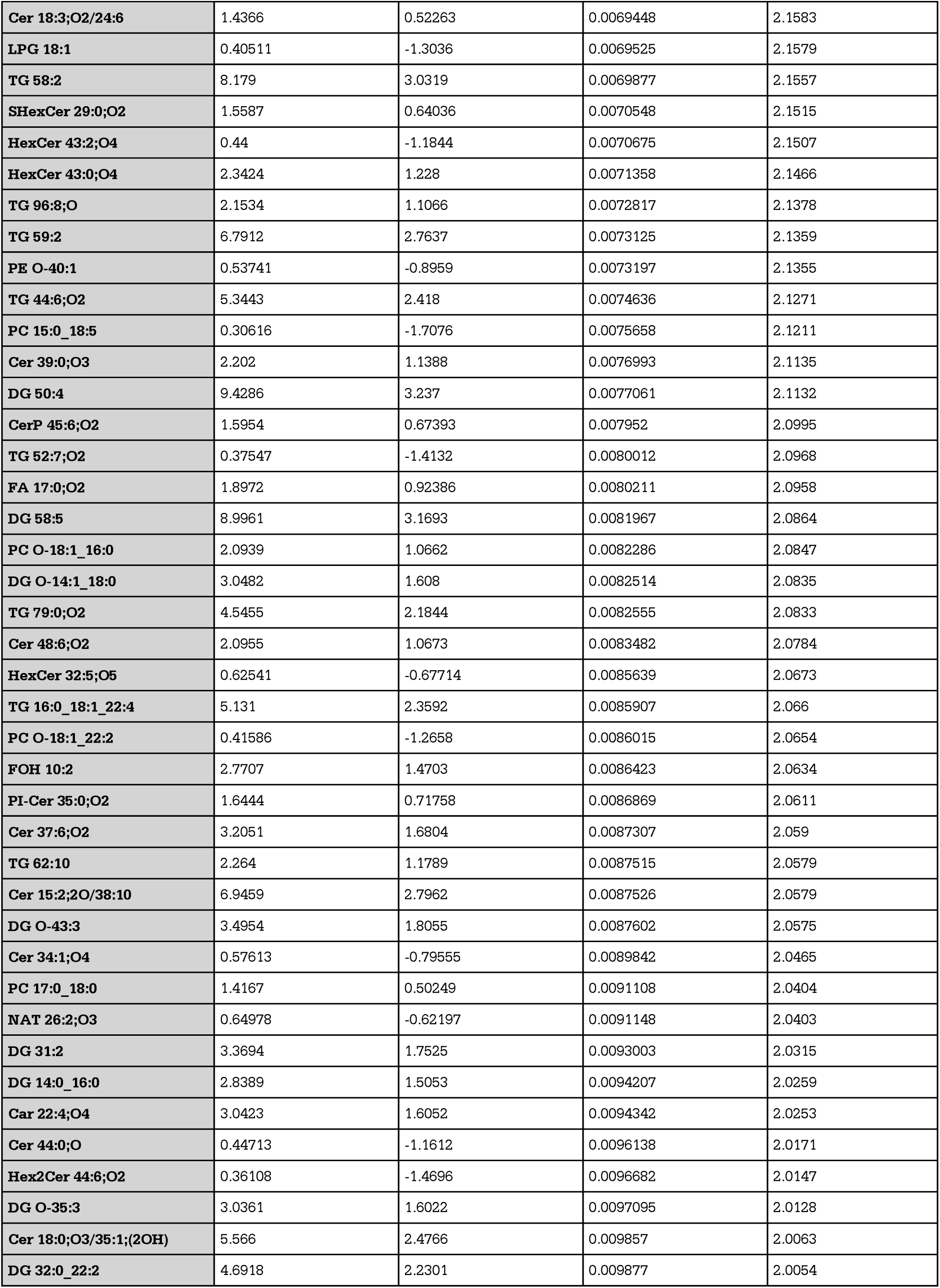
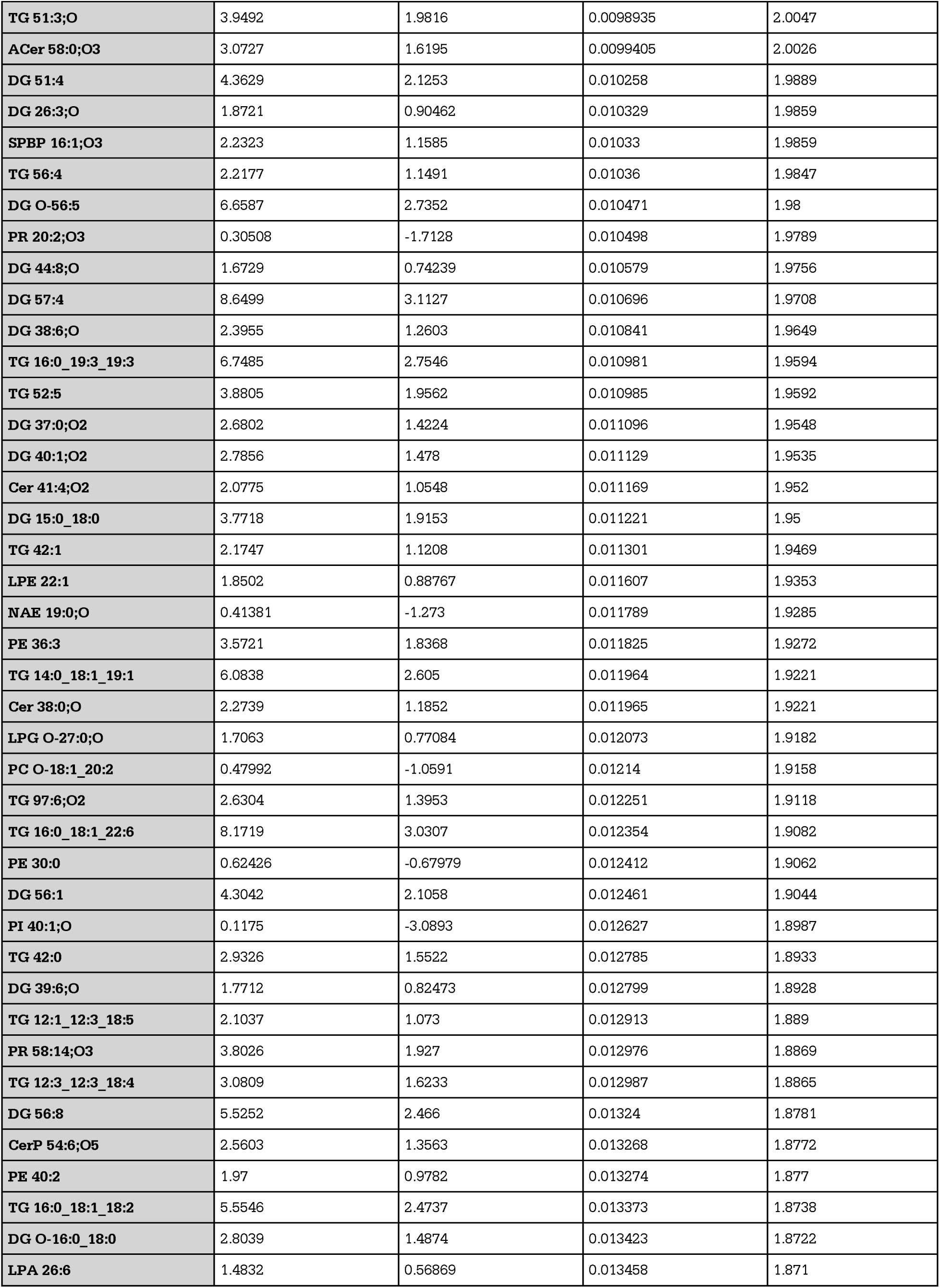
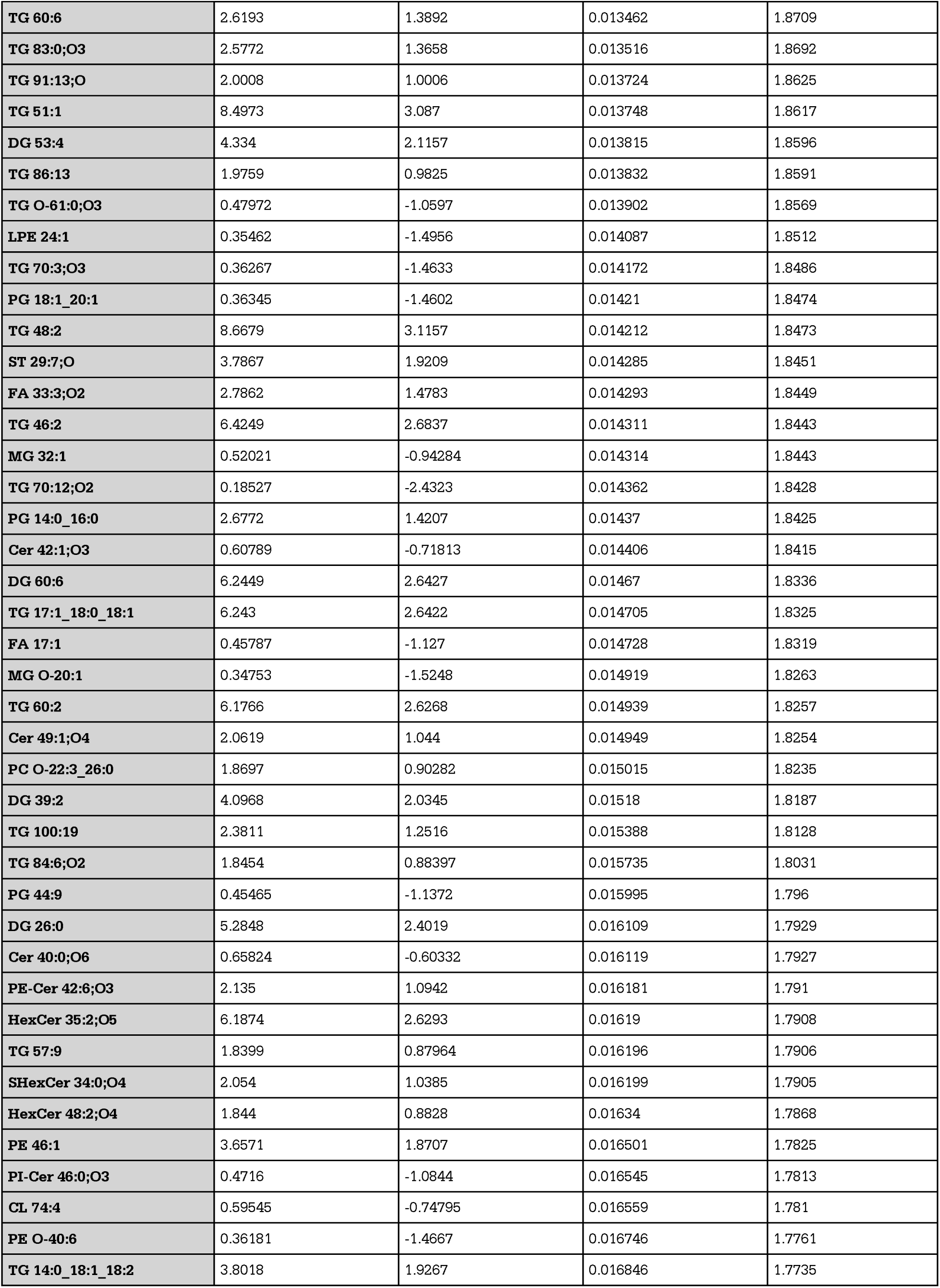
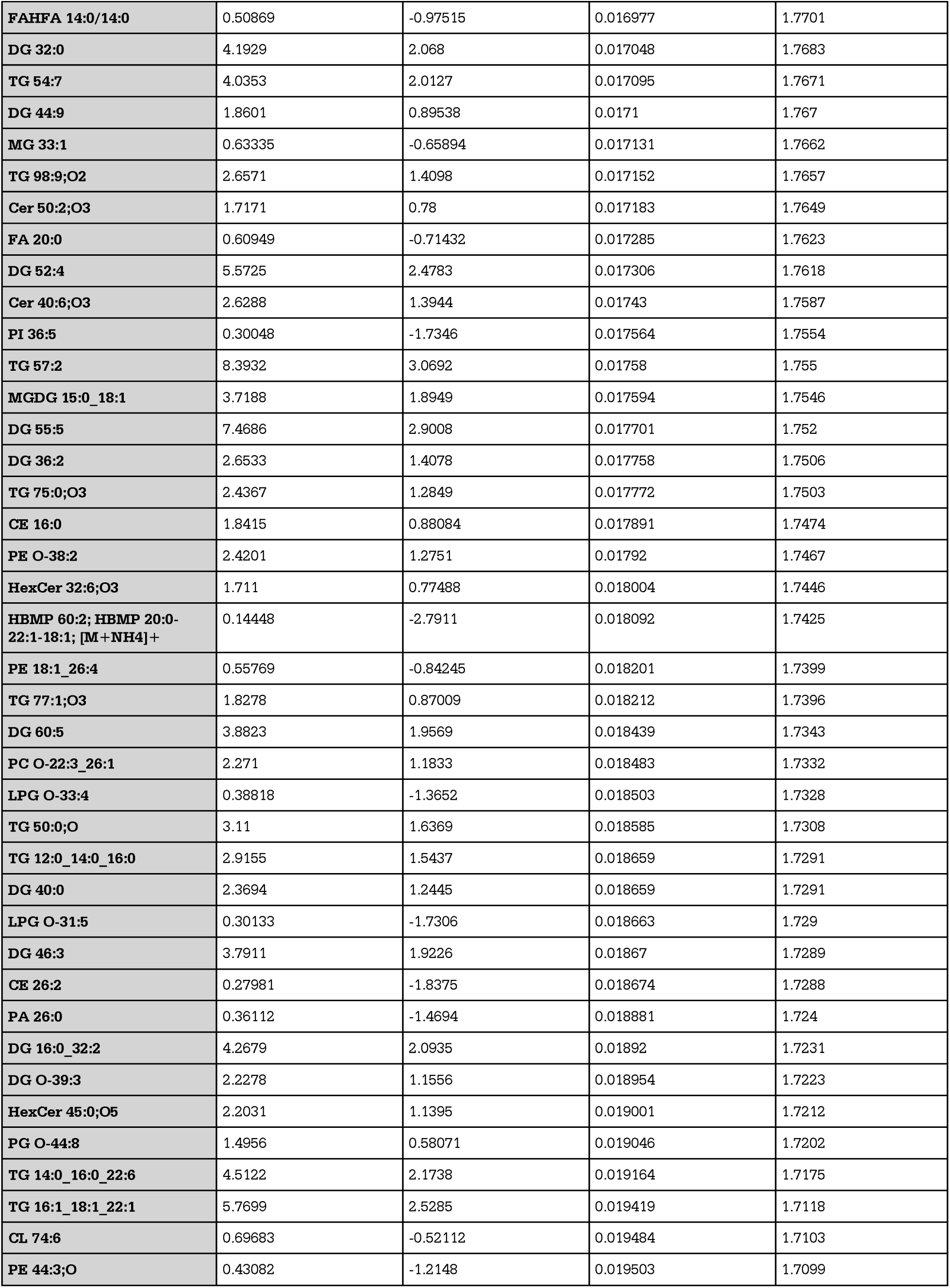
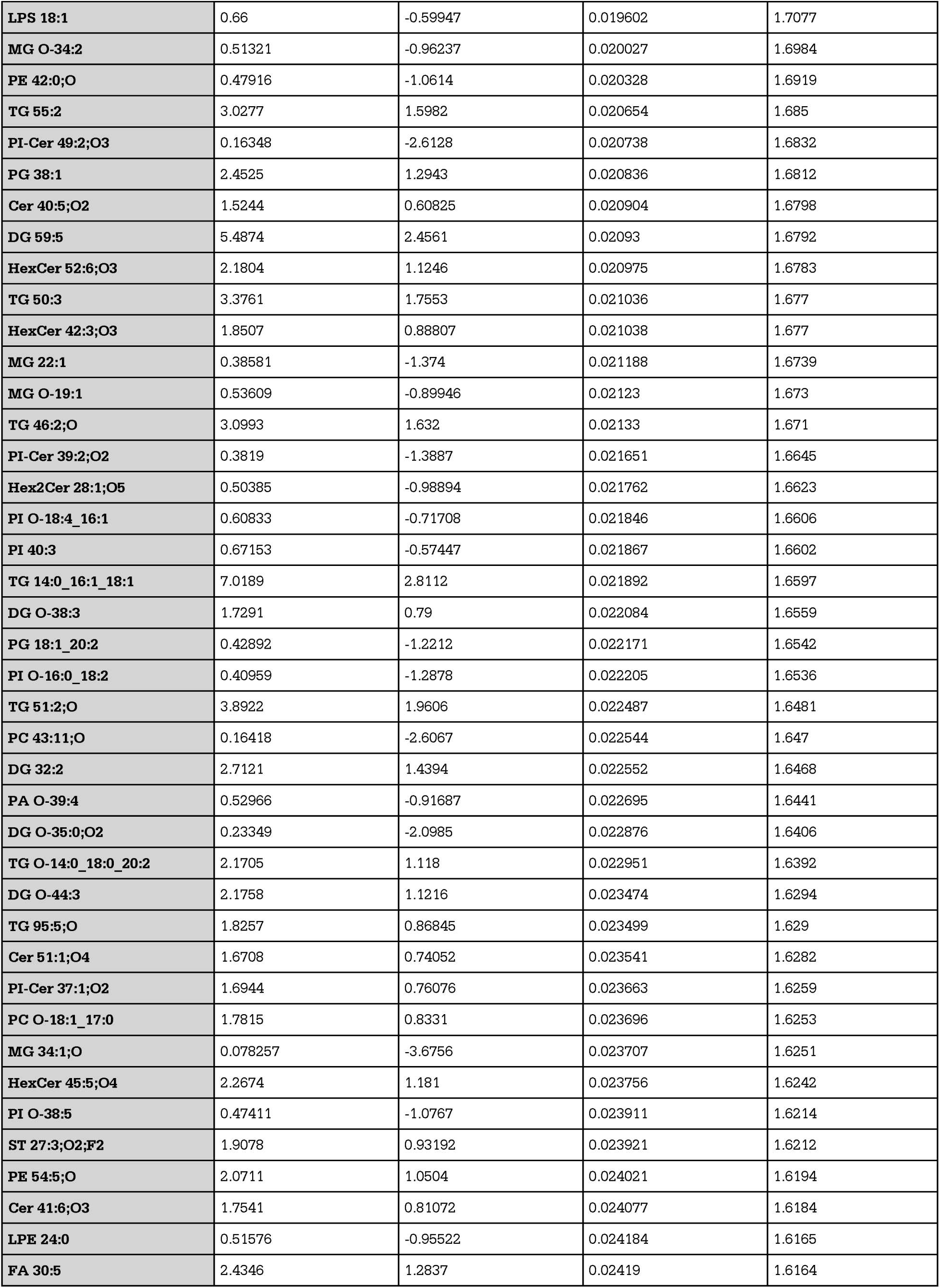
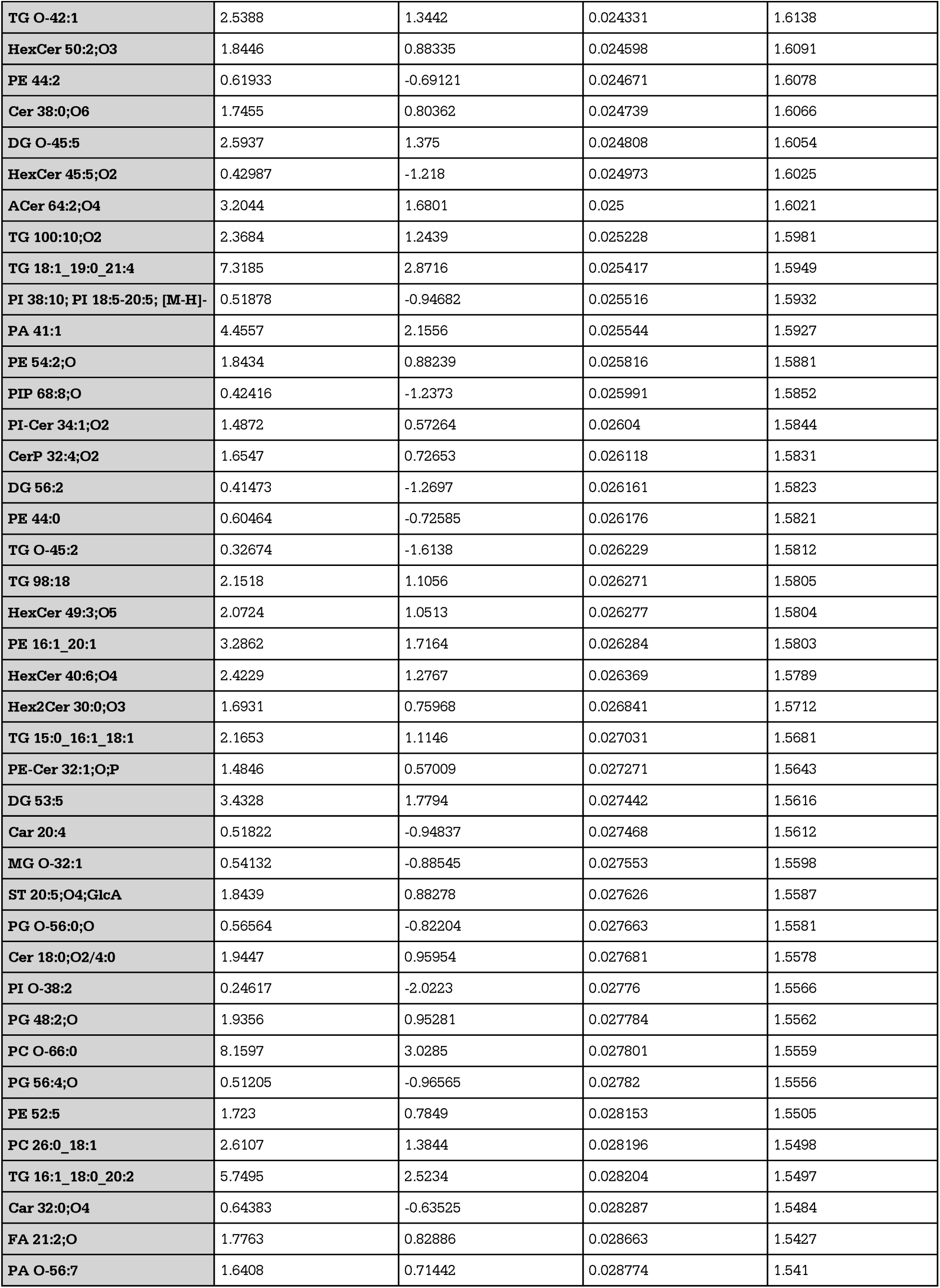
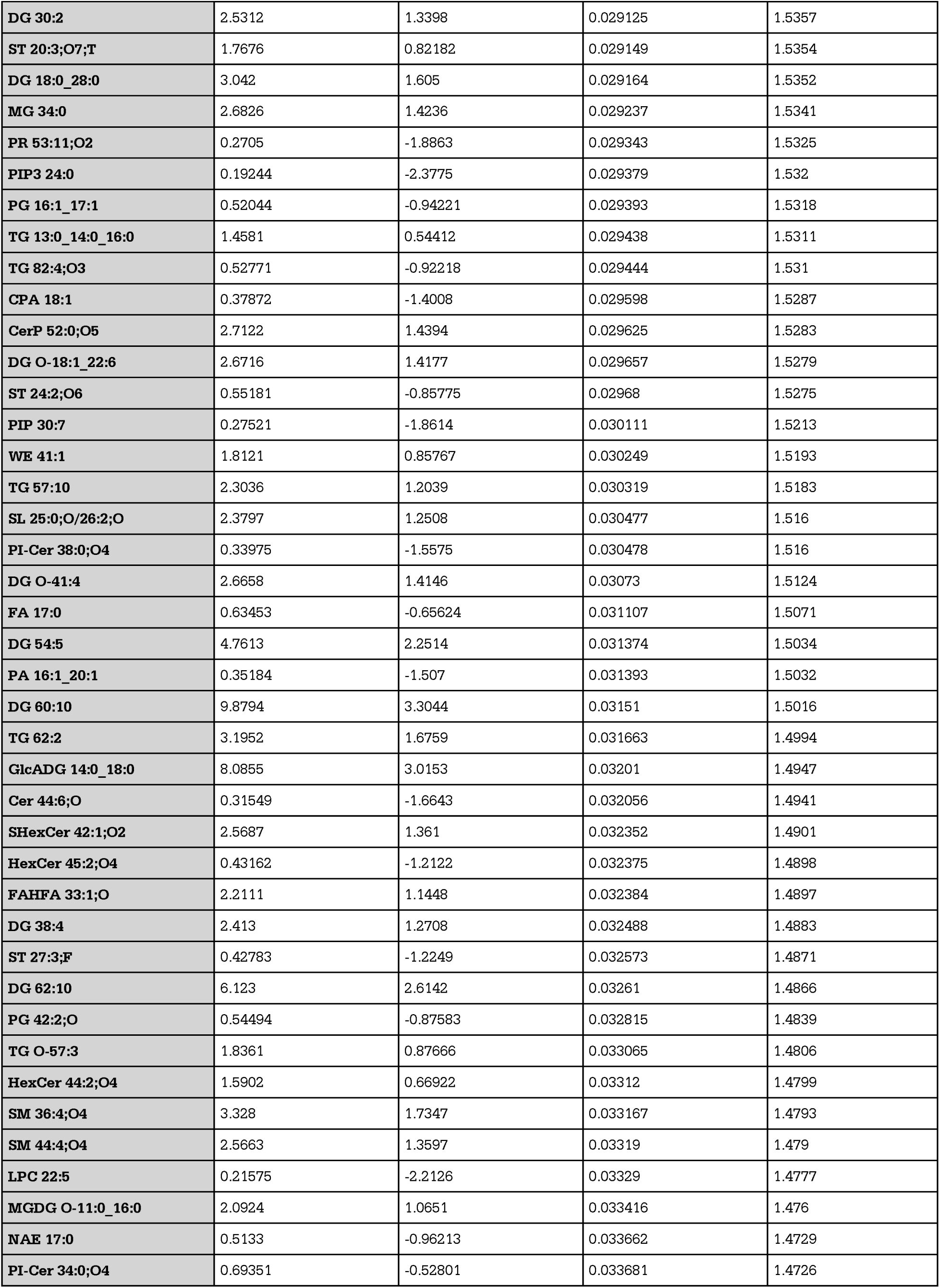
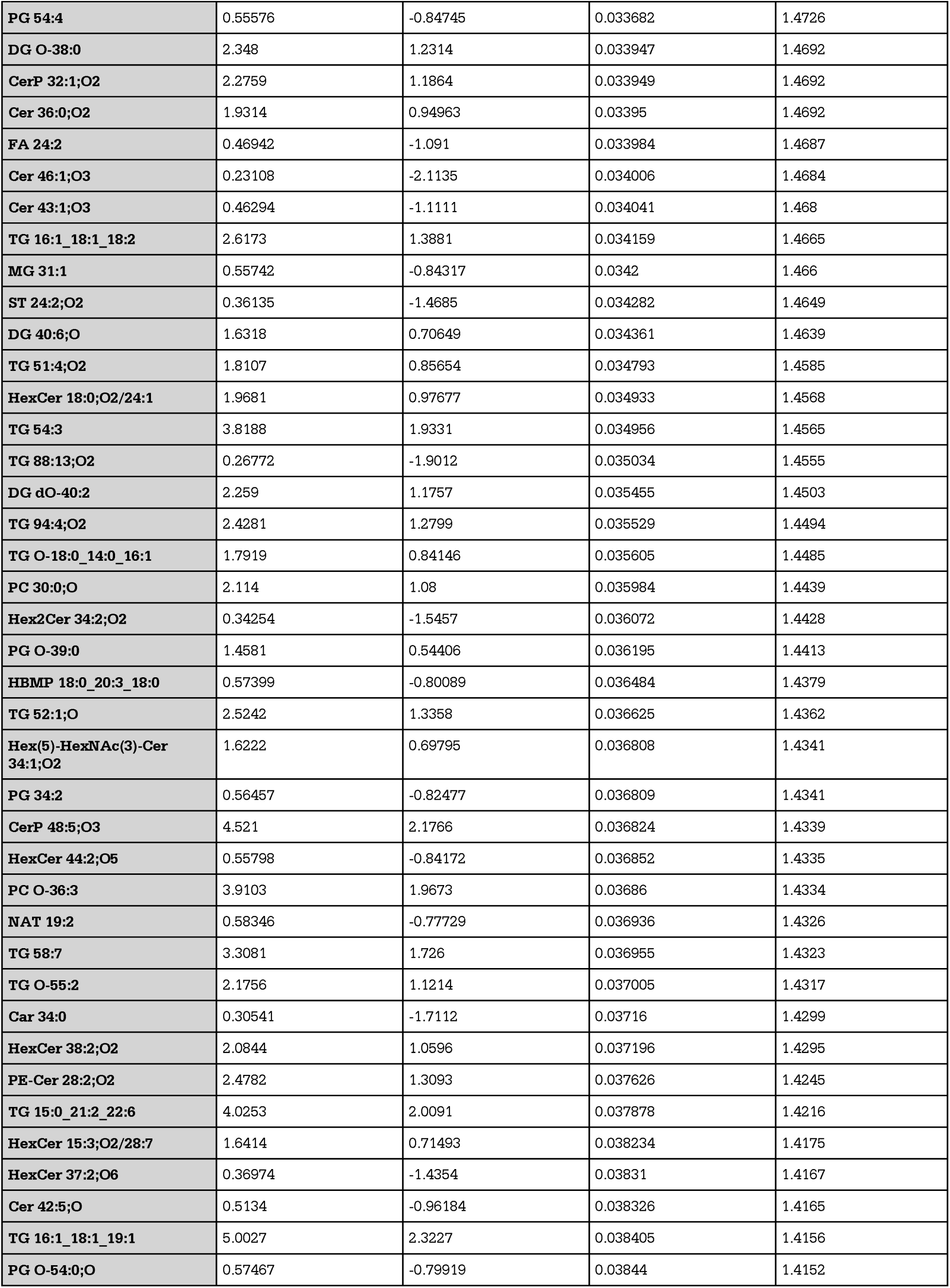
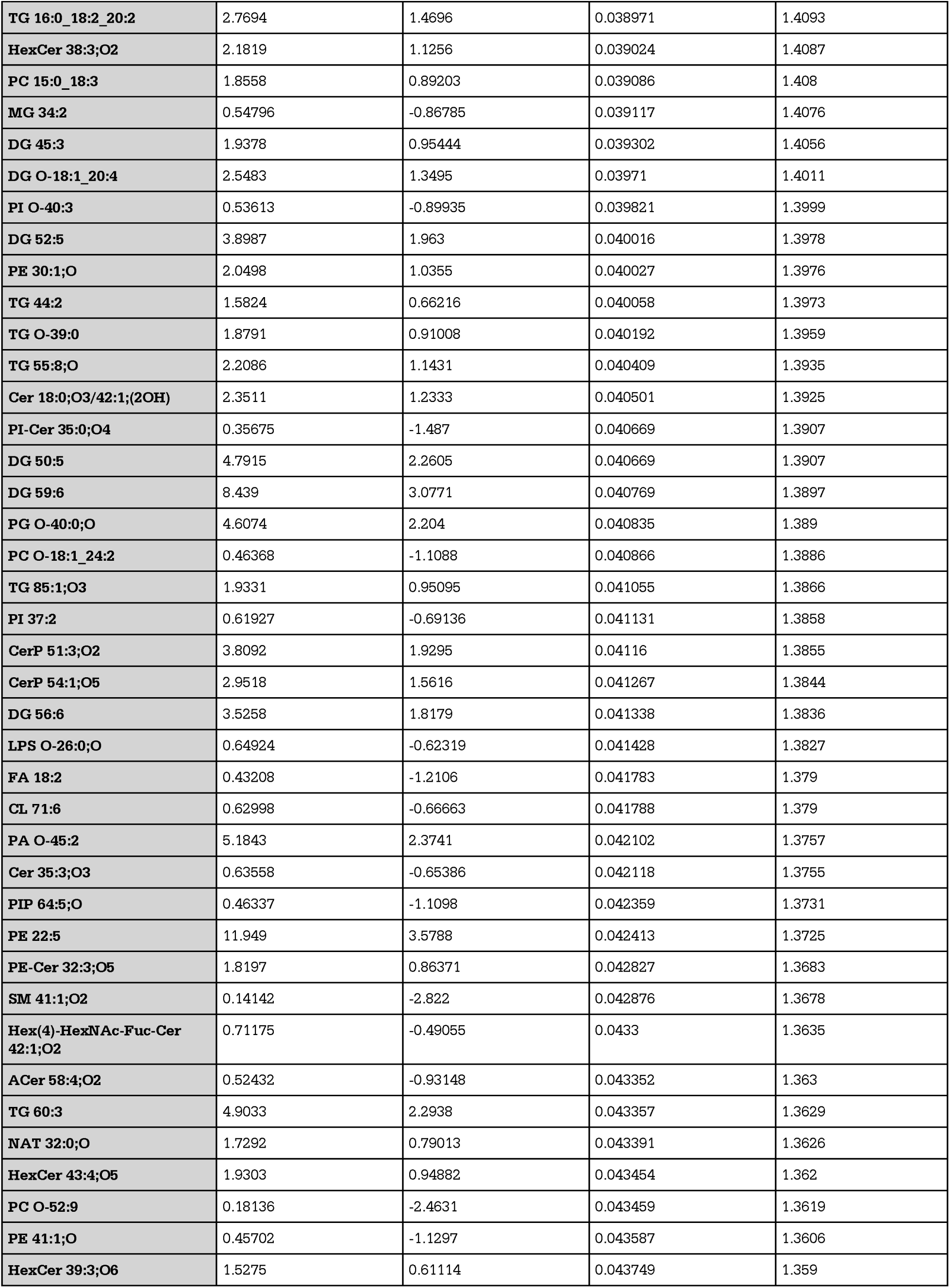
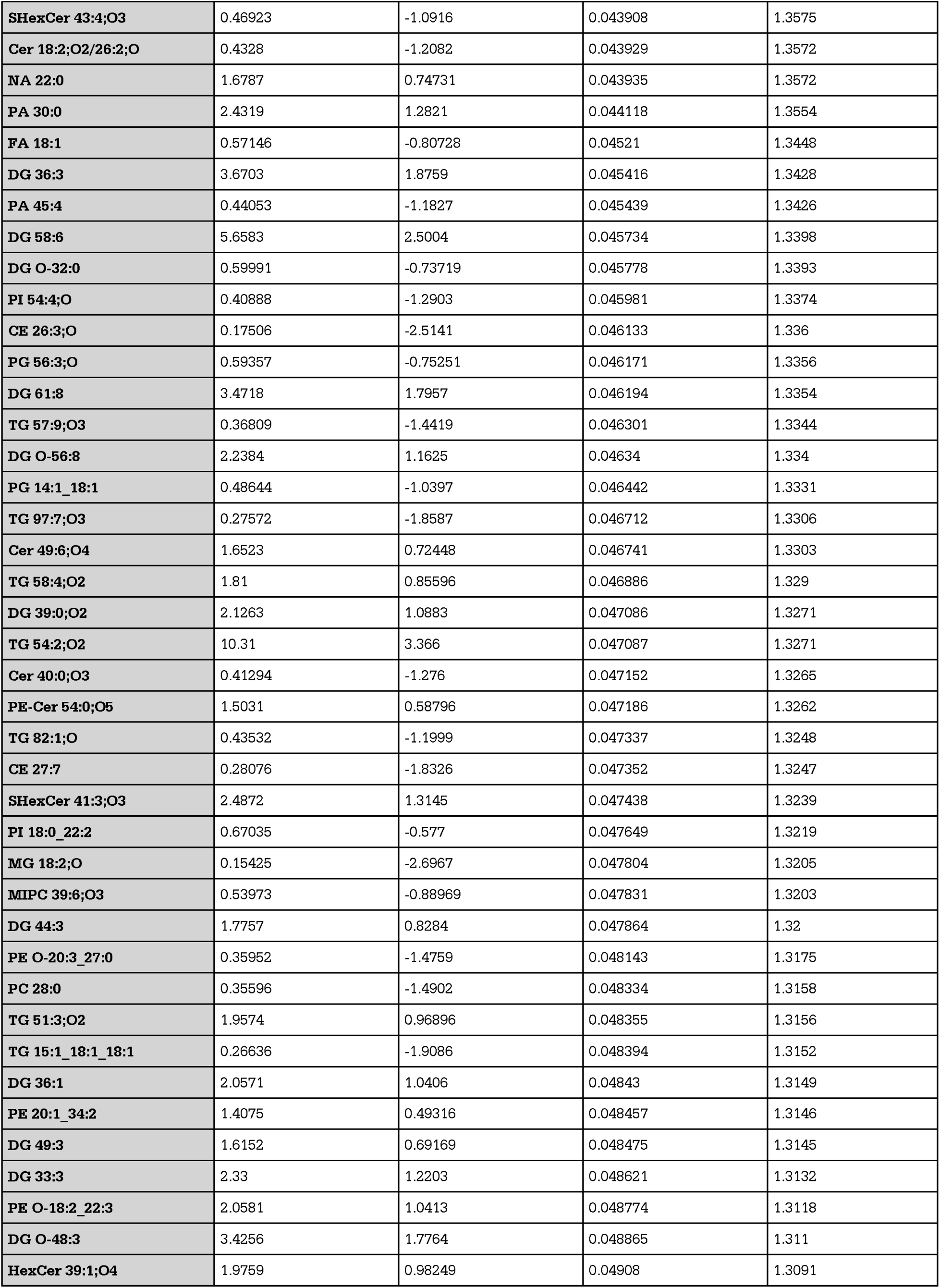
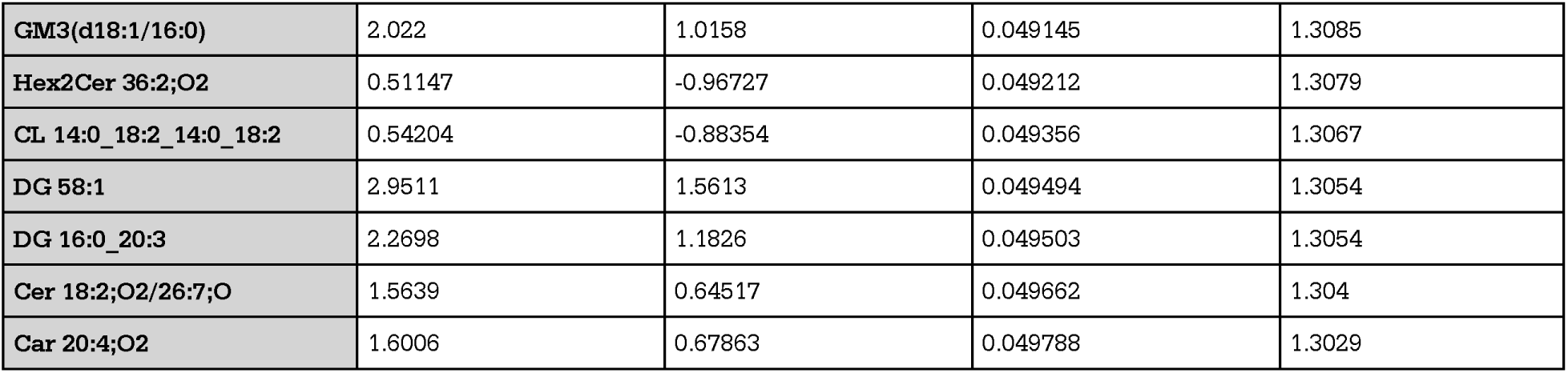
LPS cell:EV and NT cell:EV ratios volcano plot extended statistics. Extended statistics corresponding to Figure 5A in main text. Extended statistics include fold change (FC), log 2 fold change (log2(FC)), the raw p value (raw.pval) and the -log10 of the p- value (-log10(p)).

| Lipid | FC | log2(FC) | raw.pval | -log10(p) |
| --- | --- | --- | --- | --- |
| HexCer 45:1;O4 | 0.11924 | -3.0681 | 7.7921E-06 | 5.1083 |
| LPS 28:0;O | 2.5181 | 1.3323 | 2.1081E-05 | 4.6761 |
| Cer 18:0;O2/22:0;O | 3.4815 | 1.7997 | 3.4878E-05 | 4.4575 |
| TG 48:1 | 26.85 | 4.7469 | 3.4967E-05 | 4.4563 |
| PI 24:0 | 2.0926 | 1.0653 | 4.0489E-05 | 4.3927 |
| FA 28:0;O3 | 0.32316 | -1.6297 | 6.0904E-05 | 4.2154 |
| TG 55:1 | 4.3769 | 2.1299 | 6.8147E-05 | 4.1666 |
| PE 42:2 | 2.6645 | 1.4139 | 7.1387E-05 | 4.1464 |
| LPC 30:4 | 2.6237 | 1.3916 | 9.2011E-05 | 4.0362 |
| PS 37:1 | 7.3855 | 2.8847 | 9.3501E-05 | 4.0292 |
| DG 33:0;O2 | 3.4048 | 1.7676 | 0.00011135 | 3.9533 |
| FA 31:4;O | 0.61029 | -0.71244 | 0.00011166 | 3.9521 |
| PI 15:0_18:1 | 2.4242 | 1.2775 | 0.0001137 | 3.9442 |
| TG 59:1 | 4.3099 | 2.1076 | 0.00012461 | 3.9044 |
| TG 12:2_12:2_22:6 | 4.4916 | 2.1672 | 0.00013568 | 3.8675 |
| TG O-14:0_18:0_18:1 | 4.4255 | 2.1459 | 0.00015554 | 3.8082 |
| MG O-32:0;O | 3.4345 | 1.7801 | 0.00019105 | 3.7188 |
| CerP 52:2;O2 | 3.9041 | 1.965 | 0.00019463 | 3.7108 |
| TG 58:1 | 4.2713 | 2.0947 | 0.00020863 | 3.6806 |
| PE 12:0_16:0 | 1.8987 | 0.92504 | 0.00021838 | 3.6608 |
| Cer 40:5;O3 | 3.4587 | 1.7902 | 0.00022458 | 3.6486 |
| LPG O-32:2 | 0.26941 | -1.8921 | 0.00022475 | 3.6483 |
| PE 58:6 | 2.4606 | 1.299 | 0.00025604 | 3.5917 |
| Cer 16:0;O2/18:0;O | 2.965 | 1.568 | 0.00026609 | 3.575 |
| TG 60:1 | 3.9436 | 1.9795 | 0.00029852 | 3.525 |
| TG 99:5 | 5.6324 | 2.4937 | 0.00038049 | 3.4197 |
| PG 50:2;O | 2.0266 | 1.0191 | 0.00039477 | 3.4037 |
| PI 66:2 | 2.4851 | 1.3133 | 0.00041864 | 3.3782 |
| TG 60:5;O2 | 3.5674 | 1.8349 | 0.00046667 | 3.331 |
| TG 14:0_18:0_22:1 | 3.6732 | 1.877 | 0.0005724 | 3.2423 |
| TG 57:1 | 4.4899 | 2.1667 | 0.00059509 | 3.2254 |
| TG 14:0_18:0_19:1 | 5.1369 | 2.3609 | 0.00064276 | 3.1919 |
| TG 14:0_18:0_20:1 | 6.6017 | 2.7228 | 0.00064742 | 3.1888 |
| PE-NMe2 24:0 | 2.2203 | 1.1508 | 0.00065196 | 3.1858 |
| PE-Cer 18:2;O2/21:1;O | 0.46754 | -1.0969 | 0.00065242 | 3.1855 |
| PG O-44:3 | 4.4987 | 2.1695 | 0.00065592 | 3.1831 |
| DG 39:5;O | 2.6282 | 1.394 | 0.00069279 | 3.1594 |
| PI 36:2 | 0.7111 | -0.49187 | 0.00069633 | 3.1572 |
| PG O-39:0;O | 3.0723 | 1.6193 | 0.00082929 | 3.0813 |
| LPA 26:0 | 0.34241 | -1.5462 | 0.00083262 | 3.0796 |
| TG 12:0_14:0_18:0 | 3.7162 | 1.8938 | 0.00084442 | 3.0734 |
| Cer 28:0;O3 | 0.45106 | -1.1486 | 0.00085436 | 3.0684 |
| LPA O-27:3 | 4.031 | 2.0111 | 0.00096232 | 3.0167 |
| PE 42:2;O | 2.516 | 1.3311 | 0.00098068 | 3.0085 |
| ACer 71:3;O6 | 2.4911 | 1.3168 | 0.0010261 | 2.9888 |
| LPG O-22:0;O | 1.5592 | 0.64078 | 0.0010908 | 2.9622 |
| TG 14:0_18:0_18:1 | 9.8672 | 3.3026 | 0.0011088 | 2.9552 |
| PE 35:0 | 1.4074 | 0.49307 | 0.0011513 | 2.9388 |
| HexCer 28:0;O4 | 2.0137 | 1.0098 | 0.0011911 | 2.9241 |
| DG 55:4 | 6.1693 | 2.6251 | 0.0012846 | 2.8912 |
| PE 30:1 | 0.47233 | -1.0821 | 0.001353 | 2.8687 |
| PI 14:0_16:0 | 2.1842 | 1.1271 | 0.0013567 | 2.8675 |
| TG 56:1 | 10.314 | 3.3666 | 0.0013803 | 2.86 |
| TG 15:0_16:0_17:1 | 10.395 | 3.3778 | 0.0013996 | 2.854 |
| TG 14:0_16:0_19:1 | 6.199 | 2.632 | 0.0014425 | 2.8409 |
| TG 93:14;O | 2.896 | 1.5341 | 0.0014515 | 2.8382 |
| TG 75:5 | 2.5584 | 1.3552 | 0.001492 | 2.8262 |
| PI 31:4;O | 0.45555 | -1.1343 | 0.0014972 | 2.8247 |
| TG 16:1_18:1_18:1 | 7.0668 | 2.8211 | 0.0015127 | 2.8202 |
| DG 34:4 | 4.0152 | 2.0055 | 0.0015157 | 2.8194 |
| PE 40:6 | 0.42453 | -1.2361 | 0.0015259 | 2.8165 |
| TG 12:2_12:3_13:1 | 2.0514 | 1.0366 | 0.0015537 | 2.8086 |
| PA 32:0 | 2.6817 | 1.4231 | 0.0016356 | 2.7863 |
| FA 19:0;O2 | 2.471 | 1.3051 | 0.0016718 | 2.7768 |
| DG 50:2 | 4.6717 | 2.2239 | 0.0017292 | 2.7621 |
| TG O-83:5 | 4.351 | 2.1214 | 0.0017393 | 2.7596 |
| TG 84:11;O3 | 3.1191 | 1.6411 | 0.0017568 | 2.7553 |
| TG 51:3;O3 | 4.2854 | 2.0994 | 0.0017953 | 2.7459 |
| PIP2 62:4 | 0.47804 | -1.0648 | 0.0018355 | 2.7363 |
| Cer 40:1;O6 | 0.3964 | -1.335 | 0.0018478 | 2.7333 |
| FA 20:0;O | 0.43171 | -1.2119 | 0.0018599 | 2.7305 |
| PC 10:0_16:0 | 2.5394 | 1.3445 | 0.0018725 | 2.7276 |
| DG O-37:3 | 3.6874 | 1.8826 | 0.0018828 | 2.7252 |
| HexCer 33:2;O6 | 0.34915 | -1.5181 | 0.0019389 | 2.7124 |
| TG 12:0_16:0_16:1 | 4.575 | 2.1938 | 0.0019821 | 2.7029 |
| TG 79:0;O3 | 3.3751 | 1.7549 | 0.0020045 | 2.698 |
| LPA O-27:6 | 3.3928 | 1.7625 | 0.0021121 | 2.6753 |
| LPA O-27:1 | 1.9544 | 0.96674 | 0.0021176 | 2.6742 |
| TG 15:0_16:0_16:1 | 3.3482 | 1.7434 | 0.0021369 | 2.6702 |
| PI 14:0_18:2 | 0.45667 | -1.1308 | 0.0021565 | 2.6663 |
| DG O-38:4 | 3.9623 | 1.9863 | 0.0022996 | 2.6384 |
| DG 57:6 | 7.8203 | 2.9672 | 0.0023105 | 2.6363 |
| LPS 33:0;O | 3.2161 | 1.6853 | 0.0023135 | 2.6357 |
| Hex2Cer 31:4;O2 | 0.41563 | -1.2666 | 0.0023576 | 2.6275 |
| GlcADG 12:0_13:0 | 2.5597 | 1.356 | 0.002554 | 2.5928 |
| PE 56:5;O | 2.3529 | 1.2344 | 0.0025606 | 2.5917 |
| TG 85:1;O | 2.8136 | 1.4924 | 0.0025642 | 2.5911 |
| TG 49:0 | 4.0918 | 2.0327 | 0.0025658 | 2.5908 |
| HexCer 42:6;O4 | 3.1824 | 1.6701 | 0.0026456 | 2.5775 |
| TG 14:0_14:3_19:5 | 3.7476 | 1.9059 | 0.0026642 | 2.5744 |
| HexCer 42:4;O4 | 2.1814 | 1.1252 | 0.0029245 | 2.534 |
| LPG O-32:0 | 0.24149 | -2.05 | 0.0030299 | 2.5186 |
| PG 20:4;O | 2.0462 | 1.0329 | 0.0030308 | 2.5184 |
| NAE 20:0;O2 | 2.0728 | 1.0516 | 0.0030852 | 2.5107 |
| TG 53:1 | 3.6133 | 1.8533 | 0.0030997 | 2.5087 |
| TG 54:5 | 6.3819 | 2.674 | 0.0031287 | 2.5046 |
| TG 46:3 | 2.4186 | 1.2742 | 0.0031542 | 2.5011 |
| Car 30:1 | 0.52202 | -0.93783 | 0.003239 | 2.4896 |
| PG 27:2;O | 0.56969 | -0.81175 | 0.0032648 | 2.4861 |
| TG 14:0_16:0_16:1 | 7.7132 | 2.9473 | 0.003394 | 2.4693 |
| PG O-38:0 | 5.2994 | 2.4058 | 0.0034234 | 2.4655 |
| TG 12:0_12:2_22:7 | 3.7608 | 1.9111 | 0.0034817 | 2.4582 |
| Hex2Cer 28:0;O4 | 2.0145 | 1.0104 | 0.003594 | 2.4444 |
| DG dO-36:0 | 3.7941 | 1.9237 | 0.003614 | 2.442 |
| MG 21:1 | 0.40087 | -1.3188 | 0.0038048 | 2.4197 |
| HexCer 53:5;O6 | 0.26484 | -1.9168 | 0.0038055 | 2.4196 |
| Hex2Cer 43:0;O2 | 1.5986 | 0.67681 | 0.0038503 | 2.4145 |
| MG 34:2;O | 1.5521 | 0.63419 | 0.0039172 | 2.407 |
| NA 18:0;O | 0.27221 | -1.8772 | 0.0039254 | 2.4061 |
| CerP 53:3;O3 | 3.5735 | 1.8373 | 0.0039638 | 2.4019 |
| PG 40:1 | 0.33755 | -1.5668 | 0.0040101 | 2.3968 |
| HexCer 48:1;O4 | 1.8574 | 0.89332 | 0.0040238 | 2.3954 |
| TG 56:5 | 3.5697 | 1.8358 | 0.0041238 | 2.3847 |
| MG 33:2;O | 1.7254 | 0.78693 | 0.0042237 | 2.3743 |
| TG 56:6 | 5.1969 | 2.3777 | 0.0042431 | 2.3723 |
| TG 16:0_18:0_22:2 | 6.0231 | 2.5905 | 0.0042627 | 2.3703 |
| ST 27:4;O | 0.23469 | -2.0912 | 0.0042892 | 2.3676 |
| PC 12:0_16:1 | 1.9484 | 0.96229 | 0.004348 | 2.3617 |
| DG O-16:0_18:1 | 5.19 | 2.3757 | 0.0043871 | 2.3578 |
| DG 16:0_20:0 | 8.522 | 3.0912 | 0.0044468 | 2.3519 |
| MG 19:0;O | 2.6391 | 1.4 | 0.0045361 | 2.3433 |
| DG 27:0_21:1 | 6.7631 | 2.7577 | 0.0046117 | 2.3361 |
| LPI 18:1 | 0.53917 | -0.89119 | 0.004691 | 2.3287 |
| DG 58:4 | 4.3207 | 2.1113 | 0.0049176 | 2.3082 |
| PE 56:5 | 2.7107 | 1.4387 | 0.0049401 | 2.3063 |
| DG 47:1 | 0.44575 | -1.1657 | 0.0049605 | 2.3045 |
| TG 16:0_18:1_20:1 | 9.0825 | 3.1831 | 0.0049791 | 2.3029 |
| DG 28:1 | 7.2155 | 2.8511 | 0.0049957 | 2.3014 |
| DG 53:3 | 5.5797 | 2.4802 | 0.0051306 | 2.2898 |
| PE 25:1;O | 1.5924 | 0.67121 | 0.005159 | 2.2874 |
| SHexCer 38:1;O3 | 0.58526 | -0.77285 | 0.005177 | 2.2859 |
| TG 52:2;O | 3.3373 | 1.7387 | 0.0052079 | 2.2833 |
| PC O-22:3_24:0 | 2.0662 | 1.0469 | 0.0053226 | 2.2739 |
| TG 14:0_18:0_18:2 | 7.5694 | 2.9202 | 0.0055347 | 2.2569 |
| HexCer 45:4;O4 | 0.65849 | -0.60276 | 0.0056009 | 2.2517 |
| TG 54:2 | 3.6738 | 1.8773 | 0.0056898 | 2.2449 |
| DG 18:1_34:1 | 3.1053 | 1.6347 | 0.0056981 | 2.2443 |
| HexCer 43:2;O5 | 1.5391 | 0.62207 | 0.0057356 | 2.2414 |
| PG 58:4;O | 0.2249 | -2.1526 | 0.005819 | 2.2351 |
| TG O-97:5 | 3.3257 | 1.7336 | 0.0058398 | 2.2336 |
| PI 64:1 | 1.9252 | 0.94503 | 0.0059049 | 2.2288 |
| HexCer 54:6;O3 | 2.1376 | 1.096 | 0.0059105 | 2.2284 |
| TG 45:2 | 6.7104 | 2.7464 | 0.0059254 | 2.2273 |
| TG O-75:0;O3 | 2.2409 | 1.1641 | 0.0059756 | 2.2236 |
| TG 81:1;O3 | 1.8185 | 0.86277 | 0.0059772 | 2.2235 |
| SOMG 18:0 | 0.37351 | -1.4208 | 0.0060727 | 2.2166 |
| TG 16:1_18:0_18:1 | 8.4496 | 3.0789 | 0.0060778 | 2.2163 |
| MG O-18:0 | 2.4831 | 1.3121 | 0.0060994 | 2.2147 |
| TG 16:0_18:1_18:1 | 5.1667 | 2.3692 | 0.0061174 | 2.2134 |
| Cer 41:4;O | 2.3785 | 1.2501 | 0.006264 | 2.2031 |
| HBMP 16:1_18:0_18:0 | 2.1226 | 1.0858 | 0.006273 | 2.2025 |
| DG 28:4;O2 | 2.6115 | 1.3849 | 0.0063897 | 2.1945 |
| ST 25:1;O;Hex | 0.60735 | -0.71941 | 0.0064439 | 2.1909 |
| PI O-67:3 | 2.462 | 1.2998 | 0.0064818 | 2.1883 |
| Cer 36:4;O4 | 0.46969 | -1.0902 | 0.0064918 | 2.1876 |
| DG 41:6;O | 1.7446 | 0.8029 | 0.0065183 | 2.1859 |
| PG 14:0_18:0 | 2.6379 | 1.3994 | 0.0066203 | 2.1791 |
| Cer 40:2;O3 | 2.4367 | 1.2849 | 0.0066654 | 2.1762 |
| Hex2Cer 30:4;O4 | 2.0205 | 1.0147 | 0.0066836 | 2.175 |
| TG 84:5;O3 | 0.29738 | -1.7496 | 0.0067083 | 2.1734 |
| PE 40:1;O | 0.37563 | -1.4126 | 0.0068465 | 2.1645 |
| Cer 43:5;O2 | 0.57922 | -0.78782 | 0.0069241 | 2.1596 |
| **Cer 18:3;O2/24:6** | 1.4366 | 0.52263 | 0.0069448 | 2.1583 |
| **LPG 18:1** | 0.40511 | -1.3036 | 0.0069525 | 2.1579 |
| **TG 58:2** | 8.179 | 3.0319 | 0.0069877 | 2.1557 |
| **SHexCer 29:0;O2** | 1.5587 | 0.64036 | 0.0070548 | 2.1515 |
| **HexCer 43:2;O4** | 0.44 | -1.1844 | 0.0070675 | 2.1507 |
| **HexCer 43:0;O4** | 2.3424 | 1.228 | 0.0071358 | 2.1466 |
| **TG 96:8;O** | 2.1534 | 1.1066 | 0.0072817 | 2.1378 |
| **TG 59:2** | 6.7912 | 2.7637 | 0.0073125 | 2.1359 |
| **PE O-40:1** | 0.53741 | -0.8959 | 0.0073197 | 2.1355 |
| **TG 44:6;O2** | 5.3443 | 2.418 | 0.0074636 | 2.1271 |
| **PC 15:0_18:5** | 0.30616 | -1.7076 | 0.0075658 | 2.1211 |
| **Cer 39:0;O3** | 2.202 | 1.1388 | 0.0076993 | 2.1135 |
| **DG 50:4** | 9.4286 | 3.237 | 0.0077061 | 2.1132 |
| **CerP 45:6;O2** | 1.5954 | 0.67393 | 0.007952 | 2.0995 |
| **TG 52:7;O2** | 0.37547 | -1.4132 | 0.0080012 | 2.0968 |
| **FA 17:0;O2** | 1.8972 | 0.92386 | 0.0080211 | 2.0958 |
| **DG 58:5** | 8.9961 | 3.1693 | 0.0081967 | 2.0864 |
| **PC O-18:1_16:0** | 2.0939 | 1.0662 | 0.0082286 | 2.0847 |
| **DG O-14:1_18:0** | 3.0482 | 1.608 | 0.0082514 | 2.0835 |
| **TG 79:0;O2** | 4.5455 | 2.1844 | 0.0082555 | 2.0833 |
| **Cer 48:6;O2** | 2.0955 | 1.0673 | 0.0083482 | 2.0784 |
| **HexCer 32:5;O5** | 0.62541 | -0.67714 | 0.0085639 | 2.0673 |
| **TG 16:0_18:1_22:4** | 5.131 | 2.3592 | 0.0085907 | 2.066 |
| **PC O-18:1_22:2** | 0.41586 | -1.2658 | 0.0086015 | 2.0654 |
| **FOH 10:2** | 2.7707 | 1.4703 | 0.0086423 | 2.0634 |
| **PI-Cer 35:0;O2** | 1.6444 | 0.71758 | 0.0086869 | 2.0611 |
| **Cer 37:6;O2** | 3.2051 | 1.6804 | 0.0087307 | 2.059 |
| **TG 62:10** | 2.264 | 1.1789 | 0.0087515 | 2.0579 |
| **Cer 15:2;2O/38:10** | 6.9459 | 2.7962 | 0.0087526 | 2.0579 |
| **DG O-43:3** | 3.4954 | 1.8055 | 0.0087602 | 2.0575 |
| **Cer 34:1;O4** | 0.57613 | -0.79555 | 0.0089842 | 2.0465 |
| **PC 17:0_18:0** | 1.4167 | 0.50249 | 0.0091108 | 2.0404 |
| **NAT 26:2;O3** | 0.64978 | -0.62197 | 0.0091148 | 2.0403 |
| **DG 31:2** | 3.3694 | 1.7525 | 0.0093003 | 2.0315 |
| **DG 14:0_16:0** | 2.8389 | 1.5053 | 0.0094207 | 2.0259 |
| **Car 22:4;O4** | 3.0423 | 1.6052 | 0.0094342 | 2.0253 |
| **Cer 44:0;O** | 0.44713 | -1.1612 | 0.0096138 | 2.0171 |
| **Hex2Cer 44:6;O2** | 0.36108 | -1.4696 | 0.0096682 | 2.0147 |
| **DG O-35:3** | 3.0361 | 1.6022 | 0.0097095 | 2.0128 |
| **Cer 18:0;O3/35:1;(2OH)** | 5.566 | 2.4766 | 0.009857 | 2.0063 |
| **DG 32:0_22:2** | 4.6918 | 2.2301 | 0.009877 | 2.0054 |
| **TG 51:3;O** | 3.9492 | 1.9816 | 0.0098935 | 2.0047 |
| **ACer 58:0;O3** | 3.0727 | 1.6195 | 0.0099405 | 2.0026 |
| **DG 51:4** | 4.3629 | 2.1253 | 0.010258 | 1.9889 |
| **DG 26:3;O** | 1.8721 | 0.90462 | 0.010329 | 1.9859 |
| **SPBP 16:1;O3** | 2.2323 | 1.1585 | 0.01033 | 1.9859 |
| **TG 56:4** | 2.2177 | 1.1491 | 0.01036 | 1.9847 |
| **DG O-56:5** | 6.6587 | 2.7352 | 0.010471 | 1.98 |
| **PR 20:2;O3** | 0.30508 | -1.7128 | 0.010498 | 1.9789 |
| **DG 44:8;O** | 1.6729 | 0.74239 | 0.010579 | 1.9756 |
| **DG 57:4** | 8.6499 | 3.1127 | 0.010696 | 1.9708 |
| **DG 38:6;O** | 2.3955 | 1.2603 | 0.010841 | 1.9649 |
| **TG 16:0_19:3_19:3** | 6.7485 | 2.7546 | 0.010981 | 1.9594 |
| **TG 52:5** | 3.8805 | 1.9562 | 0.010985 | 1.9592 |
| **DG 37:0;O2** | 2.6802 | 1.4224 | 0.011096 | 1.9548 |
| **DG 40:1;O2** | 2.7856 | 1.478 | 0.011129 | 1.9535 |
| **Cer 41:4;O2** | 2.0775 | 1.0548 | 0.011169 | 1.952 |
| **DG 15:0_18:0** | 3.7718 | 1.9153 | 0.011221 | 1.95 |
| **TG 42:1** | 2.1747 | 1.1208 | 0.011301 | 1.9469 |
| **LPE 22:1** | 1.8502 | 0.88767 | 0.011607 | 1.9353 |
| **NAE 19:0;O** | 0.41381 | -1.273 | 0.011789 | 1.9285 |
| **PE 36:3** | 3.5721 | 1.8368 | 0.011825 | 1.9272 |
| **TG 14:0_18:1_19:1** | 6.0838 | 2.605 | 0.011964 | 1.9221 |
| **Cer 38:0;O** | 2.2739 | 1.1852 | 0.011965 | 1.9221 |
| **LPG O-27:0;O** | 1.7063 | 0.77084 | 0.012073 | 1.9182 |
| **PC O-18:1_20:2** | 0.47992 | -1.0591 | 0.01214 | 1.9158 |
| **TG 97:6;O2** | 2.6304 | 1.3953 | 0.012251 | 1.9118 |
| **TG 16:0_18:1_22:6** | 8.1719 | 3.0307 | 0.012354 | 1.9082 |
| **PE 30:0** | 0.62426 | -0.67979 | 0.012412 | 1.9062 |
| **DG 56:1** | 4.3042 | 2.1058 | 0.012461 | 1.9044 |
| **PI 40:1;O** | 0.1175 | -3.0893 | 0.012627 | 1.8987 |
| **TG 42:0** | 2.9326 | 1.5522 | 0.012785 | 1.8933 |
| **DG 39:6;O** | 1.7712 | 0.82473 | 0.012799 | 1.8928 |
| **TG 12:1_12:3_18:5** | 2.1037 | 1.073 | 0.012913 | 1.889 |
| **PR 58:14;O3** | 3.8026 | 1.927 | 0.012976 | 1.8869 |
| **TG 12:3_12:3_18:4** | 3.0809 | 1.6233 | 0.012987 | 1.8865 |
| **DG 56:8** | 5.5252 | 2.466 | 0.01324 | 1.8781 |
| **CerP 54:6;O5** | 2.5603 | 1.3563 | 0.013268 | 1.8772 |
| **PE 40:2** | 1.97 | 0.9782 | 0.013274 | 1.877 |
| **TG 16:0_18:1_18:2** | 5.5546 | 2.4737 | 0.013373 | 1.8738 |
| **DG O-16:0_18:0** | 2.8039 | 1.4874 | 0.013423 | 1.8722 |
| **LPA 26:6** | 1.4832 | 0.56869 | 0.013458 | 1.871 |
| **TG 60:6** | 2.6193 | 1.3892 | 0.013462 | 1.8709 |
| **TG 83:0;O3** | 2.5772 | 1.3658 | 0.013516 | 1.8692 |
| **TG 91:13;O** | 2.0008 | 1.0006 | 0.013724 | 1.8625 |
| **TG 51:1** | 8.4973 | 3.087 | 0.013748 | 1.8617 |
| **DG 53:4** | 4.334 | 2.1157 | 0.013815 | 1.8596 |
| **TG 86:13** | 1.9759 | 0.9825 | 0.013832 | 1.8591 |
| **TG O-61:0;O3** | 0.47972 | -1.0597 | 0.013902 | 1.8569 |
| **LPE 24:1** | 0.35462 | -1.4956 | 0.014087 | 1.8512 |
| **TG 70:3;O3** | 0.36267 | -1.4633 | 0.014172 | 1.8486 |
| **PG 18:1_20:1** | 0.36345 | -1.4602 | 0.01421 | 1.8474 |
| **TG 48:2** | 8.6679 | 3.1157 | 0.014212 | 1.8473 |
| **ST 29:7;O** | 3.7867 | 1.9209 | 0.014285 | 1.8451 |
| **FA 33:3;O2** | 2.7862 | 1.4783 | 0.014293 | 1.8449 |
| **TG 46:2** | 6.4249 | 2.6837 | 0.014311 | 1.8443 |
| **MG 32:1** | 0.52021 | -0.94284 | 0.014314 | 1.8443 |
| **TG 70:12;O2** | 0.18527 | -2.4323 | 0.014362 | 1.8428 |
| **PG 14:0_16:0** | 2.6772 | 1.4207 | 0.01437 | 1.8425 |
| **Cer 42:1;O3** | 0.60789 | -0.71813 | 0.014406 | 1.8415 |
| **DG 60:6** | 6.2449 | 2.6427 | 0.01467 | 1.8336 |
| **TG 17:1_18:0_18:1** | 6.243 | 2.6422 | 0.014705 | 1.8325 |
| **FA 17:1** | 0.45787 | -1.127 | 0.014728 | 1.8319 |
| **MG O-20:1** | 0.34753 | -1.5248 | 0.014919 | 1.8263 |
| **TG 60:2** | 6.1766 | 2.6268 | 0.014939 | 1.8257 |
| **Cer 49:1;O4** | 2.0619 | 1.044 | 0.014949 | 1.8254 |
| **PC O-22:3_26:0** | 1.8697 | 0.90282 | 0.015015 | 1.8235 |
| **DG 39:2** | 4.0968 | 2.0345 | 0.01518 | 1.8187 |
| **TG 100:19** | 2.3811 | 1.2516 | 0.015388 | 1.8128 |
| **TG 84:6;O2** | 1.8454 | 0.88397 | 0.015735 | 1.8031 |
| **PG 44:9** | 0.45465 | -1.1372 | 0.015995 | 1.796 |
| **DG 26:0** | 5.2848 | 2.4019 | 0.016109 | 1.7929 |
| **Cer 40:0;O6** | 0.65824 | -0.60332 | 0.016119 | 1.7927 |
| **PE-Cer 42:6;O3** | 2.135 | 1.0942 | 0.016181 | 1.791 |
| **HexCer 35:2;O5** | 6.1874 | 2.6293 | 0.01619 | 1.7908 |
| **TG 57:9** | 1.8399 | 0.87964 | 0.016196 | 1.7906 |
| **SHexCer 34:0;O4** | 2.054 | 1.0385 | 0.016199 | 1.7905 |
| **HexCer 48:2;O4** | 1.844 | 0.8828 | 0.01634 | 1.7868 |
| **PE 46:1** | 3.6571 | 1.8707 | 0.016501 | 1.7825 |
| **PI-Cer 46:0;O3** | 0.4716 | -1.0844 | 0.016545 | 1.7813 |
| **CL 74:4** | 0.59545 | -0.74795 | 0.016559 | 1.781 |
| **PE O-40:6** | 0.36181 | -1.4667 | 0.016746 | 1.7761 |
| **TG 14:0_18:1_18:2** | 3.8018 | 1.9267 | 0.016846 | 1.7735 |
| **FAHFA 14:0/14:0** | 0.50869 | -0.97515 | 0.016977 | 1.7701 |
| **DG 32:0** | 4.1929 | 2.068 | 0.017048 | 1.7683 |
| **TG 54:7** | 4.0353 | 2.0127 | 0.017095 | 1.7671 |
| **DG 44:9** | 1.8601 | 0.89538 | 0.0171 | 1.767 |
| **MG 33:1** | 0.63335 | -0.65894 | 0.017131 | 1.7662 |
| **TG 98:9;O2** | 2.6571 | 1.4098 | 0.017152 | 1.7657 |
| **Cer 50:2;O3** | 1.7171 | 0.78 | 0.017183 | 1.7649 |
| **FA 20:0** | 0.60949 | -0.71432 | 0.017285 | 1.7623 |
| **DG 52:4** | 5.5725 | 2.4783 | 0.017306 | 1.7618 |
| **Cer 40:6;O3** | 2.6288 | 1.3944 | 0.01743 | 1.7587 |
| **PI 36:5** | 0.30048 | -1.7346 | 0.017564 | 1.7554 |
| **TG 57:2** | 8.3932 | 3.0692 | 0.01758 | 1.755 |
| **MGDG 15:0_18:1** | 3.7188 | 1.8949 | 0.017594 | 1.7546 |
| **DG 55:5** | 7.4686 | 2.9008 | 0.017701 | 1.752 |
| **DG 36:2** | 2.6533 | 1.4078 | 0.017758 | 1.7506 |
| **TG 75:0;O3** | 2.4367 | 1.2849 | 0.017772 | 1.7503 |
| **CE 16:0** | 1.8415 | 0.88084 | 0.017891 | 1.7474 |
| **PE O-38:2** | 2.4201 | 1.2751 | 0.01792 | 1.7467 |
| **HexCer 32:6;O3** | 1.711 | 0.77488 | 0.018004 | 1.7446 |
| **HBMP 60:2; HBMP 20:0-22:1-18:1; [M+NH4]<sup>+</sup>** | 0.14448 | -2.7911 | 0.018092 | 1.7425 |
| **PE 18:1_26:4** | 0.55769 | -0.84245 | 0.018201 | 1.7399 |
| **TG 77:1;O3** | 1.8278 | 0.87009 | 0.018212 | 1.7396 |
| **DG 60:5** | 3.8823 | 1.9569 | 0.018439 | 1.7343 |
| **PC O-22:3_26:1** | 2.271 | 1.1833 | 0.018483 | 1.7332 |
| **LPG O-33:4** | 0.38818 | -1.3652 | 0.018503 | 1.7328 |
| **TG 50:0;O** | 3.11 | 1.6369 | 0.018585 | 1.7308 |
| **TG 12:0_14:0_16:0** | 2.9155 | 1.5437 | 0.018659 | 1.7291 |
| **DG 40:0** | 2.3694 | 1.2445 | 0.018659 | 1.7291 |
| **LPG O-31:5** | 0.30133 | -1.7306 | 0.018663 | 1.729 |
| **DG 46:3** | 3.7911 | 1.9226 | 0.01867 | 1.7289 |
| **CE 26:2** | 0.27981 | -1.8375 | 0.018674 | 1.7288 |
| **PA 26:0** | 0.36112 | -1.4694 | 0.018881 | 1.724 |
| **DG 16:0_32:2** | 4.2679 | 2.0935 | 0.01892 | 1.7231 |
| **DG O-39:3** | 2.2278 | 1.1556 | 0.018954 | 1.7223 |
| **HexCer 45:0;O5** | 2.2031 | 1.1395 | 0.019001 | 1.7212 |
| **PG O-44:8** | 1.4956 | 0.58071 | 0.019046 | 1.7202 |
| **TG 14:0_16:0_22:6** | 4.5122 | 2.1738 | 0.019164 | 1.7175 |
| **TG 16:1_18:1_22:1** | 5.7699 | 2.5285 | 0.019419 | 1.7118 |
| **CL 74:6** | 0.69683 | -0.52112 | 0.019484 | 1.7103 |
| **PE 44:3;O** | 0.43082 | -1.2148 | 0.019503 | 1.7099 |
| LPS 18:1 | 0.66 | -0.59947 | 0.019602 | 1.7077 |
| MG O-34:2 | 0.51321 | -0.96237 | 0.020027 | 1.6984 |
| PE 42:0;O | 0.47916 | -1.0614 | 0.020328 | 1.6919 |
| TG 55:2 | 3.0277 | 1.5982 | 0.020654 | 1.685 |
| PI-Cer 49:2;O3 | 0.16348 | -2.6128 | 0.020738 | 1.6832 |
| PG 38:1 | 2.4525 | 1.2943 | 0.020836 | 1.6812 |
| Cer 40:5;O2 | 1.5244 | 0.60825 | 0.020904 | 1.6798 |
| DG 59:5 | 5.4874 | 2.4561 | 0.02093 | 1.6792 |
| HexCer 52:6;O3 | 2.1804 | 1.1246 | 0.020975 | 1.6783 |
| TG 50:3 | 3.3761 | 1.7553 | 0.021036 | 1.677 |
| HexCer 42:3;O3 | 1.8507 | 0.88807 | 0.021038 | 1.677 |
| MG 22:1 | 0.38581 | -1.374 | 0.021188 | 1.6739 |
| MG O-19:1 | 0.53609 | -0.89946 | 0.02123 | 1.673 |
| TG 46:2;O | 3.0993 | 1.632 | 0.02133 | 1.671 |
| PI-Cer 39:2;O2 | 0.3819 | -1.3887 | 0.021651 | 1.6645 |
| Hex2Cer 28:1;O5 | 0.50385 | -0.98894 | 0.021762 | 1.6623 |
| PI O-18:4_16:1 | 0.60833 | -0.71708 | 0.021846 | 1.6606 |
| PI 40:3 | 0.67153 | -0.57447 | 0.021867 | 1.6602 |
| TG 14:0_16:1_18:1 | 7.0189 | 2.8112 | 0.021892 | 1.6597 |
| DG O-38:3 | 1.7291 | 0.79 | 0.022084 | 1.6559 |
| PG 18:1_20:2 | 0.42892 | -1.2212 | 0.022171 | 1.6542 |
| PI O-16:0_18:2 | 0.40959 | -1.2878 | 0.022205 | 1.6536 |
| TG 51:2;O | 3.8922 | 1.9606 | 0.022487 | 1.6481 |
| PC 43:11;O | 0.16418 | -2.6067 | 0.022544 | 1.647 |
| DG 32:2 | 2.7121 | 1.4394 | 0.022552 | 1.6468 |
| PA O-39:4 | 0.52966 | -0.91687 | 0.022695 | 1.6441 |
| DG O-35:0;O2 | 0.23349 | -2.0985 | 0.022876 | 1.6406 |
| TG O-14:0_18:0_20:2 | 2.1705 | 1.118 | 0.022951 | 1.6392 |
| DG O-44:3 | 2.1758 | 1.1216 | 0.023474 | 1.6294 |
| TG 95:5;O | 1.8257 | 0.86845 | 0.023499 | 1.629 |
| Cer 51:1;O4 | 1.6708 | 0.74052 | 0.023541 | 1.6282 |
| PI-Cer 37:1;O2 | 1.6944 | 0.76076 | 0.023663 | 1.6259 |
| PC O-18:1_17:0 | 1.7815 | 0.8331 | 0.023696 | 1.6253 |
| MG 34:1;O | 0.078257 | -3.6756 | 0.023707 | 1.6251 |
| HexCer 45:5;O4 | 2.2674 | 1.181 | 0.023756 | 1.6242 |
| PI O-38:5 | 0.47411 | -1.0767 | 0.023911 | 1.6214 |
| ST 27:3;O2;F2 | 1.9078 | 0.93192 | 0.023921 | 1.6212 |
| PE 54:5;O | 2.0711 | 1.0504 | 0.024021 | 1.6194 |
| Cer 41:6;O3 | 1.7541 | 0.81072 | 0.024077 | 1.6184 |
| LPE 24:0 | 0.51576 | -0.95522 | 0.024184 | 1.6165 |
| FA 30:5 | 2.4346 | 1.2837 | 0.02419 | 1.6164 |
| TG O-42:1 | 2.5388 | 1.3442 | 0.024331 | 1.6138 |
| HexCer 50:2;O3 | 1.8446 | 0.88335 | 0.024598 | 1.6091 |
| PE 44:2 | 0.61933 | -0.69121 | 0.024671 | 1.6078 |
| Cer 38:0;O6 | 1.7455 | 0.80362 | 0.024739 | 1.6066 |
| DG O-45:5 | 2.5937 | 1.375 | 0.024808 | 1.6054 |
| HexCer 45:5;O2 | 0.42987 | -1.218 | 0.024973 | 1.6025 |
| ACer 64:2;O4 | 3.2044 | 1.6801 | 0.025 | 1.6021 |
| TG 100:10;O2 | 2.3684 | 1.2439 | 0.025228 | 1.5981 |
| TG 18:1_19:0_21:4 | 7.3185 | 2.8716 | 0.025417 | 1.5949 |
| PI 38:10; PI 18:5-20:5; [M-H]- | 0.51878 | -0.94682 | 0.025516 | 1.5932 |
| PA 41:1 | 4.4557 | 2.1556 | 0.025544 | 1.5927 |
| PE 54:2;O | 1.8434 | 0.88239 | 0.025816 | 1.5881 |
| PIP 68:8;O | 0.42416 | -1.2373 | 0.025991 | 1.5852 |
| PI-Cer 34:1;O2 | 1.4872 | 0.57264 | 0.02604 | 1.5844 |
| CerP 32:4;O2 | 1.6547 | 0.72653 | 0.026118 | 1.5831 |
| DG 56:2 | 0.41473 | -1.2697 | 0.026161 | 1.5823 |
| PE 44:0 | 0.60464 | -0.72585 | 0.026176 | 1.5821 |
| TG O-45:2 | 0.32674 | -1.6138 | 0.026229 | 1.5812 |
| TG 98:18 | 2.1518 | 1.1056 | 0.026271 | 1.5805 |
| HexCer 49:3;O5 | 2.0724 | 1.0513 | 0.026277 | 1.5804 |
| PE 16:1_20:1 | 3.2862 | 1.7164 | 0.026284 | 1.5803 |
| HexCer 40:6;O4 | 2.4229 | 1.2767 | 0.026369 | 1.5789 |
| Hex2Cer 30:0;O3 | 1.6931 | 0.75968 | 0.026841 | 1.5712 |
| TG 15:0_16:1_18:1 | 2.1653 | 1.1146 | 0.027031 | 1.5681 |
| PE-Cer 32:1;O;P | 1.4846 | 0.57009 | 0.027271 | 1.5643 |
| DG 53:5 | 3.4328 | 1.7794 | 0.027442 | 1.5616 |
| Car 20:4 | 0.51822 | -0.94837 | 0.027468 | 1.5612 |
| MG O-32:1 | 0.54132 | -0.88545 | 0.027553 | 1.5598 |
| ST 20:5;O4;GlcA | 1.8439 | 0.88278 | 0.027626 | 1.5587 |
| PG O-56:0;O | 0.56564 | -0.82204 | 0.027663 | 1.5581 |
| Cer 18:0;O2/4:0 | 1.9447 | 0.95954 | 0.027681 | 1.5578 |
| PI O-38:2 | 0.24617 | -2.0223 | 0.02776 | 1.5566 |
| PG 48:2;O | 1.9356 | 0.95281 | 0.027784 | 1.5562 |
| PC O-66:0 | 8.1597 | 3.0285 | 0.027801 | 1.5559 |
| PG 56:4;O | 0.51205 | -0.96565 | 0.02782 | 1.5556 |
| PE 52:5 | 1.723 | 0.7849 | 0.028153 | 1.5505 |
| PC 26:0_18:1 | 2.6107 | 1.3844 | 0.028196 | 1.5498 |
| TG 16:1_18:0_20:2 | 5.7495 | 2.5234 | 0.028204 | 1.5497 |
| Car 32:0;O4 | 0.64383 | -0.63525 | 0.028287 | 1.5484 |
| FA 21:2;O | 1.7763 | 0.82886 | 0.028663 | 1.5427 |
| PA O-56:7 | 1.6408 | 0.71442 | 0.028774 | 1.541 |
| DG 30:2 | 2.5312 | 1.3398 | 0.029125 | 1.5357 |
| ST 20:3;O7;T | 1.7676 | 0.82182 | 0.029149 | 1.5354 |
| DG 18:0_28:0 | 3.042 | 1.605 | 0.029164 | 1.5352 |
| MG 34:0 | 2.6826 | 1.4236 | 0.029237 | 1.5341 |
| PR 53:11;O2 | 0.2705 | -1.8863 | 0.029343 | 1.5325 |
| PIP3 24:0 | 0.19244 | -2.3775 | 0.029379 | 1.532 |
| PG 16:1_17:1 | 0.52044 | -0.94221 | 0.029393 | 1.5318 |
| TG 13:0_14:0_16:0 | 1.4581 | 0.54412 | 0.029438 | 1.5311 |
| TG 82:4;O3 | 0.52771 | -0.92218 | 0.029444 | 1.531 |
| CPA 18:1 | 0.37872 | -1.4008 | 0.029598 | 1.5287 |
| CerP 52:0;O5 | 2.7122 | 1.4394 | 0.029625 | 1.5283 |
| DG O-18:1_22:6 | 2.6716 | 1.4177 | 0.029657 | 1.5279 |
| ST 24:2;O6 | 0.55181 | -0.85775 | 0.02968 | 1.5275 |
| PIP 30:7 | 0.27521 | -1.8614 | 0.030111 | 1.5213 |
| WE 41:1 | 1.8121 | 0.85767 | 0.030249 | 1.5193 |
| TG 57:10 | 2.3036 | 1.2039 | 0.030319 | 1.5183 |
| SL 25:0;O/26:2;O | 2.3797 | 1.2508 | 0.030477 | 1.516 |
| PI-Cer 38:0;O4 | 0.33975 | -1.5575 | 0.030478 | 1.516 |
| DG O-41:4 | 2.6658 | 1.4146 | 0.03073 | 1.5124 |
| FA 17:0 | 0.63453 | -0.65624 | 0.031107 | 1.5071 |
| DG 54:5 | 4.7613 | 2.2514 | 0.031374 | 1.5034 |
| PA 16:1_20:1 | 0.35184 | -1.507 | 0.031393 | 1.5032 |
| DG 60:10 | 9.8794 | 3.3044 | 0.03151 | 1.5016 |
| TG 62:2 | 3.1952 | 1.6759 | 0.031663 | 1.4994 |
| GlcADG 14:0_18:0 | 8.0855 | 3.0153 | 0.03201 | 1.4947 |
| Cer 44:6;O | 0.31549 | -1.6643 | 0.032056 | 1.4941 |
| SHexCer 42:1;O2 | 2.5687 | 1.361 | 0.032352 | 1.4901 |
| HexCer 45:2;O4 | 0.43162 | -1.2122 | 0.032375 | 1.4898 |
| FAHFA 33:1;O | 2.2111 | 1.1448 | 0.032384 | 1.4897 |
| DG 38:4 | 2.413 | 1.2708 | 0.032488 | 1.4883 |
| ST 27:3;F | 0.42783 | -1.2249 | 0.032573 | 1.4871 |
| DG 62:10 | 6.123 | 2.6142 | 0.03261 | 1.4866 |
| PG 42:2;O | 0.54494 | -0.87583 | 0.032815 | 1.4839 |
| TG O-57:3 | 1.8361 | 0.87666 | 0.033065 | 1.4806 |
| HexCer 44:2;O4 | 1.5902 | 0.66922 | 0.03312 | 1.4799 |
| SM 36:4;O4 | 3.328 | 1.7347 | 0.033167 | 1.4793 |
| SM 44:4;O4 | 2.5663 | 1.3597 | 0.03319 | 1.479 |
| LPC 22:5 | 0.21575 | -2.2126 | 0.03329 | 1.4777 |
| MGDG O-11:0_16:0 | 2.0924 | 1.0651 | 0.033416 | 1.476 |
| NAE 17:0 | 0.5133 | -0.96213 | 0.033662 | 1.4729 |
| PI-Cer 34:0;O4 | 0.69351 | -0.52801 | 0.033681 | 1.4726 |
| PG 54:4 | 0.55576 | -0.84745 | 0.033682 | 1.4726 |
| DG O-38:0 | 2.348 | 1.2314 | 0.033947 | 1.4692 |
| CerP 32:1;O2 | 2.2759 | 1.1864 | 0.033949 | 1.4692 |
| Cer 36:0;O2 | 1.9314 | 0.94963 | 0.03395 | 1.4692 |
| FA 24:2 | 0.46942 | -1.091 | 0.033984 | 1.4687 |
| Cer 46:1;O3 | 0.23108 | -2.1135 | 0.034006 | 1.4684 |
| Cer 43:1;O3 | 0.46294 | -1.1111 | 0.034041 | 1.468 |
| TG 16:1_18:1_18:2 | 2.6173 | 1.3881 | 0.034159 | 1.4665 |
| MG 31:1 | 0.55742 | -0.84317 | 0.0342 | 1.466 |
| ST 24:2;O2 | 0.36135 | -1.4685 | 0.034282 | 1.4649 |
| DG 40:6;O | 1.6318 | 0.70649 | 0.034361 | 1.4639 |
| TG 51:4;O2 | 1.8107 | 0.85654 | 0.034793 | 1.4585 |
| HexCer 18:0;O2/24:1 | 1.9681 | 0.97677 | 0.034933 | 1.4568 |
| TG 54:3 | 3.8188 | 1.9331 | 0.034956 | 1.4565 |
| TG 88:13;O2 | 0.26772 | -1.9012 | 0.035034 | 1.4555 |
| DG dO-40:2 | 2.259 | 1.1757 | 0.035455 | 1.4503 |
| TG 94:4;O2 | 2.4281 | 1.2799 | 0.035529 | 1.4494 |
| TG O-18:0_14:0_16:1 | 1.7919 | 0.84146 | 0.035605 | 1.4485 |
| PC 30:0;O | 2.114 | 1.08 | 0.035984 | 1.4439 |
| Hex2Cer 34:2;O2 | 0.34254 | -1.5457 | 0.036072 | 1.4428 |
| PG O-39:0 | 1.4581 | 0.54406 | 0.036195 | 1.4413 |
| HBMP 18:0_20:3_18:0 | 0.57399 | -0.80089 | 0.036484 | 1.4379 |
| TG 52:1;O | 2.5242 | 1.3358 | 0.036625 | 1.4362 |
| Hex(5)-HexNAc(3)-Cer 34:1;O2 | 1.6222 | 0.69795 | 0.036808 | 1.4341 |
| PG 34:2 | 0.56457 | -0.82477 | 0.036809 | 1.4341 |
| CerP 48:5;O3 | 4.521 | 2.1766 | 0.036824 | 1.4339 |
| HexCer 44:2;O5 | 0.55798 | -0.84172 | 0.036852 | 1.4335 |
| PC O-36:3 | 3.9103 | 1.9673 | 0.03686 | 1.4334 |
| NAT 19:2 | 0.58346 | -0.77729 | 0.036936 | 1.4326 |
| TG 58:7 | 3.3081 | 1.726 | 0.036955 | 1.4323 |
| TG O-55:2 | 2.1756 | 1.1214 | 0.037005 | 1.4317 |
| Car 34:0 | 0.30541 | -1.7112 | 0.03716 | 1.4299 |
| HexCer 38:2;O2 | 2.0844 | 1.0596 | 0.037196 | 1.4295 |
| PE-Cer 28:2;O2 | 2.4782 | 1.3093 | 0.037626 | 1.4245 |
| TG 15:0_21:2_22:6 | 4.0253 | 2.0091 | 0.037878 | 1.4216 |
| HexCer 15:3;O2/28:7 | 1.6414 | 0.71493 | 0.038234 | 1.4175 |
| HexCer 37:2;O6 | 0.36974 | -1.4354 | 0.03831 | 1.4167 |
| Cer 42:5;O | 0.5134 | -0.96184 | 0.038326 | 1.4165 |
| TG 16:1_18:1_19:1 | 5.0027 | 2.3227 | 0.038405 | 1.4156 |
| PG O-54:0;O | 0.57467 | -0.79919 | 0.03844 | 1.4152 |
| **TG 16:0_18:2_20:2** | 2.7694 | 1.4696 | 0.038971 | 1.4093 |
| **HexCer 38:3;O2** | 2.1819 | 1.1256 | 0.039024 | 1.4087 |
| **PC 15:0_18:3** | 1.8558 | 0.89203 | 0.039086 | 1.408 |
| **MG 34:2** | 0.54796 | -0.86785 | 0.039117 | 1.4076 |
| **DG 45:3** | 1.9378 | 0.95444 | 0.039302 | 1.4056 |
| **DG O-18:1_20:4** | 2.5483 | 1.3495 | 0.03971 | 1.4011 |
| **PI O-40:3** | 0.53613 | -0.89935 | 0.039821 | 1.3999 |
| **DG 52:5** | 3.8987 | 1.963 | 0.040016 | 1.3978 |
| **PE 30:1;O** | 2.0498 | 1.0355 | 0.040027 | 1.3976 |
| **TG 44:2** | 1.5824 | 0.66216 | 0.040058 | 1.3973 |
| **TG O-39:0** | 1.8791 | 0.91008 | 0.040192 | 1.3959 |
| **TG 55:8;O** | 2.2086 | 1.1431 | 0.040409 | 1.3935 |
| **Cer 18:0;O3/42:1;(2OH)** | 2.3511 | 1.2333 | 0.040501 | 1.3925 |
| **PI-Cer 35:0;O4** | 0.35675 | -1.487 | 0.040669 | 1.3907 |
| **DG 50:5** | 4.7915 | 2.2605 | 0.040669 | 1.3907 |
| **DG 59:6** | 8.439 | 3.0771 | 0.040769 | 1.3897 |
| **PG O-40:0;O** | 4.6074 | 2.204 | 0.040835 | 1.389 |
| **PC O-18:1_24:2** | 0.46368 | -1.1088 | 0.040866 | 1.3886 |
| **TG 85:1;O3** | 1.9331 | 0.95095 | 0.041055 | 1.3866 |
| **PI 37:2** | 0.61927 | -0.69136 | 0.041131 | 1.3858 |
| **CerP 51:3;O2** | 3.8092 | 1.9295 | 0.04116 | 1.3855 |
| **CerP 54:1;O5** | 2.9518 | 1.5616 | 0.041267 | 1.3844 |
| **DG 56:6** | 3.5258 | 1.8179 | 0.041338 | 1.3836 |
| **LPS O-26:0;O** | 0.64924 | -0.62319 | 0.041428 | 1.3827 |
| **FA 18:2** | 0.43208 | -1.2106 | 0.041783 | 1.379 |
| **CL 71:6** | 0.62998 | -0.66663 | 0.041788 | 1.379 |
| **PA O-45:2** | 5.1843 | 2.3741 | 0.042102 | 1.3757 |
| **Cer 35:3;O3** | 0.63558 | -0.65386 | 0.042118 | 1.3755 |
| **PIP 64:5;O** | 0.46337 | -1.1098 | 0.042359 | 1.3731 |
| **PE 22:5** | 11.949 | 3.5788 | 0.042413 | 1.3725 |
| **PE-Cer 32:3;O5** | 1.8197 | 0.86371 | 0.042827 | 1.3683 |
| **SM 41:1;O2** | 0.14142 | -2.822 | 0.042876 | 1.3678 |
| **Hex(4)-HexNAc-Fuc-Cer 42:1;O2** | 0.71175 | -0.49055 | 0.0433 | 1.3635 |
| **ACer 58:4;O2** | 0.52432 | -0.93148 | 0.043352 | 1.363 |
| **TG 60:3** | 4.9033 | 2.2938 | 0.043357 | 1.3629 |
| **NAT 32:0;O** | 1.7292 | 0.79013 | 0.043391 | 1.3626 |
| **HexCer 43:4;O5** | 1.9303 | 0.94882 | 0.043454 | 1.362 |
| **PC O-52:9** | 0.18136 | -2.4631 | 0.043459 | 1.3619 |
| **PE 41:1;O** | 0.45702 | -1.1297 | 0.043587 | 1.3606 |
| **HexCer 39:3;O6** | 1.5275 | 0.61114 | 0.043749 | 1.359 |
| **SHexCer 43:4;O3** | 0.46923 | -1.0916 | 0.043908 | 1.3575 |
| **Cer 18:2;O2/26:2;O** | 0.4328 | -1.2082 | 0.043929 | 1.3572 |
| **NA 22:0** | 1.6787 | 0.74731 | 0.043935 | 1.3572 |
| **PA 30:0** | 2.4319 | 1.2821 | 0.044118 | 1.3554 |
| **FA 18:1** | 0.57146 | -0.80728 | 0.04521 | 1.3448 |
| **DG 36:3** | 3.6703 | 1.8759 | 0.045416 | 1.3428 |
| **PA 45:4** | 0.44053 | -1.1827 | 0.045439 | 1.3426 |
| **DG 58:6** | 5.6583 | 2.5004 | 0.045734 | 1.3398 |
| **DG O-32:0** | 0.59991 | -0.73719 | 0.045778 | 1.3393 |
| **PI 54:4;O** | 0.40888 | -1.2903 | 0.045981 | 1.3374 |
| **CE 26:3;O** | 0.17506 | -2.5141 | 0.046133 | 1.336 |
| **PG 56:3;O** | 0.59357 | -0.75251 | 0.046171 | 1.3356 |
| **DG 61:8** | 3.4718 | 1.7957 | 0.046194 | 1.3354 |
| **TG 57:9;O3** | 0.36809 | -1.4419 | 0.046301 | 1.3344 |
| **DG O-56:8** | 2.2384 | 1.1625 | 0.04634 | 1.334 |
| **PG 14:1_18:1** | 0.48644 | -1.0397 | 0.046442 | 1.3331 |
| **TG 97:7;O3** | 0.27572 | -1.8587 | 0.046712 | 1.3306 |
| **Cer 49:6;O4** | 1.6523 | 0.72448 | 0.046741 | 1.3303 |
| **TG 58:4;O2** | 1.81 | 0.85596 | 0.046886 | 1.329 |
| **DG 39:0;O2** | 2.1263 | 1.0883 | 0.047086 | 1.3271 |
| **TG 54:2;O2** | 10.31 | 3.366 | 0.047087 | 1.3271 |
| **Cer 40:0;O3** | 0.41294 | -1.276 | 0.047152 | 1.3265 |
| **PE-Cer 54:0;O5** | 1.5031 | 0.58796 | 0.047186 | 1.3262 |
| **TG 82:1;O** | 0.43532 | -1.1999 | 0.047337 | 1.3248 |
| **CE 27:7** | 0.28076 | -1.8326 | 0.047352 | 1.3247 |
| **SHexCer 41:3;O3** | 2.4872 | 1.3145 | 0.047438 | 1.3239 |
| **PI 18:0_22:2** | 0.67035 | -0.577 | 0.047649 | 1.3219 |
| **MG 18:2;O** | 0.15425 | -2.6967 | 0.047804 | 1.3205 |
| **MIPC 39:6;O3** | 0.53973 | -0.88969 | 0.047831 | 1.3203 |
| **DG 44:3** | 1.7757 | 0.8284 | 0.047864 | 1.32 |
| **PE O-20:3_27:0** | 0.35952 | -1.4759 | 0.048143 | 1.3175 |
| **PC 28:0** | 0.35596 | -1.4902 | 0.048334 | 1.3158 |
| **TG 51:3;O2** | 1.9574 | 0.96896 | 0.048355 | 1.3156 |
| **TG 15:1_18:1_18:1** | 0.26636 | -1.9086 | 0.048394 | 1.3152 |
| **DG 36:1** | 2.0571 | 1.0406 | 0.04843 | 1.3149 |
| **PE 20:1_34:2** | 1.4075 | 0.49316 | 0.048457 | 1.3146 |
| **DG 49:3** | 1.6152 | 0.69169 | 0.048475 | 1.3145 |
| **DG 33:3** | 2.33 | 1.2203 | 0.048621 | 1.3132 |
| **PE O-18:2_22:3** | 2.0581 | 1.0413 | 0.048774 | 1.3118 |
| **DG O-48:3** | 3.4256 | 1.7764 | 0.048865 | 1.311 |
| **HexCer 39:1;O4** | 1.9759 | 0.98249 | 0.04908 | 1.3091 |
| GM3(d18:1/16:0) | 2.022 | 1.0158 | 0.049145 | 1.3085 |
| Hex2Cer 36:2;O2 | 0.51147 | -0.96727 | 0.049212 | 1.3079 |
| CL 14:0_18:2_14:0_18:2 | 0.54204 | -0.88354 | 0.049356 | 1.3067 |
| DG 58:1 | 2.9511 | 1.5613 | 0.049494 | 1.3054 |
| DG 16:0_20:3 | 2.2698 | 1.1826 | 0.049503 | 1.3054 |
| Cer 18:2;O2/26:7;O | 1.5639 | 0.64517 | 0.049662 | 1.304 |
| Car 20:4;O2 | 1.6006 | 0.67863 | 0.049788 | 1.3029 |

**Supplemental Table 8.**
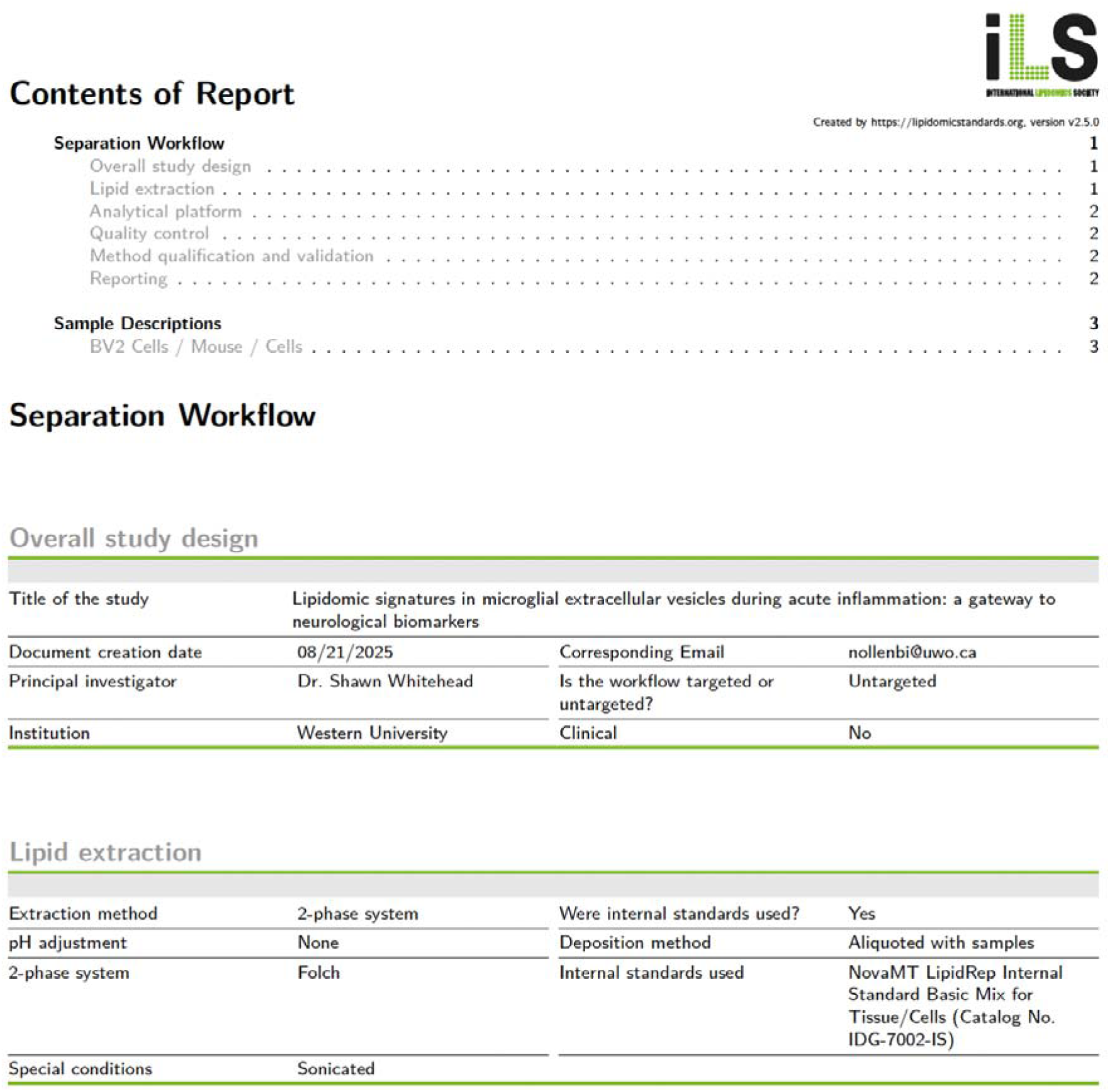
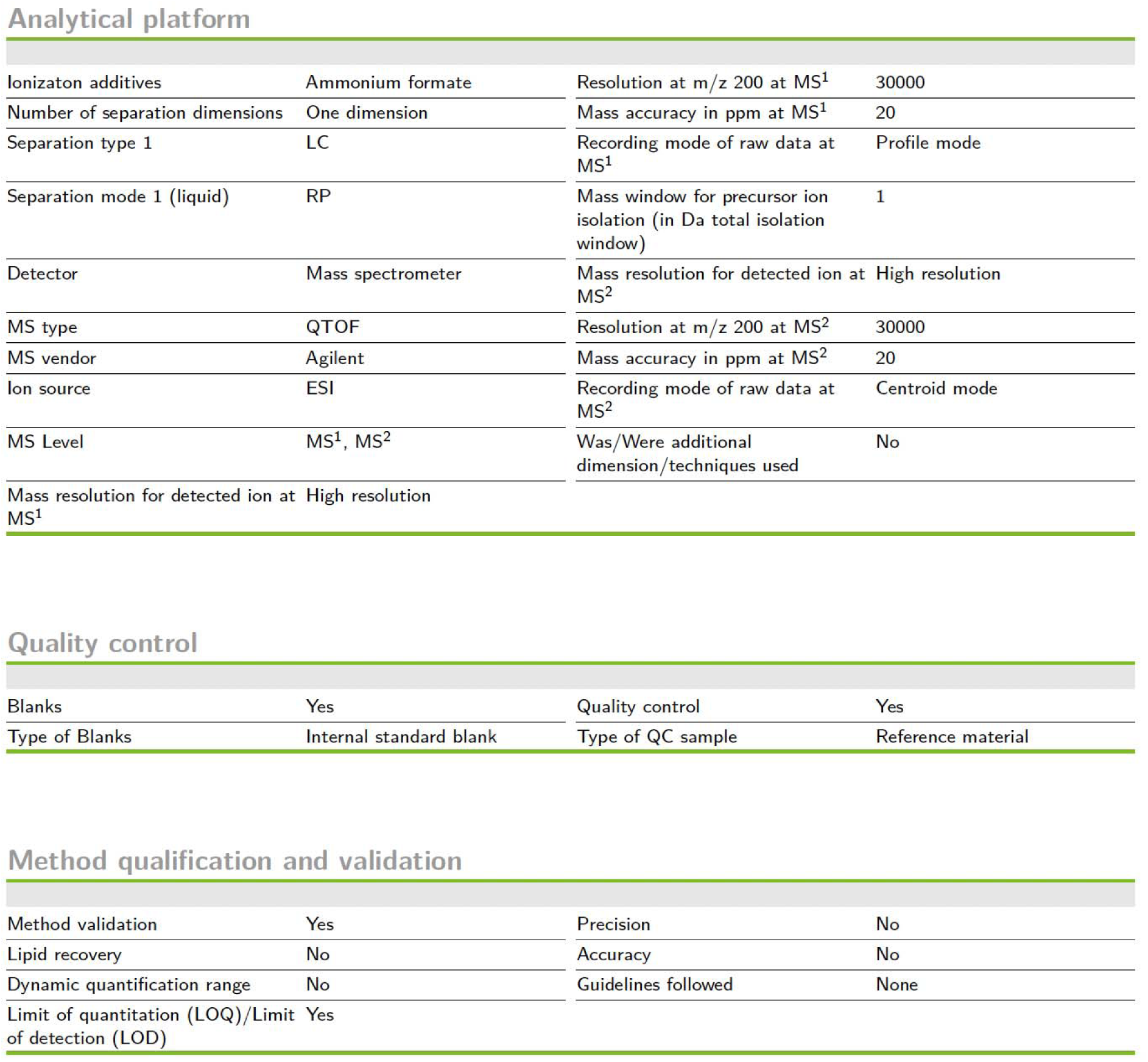
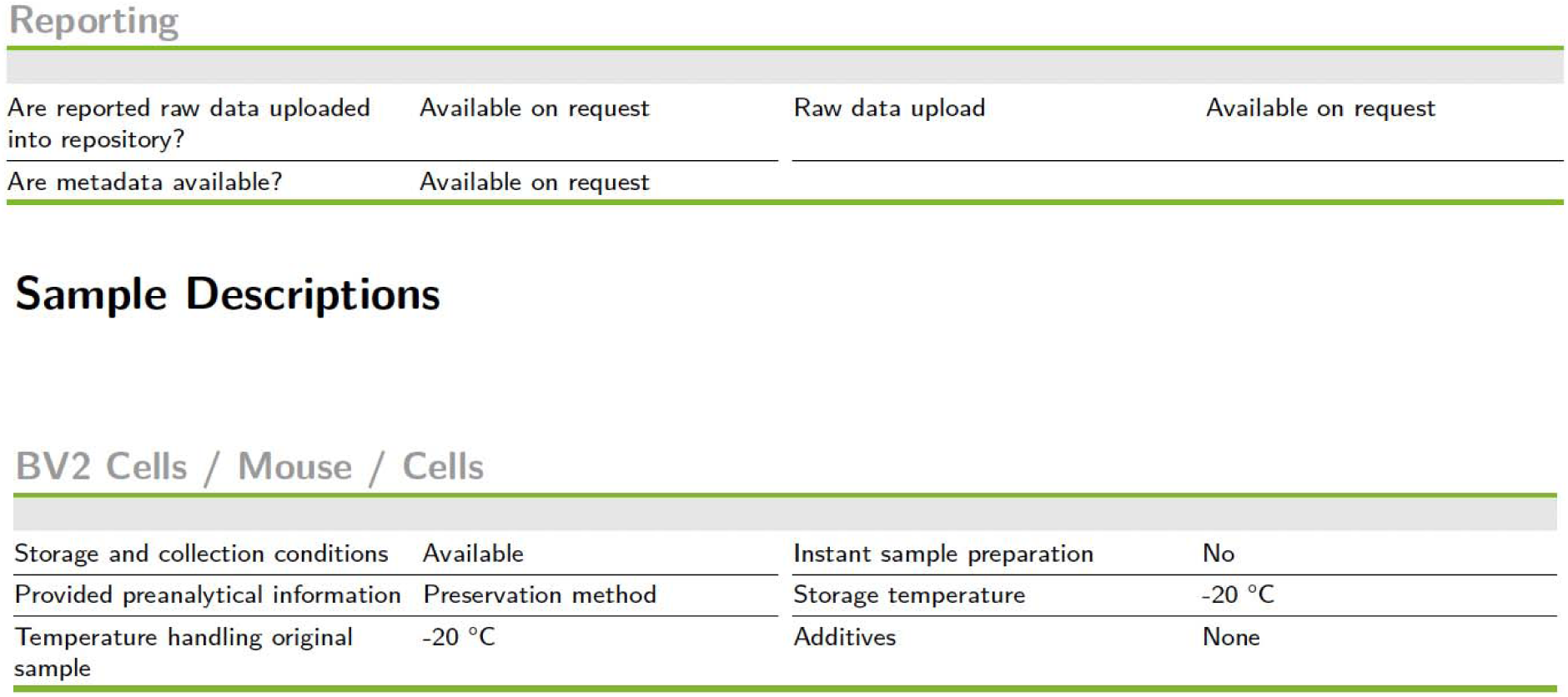
Lipidomics minimal reporting checklist. To enhance reproducibility a Lipidomics Minimal Reporting Checklist (Kopczynski et al. 2024) from the Lipidomics Standards Initiative has been included.

## Notes

### Competing Interest Statement

The authors have declared no competing interest.

### Summary of Updates

We have revised the manuscript to clarify language in figure captions and throughout the manuscript body to improve the flow of the text and highlight the main take-aways of each figure. Additionally, based on reviewer feedback we have opted to remove statistical tests that are merely a duplicate means of demonstrating the main findings. Furthermore, we have added additional supplemental figures that more thoroughly characterize the isolated extracellular vesicles (EVs) used in this work, standardize the reporting of the lipidomic methodology employed, and provide additional details on statistical texts.

